# The genetic legacy of the expansion of Bantu-speaking peoples in Africa

**DOI:** 10.1101/2023.04.03.535432

**Authors:** Cesar A. Fortes-Lima, Concetta Burgarella, Rickard Hammarén, Anders Eriksson, Mário Vicente, Cecile Jolly, Armando Semo, Hilde Gunnink, Sara Pacchiarotti, Leon Mundeke, Igor Matonda, Joseph Koni Muluwa, Peter Coutros, Terry S. Nyambe, Cirhuza Cikomola, Vinet Coetzee, Minique de Castro, Peter Ebbesen, Joris Delanghe, Mark Stoneking, Lawrence Barham, Marlize Lombard, Anja Meyer, Maryna Steyn, Helena Malmström, Jorge Rocha, Himla Soodyall, Brigitte Pakendorf, Koen Bostoen, Carina M. Schlebusch

**Author notes:** These authors contributed equally to this work: Cesar A. Fortes-Lima, Concetta Burgarella, Rickard Hammarén. (C.M.S.).

## Abstract

With the largest genomic dataset to date of Bantu-speaking populations, including newly generated data of modern-day and ancient DNA from previously unsampled regions in Africa, we shed fresh light on the expansion of peoples speaking Bantu languages that started ∼4000 years ago in western Africa. We have genotyped 1,740 participants, including 1,487 Bantu speakers from 143 populations across 14 African countries, and generated whole-genome sequences from 12 Late Iron Age individuals. Our results show that Bantu speakers received significant gene-flow from local groups in regions they expanded into. We show for the first time that genetic diversity amongst Bantu-speaking populations declines with distance from western Africa, with current-day Zambia and the DRC as possible crossroads of interaction. Using spatially explicit methods and correlating genetic, linguistic and geographical data, we provide cross-disciplinary support for a serial founder migration model. Finally, we discuss the utility of our dataset as an exhaustive modern-day African comparative dataset for ancient DNA studies. These new findings and data will be important to a wide range of disciplines from science and humanities as well as to the medical sector studying human genetic variation and health in African and African-descendant populations.

**One-sentence summary:** A comprehensive genetic analysis of the expansion of people speaking Bantu languages reveals a complex history of serial founder events, variable levels of contact with local groups, and spread-over-spread events.

## Main

African populations speaking Bantu languages (henceforth: BSP standing for “Bantu-speaking populations”) constitute almost a fourth of Africa’s total population today, where about 350 million people across nine million km^2^ speak more than 500 Bantu languages ^1,2^. Archeological, linguistic, historical, and anthropological sources attest to the complex history of the expansion of BSP across sub-equatorial Africa that fundamentally reshaped the linguistic, cultural and biological landscape of the continent with a lasting impact until today. The initial spread of Bantu languages alongside people/genes was a demic expansion across central, eastern and southern Africa (**Note.S 1**) ^3–11^. There is a broad consensus across the academic disciplines that ancestral BSP migrated first through the Congo rainforest, and only later from there to the savannas further east and south ^4–6,9,11–19^. However, debates across a wide range of disciplines persist on the precise pathways and modes of the expansion of BSP (**Fig.S 1.1, Note.S 1**) (see, for example, ^14,15^).

Whereas most major human expansions involved latitudinal movements and hence went through regions with similar climatic conditions ^20,21^, the expansion of BSP is notable for largely being longitudinal with movements through regions with very different climatic conditions, encompassing a wide range of biomes. For example, the putative BSP homeland in the highlands of Nigeria and Cameroon differs considerably from the central African rainforest, the African savannas, and the dry conditions of southwestern Africa. Yet, BSP migrated to and settled in all these different habitats and climatic conditions. Despite agreement on its demic nature, previous genetic studies on the expansion of BSP did not find the typical serial founder effect expected when small migrant groups settle in new areas, leading to a decrease in genetic diversity with increasing distance from the putative homeland ^22,23^. The signal of founder events could have been erased by a subsequent increase of genetic variation through admixture with local populations and/or via long-distance contacts involving later migrations of Bantu speaking communities, so-called spread-over-spread events ^4,16,24,25^. This underlying complexity, coupled with the different migration routes and patterns proposed by linguistics, archeology and genetics, makes the expansion of BSP interesting for exploration with newer population-genetic methods and modeling approaches that are spatio-temporally sensitive.

Although whole-genome studies of African populations have recently became available ^26–29^ and several locally focused genome-wide genotype studies exist ^9–11^, genomic data from BSP across the entirety of sub-Saharan Africa are still sparse. A better representation of populations across the landscape is needed to make in-depth inferences about the routes and modalities of their large spread. To investigate the demographic history, migration patterns, and admixture dynamics of BSP in detail, we have generated a novel genome-wide genotype dataset of 1,740 individuals (**Table.S 1**), encompassing 1,487 Bantu-speaking individuals and 253 other African individuals, from 143 populations spanning 14 sub-Saharan African countries (**Fig. 1a, Fig.S 1.2**). The genotyped dataset includes 117 populations that to our knowledge were not included in previous genomic studies. Our dataset encompasses populations speaking languages belonging to most major branches of the Bantu language family ^14^: north-western (2 NW-BSP), west-western (7 WW-BSP), south-western (13 SW-BSP plus the Damara, a Khoe-Kwadi speaking population from Namibia with a genetic profile shared with BSP), and eastern (44 E-BSP) group. In addition, we have generated genomic data for 12 ancient individuals from Late Iron Age sites of south-central and southern Africa (present-day Zambia and South Africa), spanning 97-688 years before present (BP). This dataset provides a comprehensive representation of sub-Saharan Africa’s genetic landscape. We have used allele-frequency and haplotype-based methods, genetic diversity summary statistics, and spatial modeling to gain a better understanding of the demographic history of BSP and to shed new light on complex patterns of migration, admixture and human history in Africa.

**Fig. 1.**
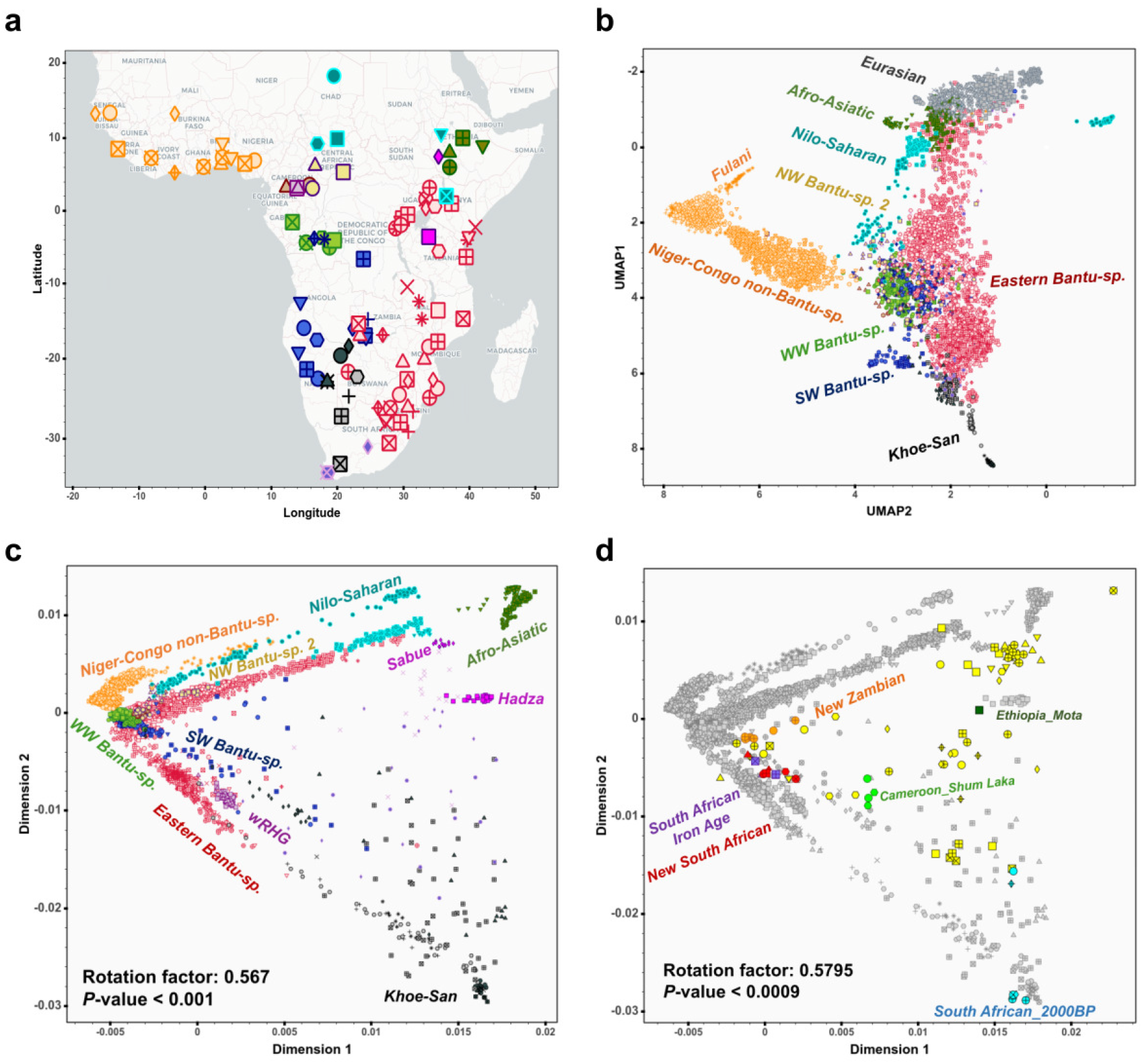
Population structure within sub-Saharan African populations. **a**, Geographical locations of the 111 sub-Saharan African populations selected for population genetics analysis within the AfricanNeo dataset. Populations with the same colour belong to the same group based on linguistic and geographic characterization (**Table.S 2**). Legend is described in **Fig.S1.5a**. Further details about the populations are presented in **TableS 2. b**, Uniform manifold approximation and projection (UMAP) analysis of sub-Saharan African populations **c**, Principal Component Analysis (PCA) of sub-Saharan African populations. PCA axes were Procrustes rotated according to geography. Further PC projections were included in **Fig.S 2.4. d**, PCA for projected ancient DNA individuals (with colours) and present-day sub-Saharan African populations (in gray, same as (**c**)) procrustes rotated according to geography.

## Results and Discussion

### Genetic admixture with local populations influenced population structure in BSP

After quality control and merging the generated data of present-day sub-Saharan African populations with publicly available data from a broad range of populations representative of different ethnolinguistic groups (**Suppl. Material**), we have assembled a dataset for 4,950 individuals from 124 populations (111 sub-Saharan African and 13 Eurasian populations with at least 10 individuals per population), hereafter referred to as the “AfricanNeo” dataset (**Fig.S 1.3–1.6, Table.S 2**). Using four dimensionality reduction methods (**Suppl. Material**), we have found evidence for fine-scale population structure between sub-Saharan African populations with a clear geographical component (Procrustes correlation to geography > 0.57, *P*-value < 10-3), and a broad correspondence with the main linguistic groups in Africa (**Fig. 1b–1c, Fig.S 2.1–2.8**; **Note.S 2**). Groups of BSP (NW-BSP, WW-BSP, SW-BSP, and E-BSP) can be distinguished, showing population substructure.

Population substructure and suggestions of admixture patterns are also apparent in model-based clustering analyses (**Fig. 2a, Fig.S 3.1–3.14**; **Note.S 2**; **Suppl. Material**), and reveal a finer picture of population ancestries with three main BSP-associated genetic components (**Fig.S 3.1–3.2, Table.S 3**): the dark-green component found in most BSP, the cyan component shared between non-Bantu Niger-Congo and western BSP (NW-BSP and WW-BSP), and the orange component mainly found in southeastern BSP.

**Fig. 2.**
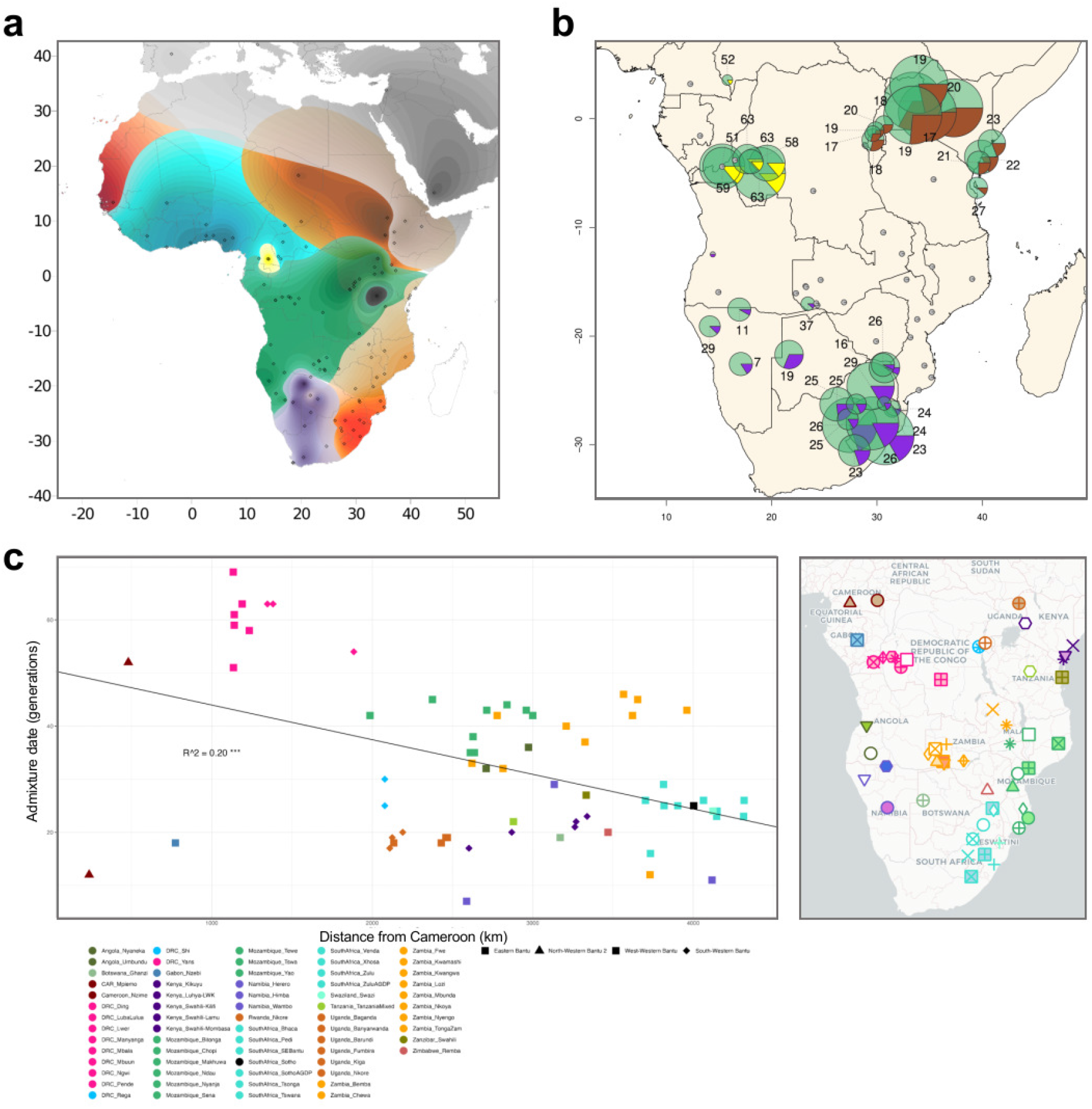
Population structure, admixture dates and fractions. **a**, Contour map of overlapping unsupervised ADMIXTURE results at K=12 created using the Kriging method for all the populations included in the AfricanNeo dataset. Ancestry components with values under 25% are not represented on the map. All the values estimated for each ancestry are shown in **Fig.S 3.3. b**, Inferred admixture dates (number of generations ago) and fractions (pie chart) for each BSP estimated using MOSAIC analyses. Each inferred population source is highlighted with a different color: west-central African-related ancestry in green; western rainforest hunter-gatherer ancestry in yellow; Afro-Asiatic-speaking ancestry in brown; and Khoe-San ancestry in purple. The size of the charts is in relation to the sample size of each population. Further details are included in **Fig.S 11.1. c**, Admixture dates of studied BSP versus geographical distances from Cameroon (colours according to country).

BSP also show differential genetic affinities with other sub-Saharan African populations (**Fig. 1b–1c, Fig.S 2.1–2.8**; **Table.S 4**). This pattern may be the result of genetic admixture with local hunter-gatherer groups and Nilo-Saharan and Afro-Asiatic speaking populations, during and/or after the expansion of BSP across sub-equatorial Africa ^5,8,30–32^. We have formally tested the hypothesis of admixture and its regional character using f3- and f4-statistics (**Suppl. Material**). The results confirm a significant and differential contribution of Afro-Asiatic ancestry in eastern BSP from Kenya and Uganda (**Fig.S 3.16**), of western rainforest hunter-gatherers (wRHG) ancestry in western BSP from the Democratic Republic of Congo (DRC) and the Central African Republic (CAR) (**Fig.S 3.17 and 3.19**), and of Khoe-San ancestry in southern BSP from South Africa, Botswana, Zambia (Fwe population), and Namibia (**Fig.S 3.18**; **Note.S 2**).

### Population structure in Bantu-speaking populations remains after removing admixture

To assess if admixture with local groups is the main process driving spatial patterns of structure in BSP (**Fig. 1c, 2a**), we have masked out admixed genomic regions in BSP ^33^ and kept only west-central African genomic components (**Suppl. Material**). This masked dataset has allowed us to minimize the influence of non-Bantu-speaker ancestries in subsequent analyses, and the comparison with our results from the original unmasked dataset allowed us to evaluate the impact of admixture on the genetic diversity and population structure of BSP. Principal component analysis (PCA) on the admixture-masked dataset (**Fig. 3, Fig.S 4.4**; **Note.S 3**) show that BSP retain a clear genetic structure that aligns with geographic (**Fig. 3b**,**d**) (Procrustes correlation > 0.649, *P*-value < 0.001; **Fig. 3c–3d**) and linguistic (**Fig. 3a**,**c**) structure, suggesting that processes other than genetic admixture influence spatial patterns of BSP diversity. However, this structure could be driven also by outlier BSP with increased genetic drift. Notably, Herero and Himba from Namibia largely influence PC2 of the admixture-masked dataset (**Fig. 3a–3b, Fig.S 4.4**). We have thus repeated the PCA on the admixture-masked dataset after excluding Himba and Herero from the analysis, and still observed population structure in the remaining BSP (**Fig.S 4.5**).

**Fig. 3.**
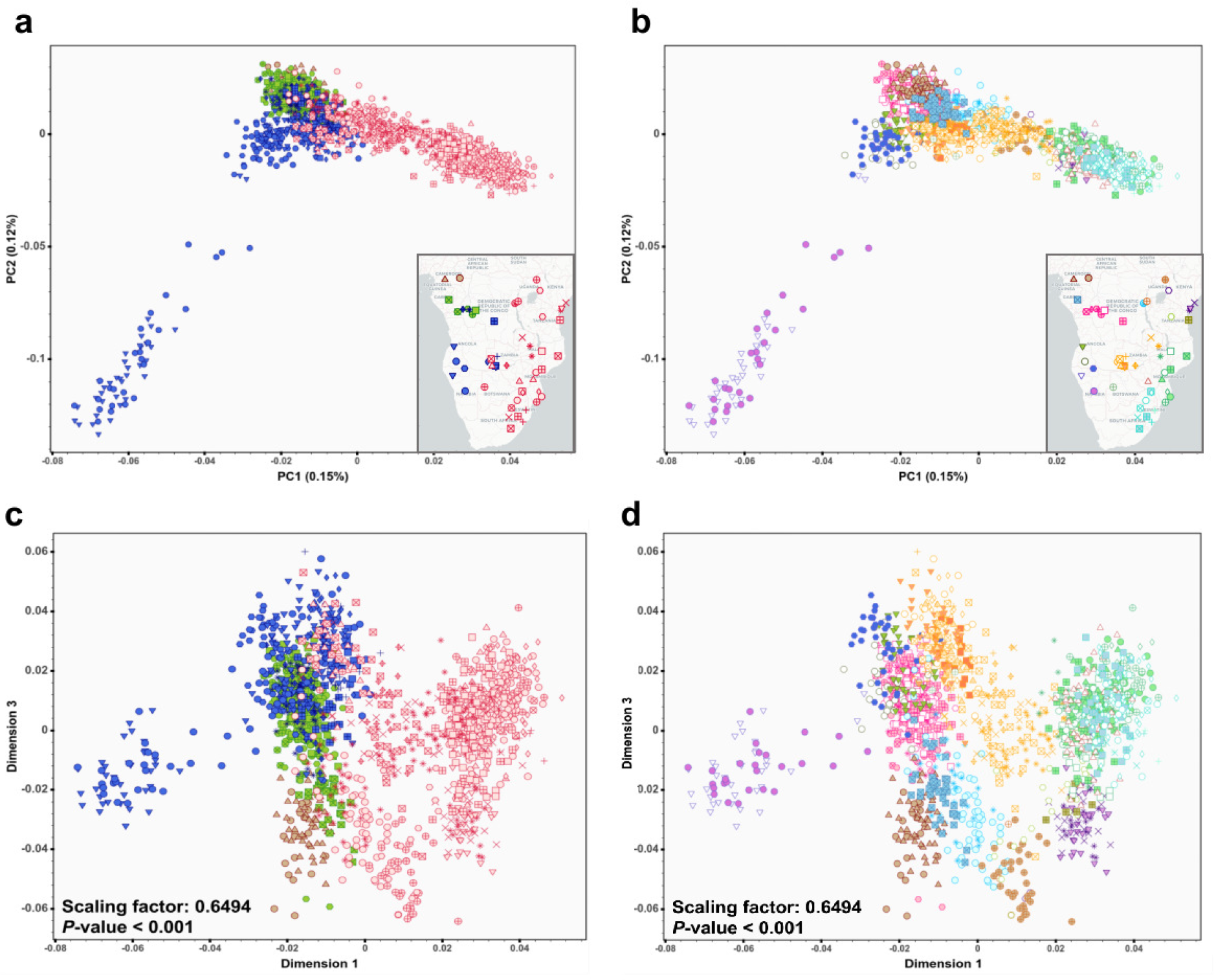
Population structure patterns in BSP after admixture masking. **a–d** PCA plot on the admixture-masked BSP dataset. Figure showing BSP coloured by linguistic groups in PC1 vs PC2 (**a**) and PC1 vs PC3 (**c**); and by geography represented by countries in PC1 vs PC2 (**b**) and PC1 vs PC3 (**d**) (also in **Fig.S 4.4**). Procrustes rotation was used for (**c**) and (**d**), and the estimated scaling factor was 0.649 (*P*-value< 0.001).

We have then investigated whether BSP harbor signatures of genetic isolation and population size changes potentially driving the observed structure. Most BSP show similar patterns of genetic drift reflected in runs of homozygosity (ROH) (**Fig.S 5.1–5.12, Table.S 6**; **Note.S 4**; **Suppl. Materials**), and changes in the effective population sizes (N_e_) that support population expansion signatures in the last 10–20 generations (**Fig.S 6.1–6.2**; **Note.S 5**; **Suppl. Materials**). The Himba and Herero populations notably deviate from the general patterns of BSP, showing higher intensities of founder events (*I*_*f*_=1.6% and 1.2%, respectively; **Fig.S 6.3, Table.S 7**), higher values of the genomic inbreeding coefficient (F_ROH_= 0.021 ±0.012SD and 0.015 ±0.007SD, respectively), and higher long ROH length averages (for ROH length categories 4 and 5; **Fig.S 5.6, 5.11–5.12, Table.S 6**) than the other studied BSP (**Note.S 4, 5**). These signatures can be the consequence of genetic isolation since their arrival in southwestern Africa and recent endogamic practices linked to cattle herding ^34^, as suggested in studies of mitochondrial DNA data ^35^, genome-wide genotype data ^36^, and exome sequencing data ^34^.

### Models underlying the structure of Bantu-speaking populations

A strong correlation of genetic relatedness among groups with geography could be explained by an isolation-by-distance (IBD) model, which assumes stepwise gene-flow between neighboring groups. Our dataset of BSP fits an IBD pattern (**Fig.S 7.1–7.4**; **Suppl. Materials**), including when admixture was removed (**Fig.S 7.5–7.8**; **Note.S 6**), consistent with previous findings based on fewer populations and a smaller dataset ^4^. However, alternative underlying models could explain IBD patterns. For instance, under a serial-founder model we also expect a strong correlation between shared genetic ancestry and geography. However, in contrast to IBD models, a serial-founder model would also show a decrease in genetic diversity from the putative region of origin. To distinguish between these two models, we have investigated the spatial distribution of three genetic diversity summary statistics (**Suppl. Materials**) suitable for array-based genotype data ^7^. All three statistics (haplotype richness, haplotype heterozygosity, and linkage disequilibrium) support a serial-founder model in which the highest genetic diversity is found in western BSP with a steady decline with distance towards eastern and southern BSP (**Fig.S 8.1–8.4**; **Note.S 7**). This pattern is stronger in the admixture-masked dataset (**Fig. 4, Fig.S 8.5–8.14**). Further evidence supports serial-founder dynamics during the BSP expansion from west-central Africa, e.g., significant demographic founder events have been inferred in 19 BSP (**Table.S 7**; **Note. S 5**), and a maximum-likelihood (ML) tree of the admixture-masked BSP dataset shows north-western BSP at the base of the ML-tree and eastern BSP forming a monophyletic group (**Fig.S 9.1**; **Note.S 8**; **Suppl. Materials**). In contrast, admixture largely drove the shape of the ML-trees for the unmasked datasets (**Fig.S 9.2–9.3**).

**Fig. 4.**
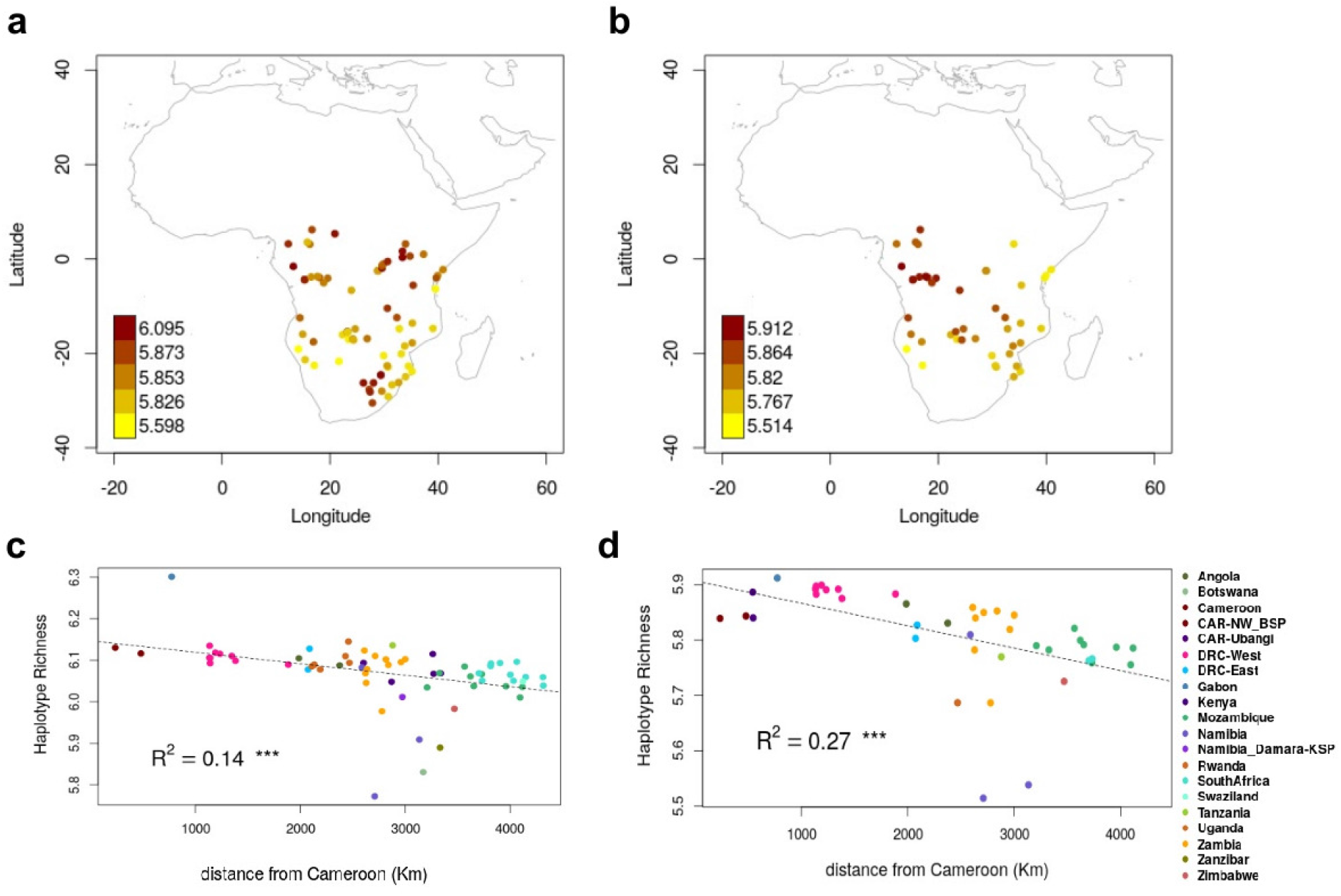
Patterns of genetic diversity in BSP. **a**, Map of haplotype richness (HR) estimated for the unmasked dataset of African populations (N= 67 populations; with a minimum sample size of 10 individuals and a maximum size of 30 individuals). **b**, HR estimated for the admixture-masked BSP dataset (N= 49 populations) that includes BSP with at least 70% of West-Central African-related (WCA) ancestry. **c**, Decrease of HR estimates with geographical distance from Cameroon to the sampling location of each BSP included in the unmasked BSP dataset of (a), and **d**, the admixture-masked BSP dataset of (b). The dotted line represents the linear relationship between HR estimates and geographical distances.

Overall, these analyses support the suggestion that the expansion of BSP started from west-central Africa and proceeded mostly via serial bottlenecks to spread throughout sub-equatorial Africa. The fact that a negative correlation between genetic diversity and distance from the source is present even in the unmasked dataset suggests that admixture had a small impact on levels of genetic diversity of BSP, either because there was not that much gene-flow with indigenous groups, or some BSP moved on before they received substantial local gene-flow. The fact that the admixture patterns are largely region-specific (**Fig. 2a**), i.e., in each population we detect non-BSP ancestry from local groups and not from elsewhere, suggests the latter.

### Routes and timing of expansions of Bantu-speaking populations

To understand how the expansion of BSP unfolded, we have investigated the spatial routes and timing of their movements. First, we have used a climate-informed spatially explicit model ^37,38^ to infer the most likely initial expansion routes of BSP with specific models of population expansion that correspond to the “Late-split” and “Early-split” hypotheses proposed by linguistic studies (**Fig.S 1.1, 10.1**). We have run one million Wright-Fisher simulations (**Suppl. Material**) to test three expansion scenarios that differ in whether BSP were allowed to spread south through the Congo rainforest (i.e. “southern route” or “Late-split”; **Fig.S 1.1b**), north of the rainforest (“northern route” or “Early-split”; **Fig.S 1.1a**), or both routes. The scenario with only the northern route receives substantially less statistical support from the data compared to scenarios in which either both routes or only the southern route were considered (R^2^= 0.19, 0.32, and 0.34, respectively). These results support the Late-split hypothesis, in agreement with the most recent linguistic, archeological and genetic evidence ^9,11,14–16,39^, and highlight the importance of the Congo rainforest in the initial expansion of BSP and in shaping present-day genetic variation in sub-equatorial Africa.

Previous studies proposed possible gene-flow between western and eastern branches of Bantu speakers ^4,6^. Even though populations speaking western and eastern Bantu languages are more separated in the PCA towards the terminal parts of the distribution, there is overlap toward the middle, in particular in BSP from current-day Zambia and the DRC (**Fig. 3**). These two countries thus represent interaction zones between different linguistic subgroupings, which is also reflected in genetic substructure that differs between eastern and western geographic regions (**Fig.S 3.14–3.15**). Our *F*_*ST*_-based model to infer expansion routes of BSP by tracing nearest *F*_*ST*_ values over the geographic landscape also indicates Zambia as a possible interaction nexus (**Fig. 5a, Fig.S 10.2–10.3**; **Note.S 9**; **Suppl. Material**). Specifically, the Lozi population from Zambia represents the proxy population of the Bantu-speaking migrants from the western DRC to Zimbabwe, Mozambique, Eswatini (former Swaziland), and South Africa (**Fig.S 10.3a, 10.3c**). However, the Lozi language, which is widely used as a *lingua franca* in Zambia’s Western Province and adjacent areas, was only introduced into the region in the 19th century CE by Sotho-speaking immigrants from what is today South Africa ^40,41^. Removing the Lozi population from the analysis moves the connection point between eastern and southeastern BSP with western BSP to what is currently the western region of the DRC (**Fig. 5b, Fig.S 10.2b**).

**Fig. 5.**
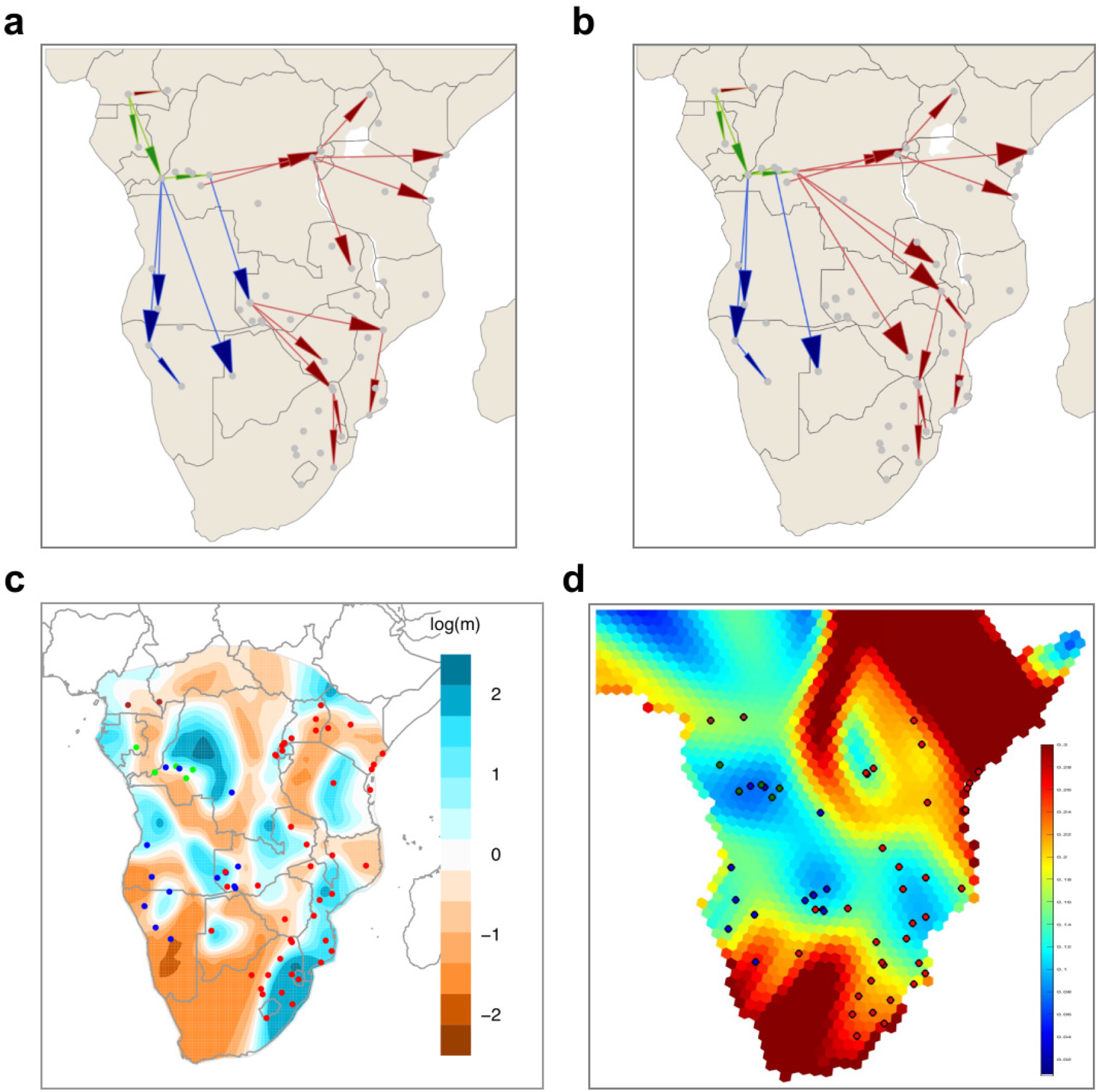
Migration routes and rates in BSP. **a**, Putative migration routes of BSP inferred using pairwise *F*_*ST*_ values (also in **Fig.S 11.3**), and **b**, after removing the Zambian Lozi population from the analyses (also in **Fig.S 11.3**). Arrow colors correspond to north-western Bantu 2 (in brown), west-western Bantu (in green), south-western Bantu (in dark blue), and eastern Bantu speakers (in red). **c**, Spatial visualization of effective migration rates (EEMS) estimated with the masked BSP dataset (further details in **Fig.S 10.6**). Populations are coloured according to Bantu-speaking linguistic groups (north-western 2 in brown, west-western in green, south-western in dark blue, and eastern in red). **d**, GenGrad analysis using *F*_*ST*_ as the genetic distance for the admixture-masked BSP dataset. Hexagons of the grid were plotted with a color scale representing the *F*_*ST*_ gradient.

Current-day Zambia as a potential point of divergence between expansion routes of BSPwas previously proposed by Choudhury et al. ^26^ using a small number of BSP not representative of the whole Bantu-speaking geographic distribution. Here, with notably better representation across the geographic region of BSP (**Fig.S 1.2, 1.5**), we have identified Zambia but also the western DRC as important nexus zones. However, spatially explicit analyses using EEMS, FEEMS, and *F*_*ST*_ analyses (**Fig. 5c–5d, Fig.S 10.4–10.11**; **Note.S 10**) (**Suppl. Material**) and clustering methods (**Fig.S 3.14–3.15**) suggest barriers to gene-flow and population structure even in this interaction zone, possibly caused by the linguistic division between BSP. Future research and model-testing methods could further establish whether these are interaction zones between populations speaking the eastern and western branches of Bantu languages or splitting points in past expansion routes. Spatial methods further indicate high effective migration rates along the Indian Ocean coast from Kenya to eastern South Africa (blue areas in **Fig. 5c, Fig.S 10.6**), as was reported previously ^11^, together with longitudinal corridors of lower migration rates in the central parts of the continent (brown areas in **Fig. 5c–d**).

Dating of admixture events in BSP^42^ strongly supports the main direction of their expansion across sub-equatorial Africa (**Fig. 2b, Fig.S 11.1, Table.S 8**; **Note.S 11**). Admixture dates significantly correlate with geographic distance from the homeland region of BSP (R^2^=0.20, *P*-value= 2.6e-05; **Fig. 2c**; **Suppl. Material**), with earlier dates of admixture in west-central Africa between BSP and wRHG, and more recent dates toward the extremes of the expansion (**Fig. 2c, Fig.S 11.1–11.2**), where admixture with Khoe-San groups ranges 7–29 generations ago in southwestern BSP from Namibia and 16–29 generations ago in southeastern BSP from South Africa (**Fig.S 11.1b**). These results also suggest that the rate of movement of BSP was more or less constant through time, despite the wide variety of environments and population interactions. Interestingly, admixture seems to be older than expected in BSP from western regions (e.g., admixture between BSP and wRHG in western DRC) and younger in certain eastern regions (e.g., admixture between BSP and different eastern African groups in Uganda and Kenya) (**Fig. 2b–2c, Fig.S 11.1**), suggesting that the rates of movement of BSP into those regions were either faster or slower than the average speed, or the admixture occurred earlier or later after arrival than in the other regions.

### Spread-over-spread events versus genetic continuity

It has been suggested that the initial expansion of BSP across sub-equatorial Africa was followed by subsequent migrations of BSP following similar routes, creating a pattern of spread-over-spread events ^43^, which in some cases may have replaced earlier settlers and their languages ^16,25,44^. Due to shifts to newcomers’ languages and the subsequent or independent death of first settlers’ languages, certain branches of the Bantu language family tree possibly no longer represent the initial expansion of BSP ^45^. In case of contact and admixture between incoming and earlier settled BSP, genetic data may represent an amalgamation of spread events, whereas linguistic data may represent the last spread event only. If so, both linguistics and genetics will correlate with geography, but not necessarily with each other.

We have tested this hypothesis using Mantel tests (**Table.S 9**; **Suppl. Material**). Pairwise-population linguistic and geographic distances are significantly correlated (r-statistics= 0.6457; *P*-value= 0.0002; **Table.S 9**) as are genetic and geographic distances (r-statistics= 0.1666; *P*-value= 0.0158). However, the correlation between genetic and linguistic distances is not significant (r-statistics= 0.0104; *P*-value= 0.4153). The correlation between genetics and geography increases after controlling for linguistic data, as well as between linguistics and geography after controlling for genetics. A marginally significant negative correlation between linguistic and genetic data is observed after controlling for geography (r-statistics= -0.1291; *P*-value= 0.0496). This overall weaker correlation between genetics and linguistics (while both correlate strongly with geography) could point to separate histories underlying genetic and linguistic data that could involve secondary, and potentially more localized, spread waves. Other explanations are also possible, for example, admixture between linguistically distantly related BSP.

To further explore the possibility of spread-over-spread events, we have compared the genetic diversity of present-day BSP and ancient (aDNA) individuals from Africa, including 12 genomes from this study (97–688 years BP) and 83 individuals (150–8895 years BP) from previous aDNA studies in Africa (**Fig.S 12.1, Table.S 5**). See Meyer et al. ^46^, Steyn et al. ^47^, **Suppl. Material**, and **Table.S 11** for the archeological and morphological descriptions and dating of the individuals for which aDNA data are newly reported here. Dimensionality reduction and clustering analyses support genetic affinities between aDNA and modern samples from the same African region (**Fig. 1d, Fig.S 12.2–12.5**; **Note.S 12**). In South Africa, aDNA individuals (since 688 BP) show homogeneity and genetic affinity with local modern BSP (**Fig. S 12.4–12.9**), thus largely supporting a scenario of genetic continuity since the Late Iron Age. On the contrary, our new aDNA individuals from Zambia (since 311 BP) have a more heterogeneous genetic makeup (**Fig.S 12.4–12.5**) and show genetic affinities with modern BSP from a wider geographical area (**Fig.S 12.7–12.8**), in line with the hypothesis that Zambia might have been a crossroad for different movements of BSP.

### Novel and comprehensive genome-wide dataset as background for future studies

Our dataset, analyzed together with published and newly generated aDNA data, shows its potential to provide an effective modern-day background genetic dataset to compare with aDNA individuals (**Fig. 1d, Fig.S 12.1–12.5**). The underlying historical patterns in BSP are very difficult to distinguish based on modern-day data only. Both IBD and serial-founder models can represent more complex underlying population histories among studied BSP, such as multiple overlapping expansions from the same location following similar routes. A clear manifestation of this pattern has been seen in the comparison of European history inferences based on modern DNA ^48^ and aDNA ^49^. Analyses such as our Mantel test correlations between linguistics, geography, and genetics tentatively point to complex histories and possible spread-over-spread events (**Table.S 9**). In agreement, recent archeological studies also found evidence supporting possible spread-over-spread events for the expansion of BSP ^16^ (**Note.S 1**). Future aDNA studies on human remains from different archeological contexts, associated with the Early, Middle, and Late Iron Age, as well as different pottery traditions in Africa, will be necessary for assessing the affinity of the Bantu-speaker-related ancestry to each other and to current-day BSP. Therefore, the availability of our extensive genomic dataset, encompassing the wide geographic range of the expansion of BSP, will enable further testing of these spread-over-spread event hypotheses using aDNA data from Africa.

## Conclusion

Our data support a large demic expansion of BSP with ancestry from western Africa spreading through the Congo rainforest of central Africa to eastern and southern Africa in a serial founder fashion. In agreement, this finding is supported by patterns of decreasing genetic diversity and increasing genetic distance (*F*_*ST*_) from their point of origin as well as admixture dates with local groups that decrease with distance from western Africa. The significant correlation of admixture times with distance from the BSP source argues for a relatively constant rate of BSP expansion despite the extremely heterogeneous nature of the landscape. While there were corridors of higher and lower effective migration rates across the African landscape, current-day Zambia and the DRC appear to be important crossroads or interaction points for the expansions of BSP. We find tentative evidence that might point to spread-over-spread events in the demographic history of BSP or genetic admixture between linguistically distantly related BSP. Future aDNA studies in Africa and new spatial modeling methods, using our newly generated dataset as comparative data, will help to refine inferences about the expansions of BSP and subsequent episodes in the population history of ancestral BSP and other African populations with which they interacted. The new findings and generated data will be useful not only to population geneticists, archeologists, historical linguists, anthropologists, and historians focusing on population history in Africa, but also to the medical and health sector studying human genetic variation and human health in African and African-descendant populations.

## Acknowledgments

We acknowledge and thank all study participants and local fieldwork teams who helped with the collection of samples. The genotyping and sequencing were performed by the SNP&SEQ Technology Platform, NGI/SciLifeLab Genomics (Sweden). The facility is part of the National Genomics Infrastructure supported by the Swedish Research Council for Infrastructures and Science for Life Laboratory (NGI-SciLifeLab), Sweden. The SNP&SEQ Technology Platform is also supported by the Knut and Alice Wallenberg Foundation. The authors would like to thank George Mudenda, Director of the Livingstone Museum, for his support and letter of approval to sample the archeological human remains housed in the Raymond A Dart collection. We are also indebted to the curators of the Raymond A. Dart Archaeological Human Remains Collection and the University of Pretoria Bone Collection for permission to study the remains. The remains were sampled and exported under SAHRA permits 2789 and 2835. Isotopic measurements were done at the Uppsala Tandem laboratories. We thank the H3Africa Consortium for sharing the whole-genome sequencing H3Africa data. The views expressed in this paper do not represent the views of either the H3Africa Consortium or their funders, the National Institutes of Health (USA) and Wellcome Trust (UK). We thank Natalia Chousou-Polydouri for technical assistance with the linguistic data matrix. The computations/data handling was enabled by resources provided by the National Academic Infrastructure for Supercomputing in Sweden (NAISS) at UPPMAX, partially funded by the Swedish Research Council through grant agreement no. 2022-06725.

## Author contributions

C.M.S. conceived the study. C.F-L, C.B., R.H., A.E., M.V. carried out the analyses. M.V., C.J., L.M., I.M., J.K.M., P.C., T.S.N., C.C., V.C., M.dC., P.E., J.D. M.S., L.B., M.L., A.M, M.S., H.S., B.P., K.B., C.M.S contributed to sample collection and preparation. C.F-L, C.B., R.H., A.E., M.V., H.G., S.P., M.S., H.M., J.R., B.P., K.B., C.M.S. contributed to the interpretation of the results. C.F-L, C.B., R.H., B.P., C.M.S took the lead in writing the manuscript. All authors provided critical feedback, discussed the results and contributed to the final manuscript. C.M.S supervised the project.

## Funding

This project was funded by the European Research Council (ERC) under the European Union’s Horizon 2020 research and innovation programme: AfricanNeo Project awarded to C.M.S. (grant agreement No. 759933). Ancient DNA work in the project is funded by the Knut and Alice Wallenberg Foundation grant to C.M.S. Sample collection in the DRC was funded by the ERC-CoG awarded to K.B. for the BantuFirst project (grant agreement No. 724275). Sample collection in Zambia was funded by the Max Planck Society. B.P. acknowledges support from the ASLAN Project of the Université de Lyon (grant agreement No. ANR-10-LABX-0081) within the French program “Investments for the Future’’ operated by the National Research Agency (ANR). C.A.F-L. received support from the Marcus Borgströms Foundation and the Sven and Lilly Lawski’s Foundation. C.B. received funding from the European Union’s Horizon 2020 research and innovation programme under the Marie Skłodowska-Curie Fellowship Programme (grant agreement No. 839643). H.G. and S.P. received support from the Research Foundation Flanders (FWO) (postdoctoral grants No. 12P8423N and 12ZV721N, respectively). H.M. was supported by the Swedish Research Council (grant agreement No. 2017-02503) and by the Riksbankens Jubileumsfond (grant agreement No. P21-0266).

## Competing interests

Authors declare no competing interests.

### Inclusion and ethics statement

This study was conducted according to the Declaration of Helsinki (World Medical Association 2013). DNA samples were collected with informed consent from participants. Ethical permits and sampling permission were obtained in African countries (Material and Methods) and the study as a whole was approved by the Swedish ethical review board (DNR-2021-01448).

## Data availability

Novel SNP array genotype data of modern-day African populations and whole-genome data of aDNA individuals (bam files) generated in this study will be made available through the European Genome-phenome Archive (EGA) data repository (EGA accessory numbers: *TBD*). Controlled access policies guided by participant consent agreements will be implemented by the AfricanNeo Data Access Committee. Authorized NIH Data Access Committee (DAC) granted data access to C.M.S. for the controlled-access genetic data analyzed in this study that were previously deposited by Scheinfeldt et al., ^32^ in the NIH dbGAP repository (accession code phs001780.v1.p1; project ID: 21567; date of approval: 2019-05-17), and for data of the Hadza and Sabue populations by Crawford et al. ^50^ in the NIH dbGAP repository (accession code phs001396.v1.p1; project ID: 19895; date of approval: 2018-12-18). For the genome-wide genotype data from the Patin et al., ^9^ study (EGA accessory number: EGAD00010001209), data access was granted via European GenomePhenome Archive (EGA) by the GEH Data Access Committee EGAC00001000139. Access to whole-genome sequencing data from the H3Africa Consortium was granted to C.M.S. (EGA accessory numbers: EGAD0000100422, EGAD00001004316, EGAD00001004334, EGAD00001004393, EGAD00001004448, EGAD00001004505, EGAD00001004533, EGAD00001004557, and EGAD00001005076).

## Code availability

Interactive plots and code used for plotting are available in an online repository (https://github.com/Schlebusch-lab/Expansion_of_BSP_Suppl_Material/blob/main/README.md).

## List of Supplementary Materials

### Materials and Methods

**Supplementary Notes.S 1 – S 12**

**Table.S 1 – S 15**

**Fig.S 1 – S 13.2**

## Notes

### Competing Interest Statement

The authors have declared no competing interest.

