## Supplementary Text for "The genetic legacy of the expansion of Bantu-speaking peoples in Africa"

**Table of contents** **1**

**1. Material and Methods** **2**
1.1. Summary of Material and Methods 2
1.2. Ethics approval 4
1.3. Genotyping procedure 5
1.4. Ancient DNA samples from sub-Saharan Africa 6
1.4.1. Description of human remains 6
1.4.2. Sampling procedure 7
1.4.3. DNA Extraction from human remains 7
1.4.4. Library building and sequencing 7
1.4.5. Sequence data processing 8
1.5. Assembling genome-wide genotype datasets 8
1.6. Population structure analyses 9
1.7. Unsupervised clustering analyses 9
1.8. Testing analysis of f3- or f4-statistics 10
1.9. Local ancestry deconvolution approach 10
1.10. Runs of homozygosity and inbreeding coefficients 11
1.11. Detection of shared segments inherited from a common ancestor 11
1.12. Estimating the timing of and strength of founder events 12
1.13. Haplotype diversity and linkage disequilibrium analyses 12
1.14. Phylogenetic analyses 13
1.15. Testing models of isolation-by-distance 13
1.16. Testing spatially explicit models of the Bantu-speaker expansion 14
1.17. Estimating effective migration rates 14
1.18. Gene-flow barriers analysis on a grid 15
1.19. Admixture timing inference 15
1.20. Correlations between linguistic, geographical, and genetic data 15
1.21. Comparisons between ancient and present-day populations 16
**2. Supplementary Notes** **16**
Note.S 1. Further background on the Bantu-speaker expansion 16
Note.S 2. Patterns of population structure and genetic diversity 17
Note.S 3. Ancestry-specific analyses in Bantu-speaking populations 19

Note.S 4. Patterns of consanguinity and founder events 19
Note.S 5. Effective population size and founder events in BSP 20
Note.S 6. Patterns of isolation-by-distance among BSP 20
Note.S 7. Patterns of haplotype diversity and linkage disequilibrium among BSP 21
Note.S 8. Phylogenetic analyses of Bantu-speaking populations 22
Note.S 9. Patterns of pairwise genetic distances and admixture graphs 23
Note.S 10. Estimation of effective migration surfaces 23
Note.S 11. Admixture dating and model-testing of the Bantu-speaker expansion 24
Note.S 12. Comparisons between ancient and present-day populations in Africa 24
**3. Supplementary References** **26**

### 1. Material and Methods

#### 1.1. Summary of Material and Methods

**Genotyping and assembled datasets.** Samples newly presented in this study were collected in fourteen sub-Saharan African countries. Ethical permits and sampling permission were obtained in African countries and the study as a whole was approved by the Swedish ethical review board (DNR-2021-01448). In total, 1,740 samples were genotyped (**Fig.S 1.2–1.3** and **Table.S 1**) on the Illumina H3Africa array (~2.4 million SNPs) using seven genotyping batches at the SNP&SEQ Technology Platform, NGI/SciLifeLab Genomics (Sweden). After merging all newly genotyped data and quality control (QC) steps using PLINK v1.90b6.4 [1], genotype data consist of 2,221,827 autosomal SNPs. After removing 67 samples due to low genotyping rate, and 105 individuals due to their first- or second-degree kinship with other samples, we obtained 1,568 individuals and 2,221,827 SNPs for the “Genotyped” dataset. After merging the Genotyped dataset with comparative data and performing QC-steps, we assembled the “Full-Genotyped” dataset that contains 482,459 SNPs and 5,341 individuals from 227 populations (including 81 populations with sample sizes lower than 10 individuals), and three sub-datasets with selected African populations. We included 4,950 individuals from 124 African and Eurasian populations in the “AfricanNeo” dataset (**Fig.S 1.4a–1.4b**); 3,902 individuals from 111 sub-Saharan African populations in the “Only-African” dataset (**Fig.S 1.5a** and **1.5c**); and 2,108 individuals from 67 populations speaking Bantu languages (henceforth “BSP”) in the “Only-BSP” dataset (**Fig.S 1.5b, 1.5d** and **Fig.S 1.6**). BSP with less than 10 individuals were removed from certain datasets. To avoid sample-size biases, for some analyses (e.g. local ancestry inference, and analyses using the masked and imputed data) populations with large sample sizes were randomly downsampled to 30 individuals, and we obtained 1,495 individuals from 124 populations in the downsampled AfricanNeo dataset.

**Dimensionality reduction and clustering methods.** To better visualize genetic variation and population structure in BSP, we applied four dimensionality reduction methods for genome-wide SNP data. We first used the uniform manifold approximation and projection (UMAP) approach [2] directly on the genotype data. Second, we applied principal component analysis (PCA) using *smartpca* [3], and then used PCA-UMAP approach to combine the information of the first ten PCs [4]. Fourth, we used the genotype convolutional autoencoder (GCAE) method [5]. In addition, we applied an unsupervised clustering-based approach using ADMIXTURE software v1.3.0 [6] from K=2 to K=25.

**Ancient DNA samples.** To compare the genetic affinities of ancient and present-day BSP, we merged the AfricanNeo dataset with 12 ancient DNA (aDNA) individuals from southern and south-central Africa (currently Zambia and South Africa), and 83 aDNA individuals from previous studies [7–13] (**Fig.S 12.1** and **Table.S 5**). We then projected aDNA samples onto a background of present-day populations using PCA. After merging haplodized modern samples and pseudo-haplodized aDNA individuals and performing LD-pruning steps, we used unsupervised ADMIXTURE analysis.

**Runs of homozygosity.** We used PLINK to calculate five parameters of runs of homozygosity (ROH) in BSP and worldwide populations: mean ROH size, total length of ROH, sum of short ROH, sum of long ROH, and ROH-based inbreeding coefficient (or $F_{ROH}$). For each population, we also calculated six ROH length classes. To estimate effective population sizes over the last 50 generations, we used IBDNe [14]. To infer both the age and strength of demographic founder events in BSP, we used ASCEND v10 [15]. To identify significant founder events, we followed the four criteria recommended by Tournebize et al. [15].

**Admixture timing analysis.** To estimate admixture dates, we applied haplotype-based admixture inference methods. First, we used MOSAIC v1.4 [16] for two- and three-way admixture models for BSP that were included in the AfricanNeo dataset. For haplotype phasing of the AfricanNeo dataset, we used SHAPEIT v2.r904 [17]. For local ancestry inference, we estimated haplotypic admixture in individuals from BSP using RFMix software v1.5.4 [18]. To avoid the influence of admixture patterns in BSP in our ancestry-specific analyses, we removed haplotypes with non-WCA-related ancestry from each haploid genome of each Bantu-speaking individual using a masking approach. For each assembled dataset, we explored patterns of population structure between and within populations using *smartPCA* [3].

**Phylogenetic analyses and correlations.** To investigate phylogenetic relationships between all the BSP, we used TreeMix v1.13 [19]. The likelihood of each proposed population-based maximum-likelihood (ML) TreeMix topology was assessed by bootstrapping blocks of 500 SNPs and assigning Juhoansi from Namibia as the root of the population tree. To test the correlation between genetic, linguistic, and geographical distances, we performed Mantel tests and partial Mantel tests using the R package ncf [20]. As genetic distances, we computed pairwise $F_{ST}$ between populations included in the ancestry-masked Only-Bantu dataset using EIGENSOFT package v6 [21]. Geographic distances were calculated as pairwise great circle distances between the studied populations using the R package *geosphere* [22]. For linguistic distances, we used a linguistic dataset from the multistate matrix of cognate sets identified by Grollemund et al. [23]. In total, 38 BSP matched the genetic dataset and the linguistic dataset of 409 Bantu languages studied by Grollemund et al. [23].

**Patterns of genetic diversity.** To investigate spatial patterns of genetic diversity of studied African populations, we calculated statistics based on haplotype and linkage disequilibrium information. Haplotype heterozygosity (HH) and haplotype richness (HR) were computed following recommendations from Schlebusch et al. [24] with homemade scripts implemented in Python. Values were calculated per chromosome and then averaged across the genome. Each calculation was also repeated 10 times. We characterized linkage-disequilibrium (LD) patterns in each population with more than 10 individuals by measuring the correlation coefficient ($r^2$) between all pairs of SNPs within 500 Kbp windows using PLINK. To assess whether LD, HH, and HR patterns in BSP were consistent with a history of expansion from the homeland of BSP, we calculated the correlation between the three summary statistics and geographical distance from Cameroon, assuming that the Bantu-speaker expansion started in that region [25].

**Pairwise genetic distances.** To reconstruct potential routes of expansion of BSP, pairwise $F_{ST}$ values were calculated between one population from Cameroon (Nzime) and each of the studied BSP for the masked and imputed Only-BSP dataset. We also applied the GenGrad method from

Pagani et al. [26], but using $F_{ST}$ as the genetic distance metric and with slightly adjusted parameters to better fit the smaller study area.

**Effective migration rates.** To further investigate spatial population structure in sub-Saharan African populations, we used EEMS software [27] and its implementation FEEMS [28]. EEMS analysis was repeated three times and an average was taken as input for the visualization as recommended in the EEMS manual. Both, EEMS and FEEMS were performed on the basis of the Only-African and Only-BSP datasets before and after using the masking approach.

**Testing isolation-by-distance models.** To test four models of migration, we used SpaceMix v0.13 [29]. The software generates geo-genetic maps where genetics rather than physical distances determine the distances between individuals/populations. The general underlying assumption evaluated with SpaceMix is that under an isolation-by-distance pattern, geographic and geo-genetic positions will be similar, which is a pattern of isolation-by-distance. The "best" fitting model was evaluated using Pearson correlations between the expected and observed data.

**Testing models of migration routes.** To test different demographic scenarios for the Bantu-speaker expansion, we used a spatiotemporally explicit population genetic framework [30,31]. Here, we adapted the extension of the model presented by Raghavan et al. [31] to apply multiple local expansions for different scenarios of expansion. We considered three demographic scenarios in which the expansion of BSP proceeded north of the rainforest, south through a rainforest corridor, or using both northern and southern routes. For each demographic scenario, we ran one million simulations with parameters drawn from an independent uniform distribution for parameters characterizing the Bantu-speaker expansion, and from a log-uniform distribution for parameters describing the initial global expansion of AMH taken from Raghavan et al. [31].

#### 1.2. Ethics approval

New samples presented in this study were collected in large-scale sampling campaigns conducted in fourteen sub-Saharan African countries: Angola (N= 34), Botswana (N= 27), Central African Republic (CAR; N= 81), the Democratic Republic of the Congo (DRC; N= 592), Lesotho (N= 8), Mozambique (N= 36), Namibia (N= 130), Rwanda (N= 25), South Africa (N= 391), Swaziland (present-day the Kingdom of Eswatini; N= 17), Tanzania (N= 16), Uganda (N= 72), Zambia (N= 242), and Zimbabwe (N= 40) (**Table.S 1**). Each participant gave informed consent before donating their samples. Where information was available, only individuals whose parents and grandparents came from the same ethnolinguistic group were included in the study.

This study was conducted according to the Declaration of Helsinki [32]. This study as a whole was approved by the Swedish Ethical Review Authority (Ministry of Education, Sweden). Biological samples for this study were in part supplied by our collaborators who obtained the original ethical permission for the sampling in African countries. Himla Soodyall was granted ethics approval by the Human Research Ethics Committee (Medical) (University of the Witwatersrand, South Africa; protocol Nr. M180656). Brigitte Pakendorf was granted ethics approval by the Biomedical Research Ethics Board (University of Zambia, Zambia; protocol number: 004-08-07). Vinet Coetzee was granted ethics approval by the Faculty of Natural and Agricultural Sciences Ethics Committee (University of Pretoria, South Africa; protocol number: EC160429-024 and 259/2016). Koen Bostoen was granted ethics approval by the Swedish National Ethics Committee (Sweden; protocol number: Dnr 2019-05244), as well as permission for sampling in DRC from the Minister of Arts and Culture (DRC; protocol number: Nr 091/CAB/MIN/CA/PKB/2018). Maryna Steyn received clearance from the Raymond A. Dart Archaeological Human Remains Collection (12/10/2017) and obtained permits to sample and export specimens from the South African Heritage Resources Agency (SAHRA).

##### 1.3. Genotyping procedure

In total, we genotyped DNA samples of 1,740 individuals for this study (**Table.S 1** and **Fig.S 1.2–1.3**), including 1,487 Bantu-speaking individuals from 143 African populations spanning 14 sub-Saharan African countries, 8 Khoe-San populations from four African countries (in Angola: Khwe (N=17) and Xun (N=17); in Botswana: GuiGhanaKgal (N=11); in Namibia: Damara (N=23), Nama (N=16), and Juhoansi (N=11); in SouthAfrica: Karretjie (N=10) and Khomani (N=32); 137 individuals in total), five Ubangi-speaking populations from CAR (Banda, DzangaShanga people, Gbaya, Nzakara, and Zande; 70 individuals in total), and South African Coloured individuals (N= 46). All the DNA samples were genotyped on the Illumina H3Africa array, designed by the H3Africa Consortium [33]. This genotyping array was specially designed for SNP-genotyping of 2,271,503 SNPs, to account for the larger genetic diversity and smaller haplotype segments in African populations [33]. Seven of the Khoe-San groups and the Coloured individuals were previously included in Schlebusch et al. [24], where they were typed on the Illumina Omni 2.5M bead chip. Here the same individuals were re-genotyped on the Illumina H3Africa array to increase overlap with the current study as well as to include additional individuals from the same populations.

Genotyping was performed at the SNP&SEQ Technology Platform, NGI/SciLifeLab Genomics (Sweden), and the results were analyzed using the software GenomeStudio (v2.0.3, Illumina Inc). Genotype data were exported from the forward strand according to polymorphism data from dbSNP v131 [34] and using the human reference genome build version 37 (or hg19).

Samples were genotyped in seven batches, as follows:

- The first batch (called "SE-2209_191110") included 2,267,346 variants and 1,157 individuals, and was generated for this study using BeadChip type: H3Africa_2017_20021485_A2. The average SNP call rate per sample was 97.11% (range: 21.58%-99.64%), and the reproducibility was 99.95% (6,201 conflicts in 13,547,660 duplicate tests). Fifty-one pairs of samples were identified as being above our threshold of 80% similarity in genotype data in the test.
- The second batch (called "TE-2567_201023_2017") included 2,267,346 variants and 4 individuals for this study and was generated using BeadChip type: H3Africa_2017_20021485_A2. The average SNP call rate per sample was 95.46% (range: 18.10-99.60%), and no samples were not identified as being above the threshold of 80% in this test.
- The third batch (called "TE-2567_201023_2019") included 2,271,503 variants and 298 individuals for this study and was generated using BeadChip type: H3Africa_2019_20037295_B1. The average SNP call rate per sample was 99.11% (range: 53.49-99.52%), and the reproducibility was 99.98% (1,796 conflicts in 9,023,008 duplicate tests). Two pairs of samples were identified as being above the threshold of 80% similarity (between 069KKT and 030KKT; and between 026KKT and 073KKT).
- The fourth batch (called "TI-2658_201112") included 2,271,503 variants and 135 individuals and was generated for this study using BeadChip type: H3Africa_2019_20037295_B1. The average SNP call rate per sample was 97.74% (range: 34.32%-99.48%). Two pairs of samples were identified as being above the threshold of 80% similarity (LD162 and LD132; SK013 and SK021).
- The fifth batch (called "RK-2011_190308") included 2,267,346 variants and 46 individuals for this study and was generated using BeadChip type: H3Africa_2017_20021485_A2. The average SNP call rate per sample was 99.31% (range: 93.46%-99.61%), and the reproducibility was 99.99% (911 conflicts in 6,710,668 duplicate tests). No pair of samples was identified as being above the threshold of 80%.
- The sixth and seventh batches (called “TC-2508_200401_A” and “TC-2508_200401_B”, respectively) were genotyped at the University of Pretoria (South Africa), and both batches included 2,267,346 variants and 100 individuals in total for this study, 41 individuals genotyped in the sixth batch and 58 genotyped in the seventh batch.

#### 1.4. Ancient DNA samples from sub-Saharan Africa

To compare the genetic diversity of ancient and present-day BSP, we merged the Only-African dataset with 95 ancient DNA (aDNA) individuals (**Fig.S 12.1** and **Table.S 5**). Among them, 12 individuals were newly sequenced for this study (**Table.S 11**) and 83 individuals were retrieved from previous studies [7–13].

##### 1.4.1. Description of human remains

The 12 new ancient human remains in this study came from various caves and rock shelters in Zambia and South Africa (**Table.S 5** and **11**). We obtained permission from the South African Heritage Resources Agency (SAHRA) to sample and export bones for ancient DNA analyses. Nine WUD samples (permit number: 2789) are from the Raymond A. Dart Archaeological Human Remains Collection (Dart Collection) located at the School of Anatomical Sciences, University of the Witwatersrand (Johannesburg, South Africa). Three UPS samples (permit number: 2804) are from the Archaeological Human remains Collection (Pretoria Bone Collection) situated within the Department of Anatomy, University of Pretoria (Pretoria, South Africa). For both collections Prof. Maryna Steyn is the permit holder. The archeological context, morphological assessments, and dating of the remains were described before for six of the samples: WUD034 and WUD037 (C1 and C9 in Meyer et al. [35]; and WUD003, WUD004, WUD008, and WUD010 [36]. WUD038b sample originated from an archeological site in KwaZulu Natal but is curated in the Dart Collection. WUD012 (Chipongwe Caves) was collected in 1930 by Raymond Dart [37]. WUD012 and WUD018 were originally collected in current-day Zambia and are curated in the Dart Collection, while UPS013, UPS017a, and UPS029 are kept in the Pretoria Bone Collection. Little is known about their archeological contexts.

Six samples (WUD038b, WUD012, WUD018, UPS013, UPS017a, and UPS029) were accelerator mass spectrometry (AMS) radiocarbon dated at the Tandem Laboratory (Department of Physics and Astronomy, Uppsala University, Sweden). Radiocarbon dates were calibrated with OxCal 4.4 [38] using the atmospheric curve SHCal20 [39] and are given at 95.4% probability (2σ), see **Table.S 5** and **Table.S 11**.

##### 1.4.2. Sampling procedure

The clean sampling of the bones was done on-site in South Africa and only the bone samples were transported to Uppsala University (Sweden). Following recommendations proposed by Schlebusch et al. [8], we used a bleach-decontaminated (RNase AWAY, ThermoScientific) enclosed sampling tent with adherent gloves (Captair Pyramid portable isolation enclosure (Erlab)). Prior to sampling and DNA extraction, the bones were cleaned to prevent contamination of the samples with modern DNA. First, the samples were ultraviolet irradiated (254 nm) for 20 minutes on each side, then the outer surface of the bones was removed by gentle scraping at low speed using a Dremel 8100 and wiped with 0.5% bleach (NaOH) and sterile water (HPLC grade, Sigma-Aldrich). The cleaned bones were ultraviolet irradiated (254 nm) again for 20 minutes per side. A piece of each bone, of between 50 and 80 mg, was cut off for aDNA analysis, using either a circular diamond cutting wheel for the tooth roots or a core drill for the petrous portion of temporal bones. Whenever possible, at least two bones from the same individual were sampled (**Table.S 11**).

##### 1.4.3. DNA Extraction from human remains

To increase the amount of endogenous aDNA, we extracted DNA from the sampled bone/tooth root pieces instead of extracting DNA from powdered bone [40]. The bone samples were pre-digested in 1 mL EDTA (0.5M, pH 8) for 30 min at 37°C, the EDTA was then discarded and an overnight digestion in 1 ml of EDTA (0.5M, pH 8.0) and 25 µl proteinase K (10 mg/ml) at 37°C were conducted.

Then, 10.4ml of Binding buffer (245 ml of PB buffer from Qiagen, 7.5 ml of sodium acetate, 3M, and 0.625 ml of sodium chloride, 5M) were added to the DNA digests, and the DNA molecules were purified using a silica-columns method and eluted in 110 µl of EB buffer, following a protocol as in Dabney et al. [41] with modifications as in Rohland et al. [42]. Two extraction negative controls were processed for every 10 samples extracted.

##### 1.4.4. Library building and sequencing

Double-indexed blunt-end DNA libraries were prepared from 20 µl of extract using P5 and P7 adapters as in Meyer and Kircher [43], with the shearing step omitted. Two library blanks were processed for every 12 samples extracted (including the extraction blanks). The optimal number of PCR cycles used for library amplification was determined using quantitative PCR (qPCR). The 25 µl qPCR reactions were set up in duplicates and contained 1 µl of DNA library, 1X Maxima SYBR Green Mastermix and 200 nM of each IS7 and IS8 primers (10 uM). To calculate the number of cycles for PCR amplification, two cycles were added to the qPCR Cq value (value at which the qPCR amplification was saturated and reached a plateau). Each library was then amplified using between 12 and 21 PCR cycles. The PCR reactions were set up in duplicates using 6µl of library DNA, Index primers P7 and P5 (10 uM) and 1µl of AmpliTaq Gold™ polymerase (Thermofisher Scientific). One negative PCR control was set up for every four reactions. The duplicate reactions were amplified according to the supplier recommendations (ThermoFisher Scientific) with an annealing temperature of 60°C. The amplified duplicates were first pooled and then purified using AMPure XP Beads (Agencourt). The resulting libraries were quantified on a TapeStation using a High Sensitivity kit (Agilent Technologies). Equimolar pools of 25-30 libraries were prepared for sequencing at the Swedish National Genomics Infrastructure (NGI)-SciLifeLab in Uppsala (Sweden) on Illumina NovaSeq 6000 SP lane (one pool per lane), with a paired-end 150 bp chemistry. The negative controls did not yield any DNA and were therefore not sequenced.

##### 1.4.5. Sequence data processing

The initial data processing of the shotgun data was as follows. Adapters were trimmed with cutadapt v. 2.3 [44] and the pair-end reads of each library were merged if the two reads overlapped at least 11 base pairs using FLASH v. 1.2.11 [45]. Bwa aln 0.7.13 [46] was then used to map them as single-end reads to the human reference genome (hg19). Non-default parameters for bwa were -l 16500 -n 0.01 -o 2 [47,48]. Reads with identical start and end positions were identified as PCR duplicates and collapsed using a modified version of FilterUniqSAMCons_cc.py [49], which ensures the random assignment of bases in a 50/50 case. Reads with less than 10% mismatches to the human reference genome and longer than 35 base pairs were retained for further analysis. Reads were authenticated by looking at read length distribution and damage patterns (**Fig.S 12.0**).

Sample contamination estimates were performed with two methods. We first ran the likelihood-based method ContamMix [50], which uses phylogenetically informative sites on the mitochondrial genome to estimate the proportion of authentic DNA ("clean" DNA). Briefly, the consensus sequence obtained from each ancient sample is aligned with 311 reference mt genomes [51] and then each read is tested against all these 312 sequences. If reads map better to one of the 311 reference mt sequences, they might be caused by contamination. Second, we run verifyBAMid [52], which checks for autosomal contamination. The method checks if the target reads match a set of known reference genotypes and if they constitute a mixture of two samples. We used the 1000 Genomes Project Phase 3 data as potential reference contaminants. Results are shown in **Table.S 11**.

To determine biological sex, we implemented the sex determination method previously described [53,54]. It uses reads with a mapping quality of at least 30 and calculates the ratio of reads mapping to the Y-chromosome and those reads mapping to both X- and Y-chromosomes. Strict mitochondrial consensus sequences were generated with SAMtools v1.5 [55] options mpileup and vcfutils.pl. Base and mapping quality scores were set to a minimum of 30 and only SNPs with at least 3-fold coverage were used. Mitochondrial haplogroups were assigned with HaploGrep v2.1.16 [56] using the mtDNA tree built by PhyloTree Build 17 [57]. To account for low coverage data, the SNPs in the ancient samples were called as follows: at each SNP site, a random read with minimum mapping and base quality 30 was drawn and the allelic status at that read was coded to be the hemizygous genotype of the individual (file-formats require diploid genotypes and we use the homozygote code for the record, but the data are treated as hemizygote in all downstream analyses). Sites showing additional alleles, indels or missing data were removed as well as transitions sites with T or A.

##### 1.5. Assembling genome-wide genotype datasets

For autosomal data, quality control (QC) steps were performed using PLINK v1.90b6.4 [1], to keep autosomal biallelic variants with a high genotyping rate (>90%). We used the same filtering before and after merging for each dataset (as follows: plink --mind 0.15 --geno 0.1 --hwe 0.0000001), and samples were removed due to their low genotyping rate (mind> 0.15). To estimate recent genetic relatedness, we used KING v1.4 [58] and the PC-Relate tool included in the R GENESIS package [59], and we removed 67 samples due to their low genotyping rate, and 106 individuals were removed due to first- or second-degree kinship. After QC steps, genotype data consist of 2,221,827 autosomal SNPs.

To merge the newly genotyped dataset with publicly available datasets, we first collected data from datasets of whole-genome sequencing studies [60], and datasets previously genotyped on Illumina HumanOmni5, Illumina HumanOmni2.5 or H3Africa arrays. Then, genome-wide SNP data were merged with reference populations presented in previous studies of our group [24,61,62] or by other previous studies [63–71] (details and accessory numbers are included in **Table.S 2**). After merging all newly genotyped data and quality control (QC) steps, we assembled the full dataset that contains 482,459 SNPs and 5,341 individuals from 227 populations, and two sub-datasets with selected populations (**Fig.S 1.6**). Due to the small sample size in some populations, BSP with less than 10 individuals were removed from the dataset, and we obtained 4,950 individuals from 124 African and Eurasian populations in the “AfricanNeo” dataset (**Table.S 2**, **Fig.S 1.4** and **Fig.S 1.6**); 3,902 individuals in the “Only-African” dataset for only 111 sub-Saharan African populations included in the AfricanNeo dataset (**Fig.S 1.5a** and **1.5c**); and 2,108 individuals in the “Only-BSP” dataset for 67 BSP included in the AfricanNeo dataset (**Fig.S 1.5b** and **1.5d**). We also used PLINK to prune SNPs under high LD (plink “--indep-pairwise 100 10 0.2”) for analyses that assume unlinked variation. After LD-pruning, the AfricanNeo LD-pruned dataset contains 223,473 variants and 268 individuals.

Of note, in our plots the Damara population from Namibia is included in the group BSP although they speak a Khoisan language (hereafter “Damara-KSP”), while the Baka population from Cameroon and Gabon is included in the group wRHG although they speak Bantu languages. This is because these groups have genetic backgrounds that distinguish them from the language they currently speak and they are likely to have undergone language shifts in the past [72–74].

To avoid sample-size biases, for some analyses (e.g., local ancestry inference, and analyses using the masked and imputed dataset) BSP with large sample sizes were randomly downsampled to 30 individuals, and we obtained 1,495 individuals from 124 populations in the downsampled AfricanNeo dataset.

##### 1.6. Population structure analyses

To explore patterns of population structure within the studied populations, we applied four dimensionality reduction methods for genotype data. First, we used the uniform manifold approximation and projection (UMAP) approach [2] directly on genome-wide SNP data. We used an in-house script (https://github.com/Hammarn/Scripts/blob/master/POPGEN/UMAP_plot_bed.py). Second, we performed PCA for the data assembled in each dataset using *smartpca* software from the Eigensoft package v7.2.1 [3]. We then plotted PCA results between PC projections from PC1 to PC10. Third, we performed PCA-UMAP to combine the information of the first ten PC projections [4]. We used an in-house Python script (script is available in GitHub; https://github.com/Hammarn/Scripts/blob/master/POPGEN/UMAP_plot.py). This method considers neighboring samples around each data point in the PCA and seeks a lower-dimensional representation that preserves the distances between the points in the neighborhood. Fourth, we applied a deep learning framework for dimensionality reduction based on genotype convolutional autoencoder (GCAE) [5]. We plotted the results of each method by using in-house Python scripts and the bokeh visualization library for interactive plots (in *.html format). To better visualize the results, we provided plots highlighting each group with different colours and markers, as well as plots for each studied population.

##### 1.7. Unsupervised clustering analyses

Admixture fractions were estimated using ADMIXTURE software v1.3.0 [6,75]. To cluster individuals based on SNP genotypes, we carried out unsupervised ADMIXTURE clustering analysis. First, we investigated African populations from the AfricanNeo dataset. To investigate the geographical distributions of ADMIXTURE results at K=4, we plotted estimated admixture proportions on a geographic map. We used the grid-based mapping Surfer software (Golden Software, LLC) and applied the Kriging method for spatial interpolation. Unsupervised ADMIXTURE analyses were also performed on the basis of only the AfricanNeo dataset from K=2 to K=25 using 10 independent runs with a random seed for each K-group, and default settings. A cross-validation (CV) test was performed for each run of each K-group. To visualize ADMIXTURE results for all the K-groups, we used PONG v1.5 [76]; however population labels were difficult to plot and read for most of the 124 labels. To better visualize the major mode of the ADMIXTURE result, we plotted the Q matrix for the major mode in the 10 runs in bar plots using the AncestryPainter graphic program v5.0 [77]. For a better comparison, the width of each population was set to be equal regardless of its sample size. We selected the results for the K-groups that are more informative for the studied populations (K=2, K=4, K=6; K=12, and K=16). For each selected K-group, we used custom R and Python scripts to plot the averages for each estimated component in each population in pie charts on geographic maps, bar plots, and for ADMIXTURE results at K=6 in the ternary diagram with a hexagonal shape.

##### 1.8. Testing analysis of f3- or f4-statistics

We formally tested the hypothesis of admixture in the present-day BSP using the f3- and f4-statistics in the form *f3(Yoruba; admixture-source*, *Target*) and *f4(Target, Yoruba; admixture-source, CHB)*, respectively. The *f3* test assessed if any of the BSP (*Target* population) had significant genetic affinities with a local non-BSP as the source of admixture. The *f4* test assessed if any of the BSP (*Target* population) was more closely related to either the outgroup Chinese-Han (CHB) or a local non-BSP as the admixture source. To disentangle the differential contribution of different hunter-gatherer groups into western and southern BSP, we also performed f3 tests in the form *f3(Ju-hoansi; Baka, Target)* and f4 tests in the form *f4(Target, Yoruba; Baka,Ju-hoansi)*. For this analysis, we used the package ADMIXTOOLS v2 (https://github.com/uqrmaie1/admixtools) under the R environment and functions *qp3pop* and *qpdstat*, respectively, with default options. To maximize the amount of information available, we run the test on the SNPs common to each triplet (respectively, quadruplet) of attested populations.

With the same tools, we also estimated *f3* for the 12 new aDNA individuals in the form *f3(Yoruba; ancient-sample, BSP)* to assess the affinity of each ancient individual to a current-day BSP. For each test, we merged each ancient individual with the unmasked AfricaNeo dataset separately, in order to maximize the number of SNPs available for each analysis.

##### 1.9. Local ancestry deconvolution approach

To estimate the continental and subcontinental haplotypic admixture along with phased chromosomal segments (ancestry tracts), we applied a local admixture inference approach using RFMix software v1.5.4 [18]. To maximize phasing accuracy, the dataset was phased using the Haplotype Reference Consortium reference panel [78], and the HapMap Phase II b37 reference map as a genetic map [17]. To deconstruct genomes of BSP into six ancestry tracts, we compared haplotypes of BSP with the haplotype diversity of six panels of putative source populations. Each panel included 30 randomly selected individuals from:

(*i)* Niger-Congo-speaking populations (8 Esan-ESN, 13 Yoruba-YRI, and 9 Igbo individuals);

(*ii)* Western hunter-gatherer populations (30 Baka individuals from Cameroon and Gabon);

(*iii)* Eastern hunter-gatherer and Nilo-Saharan-speaking populations (13 Hadza, 7 Sabue, and 10 Gumuz individuals);

(*iv)* Afro-Asiatic-speaking populations (9 Amhara, 6 Oromo, and 15 Somali individuals);

*(v)* Khoe-San groups (15 Juhoansi, 4 GuiGhanaKgal, and 11 Karretjie individuals); and

(*vi)* Eurasian populations (10 Iberian-IBS, 10 European descendants-CEU, 5 Lebanese, and 5 Yemeni individuals).

To avoid the influence of admixture patterns in BSP in our ancestry-specific analyses, we removed haplotypes without West-Central African (WCA)-related ancestry in each haploid genome of each Bantu-speaking individual using a masking approach. Haplotypes with WCA-related ancestry were previously identified using RFMix with two expectation-maximization runs (EM=2). To avoid issues with missing data after masking, we phased and imputed the masked haploid genomes of studied Bantu-speaking individuals using SHAPEIT2 for phasing, IMPUTE v2.3.2 [79,80] for imputation and Niger-Congo-speaking populations from Nigeria (YRI, ESN, and Igbo genomes; 306 individuals in total) as a reference panel for this phasing and imputation. We then merged masked, phased, and imputed data of BSP with unmasked data of reference populations from previous studies to compare the outcome of this approach for masking, phasing, and imputation (**Fig.S 13.1–13.2**).

##### 1.10. Runs of homozygosity and inbreeding coefficients

To calculate runs of homozygosity (ROH), we used a sliding-window approach and followed recommendations from Ceballos et al. [81]. First, we extracted all the individuals from each population, and we performed more restrictive filtering separately for each population (to remove SNPs due to “--maf 0.01” and “--hwe 0.001”). For each population, we then used PLINK v1.9 to apply the following parameters: 30 was the minimum number of SNPs that one ROH was required to have (“--homozyg-snp 30”); 300 was the length in kb of the sliding window (“--homozyg-kb 300”); 30 was the required minimum density to consider one ROH that means one SNP in each 30 kb (“--homozyg-density 30”); 30 was the number of SNPs that the sliding window must have (“--homozyg-window-snp 30”); 1000 was the length in kb between two SNPs to be considered in two different segments (“--homozyg-gap 1000”); 1 was the number of heterozygous SNPs allowed in each window (“--homozyg-window-het 1”); 5 was the number of missing calls allowed in a window (“--homozyg-window-missing 5”); 0.05 was the proportion of overlapping window that must be called homozygous to define a given SNP as in a “homozygous” segment (“--homozyg-window-threshold 0.05”).

For ROH longer than 1.5 Mb, we calculated the following parameters: mean ROH size, sum of long ROH, and total length of ROH. For ROH shorter than 1.5 Mb, we calculated the sum of short ROH. For each population, we also calculated six ROH length classes: class 1 (between 0.3<ROH<0.5 Mb), class 2 (between 0.5<ROH<1 Mb), class 3 (between 1<ROH<2 Mb), class 4 (between 2<ROH<4 Mb), class 5 (between 4<ROH<8 Mb), and class 6 (between 8 Mb<ROH<10Mb). The genomic inbreeding coefficient based on ROH (or $F_{ROH}$) was obtained as the total sum of ROH >1.5 Mb divided by the total length of the autosomal genome (3 Gb) [82,83].

##### 1.11. Detection of shared segments inherited from a common ancestor

To investigate recent demographic events, we analyzed identity-by-descent (IBD) segments for each population included in the AfricanNeo dataset. We employed the fastIBD algorithm in the Beagle package v4.1 [84] to detect shared identical-by-descent segments between pairwise populations. Firstly, we calculated the total amount of IBD shared between individuals in tested populations. Then we calculated the average amount of shared IBD in centimorgans (cM) between individuals in tested populations. We removed IBD segments of 3cM to avoid the conflation effect of short IBD segments [85]. To estimate effective population sizes over the last 50 generations, we used IBDNe [14].

##### 1.12. Estimating the timing of and strength of founder events

To infer both the age and strength of demographic founder events in BSP, we used ASCEND v10 [15]. This method estimates the correlation in allele sharing across the genome between pairs of individuals to recover signatures of past bottlenecks in each studied population. We performed ASCEND analysis for each population included in the AfricanNeo dataset, using default settings for all the autosomal chromosomes (i.e. distance bins from 0.1cM to 30.0cM by steps of 0.1cM) and default settings for plotting the results. We estimated the age since the founder event (*Tf*, in generations before the present, BP), the strength of the founder event (% *If*), and the 'normalized root mean squared deviation' (NRMSD) between the empirical decay curve and the theoretical decay curve. To convert the inferred dates since the founder event from generations to years, we used the following equation: 1950-(g*29), where g is the estimated number of generations and 29 is the number of years for one generation [86]. To identify significant founder events, we followed the four criteria recommended by Tournebize et al. [15]: (*i*) the 95% confidence intervals of the estimated founder age and intensity do not include 0; (*ii*) the estimated founder age is lower than 200 generations and its associated standard error is lower than 50 generations; (*iii*) the estimated founder intensity is greater than 0.5%; and (*iv*) the NRMSD is lower than 0.29.

##### 1.13. Haplotype diversity and linkage disequilibrium analyses

To further investigate the diversity of our study populations, we calculated haplotype heterozygosity (HH) and haplotype richness (HR) following recommendations from Schlebush et al. [24]. We used in-house scripts to estimate those parameters in haplotype windows ranging in size from 1 kb to 100 kb (with 1 kb increments). These metrics measure the number of different haplotypes in each population, and uneven sample size in a population can have a large influence on the calculation. To circumvent this issue, we excluded populations with fewer than 10 individuals and randomly subsampled the remaining populations to 10 individuals. Each calculation was also repeated 10 times and only the average result was reported. For each population and subsample, we removed SNPs with more than 10% missing data and minimum allele frequency (MAF) lower than the threshold of 10%. Following the recommendations of Schlebusch et al. [24], windows with 5 or more

SNP were downsampled to 5, and windows with fewer SNPs were excluded from the analysis. Values were calculated per chromosome and then averaged across the genome.

To characterize linkage disequilibrium (LD) patterns of all populations with more than 10 individuals (in total 124 populations), we measured the correlation coefficient ($r^2$) between all pairs of SNPs within 500 Kbp windows using PLINK. To control for uneven sample sizes, all populations with more than 10 individuals (119 out of 124 populations) were sub-sampled to 20 chromosomes (randomly without replacement). We repeated this process ten times, calculating each summary statistic in each replicate, and taking the average over replicates as the final estimate. For each population and subsample, we removed SNPs with more than 10% missing data and a minimum allele frequency (MAF) of less than 10%. For plotting, we computed the mean r2 and the mean distance between pairs of SNPs for all SNP pairs within bins of size 10 Kbp (50 bins in total). The effect of the choices of MAF-cutoff and bin size has previously been shown to have no impact on observed patterns of LD and relative levels of LD [87].

We repeated this procedure for the unmasked and masked AfricanNeo and Only-BSP datasets. In the unmasked dataset, populations have a minimum sample size of 10 individuals and a maximum size of 30 individuals. For the masked dataset, we computed the LD only for populations with a minimum sample size of 7.

To assess whether LD patterns of BSP were consistent with a history of expansion from the homeland in Cameroon/Nigeria, we calculated the correlation between LD and geographic distance from Cameroon, assuming that the Bantu-speaker expansion started there. The geographic distance was calculated as spherical distance with the function *distGeo* of the *geosphere* R package [22]. In Cameroon, the position around the middle of the country (longitud= 11.831477; and latitud= 5.291058) was chosen to calculate the distance.

##### 1.14. Phylogenetic analyses

To investigate phylogenetic relationships between populations included in the AfricanNeo dataset, we used the maximum-likelihood (ML)-based software TreeMix v1.13 [19]. To find the best-supported tree and infer the best number of migration events, we generated a scenario with no migration events and then tested for a range of migration events between 1 and 10. Each proposed TreeMix topology was accessed by bootstrapping blocks of 500 SNPs and assigning a Khoe-San population (Juhoansi) as the root of the population tree [24,88]. ML-based population trees were plotted using MEGA v11 [89].

##### 1.15. Testing models of isolation-by-distance

To investigate patterns of isolation-by-distance (hereafter IBD) between BSP and test four distinct population genetic models, we used SpaceMix software v0.13 [29]. SpaceMix uses geographic information coupled with genetic data to generate so-called “geo-genetic maps”. In these maps, the distance between populations is based on genetic distance rather than geographical distances between studied individuals/populations. The resulting geo-genetic maps are thus analogous to PC projections, in that variations in a 2D plane correspond to genetic variation. In short, the general underlying assumption evaluated with SpaceMix is that under an isolation-by-distance (IBD) pattern, geographic and geo-genetic positions will be similar, which is consistent with a pattern of isolation-by-distance.

We used SpaceMix to test model-testing four different isolation-by-distance models underlying the following population histories:

(*i*) Model A for populations without either migration or admixture (i.e. a full IBD model);

(*ii*) Model B where no migration, but admixture was allowed (i.e. an IBD with admixture model);

(*iii*) Model C where migration was allowed but not admixture (i.e. an IBD with migration model);

(*iv*) Model D where both migration and admixture were allowed (i.e. an IBD with both migration and admixture model).

Each tested isolation-by-distance model was run for $10^6$ iterations and 8 fast runs were performed with $10^5$ generations. The "best" fitting model was evaluated using Pearson correlations between the expected and the observed data. Only the same individuals that were masked and imputed were analysed for the "unmasked" dataset and the "masked" dataset.

##### 1.16. Testing spatially explicit models of the Bantu-speaker expansion

To test different demographic scenarios for the Bantu-speaker expansion, we used a spatiotemporally explicit population genetic framework [30,31]. Briefly, this framework represents the world as a grid of hexagonal cells, approximately 100 km wide. Each cell represents a local, panmictic population with simple population dynamics characterized by colonization of empty cells from occupied neighbors; simple population growth and exchange of migrants between occupied cells; and a maximum population size informed by palaeo-climate reconstructions of net primary productivity [30,31]. In the original version of the model, Eriksson et al. [30] considered only the initial peopling of the world by anatomically modern humans (AMH) 70–50 kya. We here adapted the extension of the model to multiple local expansions as proposed by Raghavan et al. [31] to different scenarios of Bantu-speaker expansions.

Following the initial expansion of AMH within Africa, we assume that the Bantu culture arose in a region of West Africa (**Fig.S 10.1**), from which BSP expanded demically across a wider region of sub-equatorial Africa (**Fig.S 10.1**), experiencing sequential population bottlenecks and potentially mixing with local non-BSP along the way. From this region we excluded areas of dense rainforests motivated by the harsh living conditions and relative unsuitability for farming in these areas. As described in the main text, we considered three demographic scenarios in which the Bantu-speaker expansion proceeded north of the rainforest, south through a rainforest corridor, or using both northern and southern routes. To this end, we modified the basic scenario in two ways, by either (*i*) creating a corridor through the rainforest that allowed expansion south, or (*ii*) blocking expansion north of the rainforest by excluding a section of the Bantu territory (**Fig.S 10.1**). For each demographic scenario, we ran one million simulations with parameters drawn from an independent uniform distribution for parameters characterizing the Bantu-speaker expansion, and from a log-uniform distribution for parameters describing the initial global expansion of AMH taken from Raghavan et al. [31]. The timing of Bantu-speaker culture emergence and onset of the expansion was assumed to be a randomly drawn generation, uniformly distributed in the range from 80 to 400 generations ago (corresponding to 2,000 to 10,000 years ago, respectively). In each simulation, we generated gene genealogies for 500 unlinked loci (using the Wright-Fisher model) for 131 individuals from 14 selected African populations previously published by the H3Africa Consortium [90] (**Table.S 10**), for which we have whole-genome sequencing (WGS) data (mean coverage between 10x and 30x), and calculated $R^2$ between predicted and observed genome-wide differences between pairs of individuals. The simulation framework was implemented in C++ and all the outputs were analyzed using Matlab (9.9.0.1718557 (R2020b) Update 6), and custom scripts. All code and scripts are available from the authors upon request.

##### 1.17. Estimating effective migration rates

To investigate migration rates, we used two approaches. First, we used EEMS software [27]. Analysis was performed with the following settings: diploid= true; numMCMCIter= 2000000; numBurnIter= 1000000; and numThinIter= 9999. The number of individuals, sites and demes varied between the different analyses depending on the dataset and sampling area. For the analysis of BSP, we used the following settings: “mean nIndiv= 4003”; “nSites =191292”, and “nDemes= 200”. Each analysis was repeated three times and an average was taken as input for the visualization as recommended in the EEMS manual. Then, the results were visualized in R v3.6.1 using the accompanying R library rEEMSplots [27]. For the plots of only BSP, the colors were added according to the following linguistic groups: north-western BSP, west-western BSP, south-western BSP, and eastern BSP.

Second, we used a recent implementation of EEMS called Fast Estimation of Effective Migration Surfaces software (FEEMS) [28], to further investigate spatial population structure across Africa. Unlike EEMS, FEEMS applies a Gaussian Markov Random Field (GMRF) model in a penalized-likelihood-based framework to infer whether populations are exchanging gene flow with neighboring populations in a spatial graph of a “stepping-stone” model of first migration later followed by genetic drift.

##### 1.18. Gene-flow barriers analysis on a grid

To further investigate the routes of migration for the Bantu-speaker expansion, we applied the approach from Pagani et al. [26], here called GenGrad. In short, this approach allows a visual representation of spatial genetic barriers inferred from genome-wide genetic distances and displays a gradient of spatially interpolated allele frequencies. As we are investigating a much smaller area than all of Africa, Eurasia, and Oceania, a few adjustments were made to certain parameters. Namely the sigma value was decreased to 0.1 and the color bar was adjusted to only incorporate the measured values. The calculations were performed in MatLab version 9.12.0.1884302 (R2022a), and we used $F_{ST}$ as the distance metric.

##### 1.19. Admixture timing inference

To estimate admixture dates, we applied haplotype-based admixture inference methods. First, we used MOSAIC v1.4 [16] for two- and three-way admixture models on the basis of the phased AfricanNeo dataset. For haplotype phasing the AfricanNeo dataset, we used SHAPEIT v2.r904 [17], and the Haplotype Reference Consortium as a reference panel [78,91]. Recombination maps were interpolated from the HapMap Phase 2 genetic map. To minimize switch error rates [17], we used the following parameters: 500 states, 50 MCMC main steps, 10 burnin and 10 pruning steps.

##### 1.20. Correlations between linguistic, geographical, and genetic data

To test the correlations between matrices of genetic, linguistic, and geographical distances, we performed Mantel tests and partial Mantel tests using the R package ncf [20]. For each test, we computed Pearson's correlation analysis using 100,000 permutations between the two matrices. For the matrix of genetic distances, we computed pairwise $F_{ST}$ between BSP included in the ancestry-masked Only-Bantu dataset using the EIGENSOFT package v7.2.1 [21]. For the matrix of geographic distances, pairwise distances were calculated as pairwise great circle distances (in km) between the studied populations using the R package *geosphere* [22].

To compare genetic and geographic distances between Bantu speech communities with the linguistic distances between their languages, we collected sets of basic vocabulary in 62 different varieties of 44 different Bantu languages. Basic vocabulary is part of a language’s lexicon which is considered the most stable and the least susceptible to borrowing because it bears upon concepts that are universally shared amongst human societies [92]. We compiled vocabulary for 92 such basic concepts, which have also been used in recent phylogenetic studies on BSP [23,93–95]. Most of the lexical data in our dataset have also been used in one or more of these linguistic studies. **Table.S**

**12** provides a comprehensive list of our linguistic dataset including an overview of the original sources from which the linguistic data for each variety originate and the phylogenetic studies in which they have previously been used. It also includes representative geo-coordinates for each sampled Bantu language variety. Cognates are lexical roots originating in one and the same ancestral lexeme which related languages share through inheritance from a common ancestor. We have reviewed and updated all cognacy judgments and extracted a binary-coded root-meaning association matrix using Lexedata [96]. In total, 38 BSP were matching between the linguistic dataset and our imputed Only-BSP dataset, thus each matrix of pairwise linguistic, geographical, and genetic distances were matching the subset of 38 BSP. Our complete lexical dataset including cognacy judgments is freely accessible online (at https://github.com/Schlebusch-lab/Expansion_of_BSP_Suppl_Material). Linguistic data are available in CLDF format [97].

##### 1.21. Comparisons between ancient and present-day populations

We used *smartPCA* to project ancient samples onto a background of present-day populations included in the Only-African dataset, as well as for a sub-dataset including BSP-related aDNA and Only-BSP and selected West-Central African populations (using “YES” option for the following parameters: allsnps, lsqproject, newshrink, and killr2). After merging haplodized modern samples and pseudo-haplodized aDNA samples and LD-pruning steps, we used PLINK to remove variants due to minor allele threshold (plink --maf 0.1) and LD-pruning (plink --indep-pairwise 100 10 0.2). Also, we used unsupervised ADMIXTURE analysis from K=2 to K=12 to infer clusters and ancestry proportions from the modern-day dataset that were then provided as input used to project the aDNA data using the projection approach (admixture “-P” option) of ADMIXTURE software [6,75].

#### 2. Supplementary Notes

##### Note.S 1. Further background on the Bantu-speaker expansion

The Bantu-speaker expansion was a complex pattern of movement of peoples, cultures, and languages spanning several thousands of years, and there are still many aspects and intricacies left to investigate, from the first expansion from the Bantu-speaker homeland until the last expansion into southern Africa. Preliminary suggestions of these complexities are seen in a few recent studies. Semo et al. [61] investigated Bantu language speakers in Mozambique and found a north-to-south cline in a genetic relationship, complemented by a north-to-south gradient of decreasing genetic diversity. This supports a north-to-south dispersal of Bantu-speaking groups along the Indian Ocean coast possibly associated with the Iron Age assemblages of the Kwale archeological tradition [98]. In another fine-scale study, Sengupta et al. [99] studied South African BSP and found a well-defined genetic substructure among southeastern Bantu-speaking populations from South Africa. Speakers of eight of the nine major Bantu languages in South Africa were included in the study and could genetically be distinguished from each other. The genetic substructure correlated to both geography and language. Seidensticker et al. [100] found archeological evidence for several waves of habitation by Bantu speakers in the Central African rainforest and suggested that the Bantu-expansion might have involved several spread-over-spread events [100].

Further evidence of possible spread-over-spread events from genetic data was reported by Sengupta et al. [99] for southeastern Bantu speakers from South Africa. The study reported deeper admixture dates with local San groups in the Tsonga and Venda than in all studied Sotho- and Nguni-speaking groups (including Pedi, Tswana, Southern Sotho, Swazi, Zulu, and Xhosa). This indicates complex events during the settling of Bantu speakers in southern Africa, where the Tsonga and

Venda might be the descendants of earlier waves of migration and the other BSP of later waves. Archeological sources indicate that the ancestors of Nguni speakers migrated to Southern Africa from eastern Africa about one thousand years ago (kya) [101], while the Tsonga is mentioned to be among the earlier Bantu-speaking waves that migrated along the east coast of Africa [102]. Linguistically speaking, however, all Southern Bantu (Guthrie Zone S) languages, including Tsonga, Venda, Nguni and Sotho groups, descend from a most recent common ancestor, which is unique to them within eastern Bantu speakers [95]. Thus, linguistic data is at odds with genetic and archeological inferences and might indicate that the ancestors of Tsonga and Venda shifted to more recently arrived languages (closely related to Sotho and Nguni languages). Alternatively, only part of the Tsonga and Venda ancestry might link to earlier migrations into the area, while the other part of their ancestry might have come with Tsonga and Venda speakers.

#### Note.S 2. Patterns of population structure and genetic diversity

To assess the genetic relationships between BSP, we first performed principal component analysis (PCA) for each assembled dataset. Following linguistic classifications, we grouped BSP into four major linguistic groups: north-western Bantu 2 (in brown), west-western Bantu (in green), south-western Bantu (in dark blue), and eastern Bantu speakers (in red) (**Fig.S 1.4a** and **1.5a–b**). In this classification, we also included as south-western BSP the Damara-KSP from Namibia noting that this population speak a Khoe-Kwadi language, and we excluded BSP such as the Baka sampled in Cameroon and Gabon because they were included in the western rainforest hunter-gatherer group (wRHG), following indications from previous studies [65,73,103].

To visualize genetic variation in the African continent, we applied four dimensionality reduction methods for SNP data. First, we used the uniform manifold approximation and projection (UMAP) method directly on the genotype data [2]. UMAP results highlight separate groups for African populations from western and eastern hunter-gatherer groups (e.g. Baka, Hadza, and Sabue) and Nilo-Saharan-speaking populations (e.g. Gumuz) and East Asian populations (**Fig.S 2.1a–b**). After zooming in on the results, we observed a good similarity between the position of the populations and their geographical distribution (**Fig.S 2.1c–d**). We then applied the UMAP approach to a sub-dataset after removing the outlier populations from **Fig.S 2.1a–b** (e.g., Gumuz, Hadza, Sabue, Baka, and East Asian populations). In the UMAP plot of selected populations (**Fig. 1b**), we also observed patterns of population structure and admixture. For instance, in the UMAP plot Fula individuals from Gambia are between western African populations and Eurasian populations, as previously reported [104–107]. In addition, we observed complex population structure among Nilo-Saharan-speaking populations from the north and south regions of Chad, as previously reported [69,106,107]. Interestingly, populations from southern Chad [69] might have admixture with Ubangi-speaking populations from CAR from this study.

PCA results of populations included in the Only-BSP dataset evidenced separate major Bantu-speaking groups in our dataset (**Fig.S 2.2**). We then selected sub-Saharan African populations from the African-Neo dataset. PCA results for selected populations (**Fig.S 2.3**) better highlight a strong genetic differentiation between Khoe-San-speakers and other African populations on the first principal component (PC1), between Afro-Asiatic-speaking populations and other African populations on PC2, between wRHG populations and the other African groups on PC3 and PC4, and between BSP from Namibia (Himba and Herrero) and the other African groups on PC6. Interestingly, new Ubangi-speaking populations from CAR included in the AfricanNeo dataset evidence genetic affinities with Afro-Asiatic-speaking populations and Nilo-Saharan-speaking populations from Chad and the southern region of Sudan (**Fig.S 2.3**). After rotating according to geographic sampling through Procrustes projection, the PCA for all sub-Saharan African populations better highlights the genetic diversity and differentiation among sub-Saharan African groups that have different linguistic backgrounds, lifestyles, or geographical distribution (**Fig. 1c** and **Fig.S 2.4**). We also performed PCA for all newly genotyped samples from Africa that were included in the Full-Genotyped dataset (**Table.S 1**) plus reference sub-Saharan African populations included in the AfricanNeo dataset (**Table.S 2**) after merging and quality control (**Fig.S 2.5**). PCA results of all available sub-Saharan African populations further highlight the observed patterns described before.

To investigate the genetic relationships between BSP and worldwide populations, we then performed PCA based on the the AfricanNeo dataset (**Fig.S 2.6**). As expected, the PCA plot for all the populations included in the AfricanNeo dataset highlights continental ancestries of African versus Eurasian populations on each side of PC1, and between European and East Asian populations on PC2, and individuals from each major Bantu-speaking group are close in the multidimensional space (**Fig.S 2.6**). On PC3, we observed the split of Khoe-San populations from southern Africa. We then used the UMAP algorithm to combine the information of the first 10 PCs of the PCA estimated for the AfricanNeo dataset (**Fig.S 2.6**). PCA-UMAP results (**Fig.S 2.7**) showed Bantu-speaking populations grouping together, and splitting away from other African populations that were separated into different groups in the multidimensional space, except for BSP which seem to have a continuous connection that is in agreement with the geographical locations of the populations (see the zoomed region in **Fig.S 2.7c–d**).

Lastly, we applied a deep learning framework for dimensionality reduction based on genotype convolutional autoencoder (GCAE) [5]. GCAE results further support the clustering of populations at the continental level between all studied worldwide populations included in the AfricanNeo database, and also some degree of differentiation among sub-Saharan African populations (**Fig.S 2.8a–b**), notably different than the lack of differentiation between African populations on PC1 in **Fig.S 2.6a**. In addition, we also observed a better split of populations at the sub-continental level in Africa for the GCAE analysis on the basis of sub-Saharan African populations (**Fig.S 2.8c–d**). GCAE results highlighted patterns of admixture and population structure that further supported our previous results. In general, dimensionality reduction methods, such as UMAP, PCA-UMAP, and GCAE, are shown to be able to capture finer population substructure among African populations than the first two projections of the PCA alone. Taken altogether, in the multidimensional space of all these results we observed genetic differentiation between and within Bantu-speaking groups. In addition, there are different genetic patterns among eastern BSP who are geographically distant but who belong to the same linguistic group. Among northeastern BSP we observed the highest amounts of Afro-Asiatic-related admixture, while among southeastern BSP we observed the highest amounts of Khoe-San-related admixture. North-western, west-western, and south-western BSP were more similar genetically, except for three populations: the Himba and Herero populations, which have undergone more genetic drift, and the Damara-KSP, which has individuals with higher proportions of Khoe-San-related admixture (**Fig.S 2.3c**).

To further assess the human genetic landscape in Africa, we investigated the genetic diversity across worldwide populations using unsupervised ADMIXTURE analyses. First, we performed ADMIXTURE analyses at K=4 for the worldwide populations included in the AfricanNeo dataset (**Fig.S 3.6**, and **3.13**). We observed population substructure among sub-Saharan African populations, in agreement with previous research [64,104,108]. Eastern BSP had indications of admixture with eastern African populations (brown component; **Fig.S 3.2f**), Middle Eastern populations (gray component; **Fig.S 3.2f**), and eastern hunter-gatherer groups in several regions (black component; **Fig.S 3.2i**). We also observed indications of admixture between South African Coloured groups and Eurasian populations (dark-green component; **Fig.S 3.13**), as previously reported [24,68,109].

We then performed ADMIXTURE analyses from K=2 to K=25 on the basis of the AfricanNeo dataset. To better visualize patterns of population structure, we plotted ADMIXTURE results for selected K-groups using pie charts (for K=2, K=4, K=6, K=12, and K=16; **Fig.S 3.5–3.11**), ternary diagrams (for K=6; **Fig.S 3.12**), and bar plots (for K=4 and K=16; **Fig.S 3.11–3.12**). ADMIXTURE results at K=2 and K=4 (**Table.S 3**) further indicated possible gene-flow from Eurasian ancestry into eastern and southern African populations (**Fig.S 3.5–3.6**), in particular in eastern African populations such as Swahili-speakers from eastern Kenya [63] and in the South African Coloured groups [24,109]. In ternary diagrams of ADMIXTURE results at K=6 (**Fig.S 3.12**), we also observed a genetic cline between Niger-Congo-speaking populations from western to west-central African countries. ADMIXTURE results at K=12 (**Fig.S 3.1–3.3**) better highlight different components for hunter-gatherer populations in western Africa (yellow component), eastern Africa (black component), and southern Africa (purple component) (**Fig.S 3.8**). Those components are also present in BSP from different regions with different proportions of possible admixture.

At K=16, the K-group with the lowest cross-validation (CV) value (**Fig.S 3.4**), we observed different clusters between and within groups of western and eastern BSP (e.g. cyan, green and orange components in **Fig.S 3.14**), in agreement with their genetic cluster sharing with other sub-Saharan African groups (**Table.S 4**, **Fig. 2b, Fig.S 3.9**, **3.11** and **3.14**). We formally tested the hypothesis of admixture and its regional character using f3- and f4-statistics (**Fig.S 3.16–3.19**). We detected evidence of admixture in some BSP with Afro-Asiatic speaking groups (**Fig.S 3.16**), western RHG groups (**Fig.S 3.17** and **3.19**), or Khoe-San groups (**Fig.S 3.18** and **3.19**).

#### Note.S 3. Ancestry-specific analyses in Bantu-speaking populations

To eliminate the influence of admixture on our analyses of BSP, we only analyzed haplotype segments with West-Central African-related (WCA) ancestry previously estimated using a local ancestry inference approach. Ancestry-specific (AS-)PCA results based on the masked, phased, and imputed dataset of Bantu-speaking individuals with haplotypes with at least 70% WCA ancestry (thereafter “masked Only-BSP” dataset) together with the unmasked dataset of reference panels evidence the null influence of admixture patterns in BSP that were grouped in all PC projections (**Fig.S 4.1**). AS-PCA results on the basis of the Only-BSP dataset evidenced fine population structure among BSP, with Himba and Herero populations from Namibia separate from other studied BSP on PC1 and PC2, while PC3 split northwestern BSP and western BSP (**Fig.S 4.1a–b**). After removing Himba and Herrero individuals from the analysis, AS-PCA better highlighted the geographical distribution of BSP in the multidimensional space without the influence of the admixture (**Fig.S 4.2**). Interestingly, after masking admixture in studied BSP, the Himba and Herrero populations present the highest values of ROH-based genomic inbreeding coefficient (on average $F_{ROH}$= 0.02), notably different from other BSP (**Fig.S 4.3**).

#### Note.S 4. Patterns of consanguinity and founder events

To shed additional light on the demographic history and cultural practices of BSP, we analyzed eleven patterns of runs of homozygosity (ROH) (**Table.S 6** and **Fig.S 5.1–5.12**). In general, BSP have low values of the total sum of short ROH (**Fig.S 5.1**). The kurtosis and skewness of the violin plots also provide additional information. BSP are relatively homogeneous with very short tails and an almost normal distribution, while other African groups like Afro-Asiatic-speaking populations and Khoe-San-speaking populations, present other types of shapes. For long ROH segments (**Fig.S 5.2**), the highest values among all studied worldwide populations were detected in the Hadza population in Tanzania and the Sabue population in Ethiopia (on average: 0.48; **Table.S 6**). Among BSP, the Himba and Herero populations in Namibia have the highest values of ROH-based inbreeding coefficient ($F_{ROH}$= 0.021 ±0.012SD and 0.015 ±0.007SD, respectively; see **Table.S 6**, **Fig.S 5.5**), highlighting genetic isolation in this population likely after the Bantu-speaker expansion to southwestern Africa. We also analyzed in each population their average for six ROH length categories (**Fig.S 5.6–5.12**). Interestingly, the highest averages among BSP were detected in the Himba and Herero populations for categories 4 and 5 of ROH length (**Fig.S 5.10–5.11**). In agreement with previous studies [110,111], those populations have the highest values of the mean long ROH (**Fig.S 5.3**), total length of long ROH (**Fig.S 5.4**), and genomic inbreeding coefficient (**Fig.S 5.5**), highlighting patterns of strong genetic isolation or genetic drift.

##### Note.S 5. Effective population size and founder events in BSP

To infer population changes over the last millennium, we estimated effective population sizes (Ne) for the last 50 generations using IBDNe (ibdne.19Sep19.268.jar) [14]. In general, IBDNe results highlight population expansions in the last 10 generations for all the studied populations (**Fig.S 6.1–6.2**). As expected, some hunter-gatherer populations also experienced demographic bottlenecks, in particular the Sabue population from Ethiopia [111]. Among BSP, we detected different patterns between populations from different countries or within populations from the same country (**Fig.S 6.2**).

Founder events were investigated using ASCEND v10.0 [15]. Following the four criteria recommended by Tournebize et al. [15], among BSP the results evidenced significant founder events in 19 populations in Namibia (Himba and Herero), Eswatini (Swazi), Zambia (Fwe), Tanzania (Swahili population in Zanzibar), Botswana (Ghanzi), South Africa (Bhaca, SEBantu, Xhosa, Zulu, and ZuluAGDP), and Mozambique (Bitonga, Chopi, Makhuwa, Ndau, Nyanja, Tewe, Tswa, and Yao) (**Table.S 7**). Among BSP, the highest intensity was found for the founder event in the Himba ($I_f$=1.6%, 95% CI:1.5–1.7%; $T_f$= 21 generations, 95% CI: 18.7–24.1) and Herero populations ($I_f$=1.2%, 95% CI:1.1–1.3%; $T_f$= 29 generations, 95% CI: 25.6–31.6) (**Fig.S 6.3** and **Table.S 7**), suggesting that those populations have been more genetically isolated from other groups (**Fig.S 6.4**). Among the BSP with significant founder events, the oldest date was estimated in the Chopi population from Mozambique ($T_f$= 87 generations; 95% CI: 73–100.5), followed by other populations in Mozambique (**Table.S 7**).

##### Note.S 6. Patterns of isolation-by-distance among BSP

One of the simplest models for relationships between populations is the isolation-by-distance model (IBD in this section). Under an IBD model, populations close to each other are genetically more similar than populations far apart and this differentiation should increase with greater geographic distance. To investigate these patterns we used the software SpaceMix [29] on both the unmasked and masked Only-BSP datasets. Four models are tested: (a) No migration and no admixture; (b) No migration but admixture; (c) Migration but no admixture; and (d) Both migration and admixture. For all four models, including the IBD model, there was a high correlation between the observed data and the data estimated from the model (**Fig.S 7.4** and **Fig.S 7.8**). In the Only-BSP dataset, we see a strong adherence to IBD patterns although the model with admixture (b) or migration (c) has slightly higher correlations (**Fig.S 7.4**). In the dataset where we masked out the admixed parts of the genomes of BSP (**Fig.S 7.8**) the IBD model receives the highest likelihood. The Herero population is outside the plot area in for instance **Fig.S 7.6c** but can be seen in the wider plot in **Fig.S 7.7c**. This is likely due to the demographic bottleneck observed in this population (**Fig.S 6.3**).

In addition, SpaceMix analyses suggested patterns of IBD within the BSP in our dataset, even when masking out admixture in BSP. Previous genetic studies have indicated that the Khoe-San populations of Southern Africa follow a very strong pattern of IBD [62] and that these IBD patterns extend to hunter-gatherer populations across the African continent [112]. It might thus be suggested that IBD patterns observed within BSP can be influenced or even explained by admixture with local hunter-gatherer groups. SpaceMix results however indicate that IBD patterns between Bantu-speaking groups remain and even become stronger when admixed parts of the genome were removed after masking and only the Bantu-speaking-related component was considered.

#### Note.S 7. Patterns of haplotype diversity and linkage disequilibrium among BSP

To further investigate genetic diversity within our dataset, we calculated haplotype richness (HR) and haplotype heterozygosity (HH) following recommendations from Schlebusch et al. [24]. These statistics were calculated for increasing window sizes starting at 2000 bp and increasing by 1000 up to a window size of 100,000. As previously reported [24], sub-Saharan African populations are the most genetically diverse human populations in the world (**Fig.S 8.1–8.3**). Among BSP, HH and HR followed a pattern of decreasing diversity along the suggested paths of the Bantu-speaker expansions, with increases in diversity in regions with high patterns of genetic admixture, such as South African BSP and around the Central African BSP (**Fig.S 8.9**). South African BSP generally have a high level of Khoe-San admixture, while RHG admixture is present in BSP close to the Congo rainforest in Central Africa.

Population structure and genetic drift have large effects on these diversity metrics, as is apparent when looking at the Himba population of Namibia (**Fig.S 8.10**). Furthermore, the masking, phasing and imputation approach to individually analyze west-central African-related haplotypes comes at the cost of a lack of haplotype diversity (**Fig.S 8.9**), as the admixed haplotypes are replaced with West African ones.

To further examine the genetic patterns of the Bantu-speaker expansion, we correlated these statistics with the distance from the Bantu-speaker homeland (**Fig.S 8.10–8.11**). We expect that these metrics would generally decrease the further a population is from the Bantu-speaker homeland as genetic variation is lost along with the expansion/migration i.e. a serial-founder effect. For the haplotype heterozygosity metric estimated on the basis of the Only-BSP dataset without masking and imputation, we observe a correlation with the distance from Cameroon ($R^2$= 0.126, *P*-value= 0.0006; **Fig.S 8.11a**), as well as for the masked and imputed Only-BSP dataset ($R^2$= 0.165, *P*-value= 0.0040; **Fig.S 8.11b**). In agreement, for the haplotype richness metric, we also observed a correlation ($R^2$= 0.136; *P*-value= 0.0002; **Fig.S 8.10a**) for the original data, and a correlation of 0.267 (*P*-value= 0.0002; **Fig.S 8.10b**) for the masked and imputed Only-BSP dataset. The predicted genetic pattern is in general observed in BSP with high admixture with hunter-gatherer groups (higher than expected) or genetic drift (lower than expected values). Masking out admixture thus increases the correlation between distance from the Bantu-speaker homeland and a decrease in genetic variation.

In general, short-range LD patterns provide information on ancient events (long-term $N_e$ and LD) [113]. When looking at short-range LD in the unmasked dataset (**Fig.S 8.4**), East Asian populations have the highest values, followed by European and Middle-Eastern populations and lastly African populations. Among sub-Saharan African populations, South African Khoe-San populations showed the lowest LD, as previously shown [24]. Long-range LD (up to 500 Kbp) reflects more recent events. When looking at long-range LD, a different pattern was visible in some studied populations. Interestingly, some sub-Saharan African populations (e.g. Sabue in Ethiopia and Hadza in Tanzania) have long-range LD slightly higher than European populations. Sabue and Hadza have notably higher LD (**Fig.S 10.11**), due to their recent population size decrease (**Fig.S 6.1**) and their observed high genomic inbreeding (**Fig.S 5.5**).

An LD gradient was observed among BSP (**Fig.S 8.4a** and **8.12a**), in which South African BSP had the lowest short-range LD and BSP in Mozambique had the highest one. The admixture with hunter-gatherer groups could account for this pattern in South Africa, while BSP in Mozambique show very small evidence of RHG admixture, in agreement with a previous study [61]. To evaluate the effect of admixture on LD patterns in BSP, LD calculations were performed on the masked Only-BSP dataset. We selected BSP with more than 70% of West-Central African-related ancestry and plotted them together with six reference populations (**Fig.S 8.4b** and **8.12b**). After masking, LD was higher in several BSP (max value= 0.275 in the Pedi population from South Africa, while 0.155 was estimated in the unmasked dataset). Also, we see a spatial pattern of LD increasing from west-central Africa towards the east and the south (**Fig.S 8.12b**), which is more coherent with a history of expansion from their homeland between the border of Nigeria and Cameroon than the patterns observed in the unmasked dataset (**Fig.S 8.12a**). Assuming that Bantu-speakers started their migration in this region, we would expect a genome-wide increase of LD in BSP situated farther away from the origin of the expansion in Cameroon. For that reason, we estimated the correlations between LD patterns and geographic distances in BSP (**Fig.S 8.13**). The correlation was positive and significant. LD at 50 Kb increased with spatial distance from Cameroon in BSP (**Fig.S 8.14a**; adjusted R-squared: 0.2258; *P*-value= 0.00002). The correlation is still significant, albeit less strong, for the masked dataset (**Fig.S 8.14b**; adjusted R-squared= 0.0946; *P*-value= 0.0180). The difference between the unmasked and the masked dataset seems to be mainly due to Mozambican populations, whose LD decreases with imputation, and South African BSP, largely absent from the masked dataset because of their small sample sizes. BSP in Mozambique showed higher values of homozygosity than populations from nearby regions (visible in short ROH length **Fig.S 5.1** and **Table.S 6**), so it is reasonable to think that the imputation introduced some diversity disrupting the range of LD. By comparison, the correlation was not significant when all African populations (N= 111) were included in the analysis (adjusted R-squared= 0.0039; *P*-value= 0.2329).

#### Note.S 8. Phylogenetic analyses of Bantu-speaking populations

To investigate patterns of population splits and migration events among BSP, we used TreeMix-based phylogenetic trees [19]. For the AfricanNeo dataset, African and Eurasian populations split into separate branches in the phylogenetic tree, Khoe-San populations root the tree, and sub-Saharan African populations separate into different branches in agreement with their geographical distributions, lifestyles and linguistic backgrounds (**Fig.S 9.1–9.2**). For the unmasked Only-BSP dataset (**Fig.S 9.3**), northwestern BSP were at the base of the phylogenetic tree, while western and eastern BSP split from each other. There are two main branches among western BSP, the northwestern BSP (in DRC) and the southwestern BSP, as well as among eastern BSP, the northeastern BSP and the southeastern BSP. Therefore, the separation of BSP was in agreement with their observed admixture patterns with other African groups described above, and also with the linguistic classification of BSP [114]. For the masked Only-BSP dataset (**Fig.S 9.3**), eastern BSP were together in the same branch without the subdivision due to admixture patterns with other African groups.

#### Note.S 9. Patterns of pairwise genetic distances and admixture graphs

For the masked Only-BSP dataset, $F_{ST}$ values evidenced the lowest values among BSP from West-Central Africa to the rest of sub-Saharan Africa (**Fig.S 10.2**). We used the lowest $F_{ST}$ values between neighboring BSP to infer the putative migration routes during the Bantu-speaker expansion (**Fig. 5a** and **Fig.S 10.3a–b**). To do so, for each African country we selected one BSP that is most (geographically) distant from the BSP in Cameroon and by depicting one arrow from the target population to the BSP with the lowest pairwise $F_{ST}$ value in our dataset (**Fig.S 10.3**). After finding the direction of the first arrow, we continue the analysis using as a target the destination of the arrow and excluding for the new analysis the previous target population. For instance, we started with the Zulu population from South Africa, and the lowest pairwise $F_{ST}$ value of this BSP was with the Tsonga population from South Africa, and for the Tsonga population the lowest value was with the Lozi population from Zambia (after excluding the Zulu for the analysis), and we repeat the same procedure until reaching the Nzime population in Cameroon. Using this approach repeatedly, we depicted the arrows connecting the BSP that are more distant from the Bantu-speaker homeland, and we reconstructed the putative migration routes (**Fig. 5a** and **Fig.S 10.3a**). We evaluated these migration routes using qpGraph to build an admixture graph of the putative routes (**Fig.S 10.3c**). Bantu-speaking populations from western DRC seem to be the starting point of the migration route of BSP to southern Africa, in particular the Lozi population from Zambia and from the Lozi to South Africa, Mozambique and Zimbabwe (**Fig.S 10.3a**). We took this exploratory approach with caution, and we repeated the analysis after removing the Lozi population from Zambia due to the wide distribution of this population in Zambia (**Fig.S 10.3b**). With or without the Lozi (**Fig.S 10.3a** and **10.3b**, respectively), in both analyses the results showed one migration wave from Cameroon to western DRC, and different migration waves from western DRC to eastern and southern BSP. The connection between BSP from western DRC and southwestern BSP was also observed, suggesting that western DRC is likely to have been an important crossroad region for the expansion of BSP to most of the sub-equatorial African regions.

#### Note.S 10. Estimation of effective migration surfaces

To further investigate migration rates over geographic space, we applied EEMS and FEEMS [27,28] (**Fig.S 10.4–10.10**). In EEMS analyses with the masked Only-Africa dataset (**Fig. 5c**), low effective migration rates were observed at geographical barriers such as the eastern African Great Lakes regions, the southern African Kalahari desert, and the west-central African rainforest (**Fig.S 10.4–10.5**). These areas also coincide with locations where groups with very different genetic ancestries co-exist with BSP. High genetic differentiation between BSP and non-Bantu-speaking groups likely drives the patterns of low migration rates in these regions. EEMS analysis was also carried out on the unmasked and masked Only-BSP dataset (**Fig.S 10.6–10.7**). The unmasked Only-BSP dataset recapitulates the findings of the masked Only-Africa dataset, where admixture within Bantu-speakers from genetically distant neighboring groups is likely associated with inferred low effective migration rates. In the masked Only-BSP dataset (**Fig. 5c**), patterns of relationships restricted to the BSP genetic component are highlighted. There are regions of high migration rates along the Indian Ocean coast from Kenya to eastern South Africa. In contrast, we identify a region of low migration in Tanzania between three eastern African lakes (Lake Victoria, Lake Tanganyika, and Lake Malawi). Another corridor of low migration stretches up from South Africa and Zimbabwe, through Zambia and the DRC, possibly associated with a separation between western and eastern Bantu speakers. In Zambia, the barrier is between the western and the eastern regions and in between the western and eastern branches of the Bantu languages (**Fig. 5c**).

Taken together, EEMS results identified an eastern coastal path as a region of high migration rates for the Bantu-speaker expansions, together with longitudinal corridors of lower migration rates in the central parts of the continent.

#### Note.S 11. Admixture dating and model-testing of the Bantu-speaker expansion

To estimate the population sources and admixture dates in BSP, we used two approaches. First, we used MOSAIC [16]. Admixture testing using MOSAIC inferred the dates of admixture events between BSP and local populations from different regions (**Fig. 2b, Fig.S 11.1**, and **Table.S 8**). As suggested by previous results, southwestern BSP have admixture with local populations of Khoe-San-related ancestry (ranging from 11 to 20 generations ago). Northeastern BSP have higher amounts of admixture with populations of Afro-Asiatic-speaking related ancestry than southeastern BSP (**Fig.S 11.1**). In South Africa, southeastern BSP have admixture with populations of Khoe-San-related ancestry and the admixture dates are more recent (range: 20-24) than the dates inferred in BSP from Mozambique (range: 28-41) with low amounts of gene-flow with eastern African populations. Admixture dates correlate significantly with geographic distance from Cameroon ($R^2$=0.13, *P*-value= 2.6e-05) with earlier dates in the Bantu-core region and more recent dates toward the extremes of the expansion (**Fig. 2b** and **Fig.S 11.2**).

#### Note.S 12. Comparisons between ancient and present-day populations in Africa

We produced whole-genome sequencing data of 12 ancient samples from six archeological sites in South Africa and three in Zambia (**Table.S 11**). The age of the samples spanned the last 800 years according to C14 carbon-dating (**Table.S 11**; [35,36]). Ancient DNA sequences have a high frequency of cytosine to thymine (C to T) transitions at the 5' ends and of guanidine to adenine (G to A) at 3' ends due to post-mortem deamination. These typical damage patterns increase with the age of the sample and its conservation status. The nucleotide misincorporation patterns of our samples are shown in **Fig. S12.0** and **Table.S 11**. In general, the damage patterns of our samples are consistent with their age, with seven out of 12 samples showing damage patterns higher than 10%. Mitochondrial contamination estimates were low for all samples, ranging from 0 to 0.03 (**Table.S 11**), and showed no relation with damage patterns. Coverage ranged from 0.02X to 1X for the autosomal genome and 0.8 to 71X for the mitochondrial genome. For all but one sample, we were able to retrieve the mitochondrial haplogroups, which all belonged to the African lineages L0a, L0d, L1c, L2a, L3b and L3e (**Table.S 11**). All the identified mtDNA-haplogroups are haplogroups associated with Bantu-speakers except the L0d haplogroup [115]. L0d is a Khoe-San associated haplogroup [116] and was carried by three out of the six ancient individuals from current-day South Africa. The presence of this haplogroup in the Iron Age individuals is likely due to Khoe-San admixture, as was observed in previous ancient DNA studies [8] and studies on modern-day groups [115,116].

To compare the autosomal genetic diversity of ancient and present-day BSP, we merged the AfricanNeo dataset with 83 ancient DNA (aDNA) individuals from previous studies in Africa [8–13] (**Table.S 5**). Both, PCA and PCA-UMAP results evidenced genetic affinities between aDNA and modern African samples from the same geographic location (**Fig. 1d, Fig.S 12.1–12.5**), suggesting similar patterns of shared ancestry between samples from different time periods. Among BSP (**Fig.S 4.3**), we also observed population structure in DRC between the ancient samples from western DRC with more Niger-Congo-speaking-related ancestry (in the two Kindoki samples from 150-230BP) and the eastern DRC sample with more eastern African-related ancestry (Matangai Turu from 750BP), as previously reported [9]. ADMIXTURE results at K=4 (**Fig.S 12.4**) and K=12 (**Fig.S 12.5**) for modern-day populations and aDNA individuals evidence complex patterns of admixture between Niger-Congo-speaking populations and ancient Pastoral populations in Kenya [10,11]. In southeastern Africa, Iron Age aDNA samples and modern-day BSP provide evidence for genetic continuity in this region (**Fig.S 12.13**), as previously reported [99].

In our newly reported ancient individuals, we observe different cluster assignment profiles for the samples originating from current-day South Africa and current-day Zambia (**Fig.S 12.13**). The South African samples are more homogenous and contain more of the "green" component maximized in modern-day southeast Bantu-speakers from South Africa (SouthAfrica_SEBantu). Their cluster assignments are also similar to previously reported Iron Age individuals from the region, including Mfongosi, ElandCave, Newcastle and ChampagneCastle [8]. The four previously published and six new Iron Age samples from current-day South Africa also show more indications of Khoe-San admixture (gray component) compared to the six Zambia Iron Age samples. Compared to the South African Iron Age samples, the Zambian samples seem to have a more heterogeneous cluster assignment and they contain more of the "dark blue" component maximized in the BSP Shi population from the eastern DRC as well as the "dark yellow" component maximized in the Niger Congo speaking Esan population from Nigeria.

To quantify their affinity to present-day Bantu-speaking populations, we estimated the f3-statistics for all our 12 new ancient samples (**Fig.S 12.6–12.9**). Among the aDNA samples from current-day South Africa, five out of six aDNA samples showed the two highest affinity values for the BSP from Ghanzi in Botswana and the Sotho population from South Africa, while one aDNA sample (WUD038b) had its highest affinity for the South African Sotho and Bhaca populations. For the six samples from current-day South Africa the top 4-6 highest affinity values were always to a current-day South African population or the Botswana Ghanzi population. The ethnolinguistic affiliation of the Ghanzi population is not available - however they were likely Sotho-Tswana speakers, closely related to Sotho speakers from South Africa.

Samples originating from modern-day Zambia, instead, showed more heterogeneous affinities, consistent with the notion that the region was a crossroad of interactions. For the six samples, two (WUD008, WUD0012) showed the strongest affinities with Swahili from Kenya, and the other four with the Chopi population from Mozambique (WUD003), Tonga- and Nyengo populations from Zambia (WUD004 and WUD018), and the Sotho population from South Africa (WUD010). The top 4-6 populations with highest affinity to the six samples originating from current-day Zambia also vary substantially more in terms of geographic location, relative to the results for the six samples from current-day South Africa.

We reported morphological analyses, C14-dating and dietary isotope analyses before for the four samples originating from Mumbwa Cave in Zambia, WUD003, WUD004, WUD008, WUD010 [36]. For WUD003, WUD004 and WUD010 (museum IDs A339, A340 and A346) the morphological sex could not be determined. Their biological sex based on DNA analyses were all XY. WUD008 (A344) was predicted to be male based on morphology and this was confirmed by the DNA analyses (**Table.S 11**). The dietary isotope analyses found that most of the individuals from Mumbwa had similar diets to recent southern African farmers who depended largely on C4 crops and/or plant foods with relatively limited dairy and meat supplements. The WUD008 (A344) dietary isotope profile was different from the other individuals in the study in that his d13C values indicated that he had a higher C3 plant intake, more similar to inland hunter-gatherer groups from southern Africa. His d13N values however were not as high as hunter-gatherer groups[36]. Our genetic analyses confirmed that this individual is genetically associated with Bantu-speakers and not with hunter-gatherers. Within the heterogeneous profiles from the Zambian ancient individuals his cluster assignment profile (**Fig.S 12.13**) shows more of the eastern African BSP associated “blue” and west African associated “dark yellow” component and less of the southeast BSP associated “green” component. His f3 statistics (**Fig.S 12.8**) associate him with Swahili populations from Kenya.

We also reported C14 dates and archeological descriptions for two ancient individuals from current-day South Africa (WUD034 - museum ID A294 and WUD037 - museum ID A302) [35]. The A294 individual was partially mummified and buried in a cave, Kaybar’s Cave, near Cathkin Peak in the Drakensberg. Morphological analyses on pelvis bones indicated a male individual and this is confirmed by the DNA analyses. The A302 individual was buried in Robinson’s Shelter 2, not far from Kaybar’s Cave and composed a near complete skeleton. This individual was also classified as male based on morphological analyses, which is confirmed here by DNA analyses. The details around the two burials are fully described in [35]. The two individuals had similar cluster assignment profiles (**Fig.S 12.13**) that show a large proportion of southeast BSP associated “green” component. Their f3 statistics (**Fig.S 12.8**) associate them with the South African Sotho population and the Botswana Ghanzi group.
