## Supplementary Figures for "The genetic legacy of the expansion of Bantu-speaking peoples in Africa"

**Table of Contents 1**
**1- Extended introduction and geographical locations of studied populations 4**
Fig.S 1.1 | Linguistic hypotheses proposed to explain the expansion of BSP. 4
Fig.S 1.2 | Geographical locations and sample sizes of African populations. 5
Fig.S 1.3 | Geographical locations of African and Eurasian populations. 5
Fig.S 1.4 | Labels of all the groups and populations included in the AfricanNeo dataset. 7
Fig.S 1.5 | Labels of the populations included in the Only-Africa and Only-BSP datasets. 8
Fig.S 1.6 | Workflow for the assembled datasets. 10
**2- Dimensionality reduction methods 11**
Fig.S 2.1 | UMAP approach on the basis of genotype data. 11
Fig.S 2.2 | PCA plots for only Bantu-speaking populations. 12
Fig.S 2.3 | PCA plots for selected sub-Saharan African populations. 13
Fig.S 2.4 | Procrustes rotated PCA for the Only-African dataset. 15
Fig.S 2.5 | PCA for the Full-Genotyped dataset and reference sub-Saharan African populations. 16
Fig.S 2.6 | PCA for each group included in the AfricanNeo dataset. 17
Fig.S 2.7 | PCA-UMAP approach for the AfricanNeo dataset. 18
Fig.S 2.8 | GCAE approach for the AfricanNeo and Only-African datasets. 19
**3- Unsupervised clustering analyses and F-statistics analyses 20**
Fig.S 3.1 | Surfer map of ADMIXTURE at K=12. 20
Fig.S 3.2 | Surfer map of each ADMIXTURE result at K=12. 22
Fig.S 3.3 | Bar plots of ADMIXTURE results for K=12. 23
Fig.S 3.4 | Cross-validation test from the ADMIXTURE analyses for each K-group. 24
Fig.S 3.5 | Pie charts of ADMIXTURE results for K=2. 25
Fig.S 3.6 | Pie charts of ADMIXTURE results for K=4. 26
Fig.S 3.7 | Pie charts of ADMIXTURE results for K=6. 27
Fig.S 3.8 | Pie charts of ADMIXTURE results for K=12. 28
Fig.S 3.9 | Pie charts of ADMIXTURE results for K=16. 29
Fig.S 3.10 | Pie charts of ADMIXTURE results for only studied BSP. 30
Fig.S 3.11 | Pie charts of ADMIXTURE results for K=16 only for BSP. 31
Fig.S 3.12 | Ternary diagram of ADMIXTURE results at K=6. 33
Fig.S 3.13 | Bar plots of ADMIXTURE results for K=4. 34
Fig.S 3.14 | Bar plots of ADMIXTURE results for K=16. 35
Fig.S 3.15 | Pie charts of ADMIXTURE results for K=16 only for DRC and Zambia. 36
Fig.S 3.16 | Test of Afro-Asiatic admixture in BSP estimated using f3- and f4-statistics. 37
Fig.S 3.17 | Test of western RGH admixture in BSP estimated using f3- and f4-statistics. 38
Fig.S 3.18 | Test of Khoe-San admixture in BSP estimated using f3- and f4-statistics. 39
Fig.S 3.19 | Tests of hunter-gatherer admixture in BSP estimated using f3- and f4-statistics. 40
**4- Ancestry-specific analyses 41**
Fig.S 4.1 | Ancestry-specific (AS-)PCA for the masked Only-BSP dataset. 41
Fig.S 4.2 | Ancestry-specific (AS-)-PCA after removing Himba and Herero. 43

Fig.S 4.3 | FROH after masking the AfricanNeo dataset. 44
**5- Genome-wide runs of homozygosity (ROH) estimates 45**
Fig.S 5.1 | Total sum of short ROH length for the AfricanNeo dataset. 45
Fig.S 5.2 | Total sum of long ROH length for the AfricanNeo dataset. 46
Fig.S 5.3 | Mean of long ROH for the AfricanNeo dataset. 47
Fig.S 5.4 | Total length of long ROH for the AfricanNeo dataset. 48
Fig.S 5.5 | Genomic inbreeding coefficient (FROH) for the AfricanNeo dataset. 49
Fig.S 5.6a | All categories of ROH length for the AfricanNeo dataset. 50
Fig.S 5.6b | All categories of ROH length for the Only-BSP dataset. 51
From Fig.S 5.7 to Fig.S 5.12 | Violin plots for six categories of ROH length. 52
**6- Estimated effective population sizes and demographic founder events 58**
Fig.S 6.1 | IBDNe results for the AfricanNeo dataset. 58
Fig.S 6.2 | IBDNe results for BSP from six African regions. 59
Fig.S 6.3 | Intensity of founder events in sub-Saharan African populations. 60
Fig.S 6.4 | Timing of founder ages in sub-Saharan African populations. 61
**7- Patterns of isolation-by-distance for the unmasked and masked datasets 62**
Fig.S 7.1 | SpaceMix results with no population text overlays. 62
Fig.S 7.2 | SpaceMix results with text instead of dots. 63
Fig.S 7.3 | SpaceMix results with sources of admixture. 64
Fig.S 7.4 | SpaceMix results with correlations of each tested model. 65
Fig.S 7.5 | SpaceMix results with no population text overlays. 66
Fig.S 7.6 | SpaceMix results with text instead of dots. 67
Fig.S 7.7 | SpaceMix results with sources of admixture. 68
Fig.S 7.8 | SpaceMix results with correlations of each model. 69
**8- Patterns of haplotype diversity 70**
Fig.S 8.1 | Haplotype richness (HR) for the AfricanNeo dataset. 70
Fig.S 8.2 | Haplotype heterozygosity (HH) for the AfricanNeo dataset. 71
Fig.S 8.3 | Maps of haplotype heterozygosity and haplotype richness for the AfricanNeo data. 72
Fig.S 8.4 | Linkage-disequilibrium (LD)-decay of the unmasked AfricanNeo dataset. 73
Fig.S 8.5 | Haplotype richness (HR) with masked AfricanNeo dataset. 74
Fig.S 8.6 | Haplotype heterozygosity (HH) with masked AfricanNeo dataset. 75
Fig.S 8.7 | Haplotype richness (HR) for the masked Only-BSP dataset. 76
Fig.S 8.8 | Haplotype heterozygosity (HH) for the masked Only-BSP. 77
Fig.S 8.9 | Maps of haplotype heterozygosity and haplotype richness for only BSP. 78
Fig.S 8.10 | Haplotype richness plotted against distance from Cameroon. 79
Fig.S 8.11 | Haplotype heterozygosity plotted against distance from Cameroon. 80
Fig.S 8.12 | Linkage-disequilibrium (LD)-decay of Only-BSP dataset. 81
Fig.S 8.13 | Spatial distribution of LD-decay in each studied dataset. 82
Fig.S 8.14 | Increase of LD-decay with geographical distances in studied populations. 83
**9- Maximum likelihood trees based on population allele frequencies 84**
Fig.S 9.1a | TreeMix for the masked Only-BSP dataset in rectangular shape. 84
Fig.S 9.1b | Coancestry matrix for the masked and imputed Only-BSP dataset. 85
Fig.S 9.2a | Population tree of the unmasked AfricanNeo dataset. 86

Fig.S 9.2b | TreeMix results for the unmasked AfricanNeo dataset. 87
Fig.S 9.3a | TreeMix for the unmasked Only-BSP dataset in rectangular shape. 88
Fig.S 9.3b | Coancestry matrix for the unmasked Only-BSP dataset. 89
**10- Visualizing migration routes in sub-Saharan Africa 90**
Fig.S 10.1 | Spatially explicit framework 90
Fig.S 10.2 | FST matrix for the masked Only-BSP dataset. 91
Fig.S 10.3 | FST map for the masked Only-BSP dataset. 92
Fig.S 10.4 | EEMS for the Only-African dataset. 93
Fig.S 10.5 | EEMS for the Only-African dataset after masking data of BSP. 94
Fig.S 10.6 | EEMS for the unmasked Only-BSP dataset. 95
Fig.S 10.7 | Comparisons between EEMS and ADMIXTURE results 96
Fig.S 10.8 | EEMS on the basis of the masked Only-Bantu dataset. 97
Fig.S 10.9 | FEEMS of the unmasked AfricanNeo dataset. 98
Fig.S 10.10 | FEEMS on the basis of the unmasked Only-BSP dataset. 99
Fig.S 10.11 | Spatial visualization of genetic barriers analysis on a grid. 100
**11- Estimated admixture dates in BSP 101**
Fig.S 11.1 | MOSAIC results for BSP with admixture. 101
Fig.S 11.2 | Admixture dates versus geographical distances from Cameroon. 102
**12- Comparisons between aDNA samples and modern-day African populations 103**
Fig.S 12.0 | Misincorporation patterns for the twelve ancient samples newly sequenced. 103
Fig.S 12.1 | Geographical locations of aDNA individuals included in this study. 104
Fig.S 12.2 | PCA of aDNA and modern African populations. 105
Fig.S 12.3 | PCA-UMAP of aDNA individuals and present-day African populations. 107
Fig.S 12.4 | ADMIXTURE results at K=4 of ancient and modern African populations. 108
Fig.S 12.5 | ADMIXTURE results at K=12 of ancient and modern African populations. 109
Fig.S 12.6 | Genetic affinity of ancient samples UPS013, UPS017a, UPS029 to modern BSP estimated using the f3-statistics. 110
Fig.S 12.7 | Genetic affinity of ancient samples WUD034, WUD037, WUD038b to modern BSP estimated using the f3-statistics. 111
Fig.S 12.8 | Genetic affinity of ancient samples WUD003, WUD004, WUD008 to modern BSP estimated using the f3-statistics. 112
Fig.S 12.9 | Genetic affinity of ancient samples WUD010, WUD012, WUD018 to modern BSP estimated using the f3-statistics. 113
Fig.S 12.11 | PCA of Only-Zambia database 114
Fig.S 12.12 | PCA of ancient samples and modern BSP from Zambia. 115
Fig.S 12.13 | PCA of ancient samples and modern BSP from South Africa. 116
Fig.S 12.14 | PCA-UMAP of ancient samples and modern BSP from South Africa. 117
**13- PCA results after masking datasets 118**
Fig.S 13.1 | PCA of BSP and six selected reference panels. 118
Fig.S 13.2 | Ancestry-specific (AS-)PCA of BSP and six selected reference panels. 119
**14. List of Supplementary Tables included in the Excel file 120**

#### 1- Extended introduction and geographical locations of studied populations

**a** **b**

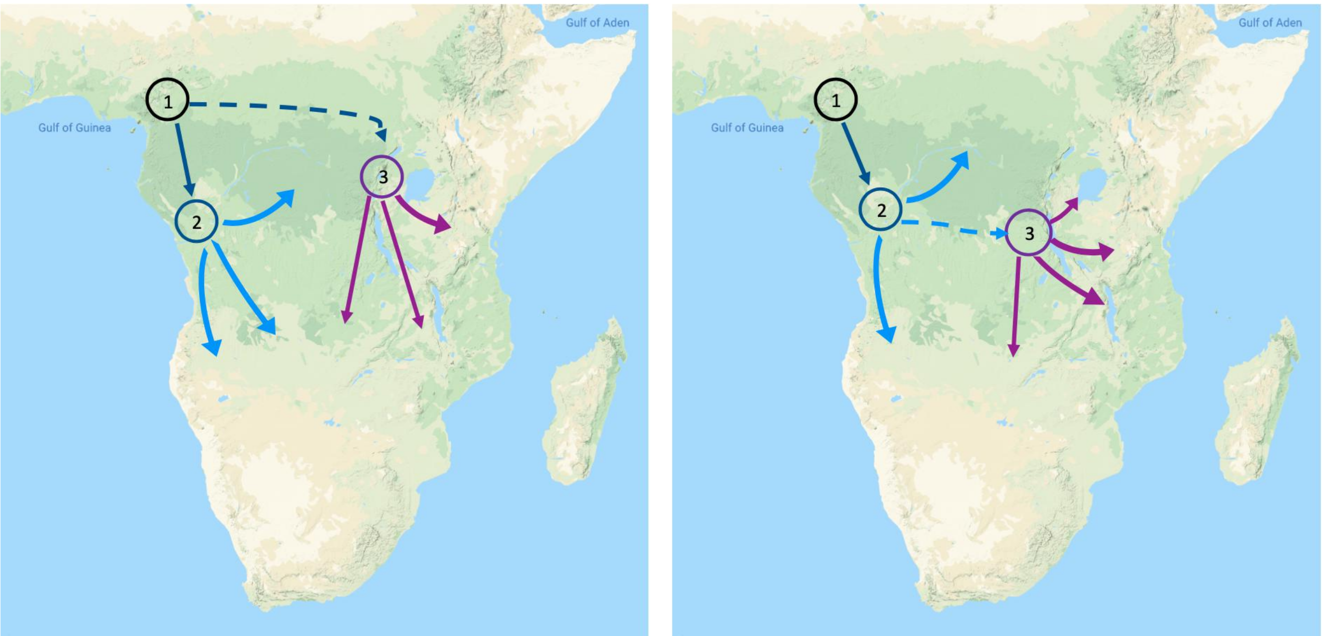


**Fig.S 1.1 | Linguistic hypotheses proposed to explain the expansion of BSP.**
Illustration of the (**a**) “Early-split” and (**b**) “Late-split” hypotheses based on linguistic sources. The main difference between the two is whether the eastern Bantu branch is a direct off-shoot of the Narrow Bantu languages (left) or a sub-branch of western Bantu branch (right). The circles represent the presumed locations of proto-Bantu (1), the nucleus of the western Bantu branch (2), and the nucleus of the eastern branch of the Bantu languages (3). Map data by Google ©2022.

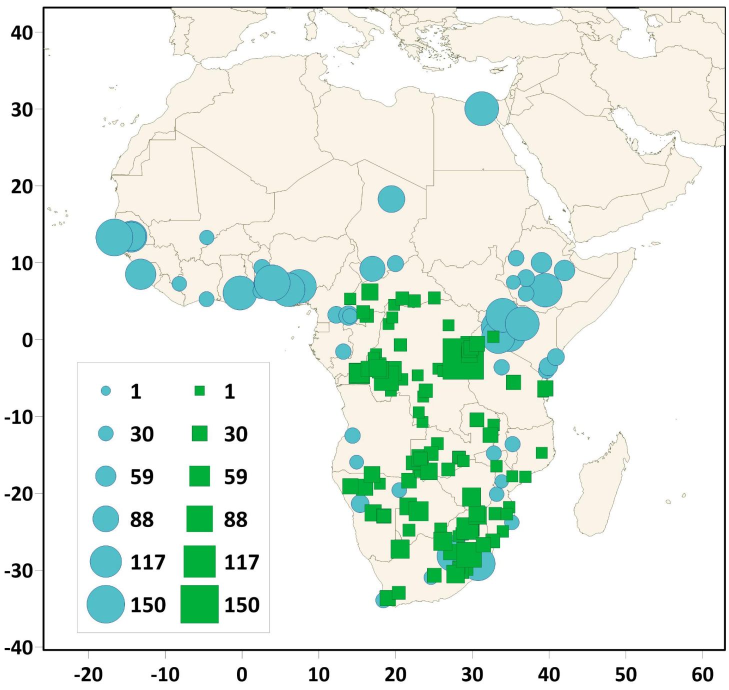


**Fig.S 1.2 | Geographical locations and sample sizes of African populations.**
Figure showing populations that were included in the newly genotyped dataset (green squares) and reference populations from previous studies (blue circles). The size of the markers is in relation to the sample size of each population. Details about the populations were presented in **Table.S 1–2**.

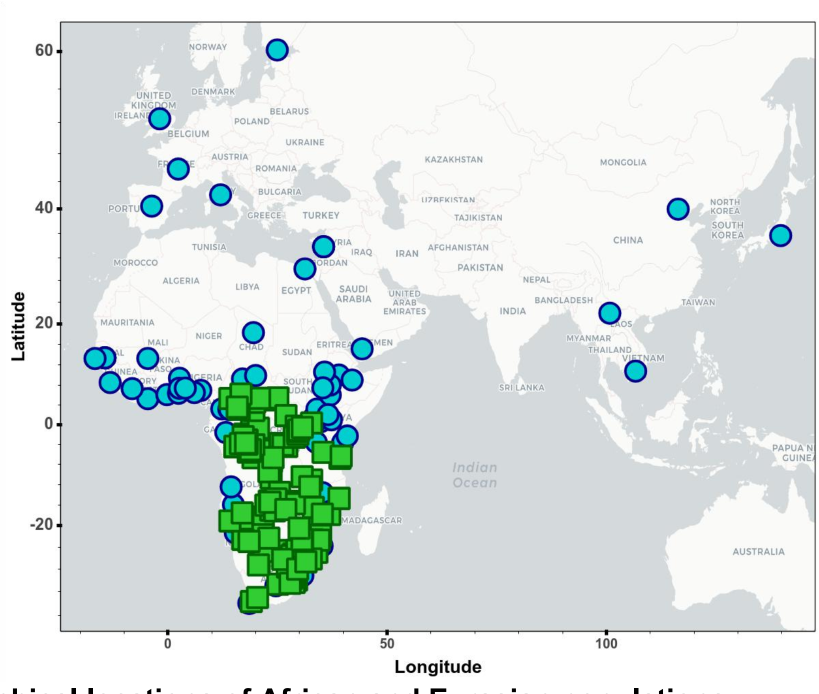


**Fig.S 1.3 | Geographical locations of African and Eurasian populations.**
Figure showing locations of 227 populations (5,341 individuals) included in the dataset after quality filtering and merging. This included the genotyped dataset (green squares) and reference populations from previous studies (blue circles) for a total of 214 African and 13 Eurasian populations. To better visualize the locations of each population we created interactive plots (see **Fig.S_1.3_Map.html**).

a

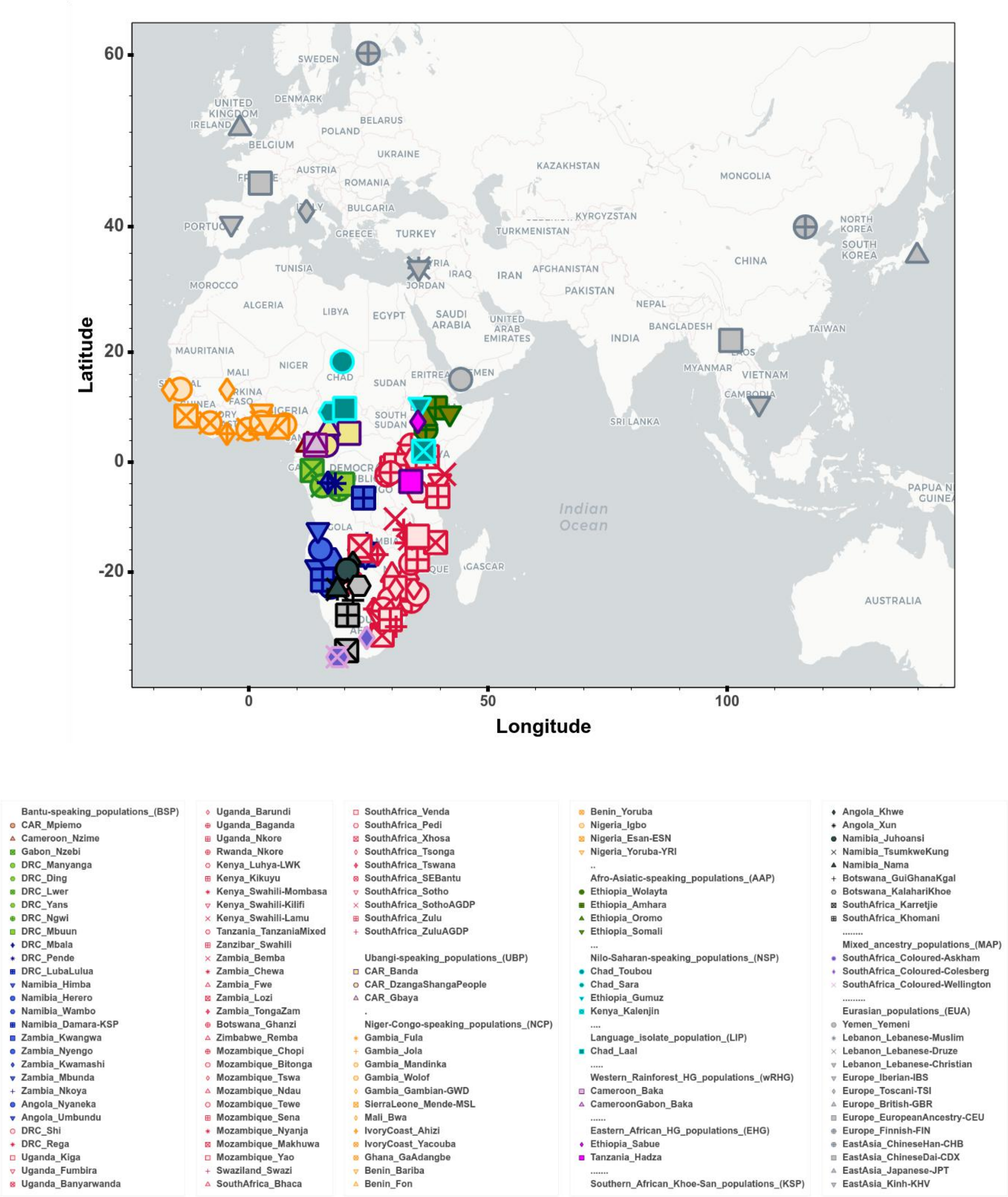
60
40
20
0
-20
0
50
100
Latitude
Longitude
SWEDEN
DENMARK
UNITED KINGDOM
IRELAND
BELARUS
POLAND
BELGIUM
UKRAINE
AUSTRIA
KAZAKHSTAN
MONGOLIA
ROMANIA
BULGARIA
KYRGYZSTAN
GREECE
TURKEY
TURKMENISTAN
NORTH KOREA
SOUTH KOREA
CHINA
TUNISIA
IRAQ
IRAN
AFGHANISTAN
MOROCCO
JORDAN
PAKISTAN
ALGERIA
LIBYA
EGYPT
SAUDI ARABIA
UNITED ARAB EMIRATES
NEPAL
BANGLADESH
INDIA
TAIWAN
MAURITANIA
NIGER
MALI
CHAD
SUDAN
MYANMAR
VIETNAM
SOUTH SUDAN
SRI LANKA
Indian Ocean
AUSTRALIA
Bantu-speaking_populations_(BSP)
CAR_Mpiemo
Cameroon_Nzime
Gabon_Nzebi
DRC_Manyanga
DRC_Ding
DRC_Lwer
DRC_Yans
DRC_Ngwi
DRC_Mbuun
DRC_Mbala
DRC_Pende
DRC_LubaLulua
Namibia_Himba
Namibia_Herero
Namibia_Wambo
Namibia_Damara-KSP
Zambia_Kwangwa
Zambia_Nyengo
Zambia_Kwamashi
Zambia_Mbunda
Zambia_Nkoya
Angola_Nyaneka
Angola_Umbundu
DRC_Shi
DRC_Rega
Uganda_Kiga
Uganda_Fumbira
Uganda_Banyarwanda
Uganda_Barundi
Uganda_Baganda
Uganda_Nkore
Rwanda_Nkore
Kenya_Luhya-LWK
Kenya_Kikuyu
Kenya_Swahili-Mombasa
Kenya_Swahili-Kilifi
Kenya_Swahili-Lamu
Tanzania_TanzaniaMixed
Zanzibar_Swahili
Zambia_Bemba
Zambia_Chewa
Zambia_Fwe
Zambia_Lozi
Zambia_TongaZam
Botswana_Ghanzi
Zimbabwe_Remba
Mozambique_Chopi
Mozambique_Bitonga
Mozambique_Tswa
Mozambique_Ndau
Mozambique_Tewe
Mozambique_Sena
Mozambique_Nyanja
Mozambique_Makhuwa
Mozambique_Yao
Swaziland_Swazi
SouthAfrica_Bhaca
SouthAfrica_Venda
SouthAfrica_Pedi
SouthAfrica_Xhosa
SouthAfrica_Tsonga
SouthAfrica_Tswana
SouthAfrica_SEBantu
SouthAfrica_Sotho
SouthAfrica_SothoAGDP
SouthAfrica_Zulu
SouthAfrica_ZuluAGDP
Ubangi-speaking_populations_(UBP)
CAR_Banda
CAR_DzangaShangaPeople
CAR_Gbaya
Niger-Congo-speaking_populations_(NCP)
Gambia_Fula
Gambia_Jola
Gambia_Mandinka
Gambia_Wolof
Gambia_Gambian-GWD
SierraLeone_Mende-MSL
Mali_Bwa
IvoryCoast_Ahizi
IvoryCoast_Yacouba
Ghana_GaAdangbe
Benin_Bariba
Benin_Fon
Benin_Yoruba
Nigeria_Igbo
Nigeria_Esan-ESN
Nigeria_Yoruba-YRI
Afro-Asiatic-speaking_populations_(AAP)
Ethiopia_Wolayta
Ethiopia_Amhara
Ethiopia_Oromo
Ethiopia_Somali
Nilo-Saharan-speaking_populations_(NSP)
Chad_Toubou
Chad_Sara
Ethiopia_Gumuz
Kenya_Kalenjin
Language_isolate_population_(LIP)
Chad_Laal
Western_Rainforest_HG_populations_(wRHG)
Cameroon_Baka
CameroonGabon_Baka
Eastern_African_HG_populations_(EHG)
Ethiopia_Sabue
Tanzania_Hadza
Southern_African_Khoe-San_populations_(KSP)
Angola_Khwe
Angola_Xun
Namibia_Juhoansi
Namibia_TsumkweKung
Namibia_Nama
Botswana_GuiGhanaKgal
Botswana_KalahariKhoe
SouthAfrica_Karretjie
SouthAfrica_Khomani
Mixed_ancestry_populations_(MAP)
SouthAfrica_Coloured-Askham
SouthAfrica_Coloured-Colesberg
SouthAfrica_Coloured-Wellington
Eurasian_populations_(EUA)
Yemen_Yemeni
Lebanon_Lebanese-Muslim
Lebanon_Lebanese-Druze
Lebanon_Lebanese-Christian
Europe_Iberian-IBS
Europe_Toscani-TSI
Europe_British-GBR
Europe_EuropeanAncestry-CEU
Europe_Finnish-FIN
EastAsia_ChineseHan-CHB
EastAsia_ChineseDai-CDX
EastAsia_Japanese-JPT
EastAsia_Kinh-KHV

**b**

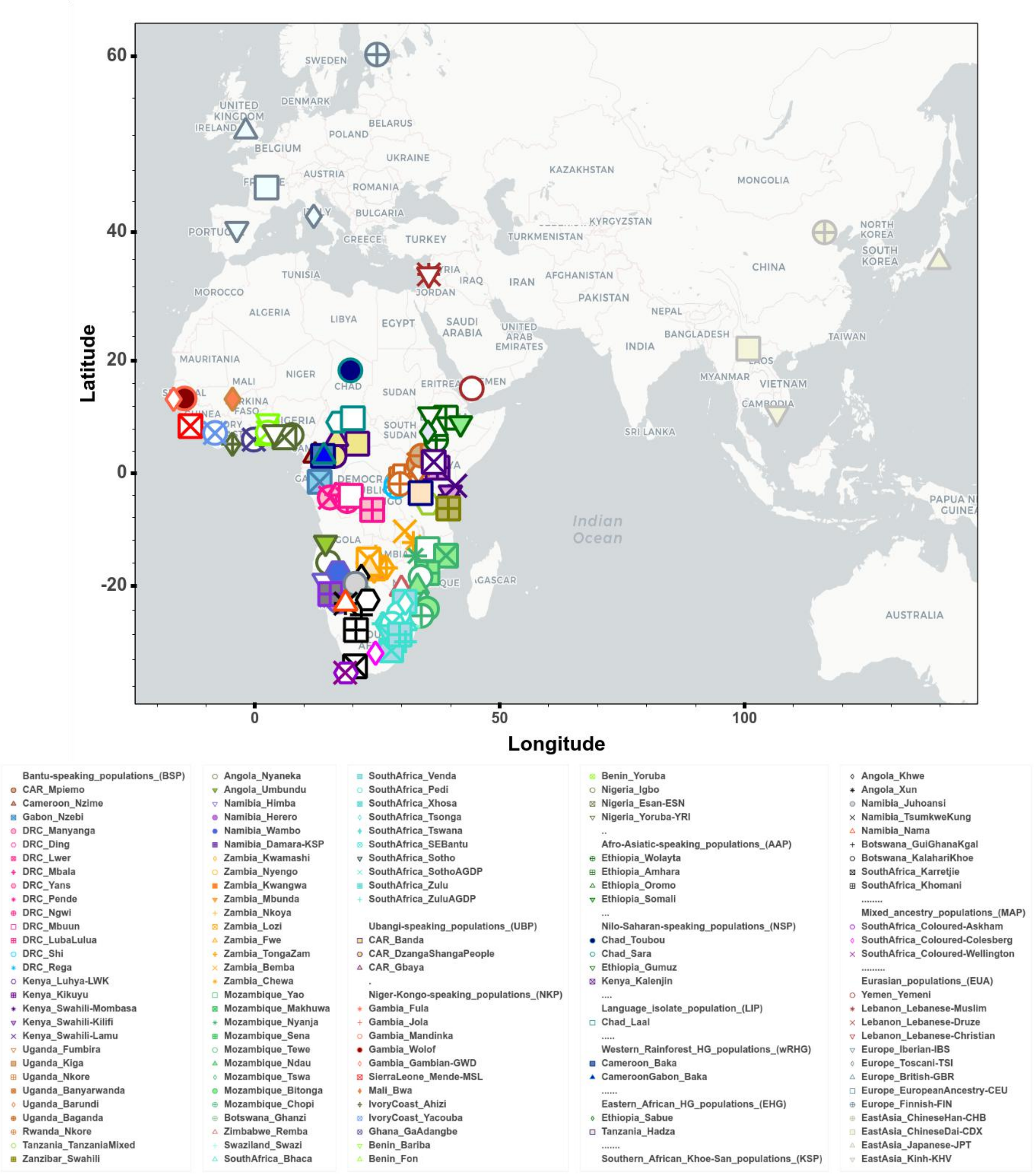


**Fig.S 1.4 | Labels of all the groups and populations included in the AfricanNeo dataset.**
Figure showing the labels of (**a**) each group and (**b**) each population that was included in the AfricanNeo dataset for populations with at least 10 individuals. Further details about the populations are presented elsewhere (**Table.S 2**). (**a**) BSP were grouped into four major linguistic groups: north-western Bantu 2 (in brown), west-western Bantu (in green), south-western Bantu (in dark blue), and eastern Bantu speakers (in red). To better visualize the locations of each group and each population we created interactive plots (see **FigS_1.4a_Map.html** and **FigS_1.4b_Map.html**, respectively).

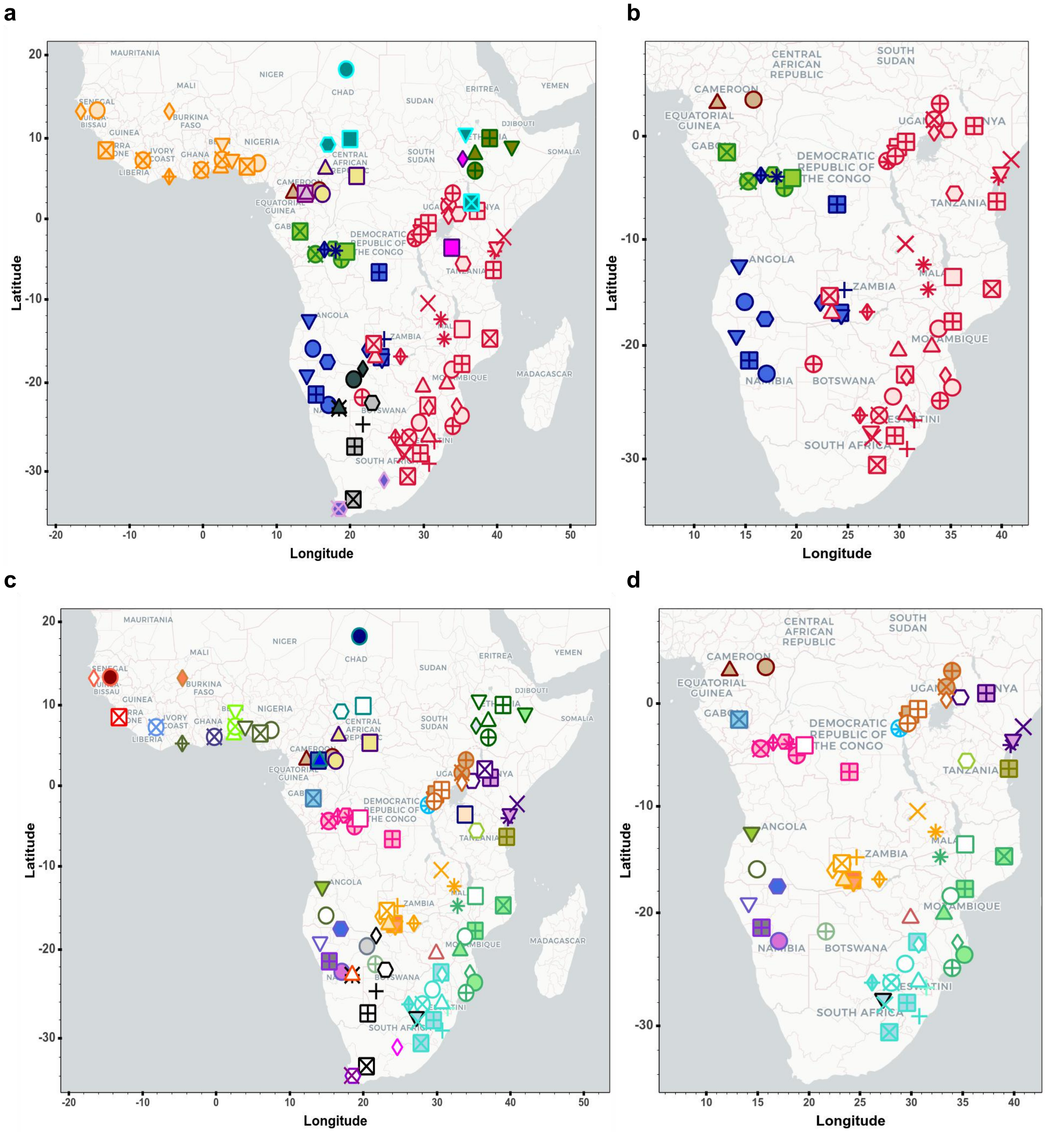


**Fig.S 1.5 | Labels of the populations included in the Only-Africa and Only-BSP datasets.**
Panel figure showing the labels of the populations that were included in the Only-Africa dataset for (**a**) each group (also **Fig. 1a**) and (**c**) each population; and populations included in the Only-BSP dataset for (**b**) each group and (**d**) each population. Population labels are matching the labels presented in **Fig.S 1.4**. Further details about the populations are presented elsewhere (**Table.S 2**). To better visualize the locations of each population we created an interactive plot (by zooming into **Fig.S_1.5a_Map.html**, **Fig.S_1.5b_Map.html**, **Fig.S_1.5c_Map.html** and **Fig.S_1.5d_Map.html**).

Legend for each Bantu-speaking group in **Fig.S 1.5a**.

Bantu-speaking_populations_(BSP)
CAR_Mpiemo
Cameroon_Nzime
Gabon_Nzebi
DRC_Manyanga
DRC_Ding
DRC_Lwer
DRC_Yans
DRC_Ngwi
DRC_Mbuun
DRC_Mbala
DRC_Pende
DRC_LubaLulua
Namibia_Himba
Namibia_Herero
Namibia_Wambo
Namibia_Damara-KSP
Zambia_Kwangwa
Zambia_Nyengo
Zambia_Kwamashi
Zambia_Mbunda
Zambia_Nkoya
Angola_Nyaneka
Angola_Umbundu
DRC_Shi
DRC_Rega
Uganda_Kiga
Uganda_Fumbira
Uganda_Banyarwanda
Uganda_Barundi
Uganda_Baganda
Uganda_Nkore

Rwanda_Nkore
Kenya_Luhya-LWK
Kenya_Kikuyu
Kenya_Swahili-Mombasa
Kenya_Swahili-Kilifi
Kenya_Swahili-Lamu
Tanzania_TanzaniaMixed
Zanzibar_Swahili
Zambia_Bemba
Zambia_Chewa
Zambia_Fwe
Zambia_Lozi
Zambia_TongaZam
Botswana_Ghanzi
Zimbabwe_Remba
Mozambique_Chopi
Mozambique_Bitonga
Mozambique_Tswa
Mozambique_Ndau
Mozambique_Tewe
Mozambique_Sena
Mozambique_Nyanja
Mozambique_Makhuwa
Mozambique_Yao
Swaziland_Swazi
SouthAfrica_Bhaca
SouthAfrica_Venda
SouthAfrica_Pedi
SouthAfrica_Xhosa
SouthAfrica_Tsonga
SouthAfrica_Tswana
SouthAfrica_SEBantu

SouthAfrica_Sotho
SouthAfrica_SothoAGDP
SouthAfrica_Zulu
SouthAfrica_ZuluAGDP

Ubangi-speaking_populations_(UBP)
CAR_Banda
CAR_DzangaShangaPeople
CAR_Gbaya
.
Niger-Congo-speaking_populations_(NCP)
Gambia_Fula
Gambia_Jola
Gambia_Mandinka
Gambia_Wolof
Gambia_Gambian-GWD
SierraLeone_Mende-MSL
Mali_Bwa
IvoryCoast_Ahizi
IvoryCoast_Yacouba
Ghana_GaAdangbe
Benin_Bariba
Benin_Fon
Benin_Yoruba
Nigeria_Igbo
Nigeria_Esan-ESN
Nigeria_Yoruba-YRI
..
Afro-Asiatic-speaking_populations_(AAP)
Ethiopia_Wolayta
Ethiopia_Amhara
Ethiopia_Oromo

Ethiopia_Somali
...
Nilo-Saharan-speaking_populations_(NSP)
Chad_Toubou
Chad_Sara
Ethiopia_Gumuz
Kenya_Kalenjin
....
Language_isolate_population_(LIP)
Chad_Laal
.....
Western_Rainforest_HG_populations_(wRHG)
Cameroon_Baka
CameroonGabon_Baka
......
Eastern_African_HG_populations_(EHG)
Ethiopia_Sabue
Tanzania_Hadza
.......
Southern_African_Khoe-San_populations_(KSP)
Angola_Khwe
Angola_Xun
Namibia_Juhoansi
Namibia_TsumkweKung
Namibia_Nama
Botswana_GuiGhanaKgal
Botswana_KalahariKhoe
SouthAfrica_Karretjie
SouthAfrica_Khomani
........
Mixed_ancestry_populations_(MAP)
SouthAfrica_Coloured-Askham
SouthAfrica_Coloured-Colesberg
SouthAfrica_Coloured-Wellington

Legend for each Bantu-speaking population in **Fig.S 1.5b**.

Bantu-speaking_populations_(BSP)
CAR_Mpiemo
Cameroon_Nzime
Gabon_Nzebi
DRC_Manyanga
DRC_Ding
DRC_Lwer
DRC_Mbala
DRC_Yans
DRC_Pende
DRC_Ngwi
DRC_Mbuun
DRC_LubaLulua
DRC_Shi
DRC_Rega
Kenya_Luhya-LWK
Kenya_Kikuyu
Kenya_Swahili-Mombasa
Kenya_Swahili-Kilifi
Kenya_Swahili-Lamu
Uganda_Fumbira
Uganda_Kiga
Uganda_Nkore
Uganda_Banyarwanda
Uganda_Barundi
Uganda_Baganda
Rwanda_Nkore
Tanzania_TanzaniaMixed
Zanzibar_Swahili
Angola_Nyaneka
Angola_Umbundu
Namibia_Himba

Namibia_Herero
Namibia_Wambo
Namibia_Damara-KSP
Zambia_Kwamashi
Zambia_Nyengo
Zambia_Kwangwa
Zambia_Mbunda
Zambia_Nkoya
Zambia_Lozi
Zambia_Fwe
Zambia_TongaZam
Zambia_Bemba
Zambia_Chewa
Mozambique_Yao
Mozambique_Makhuwa
Mozambique_Nyanja
Mozambique_Sena
Mozambique_Tewe
Mozambique_Ndau
Mozambique_Tswa
Mozambique_Bitonga
Mozambique_Chopi
Botswana_Ghanzi
Zimbabwe_Remba
Swaziland_Swazi
SouthAfrica_Bhaca
SouthAfrica_Venda
SouthAfrica_Pedi
SouthAfrica_Xhosa
SouthAfrica_Tsonga
SouthAfrica_Tswana
SouthAfrica_SEBantu

SouthAfrica_Sotho
SouthAfrica_SothoAGDP
SouthAfrica_Zulu
SouthAfrica_ZuluAGDP

Ubangi-speaking_populations_(UBP)
CAR_Banda
CAR_DzangaShangaPeople
CAR_Gbaya
.
Niger-Congo-speaking_populations_(NCP)
Gambia_Fula
Gambia_Jola
Gambia_Mandinka
Gambia_Wolof
Gambia_Gambian-GWD
SierraLeone_Mende-MSL
Mali_Bwa
IvoryCoast_Ahizi
IvoryCoast_Yacouba
Ghana_GaAdangbe
Benin_Bariba
Benin_Fon
Benin_Yoruba
Nigeria_Igbo
Nigeria_Esan-ESN
Nigeria_Yoruba-YRI
..
Afro-Asiatic-speaking_populations_(AAP)
Ethiopia_Wolayta
Ethiopia_Amhara
Ethiopia_Oromo

Ethiopia_Somali
...
Nilo-Saharan-speaking_populations_(NSP)
Chad_Toubou
Chad_Sara
Ethiopia_Gumuz
Kenya_Kalenjin
....
Language_isolate_population_(LIP)
Chad_Laal
.....
Western_Rainforest_HG_populations_(wRHG)
Cameroon_Baka
CameroonGabon_Baka
......
Eastern_African_HG_populations_(EHG)
Ethiopia_Sabue
Tanzania_Hadza
.......
Southern_African_Khoe-San_populations_(KSP)
Angola_Khwe
Angola_Xun
Namibia_Juhoansi
Namibia_TsumkweKung
Namibia_Nama
Botswana_GuiGhanaKgal
Botswana_KalahariKhoe
SouthAfrica_Karretjie
SouthAfrica_Khomani
........
Mixed_ancestry_populations_(MAP)
SouthAfrica_Coloured-Askham
SouthAfrica_Coloured-Colesberg
SouthAfrica_Coloured-Wellington

- **"Genotyped" dataset:** 1,740 individuals in total for this study using the Illumina H3Africa array, 2,271,503 SNPs (**Table.S 1**).

*Quality control (QC) using PLINK (--mind 0.15 --geno 0.1 --hwe 0.0000001)*

- **Database after QC 2,221,827 autosome SNPs and 1,567 individuals.**

*Merging with datasets previously genotyped in:*

*Whole-genome sequencing data (1000 Genomes Project Consortium et al. 2015),*

*Illumina HumanOmni5 array (Fortes-Lima et al. 2017),*

*Illumina HumanOmni2.5 array (Schlebusch et al. 2012; Vicente et al. 2019; Semo et al. 2020; Fortes-Lima et al. 2017; Gurdasani et al. 2015; Pagani et al. 2015; Haber et al. 2016; Crawford et al. 2017; Brucato et al. 2018; Sjöstrand et al. 2021)*

- "**Full-Genotyped" dataset**: 5,341 individuals from 227 populations and 482,459 SNPs.

*Removed 103 populations with less than 10 individuals per population.*

- "**AfricanNeo" dataset**: 4,950 individuals from 124 populations and 482,459 SNPs. [AfricanNeo LD-pruned dataset: 223,473 SNPs]

*Removed 13 Eurasian populations.*

- "**Only-African" dataset**: 3,902 individuals from 111 sub-Saharan African populations and 482,459 SNPs.

*Removed 57 non-Bantu-speaking populations.*

- "**Only-BSP" dataset**: 2,108 individuals from 67 BSP and 482,459 SNPs.

*Large populations randomly downsampled to 30 individuals.*

- "**Masked Only-BSP" dataset**: 1,495 individuals from 67 BSP and 482,459 SNPs.

- "**Africa-aDNA" dataset**: Only-African dataset, 12 new aDNA individuals, and 83 ancient DNA (aDNA) individuals from previous studies (Skoglund et al. 2017; Schlebusch et al. 2017; Prendergast et al. 2019; Lipson et al. 2020, 2022; Wang et al. 2020; Coutinho et al. 2021). 5,036 individuals and 482,459 SNPs.

*Removed 57 non-Bantu-speaking populations (BSP).*

- "**BSP-aDNA" dataset**: 2,194 individuals and 482,459 SNPs.

**Fig.S 1.6 | Workflow for the assembled datasets.**
Panel figure summarizing the databases assembled for this study.

#### 2- Dimensionality reduction methods

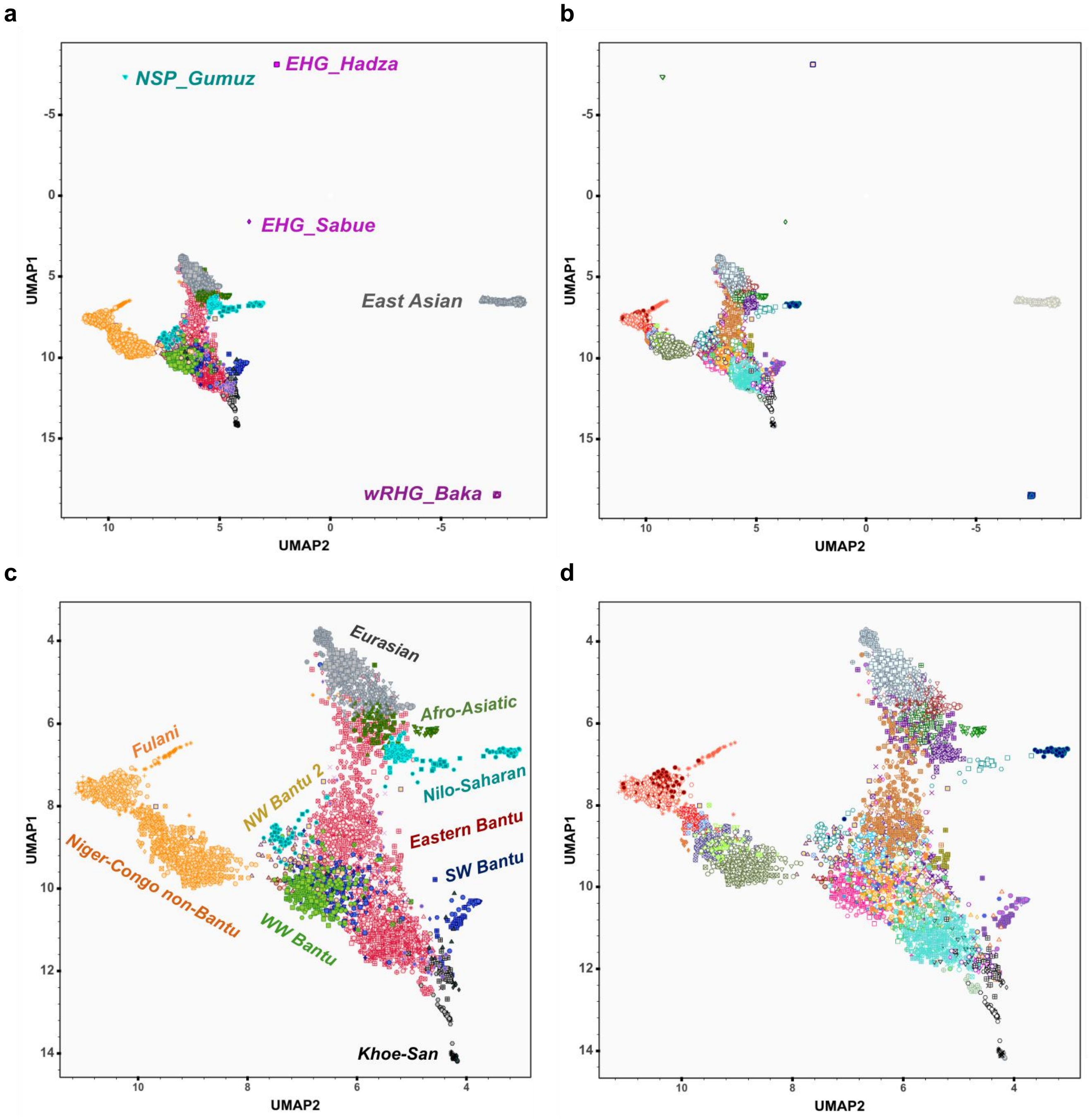


**Fig.S 2.1 | UMAP approach on the basis of genotype data.**
Panel figure showing UMAP results for all African and Eurasian populations included in the AfricanNeo dataset (**Table.S 1**) using the Uniform Manifold Approximation and Projection (UMAP) algorithm directly on the genotype data without performing a PCA. This panel figure highlights results for (**a**) each group listed in **Fig.S 1.4a** and (**b**) each population listed in **Fig.S 1.4b**. The legend of this figure is the same legend as **Fig.S 1.4**. To better see the results for the studied BSP, we zoom in on each plot highlighting (**c**) each group (also **Fig. 1B**) and (**d**) each population. To better visualize the results of each group or population, plots were included as interactive plots (see **Fig.S_2.1a_UMAP_plot.html** and **Fig.S_2.1b_UMAP_plot.html**).

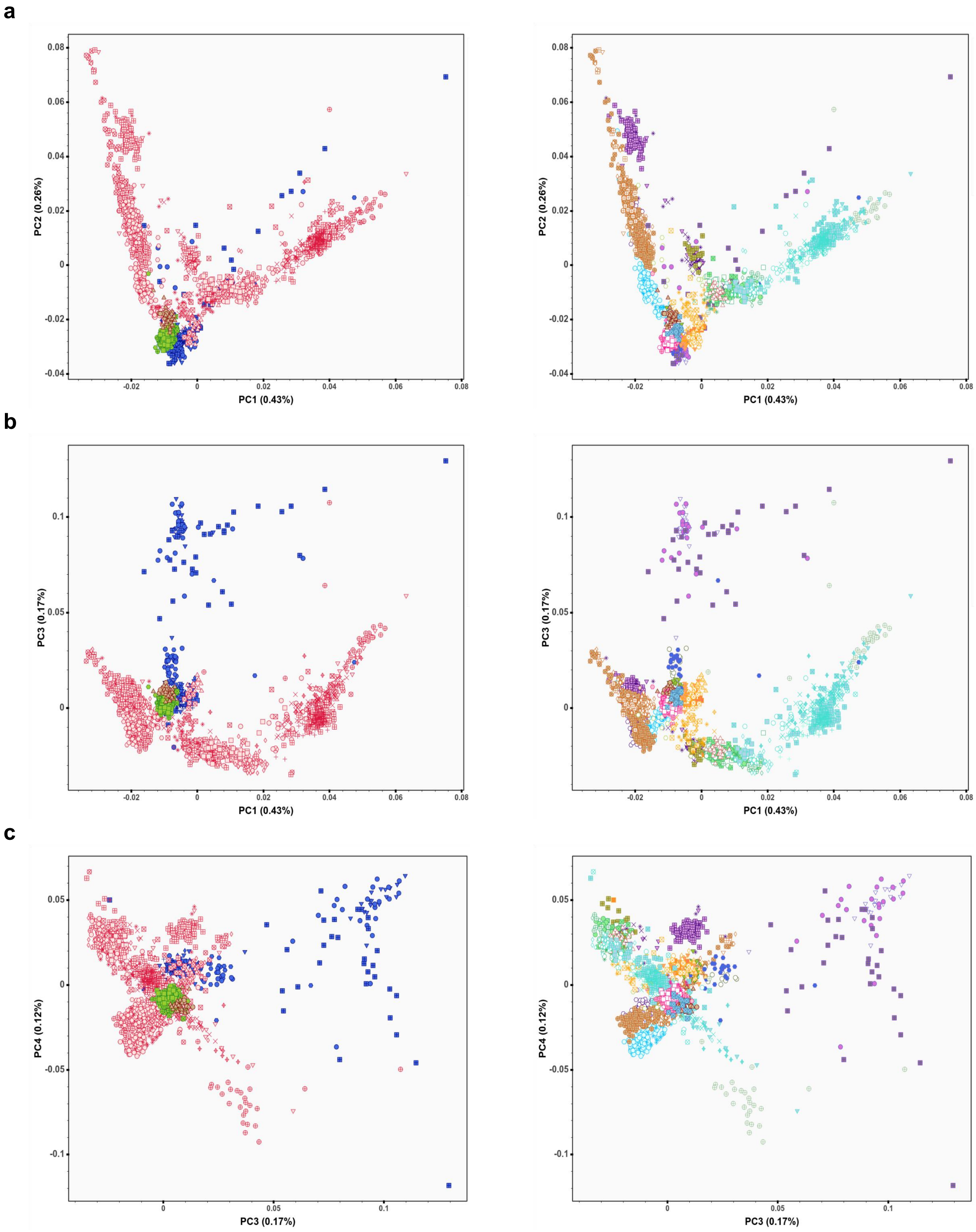


**Fig.S 2.2 | PCA plots for only Bantu-speaking populations.**
Panel figure showing PCA results of only individuals from the Only-BSP dataset (2,108 Bantu-speaking individuals; **Table.S 2**), for PC projections obtained for each group (left column) and for each population (right column) between: (**a**) PC1 vs PC2; (**b**) PC1 vs PC3; and (**c**) PC3 vs PC4. The legend is the same as in **Fig.S 6c** (left column) and **Fig.S 6d** (right column). The first ten PC projections were also included in interactive plots (**Fig.S_2.2_PCA_Only-BSP_Groups.html** and **Fig.S_2.2_PCA_Only-BSP_Populations.html**).

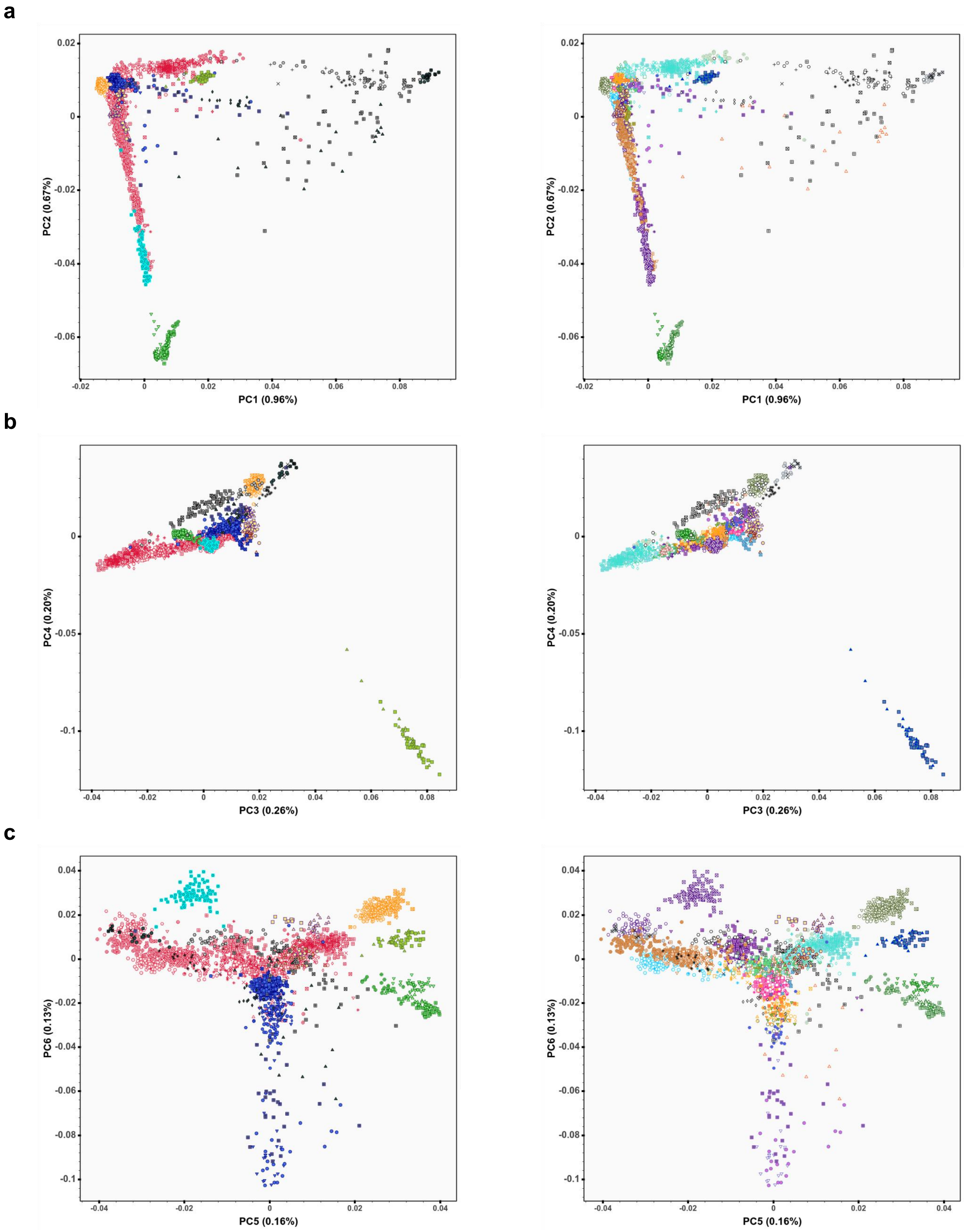


**Fig.S 2.3 | PCA plots for selected sub-Saharan African populations.**
Panel figure showing PCA results of selected sub-Saharan African populations that were included in the AfricanNeo dataset (**Table.S 2**), for PC projections obtained for each group (left column) and for each population (right column) between: (**a**) PC1 vs PC2; (**b**) PC3 vs PC4; and (**c**) PC5 vs PC6. To better visualize the results of each population, the first ten PC projections were also included in interactive plots (see **Fig.S_2.3_PCA_SSA_Groups.html** and **Fig.S_2.3_PCA_SSA_Groups.html**).

Legend for each group (left column) in panel **Fig.S 2.3**.

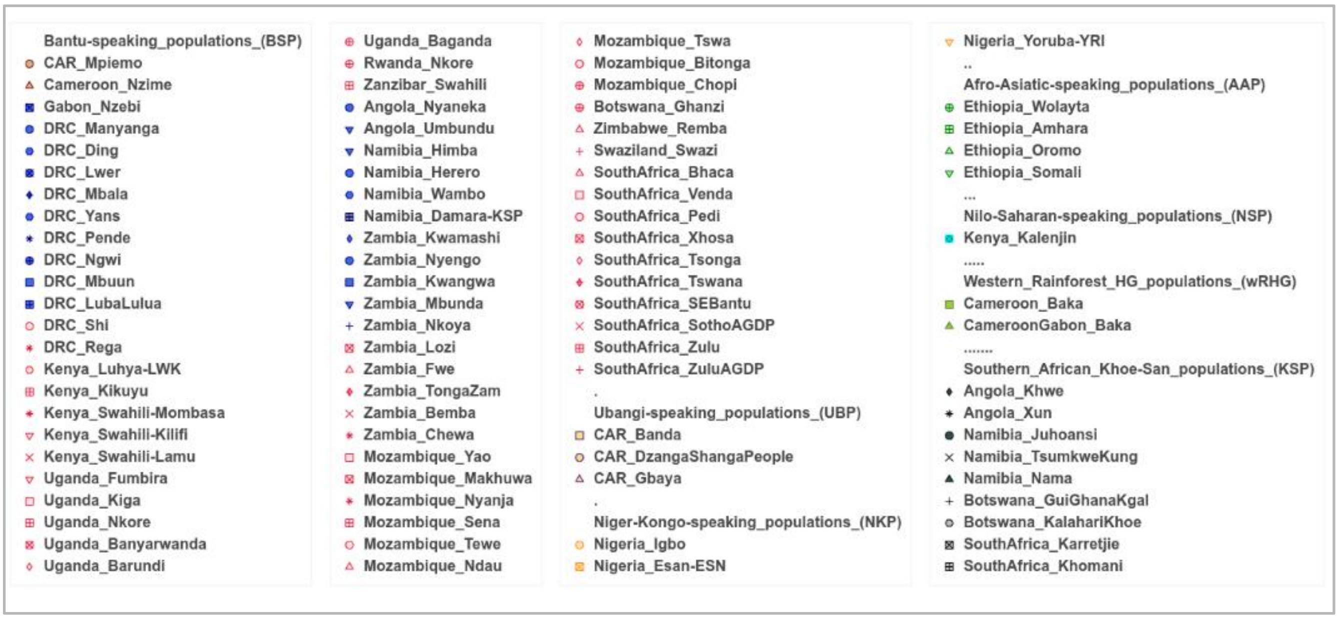


Legend for each population (right column) in panel **Fig.S 2.3**.

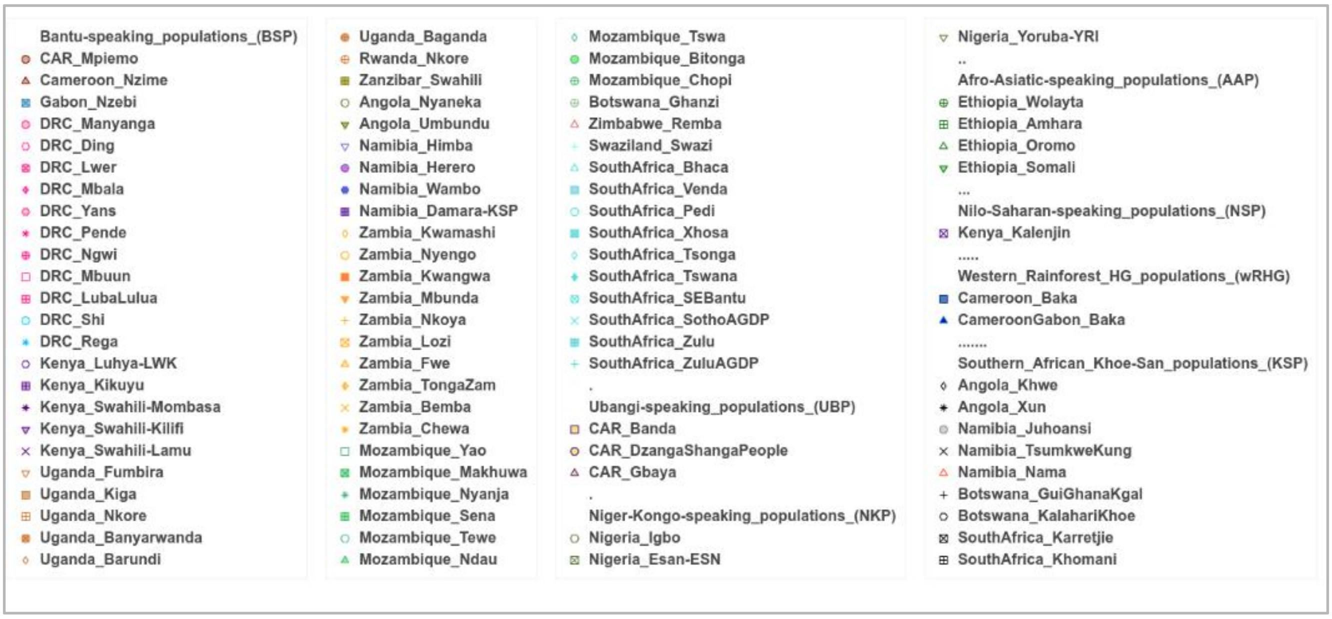

a

b

c

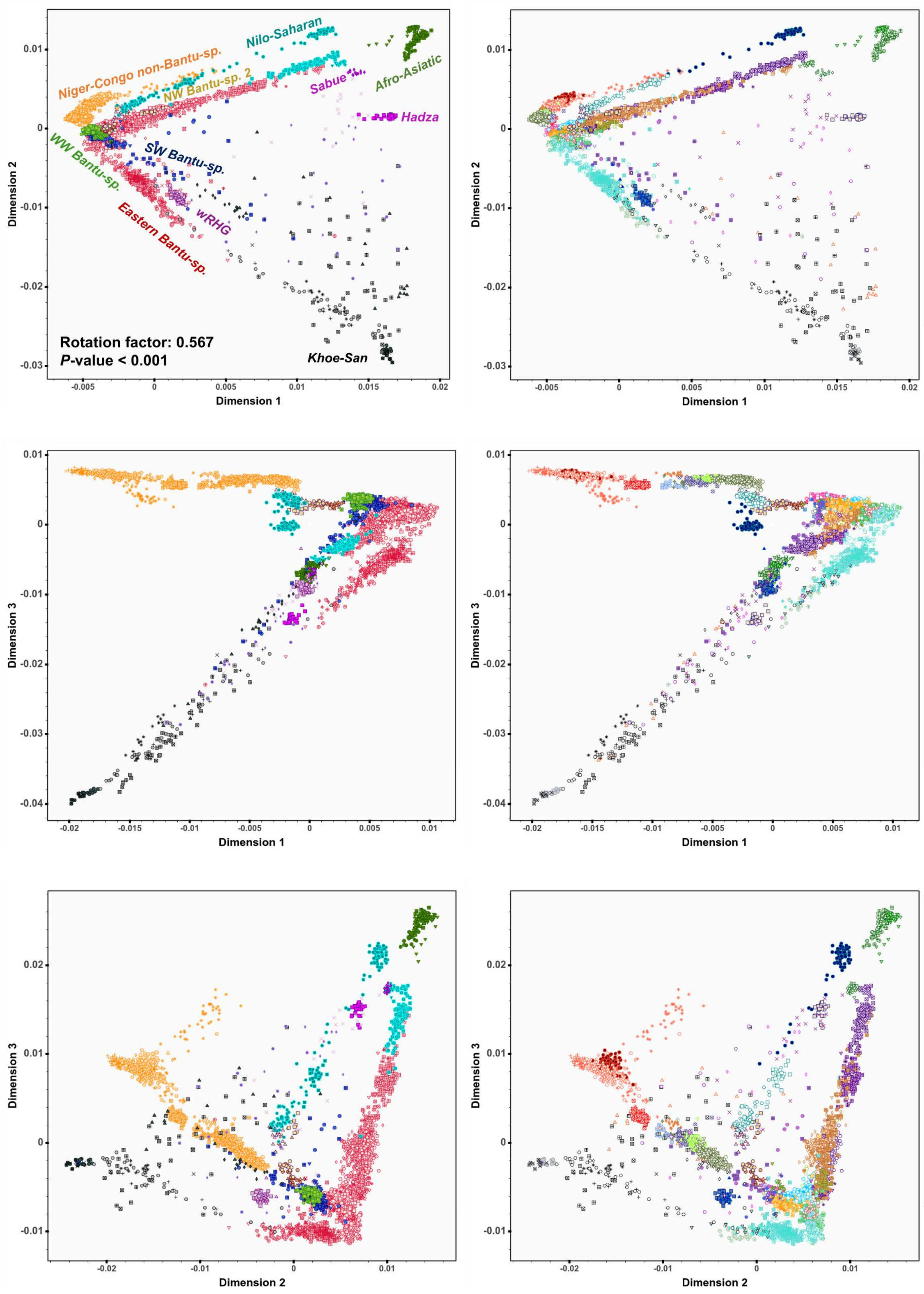


**Fig.S 2.4 | Procrustes rotated PCA for the Only-African dataset.**
Panel figure showing procrustes rotated PCA for sub-Saharan African populations (3,902 individuals) included in the AfricanNeo dataset (**Table.S 2**), for projections obtained for each group (left column; **Fig.S 1.4a**) and for each population (right column; **Fig.S 1.4B**) between: (**a**) Dim1 vs Dim2; (**b**) Dim1 vs Dim3; and (**c**) Dim2 vs Dim3. Estimated correlations in a symmetric Procrustes rotation were: 0.5671, 0.7105 and 0.7676, respectively; and all of them were significant (*P*-value < 0.001). Interactive plots were included **Fig.S_2.4*_Procrustes_PCA_*.html**).

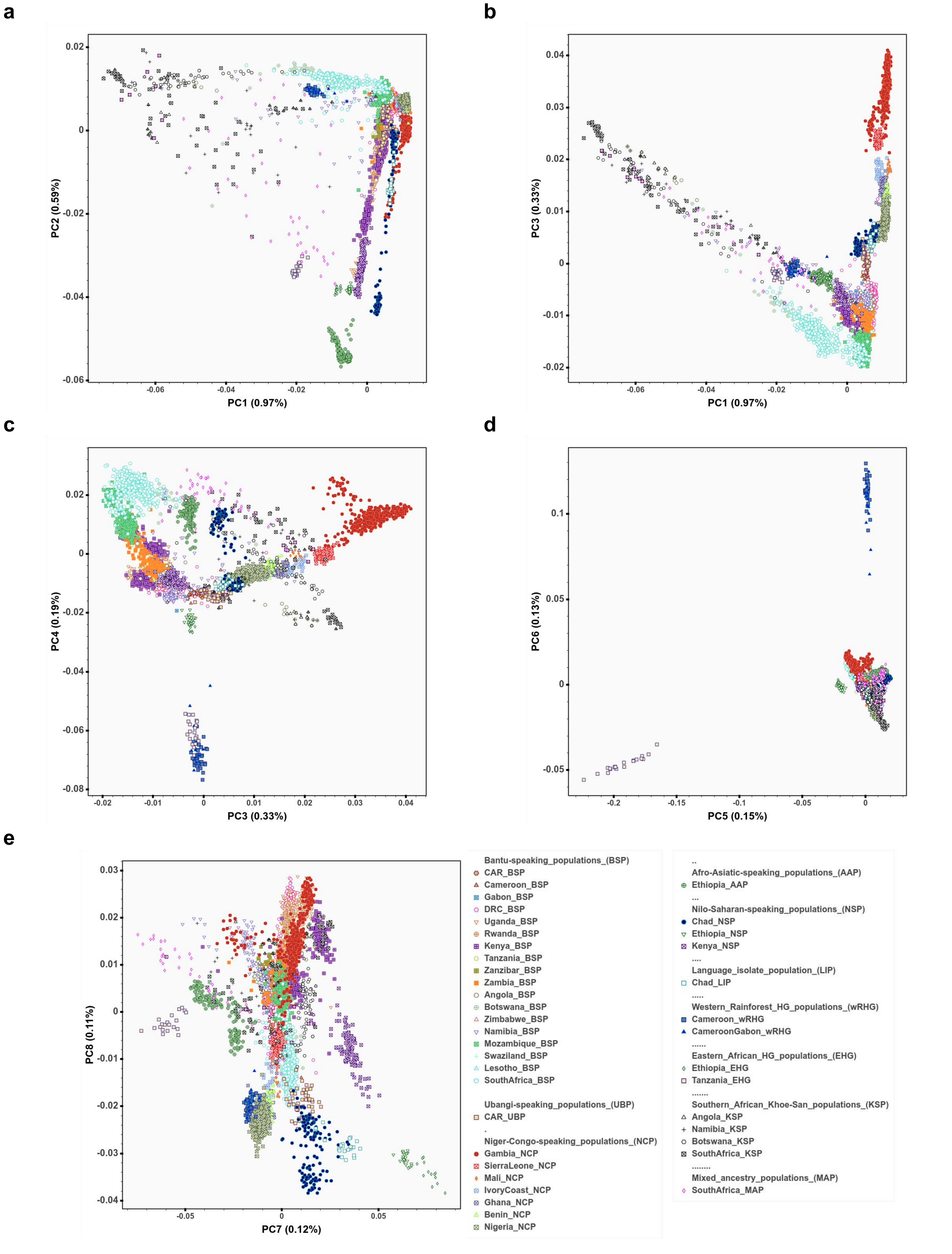


**Fig.S 2.5 | PCA for the Full-Genotyped dataset and reference sub-Saharan African populations.** Panel figure showing PCA results of all newly genotyped samples from Africa that were included in the Full-Genotyped (**Table.S 1**) plus reference sub-Saharan African populations included in the AfricanNeo dataset (**Table.S 2**) after quality control and merging. Panel figure showing each PC projection: (**a**) PC1 vs PC2; (**b**) PC1 vs PC3; (**c**) PC3 vs PC4; (**d**) PC5 vs PC6; and (**e**) PC7 vs PC8.

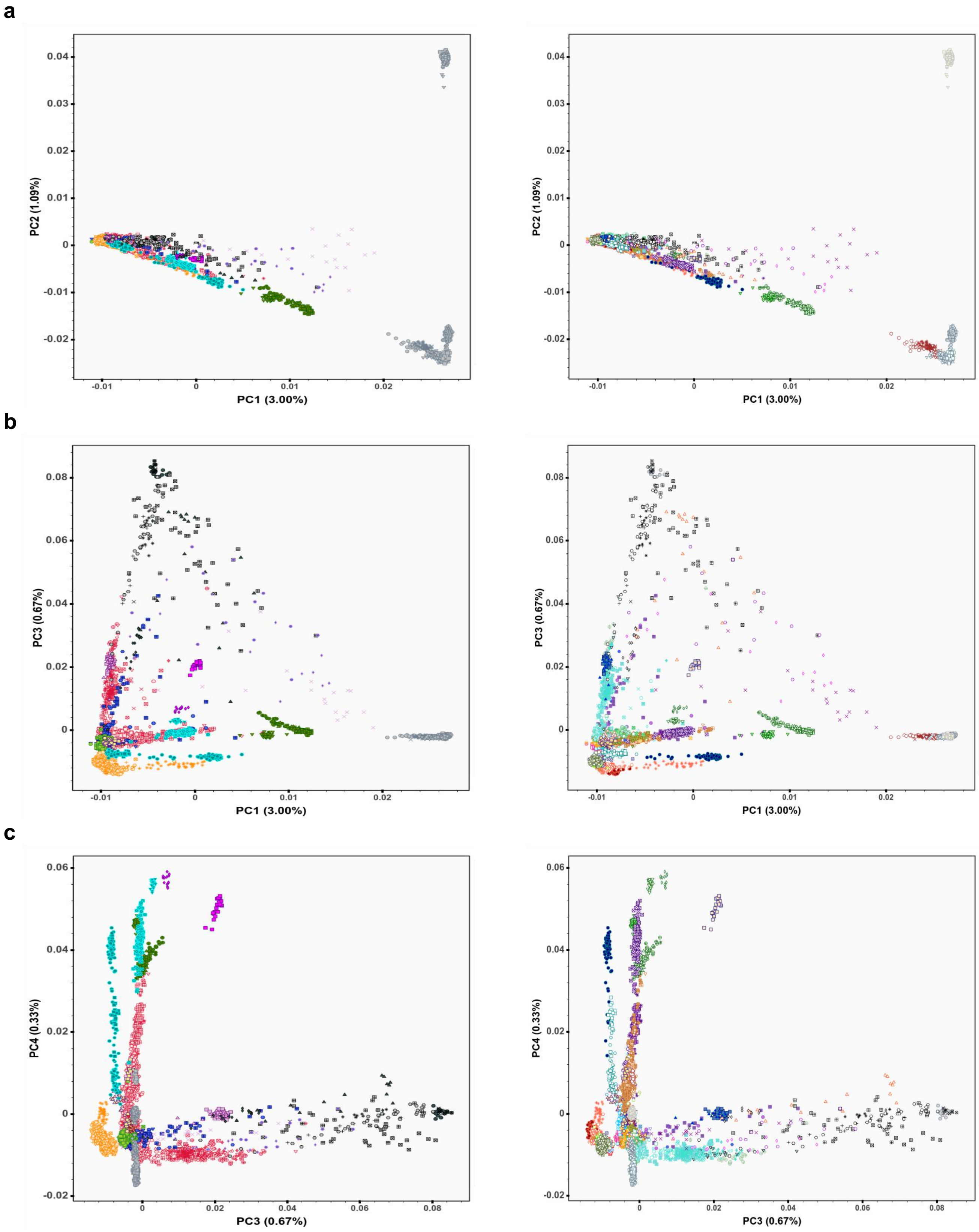


**Fig.S 2.6 | PCA for each group included in the AfricanNeo dataset.**
Panel figure showing PCA of worldwide groups included in the AfricanNeo dataset (4,950 individuals; in total from 124 populations; **Fig.S 1.4b** and **Table.S 2**), for PC projections obtained for each group (left column) and for each population (right column) between: (**a**) PC1 vs PC2; (**b**) PC1 vs PC3; and (**c**) PC3 vs PC4. The first ten PC projections were included in interactive plots (see

**Fig.S_2.6_PCA_AfricanNeo_Groups.html** and **Fig.S_2.6_PCA_AfricanNeo_Populations.html**). The legend of this figure is the same legend as in **Fig.S 1.4**.

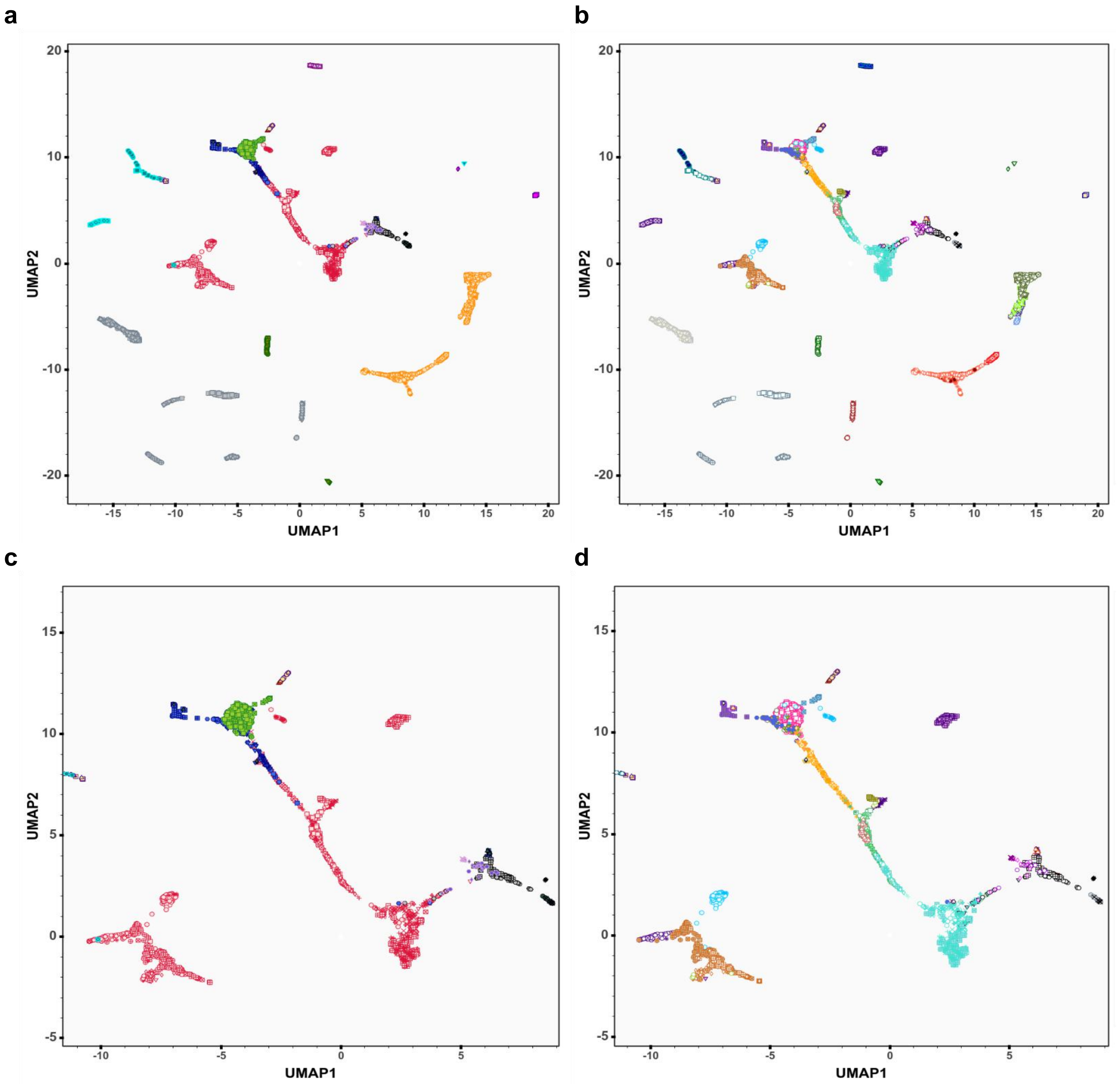


**Fig.S 2.7 | PCA-UMAP approach for the AfricanNeo dataset.**

Panel figure showing PCA-UMAP results of populations included in the AfricanNeo dataset using UMAP algorithm to combine the information of the first 10 PCs of the PCA (**Fig.S 2.6**). This panel figure highlights results for (**a**) each group listed in **Fig.S 1.4a** and (**b**) each population listed in **Fig.S 1.4b**. To better see the results for the studied BSP, we zoom in on each plot highlighting (**c**) each Bantu-speaking group and (**d**) each Bantu-speaking population. To better visualize the result, plots were included as interactive plots (see **Fig.S_2.7a_PCA-UMAP_plot.html** and **Fig.S_2.7b_PCA-UMAP_plot.html**). The legend of this figure is the same legend as in **Fig.S 1.4**.

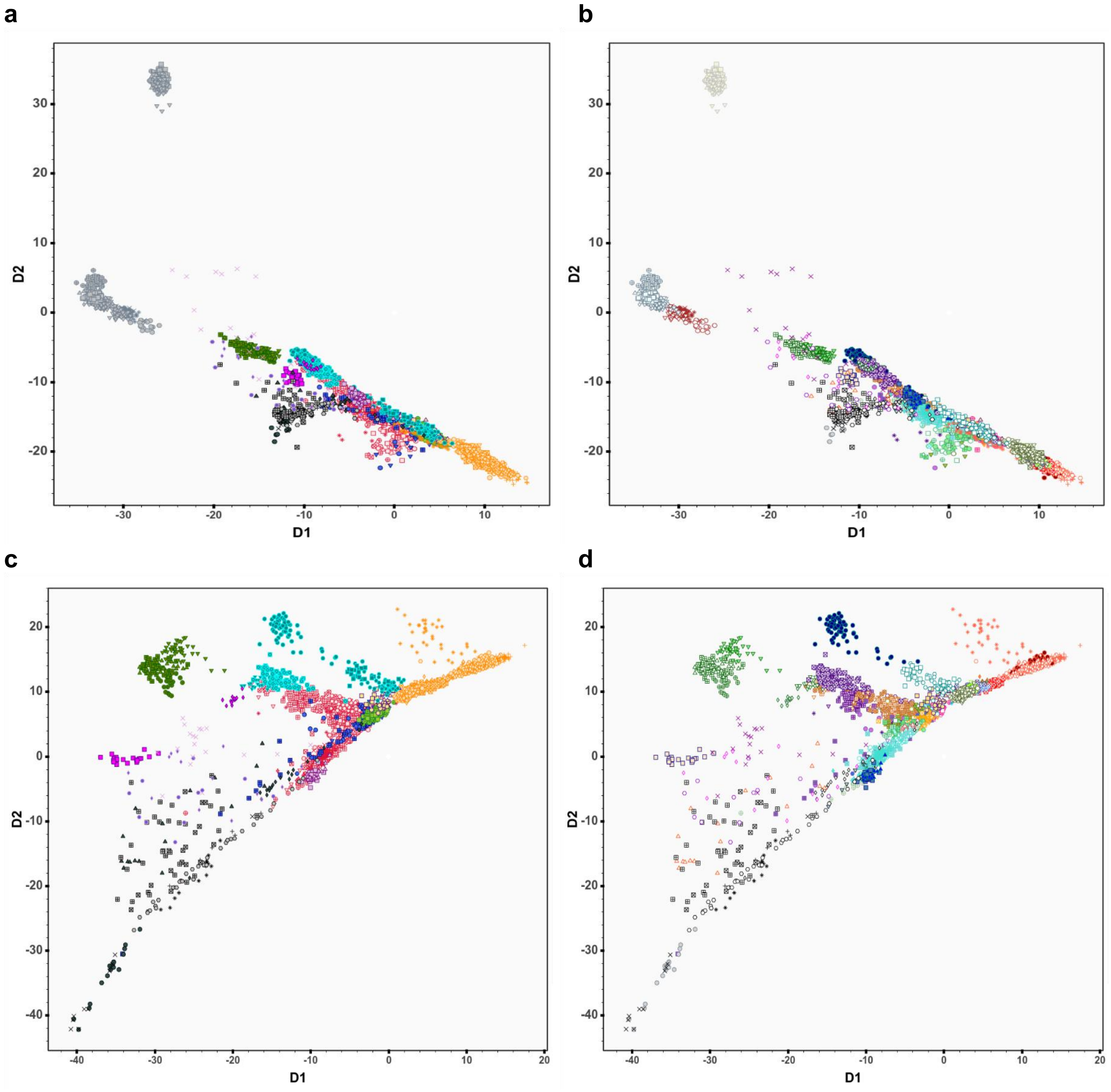


**Fig.S 2.8 | GCAE approach for the AfricanNeo and Only-African datasets.**
Panel figure showing GCAE results of all African and Eurasian populations included in the AfricanNeo dataset (4,950 individuals; in total from 124 populations; **Table.S 2**) were estimated using Genotype Convolutional Autoencoder (GCAE) approach, a deep learning framework for dimensionality reduction. This panel figure highlights (**a**) each group listed in **Fig.S 1.4a** and (**b**) each population listed in **Fig.S 1.4b**. Figure also shows GCAE results of all populations included in the Only-African dataset (**Table.S 2**), (**c**) for each group and (**d**) for each population. The legend of this figure is the same legend as in **Fig.S 1.4**.To better visualize the results of each group or population, the results were included in interactive plots (see **Fig.S_2.8a_GCAE_plot.html**, **Fig.S_2.8b_GCAE_plot.html**, **Fig.S_2.8c_GCAE_plot.html**, and **Fig.S_2.8d_GCAE_plot.html**).

#### 3- Unsupervised clustering analyses and F-statistics analyses

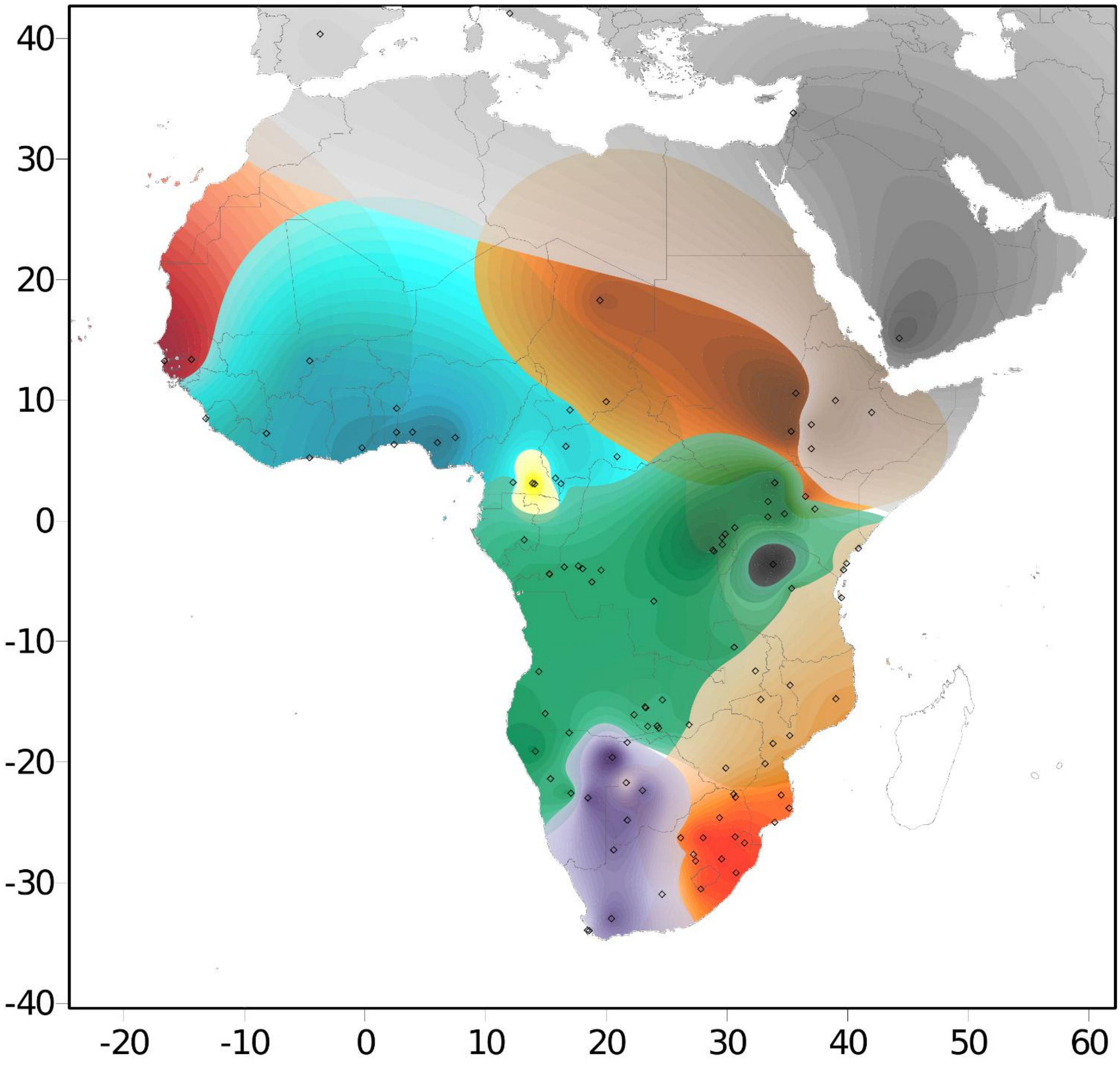


**Fig.S 3.1 | Surfer map of ADMIXTURE at K=12.**
Panel figure showing a contour map overlapping unsupervised ADMIXTURE results at K=12 created using the Kriging method for all the populations included in the AfricanNeo dataset (**Fig.S 1.4**). The geographical distributions of average admixture proportions were represented for nine ancestries found in high frequencies in African populations. Ancestry components with values under 25% are not represented in this figure, while all the estimated values (from 0% to 100%) for each represented ancestry component are shown **Fig.S 3.2**.

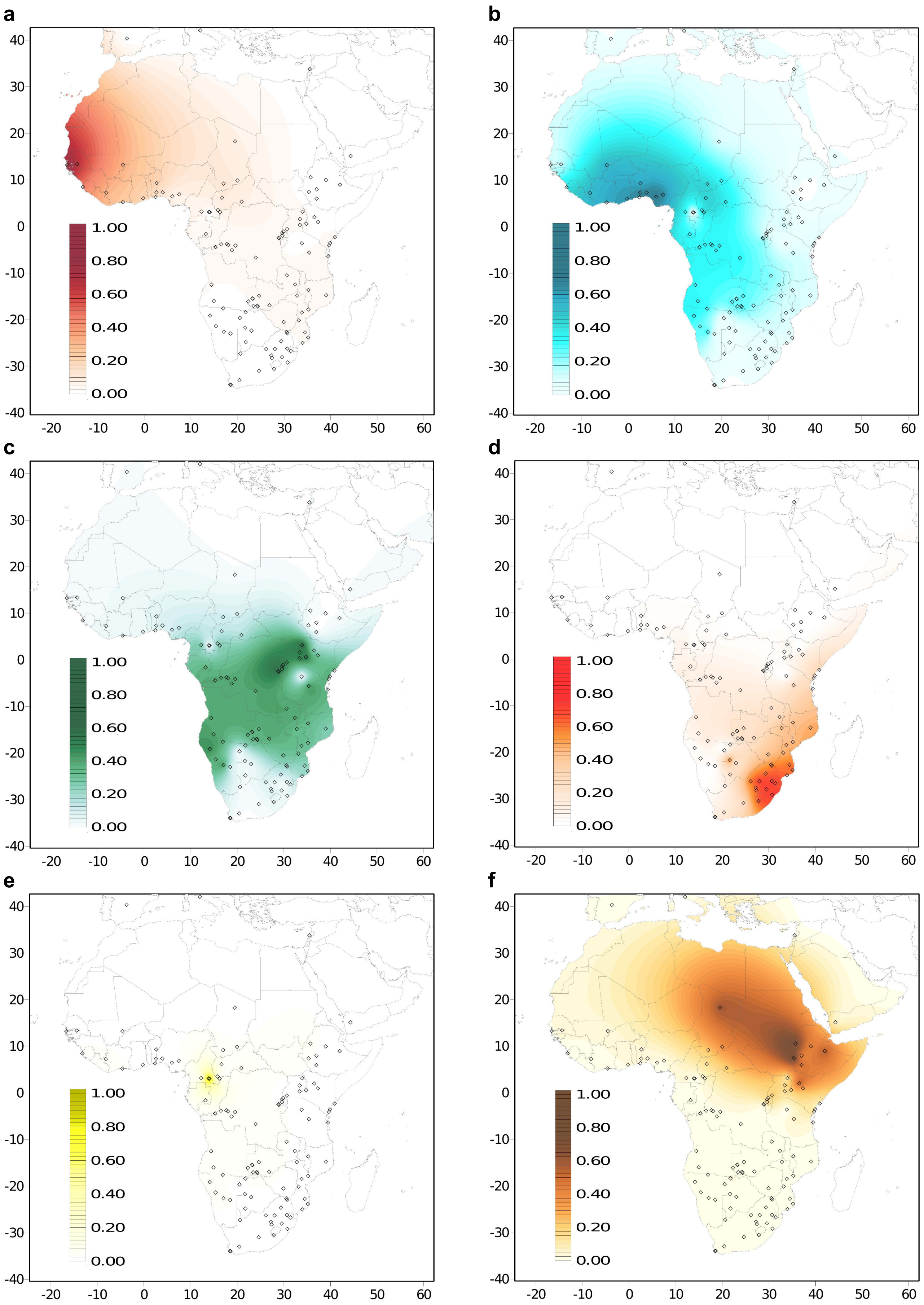
a
b
c
d
e
f
40
30
20
10
0
-10
-20
-30
-40
-20
-10
0
10
20
30
40
50
60
1.00
0.80
0.60
0.40
0.20
0.00

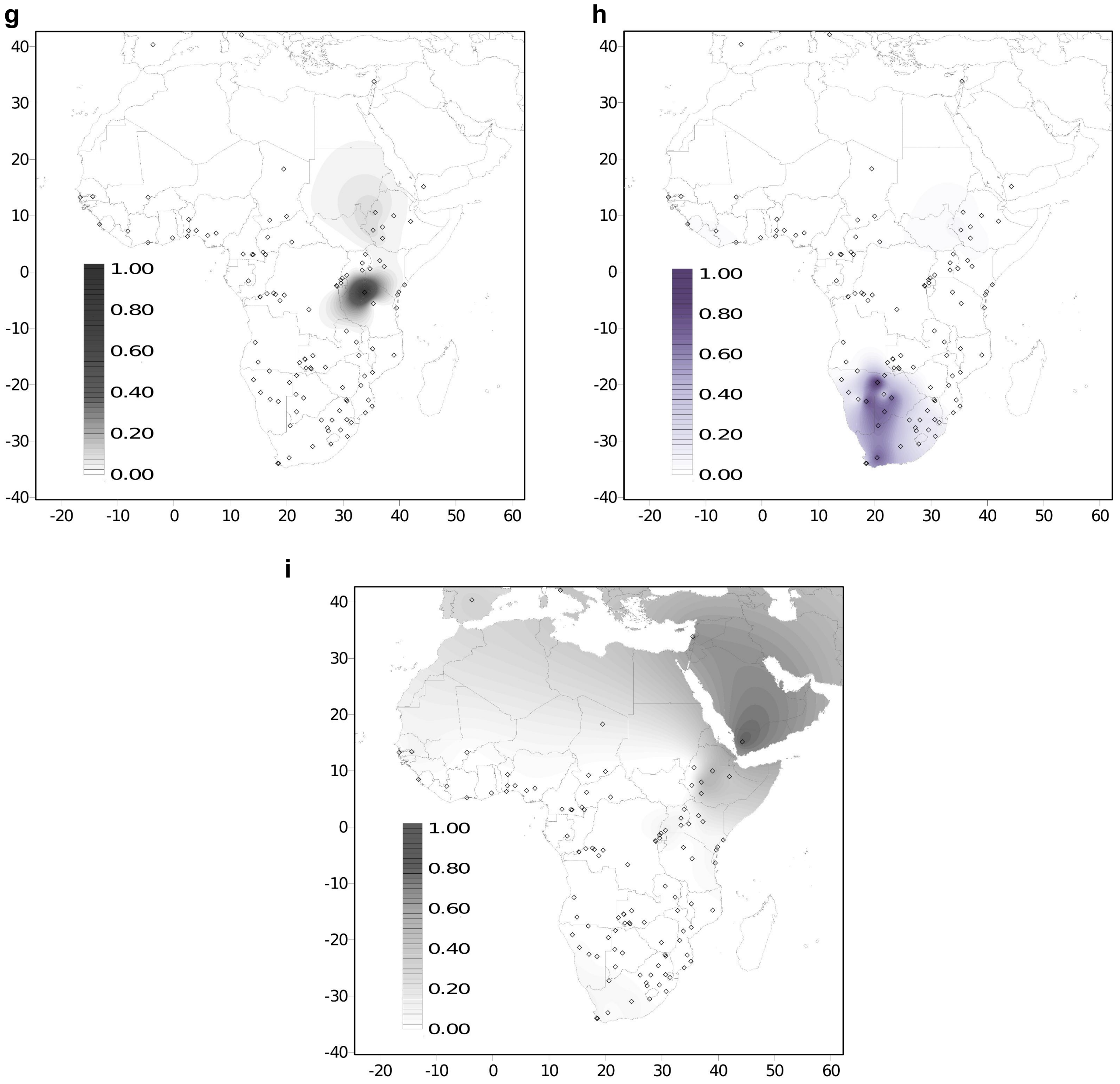


**Fig.S 3.2 | Surfer map of each ADMIXTURE result at K=12.**
Panel figure showing contour maps of unsupervised ADMIXTURE results at K=12 were created using the Kriging method for all the populations included in the AfricanNeo dataset. Average admixture proportions across Africa were depicted for nine main components across the African continent.
(**a**) red component or west African-related ancestry, (**b**) blue component or west-central African-related ancestry, (**c**) green component or Bantu-speaking-related ancestry, (**d**) orange component or south-eastern Bantu-speaking-related ancestry, (**e**) yellow component or western RHG-related ancestry, (**f**) brown component or eastern African-related ancestry, (**g**) black component or eastern RHG-related ancestry, (**h**) purple component or Khoi-San-speaking-related ancestry, and (**i**) gray component or Middle Eastern-related ancestry.

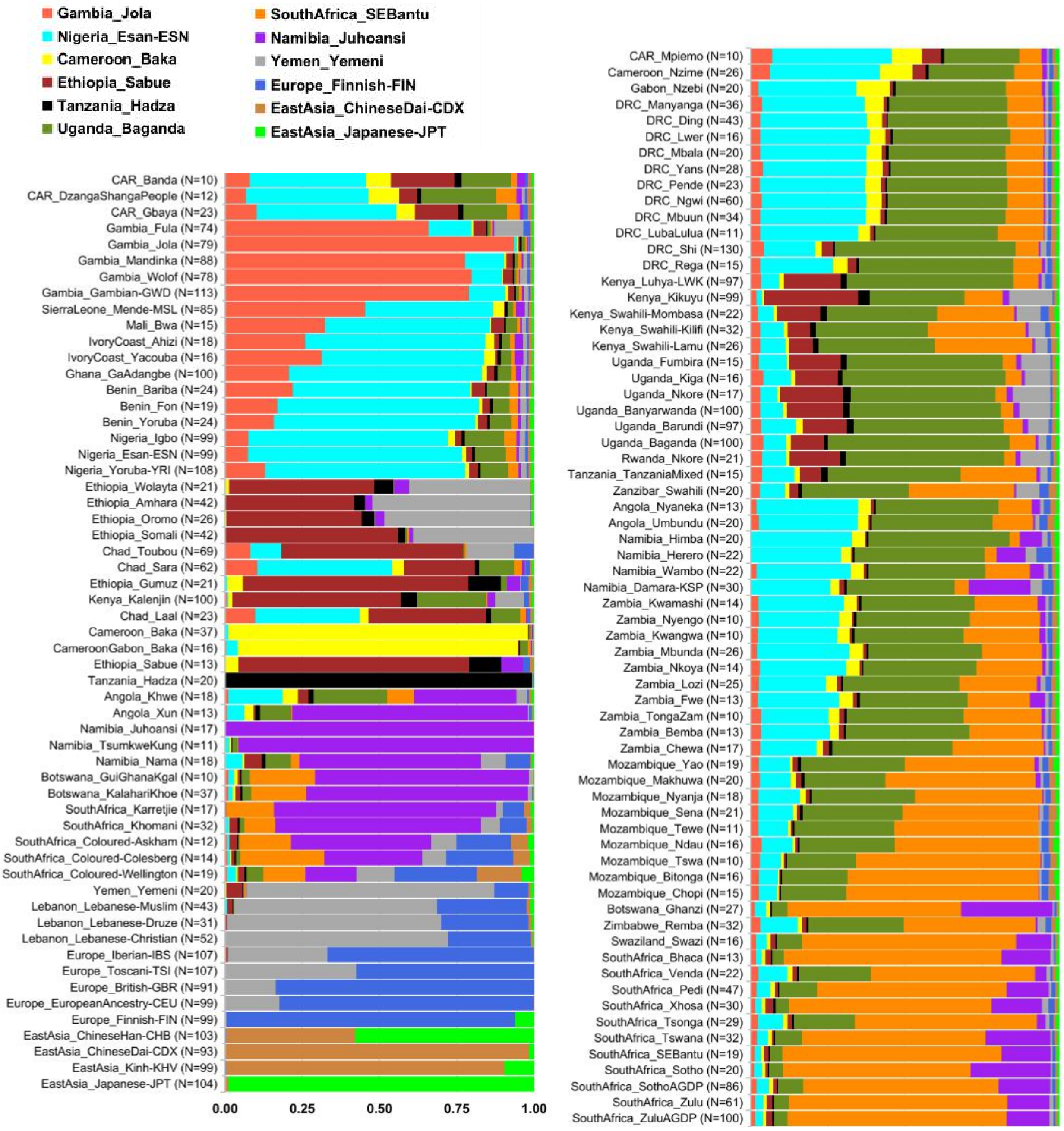


**Fig.S 3.3 | Bar plots of ADMIXTURE results for K=12.**
Panel figure showing average admixture results estimated using unsupervised ADMIXTURE plot at K=12 for all the populations included in the AfricanNeo dataset.

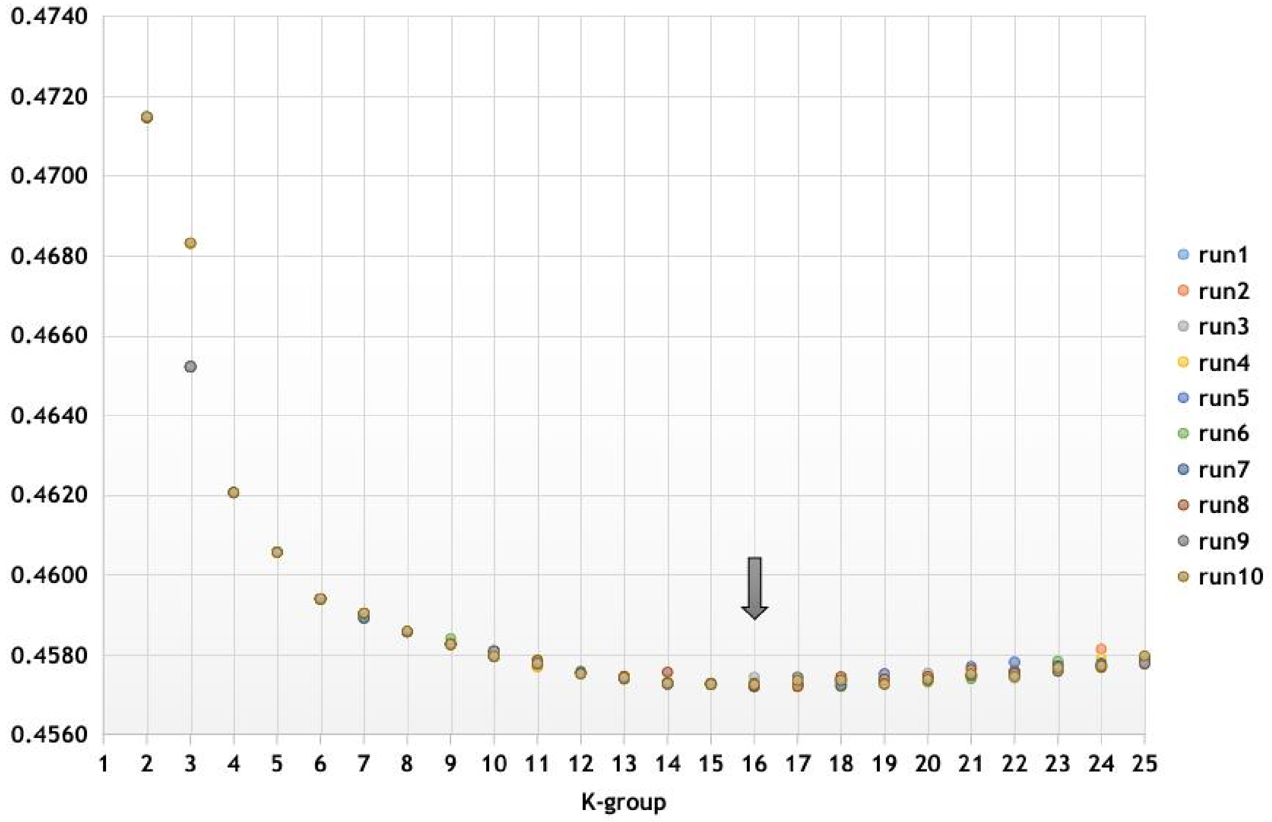


**Fig.S 3.4 | Cross-validation test from the ADMIXTURE analyses for each K-group.**
Figure showing cross-validation test from the ADMIXTURE analyses from K=2 to K=25, after using 10 independent runs with a random seed for each K-group. We highlighted with a black arrow the K-group with the lowest CV value (K=16). Each run from each K-group is one dot, and the dots from the same run have the same colour.

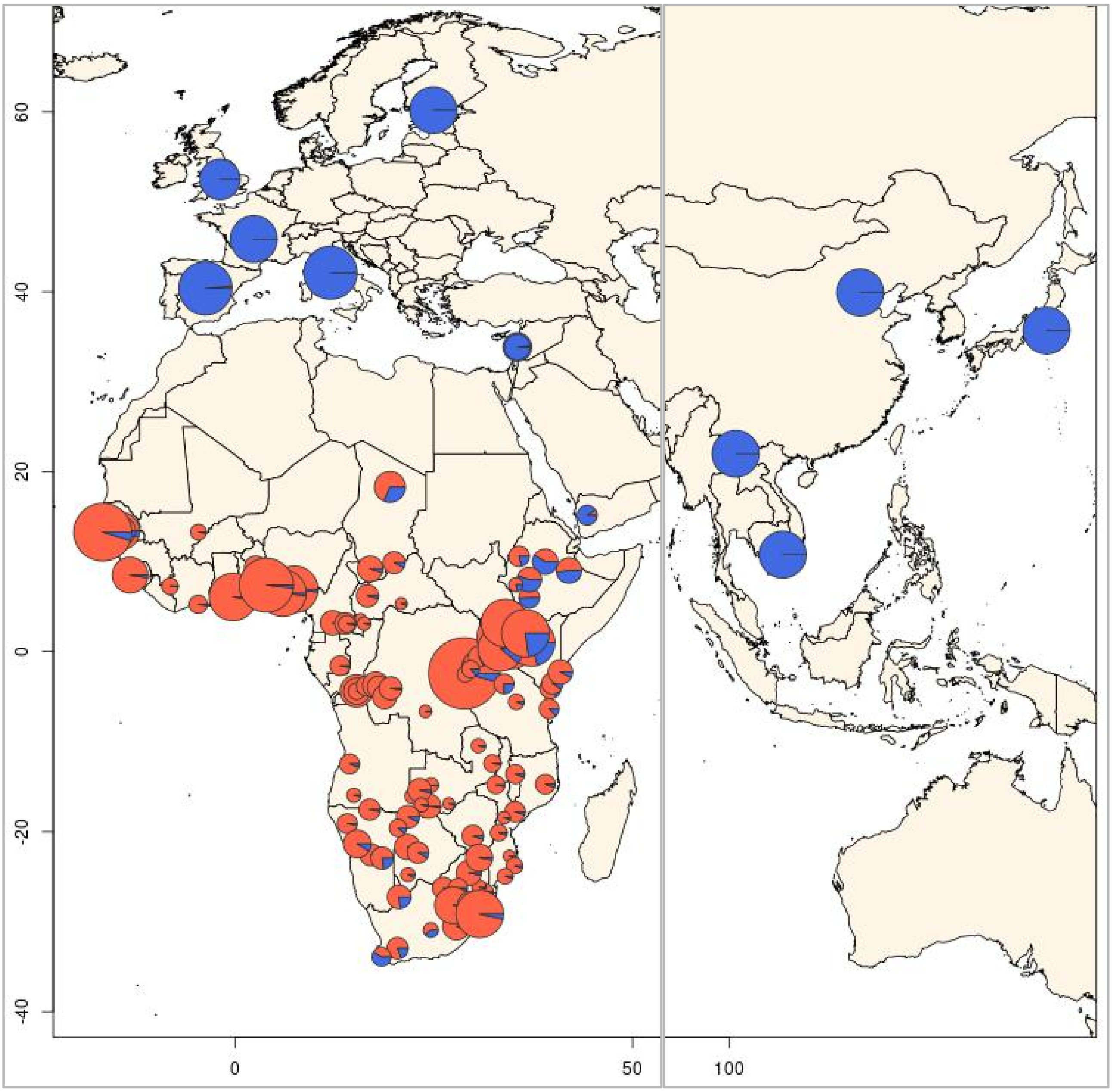


**Fig.S 3.5 | Pie charts of ADMIXTURE results for K=2.**
Panel figure showing unsupervised ADMIXTURE plot at K=2 for all the populations included in the AfricanNeo dataset. Results highlight the African-related component (in red) and the Eurasian-related component (in blue). The size of the pie charts is in relation to the sample size of each studied population. To better visualize the plot, central Asia was removed from the map.

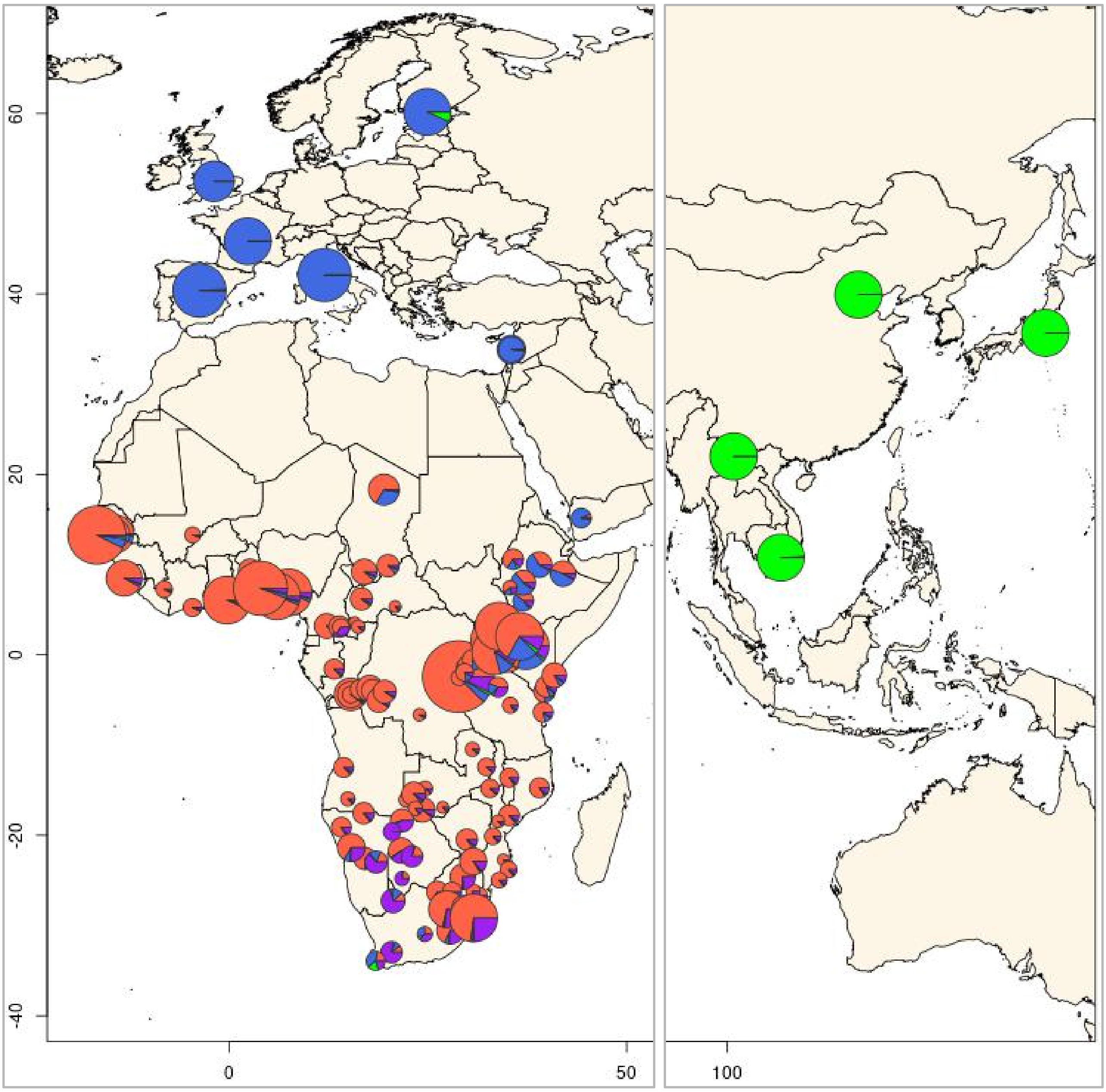


**Fig.S 3.6 | Pie charts of ADMIXTURE results for K=4.**
Panel figure showing unsupervised ADMIXTURE results at K=4 for all the populations included in the AfricanNeo dataset. Average admixture proportions of each African population are presented in **Table.S 3**. Results highlight the African-related component (in red), the hunter-gatherer-related component (in purple), the European-related component (in blue), and the East Asian-related component (in green; as expected in the Coloured populations in South Africa and the Finnish population). The size of the pie charts is in relation to the sample size of each studied population. Further details of the results of each individual in each population were included in **Fig.S 3.13**.

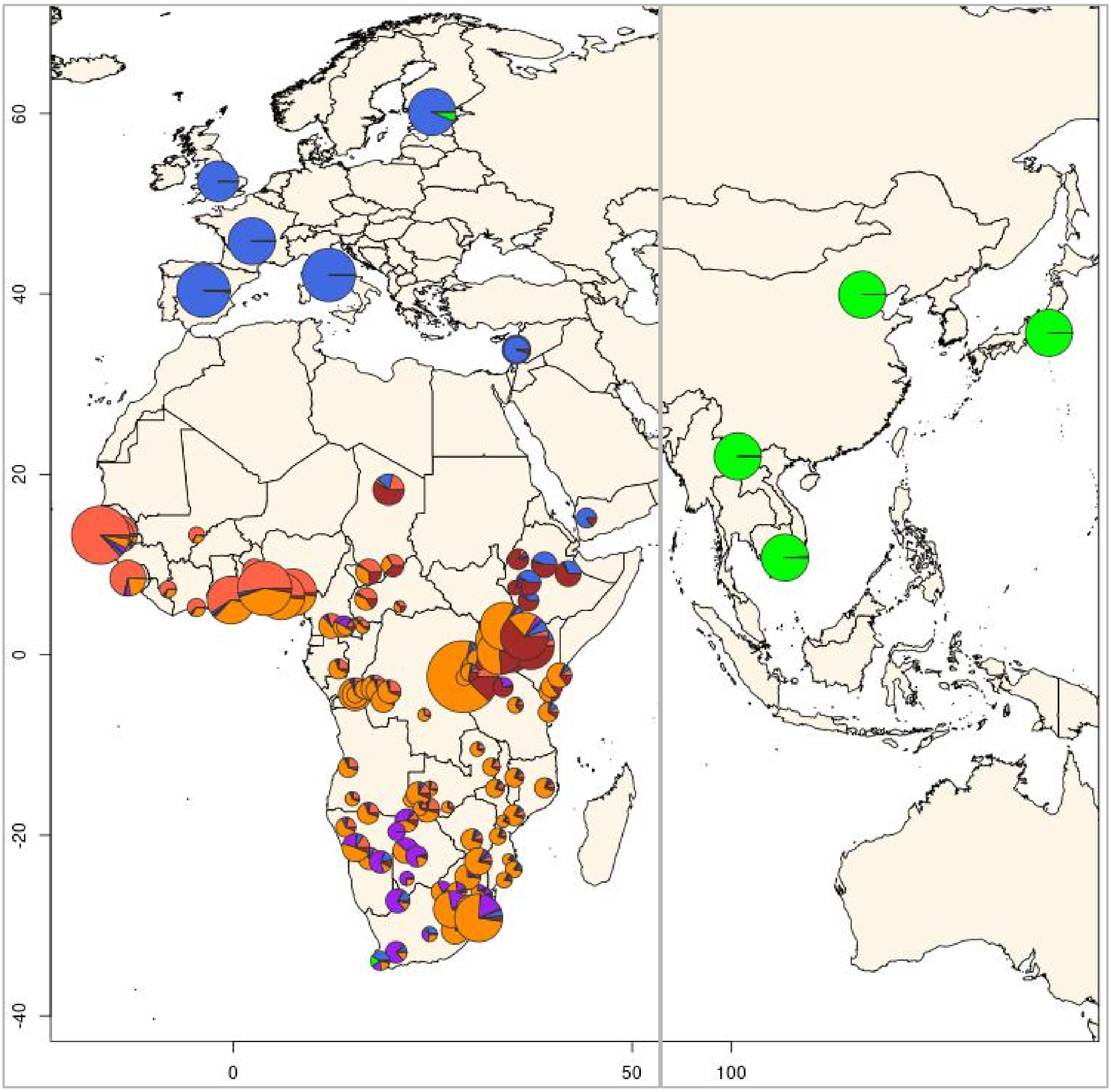


**Fig.S 3.7 | Pie charts of ADMIXTURE results for K=6.**
Panel figure showing unsupervised ADMIXTURE plot at K=6 for all the populations included in the AfricanNeo dataset. The size of the pie charts is in relation to the sample size of each studied population. To better understand the results, ADMIXTURE results at K=6 were also plotted the results in ternary diagrams, see **Fig.S 3.12**.

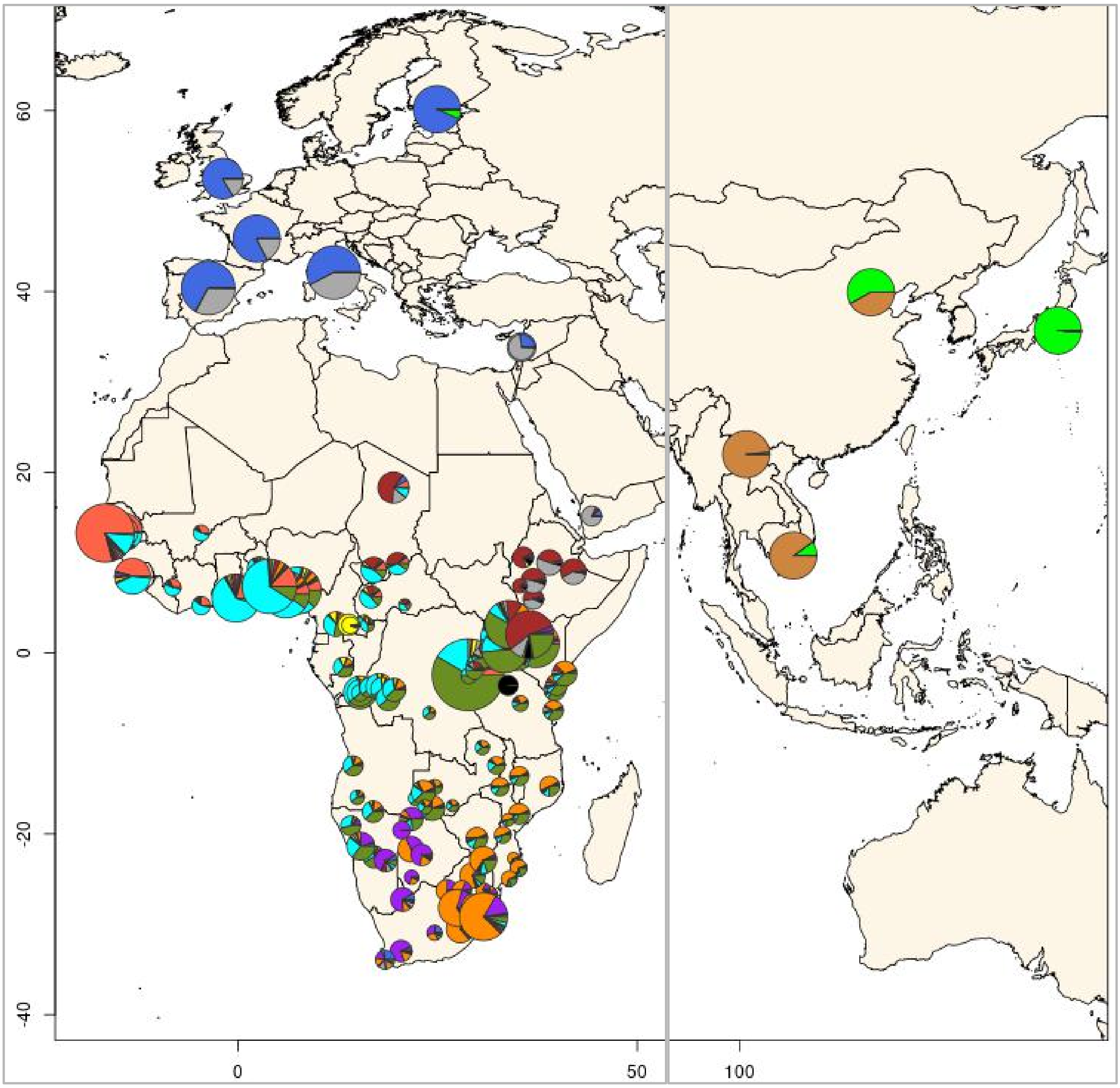


**Fig.S 3.8 | Pie charts of ADMIXTURE results for K=12.**
Panel figure showing unsupervised ADMIXTURE plot at K=12 for all the populations included in the AfricanNeo dataset. The size of the pie charts is in relation to the sample size of each studied population.

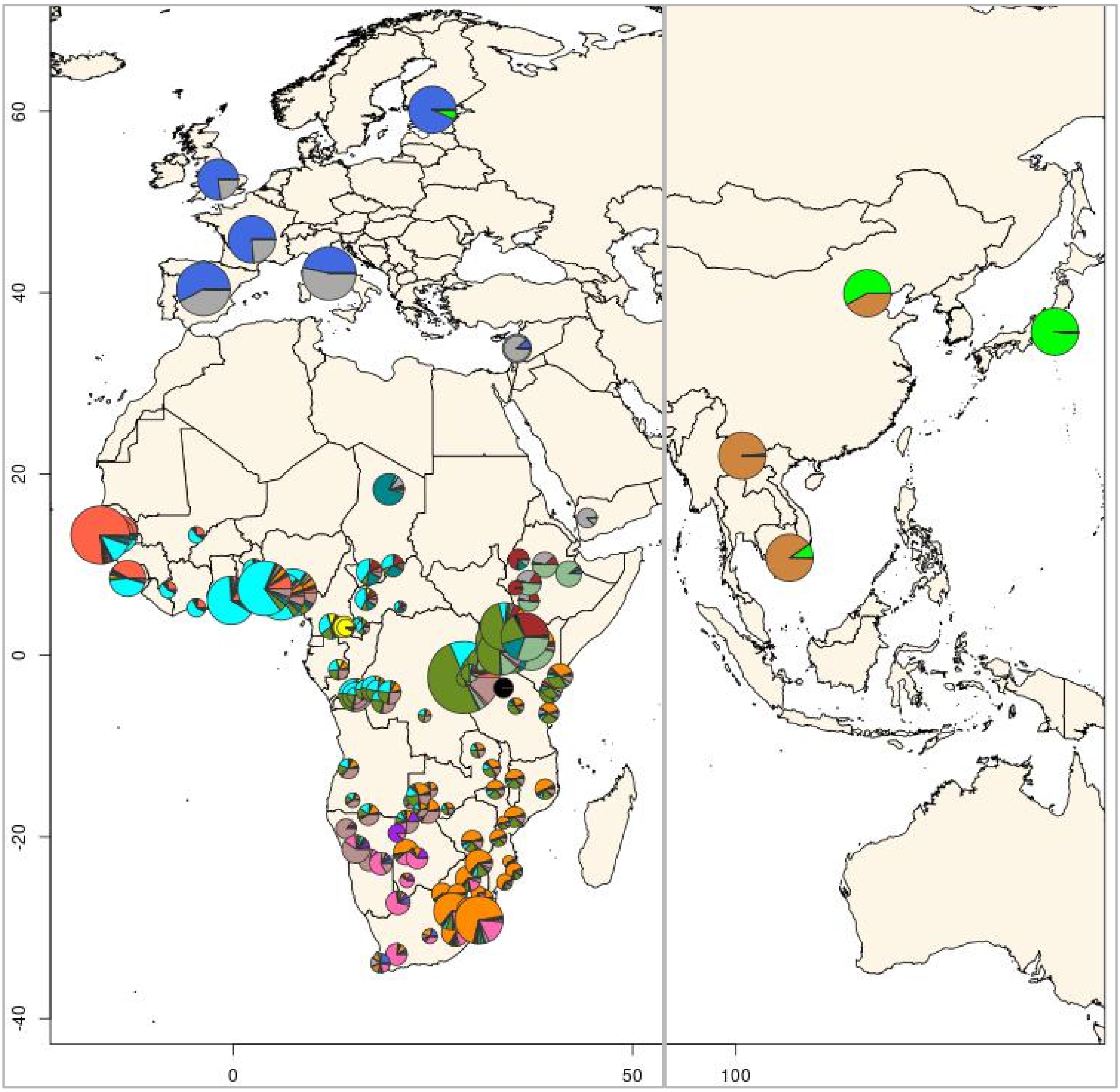


**Fig.S 3.9 | Pie charts of ADMIXTURE results for K=16.**
Unsupervised ADMIXTURE plot at K=16 for all the populations included in the AfricanNeo dataset. Further details of the results of each individual in each population were included in **Fig.S 3.11** and **Table.S 4**. The size of the pie charts is in relation to the sample size of each studied population.

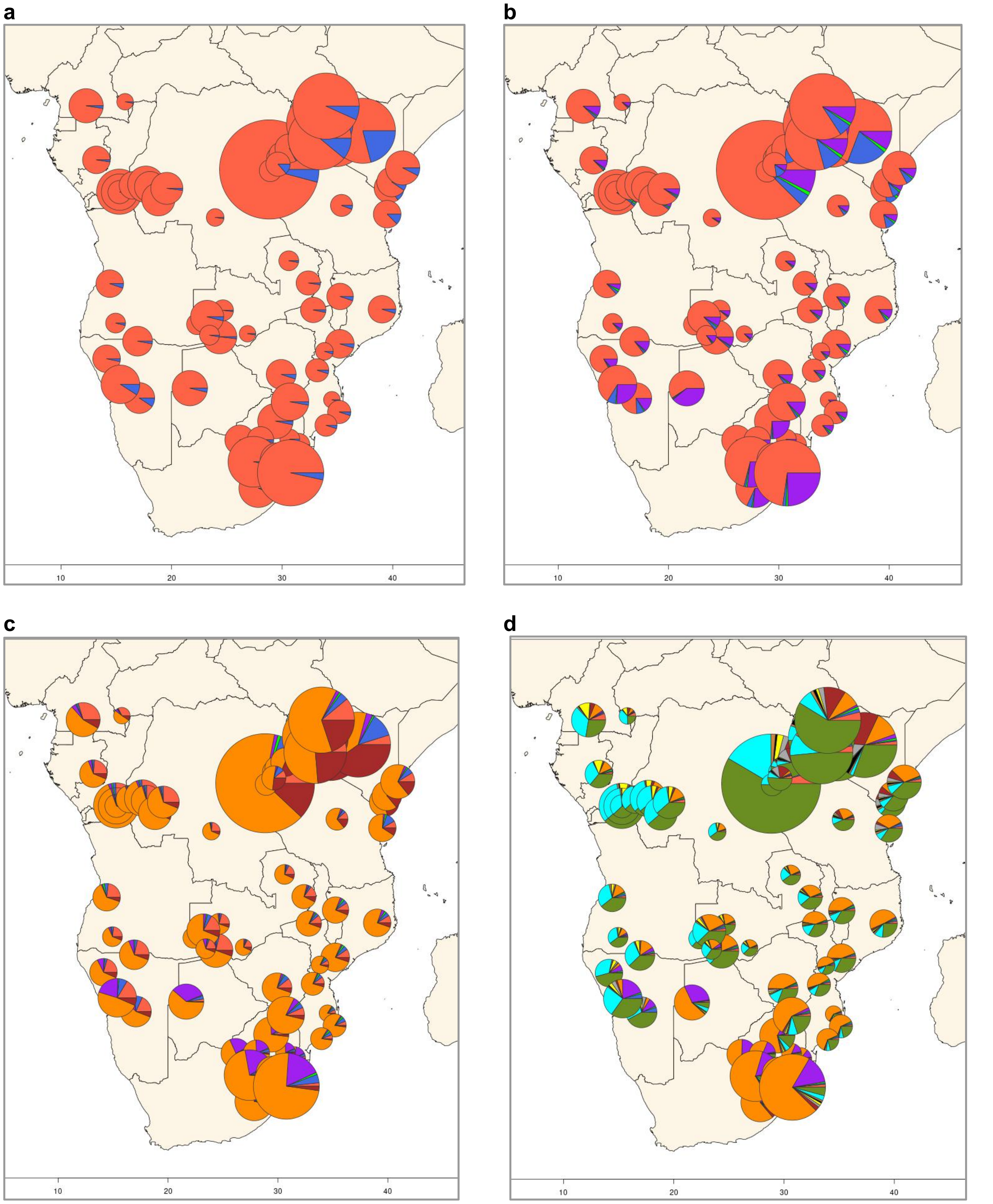


**Fig.S 3.10 | Pie charts of ADMIXTURE results for only studied BSP.**
Panel figure showing unsupervised ADMIXTURE results only for studied BSP for (**a**) K-group= 2; (**b**) K-group= 4; (**c**) K-group= 6; and (**d**) K-group= 12. Results for BSP and all comparative populations were included in previous figures (see **Fig.S 3.5–3.8**).

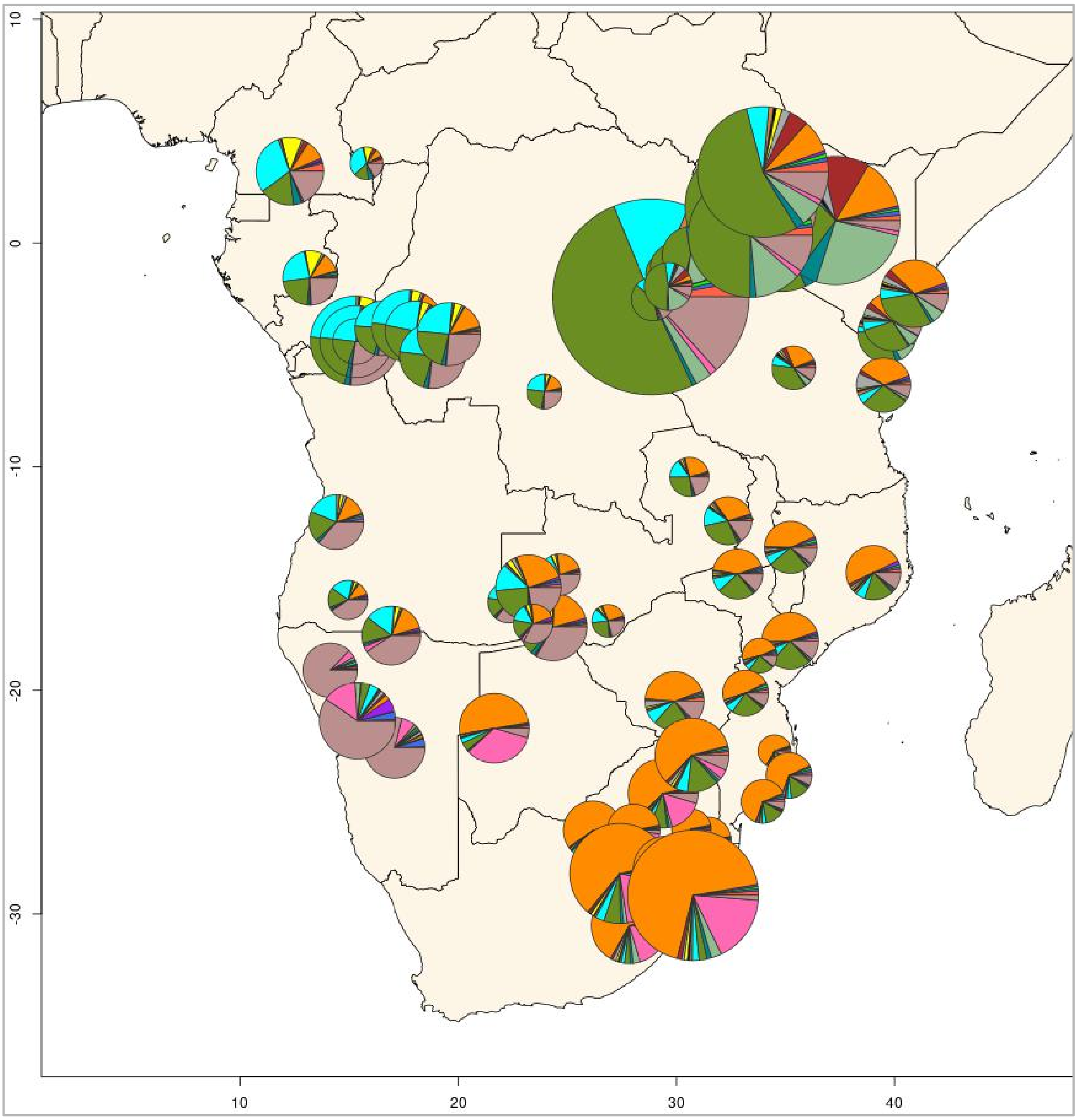


**Fig.S 3.11 | Pie charts of ADMIXTURE results for K=16 only for BSP.**
Panel figure showing unsupervised ADMIXTURE plot at K=16 only for BSP included in the AfricanNeo dataset, and comparative populations were not included in this plot. Results for BSP and all comparative populations were included in **Fig.S 3.9**.

a

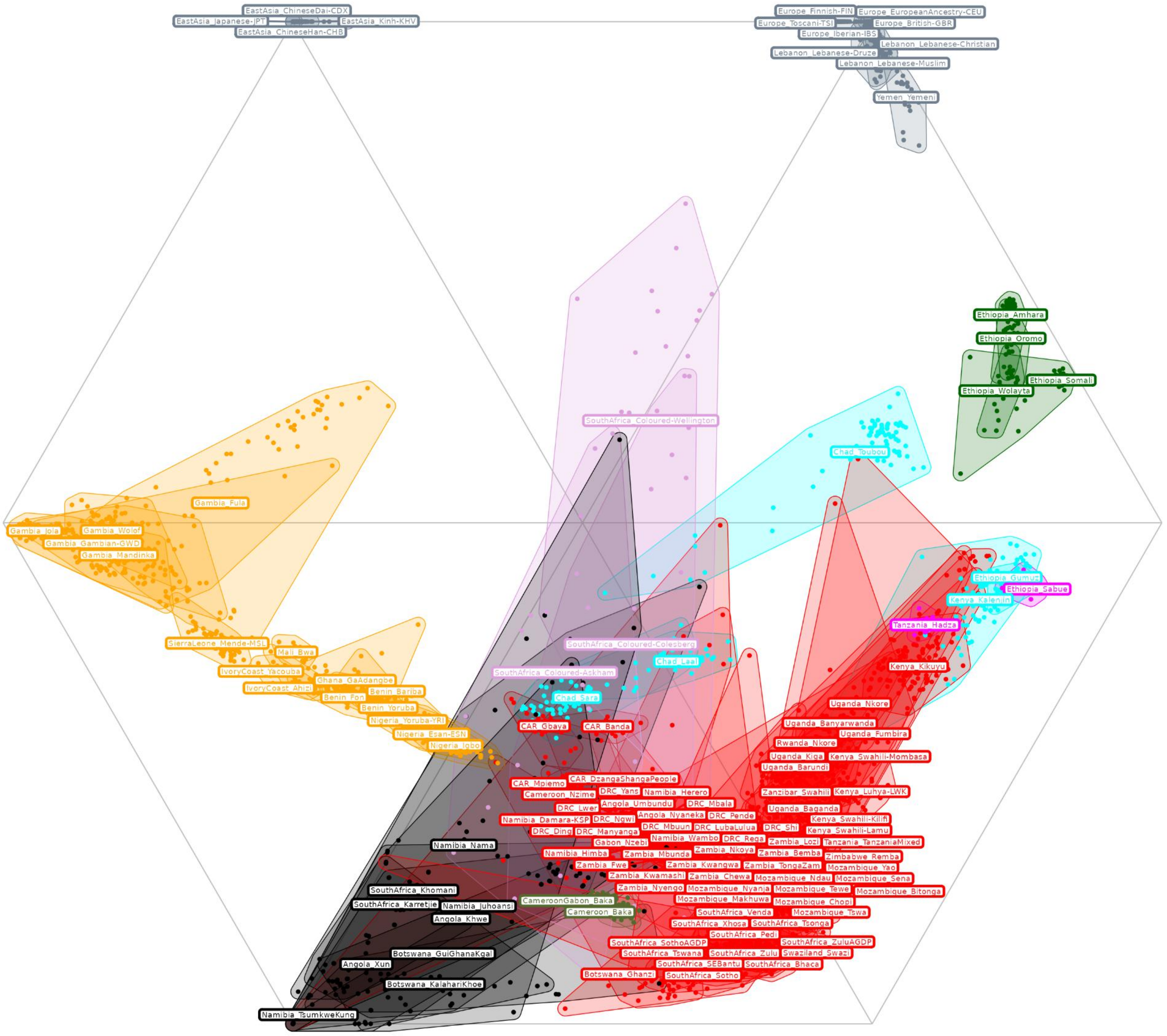
EastAsia_ChineseDai-CDX
EastAsia_Japanese-JPT
EastAsia_Kinh-KHV
EastAsia_ChineseHan-CHB
Europe_Finnish-FIN
Europe_EuropeanAncestry-CEU
Europe_Toscani-TSI
Europe_British-GBR
Europe_Iberian-IBS
Lebanon_Lebanese-Christian
Lebanon_Lebanese-Druze
Lebanon_Lebanese-Muslim
Yemen_Yemeni
Ethiopia_Amhara
Ethiopia_Oromo
Ethiopia_Somali
Ethiopia_Wolayta
SouthAfrica_Coloured-Wellington
Chad_Toubou
Gambia_Fula
Gambia_Jola
Gambia_Wolof
Gambia_Gambian-GWD
Gambia_Mandinka
Ethiopia_Gumuz
Ethiopia_Sabue
Kenya_Kalenjin
Tanzania_Hadza
SouthAfrica_Coloured-Colesberg
SierraLeone_Mende-MSL
Mali_Bwa
Chad_Laal
Kenya_Kikuyu
IvoryCoast_Yacouba
SouthAfrica_Coloured-Askham
Ghana_GaAdangbe
IvoryCoast_Ahizi
Benin_Bariba
Benin_Fon
Chad_Sara
Uganda_Nkore
Benin_Yoruba
Nigeria_Yoruba-YRI
Uganda_Banyarwanda
CAR_Gbaya
CAR_Banda
Nigeria_Esan-ESN
Uganda_Fumbira
Nigeria_Igbo
Rwanda_Nkore
Uganda_Kiga
Kenya_Swahili-Mombasa
Uganda_Barundi
CAR_DzangaShangaPeople
CAR_Mpiemo
Cameroon_Nzime
DRC_Yans
Namibia_Herero
Zanzibar_Swahili
Kenya_Luhya-LWK
Angola_Umbundu
DRC_Mbala
DRC_Lwer
Uganda_Baganda
Angola_Nyaneka
DRC_Pende
Kenya_Swahili-Kilifi
Namibia_Damara-KSP
DRC_Ngwi
DRC_Mbuun
DRC_LubaLulua
DRC_Shi
Kenya_Swahili-Lamu
DRC_Ding
DRC_Manyanga
Namibia_Wambo
DRC_Rega
Zambia_Lozi
Tanzania_TanzaniaMixed
Gabon_Nzebi
Namibia_Nama
Zambia_Nkoya
Namibia_Himba
Zambia_Mbunda
Zambia_Bemba
Zimbabwe_Remba
Zambia_Fwe
Zambia_Kwangwa
Zambia_TongaZam
Mozambique_Yao
Zambia_Kwamashi
Zambia_Chewa
Mozambique_Ndau
Mozambique_Sena
SouthAfrica_Khomani
Zambia_Nyengo
Mozambique_Nyanja
Mozambique_Tewe
Mozambique_Bitonga
CameroonGabon_Baka
Mozambique_Makhuwa
Mozambique_Chopi
SouthAfrica_Karretjie
Namibia_Juhoansi
Cameroon_Baka
SouthAfrica_Venda
Mozambique_Tswa
Angola_Khwe
SouthAfrica_Xhosa
SouthAfrica_Tsonga
SouthAfrica_Pedi
SouthAfrica_SothoAGDP
SouthAfrica_ZuluAGDP
Botswana_GuiGhanaKgal
SouthAfrica_Tswana
SouthAfrica_Zulu
Swaziland_Swazi
Angola_Xun
SouthAfrica_SEBantu
SouthAfrica_Bhaca
Botswana_Ghanzi
SouthAfrica_Sotho
Botswana_KalahariKhoe
Namibia_TsumkweKung

**b**

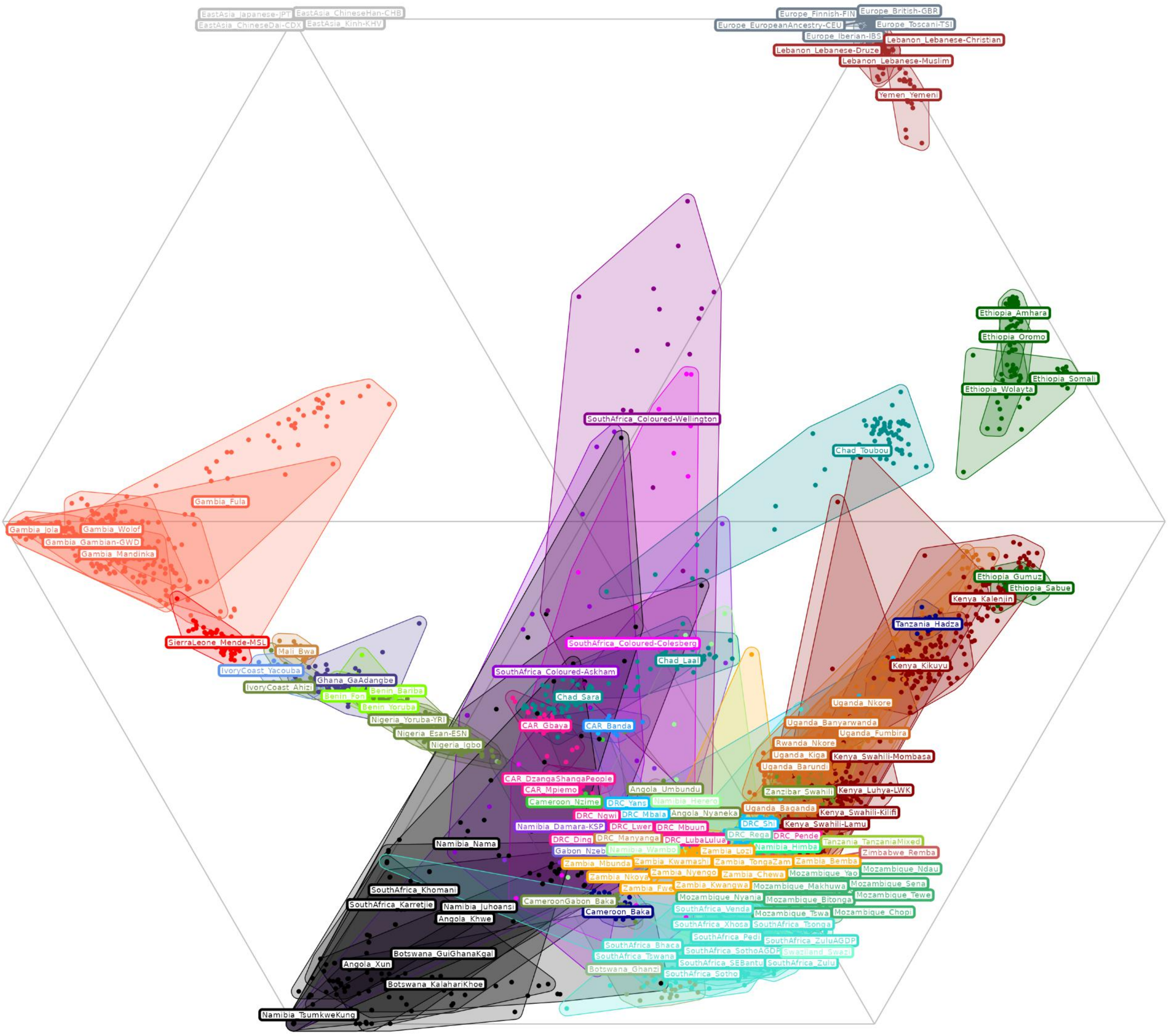


**Fig.S 3.12 | Ternary diagram of ADMIXTURE results at K=6.**
Figure showing a ternary diagram in hexagonal shape for ADMIXTURE results at K=6 (**Fig.S 3.7**) in (**a**) each group and (**b**) each population included in the AfricanNeo dataset. Each corner of the hexagon corresponds to each ancestry as follows: East Asian-related ancestry on the top-left corner; European-related ancestry on the top-right corner; eastern African-related ancestry on the central-right corner; Bantu-speaking-related ancestry on the bottom-right corner; Khoisan-speaking-related ancestry on the bottom-left corner; and western African-related ancestry on the central-left corner. Each individual is one dot from each population (shadow area), and individuals close to each corner suggest high ancestry proportions for that ancestry.

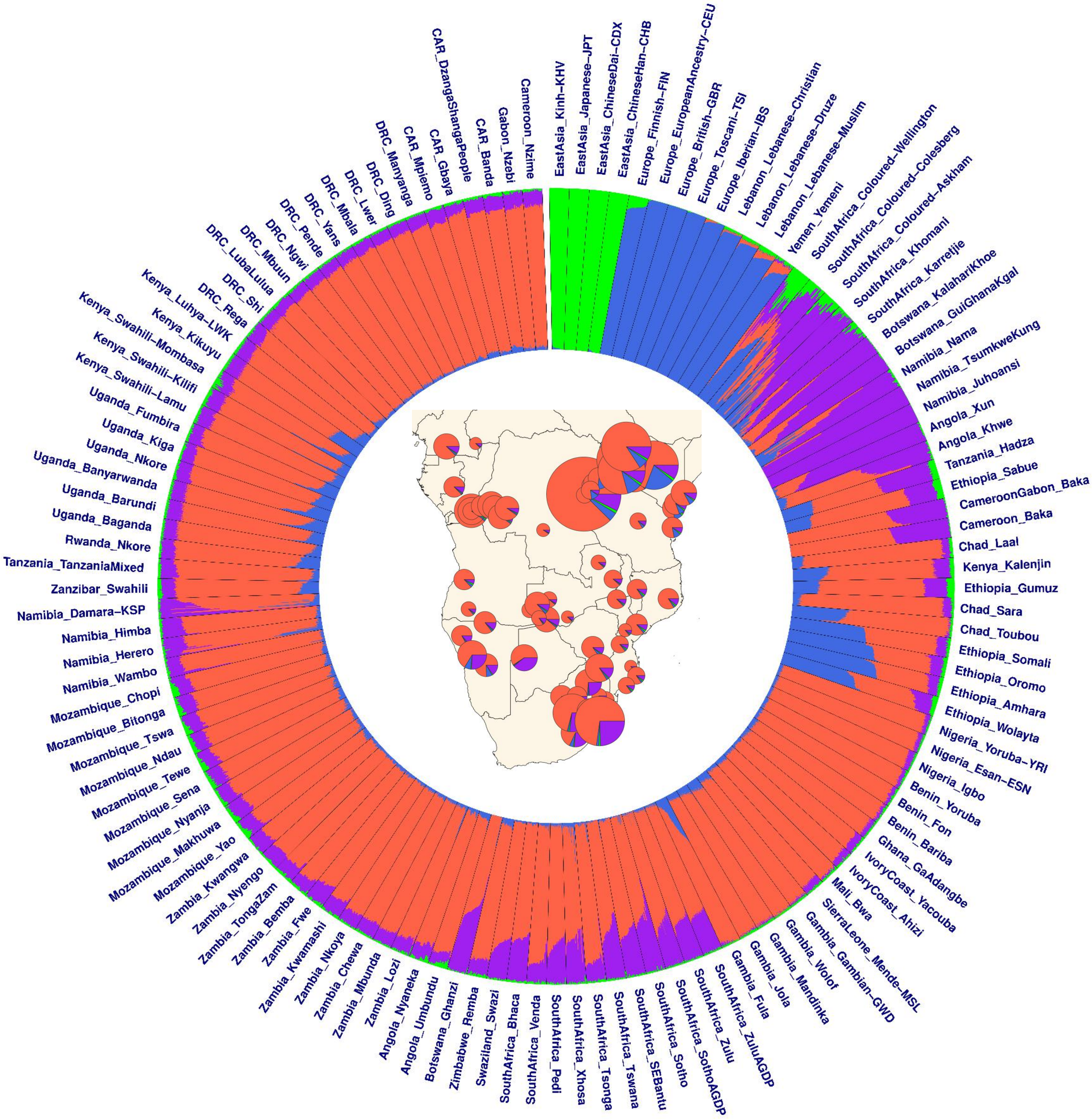


**Fig.S 3.13 | Bar plots of ADMIXTURE results for K=4.**
Panel figure showing average admixture proportions estimated using unsupervised ADMIXTURE analyses at K=4 on the basis of all the populations included in the AfricanNeo dataset (**Table. S3**). BSP are in the left part of the plot while comparative populations are in the right part of the plot. The central part of the figure included the map presented in **Fig.S 3.10a**.

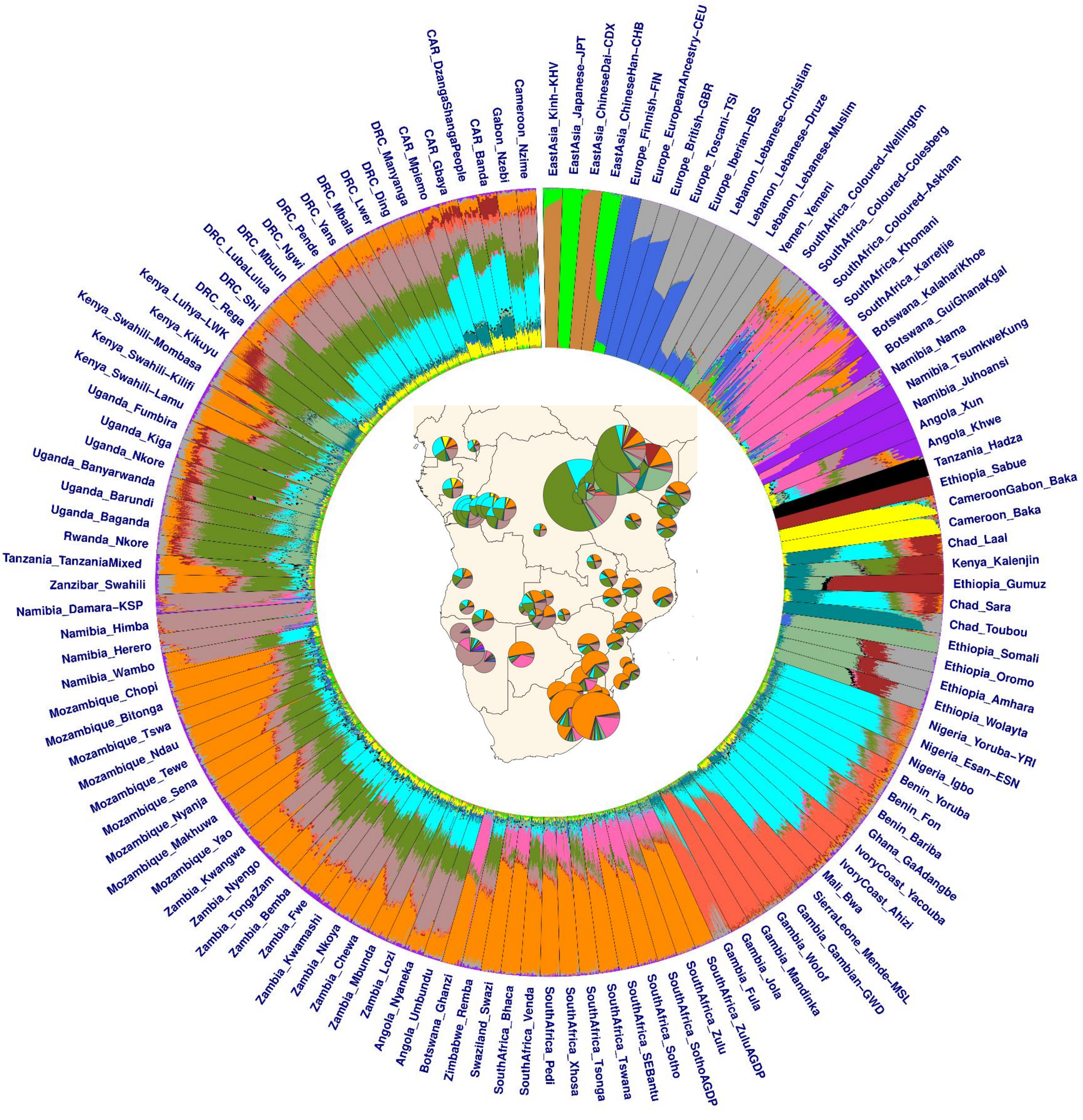


**Fig.S 3.14 | Bar plots of ADMIXTURE results for K=16.**

Panel figure showing average admixture proportions estimated using unsupervised ADMIXTURE analyses at K=16 on the basis of all the populations included in the AfricanNeo dataset (**Table. S4**). The colors of each K-group are matching the same colors in **Fig.S 3.9**. To better visualize the figure, the width of each population was set to be equal regardless of its sample size using AncestryPainter. BSP are in the left part of the plot while comparative populations are in the right part of the plot. The central part of the figure included the map presented in **Fig.S 3.10d**.

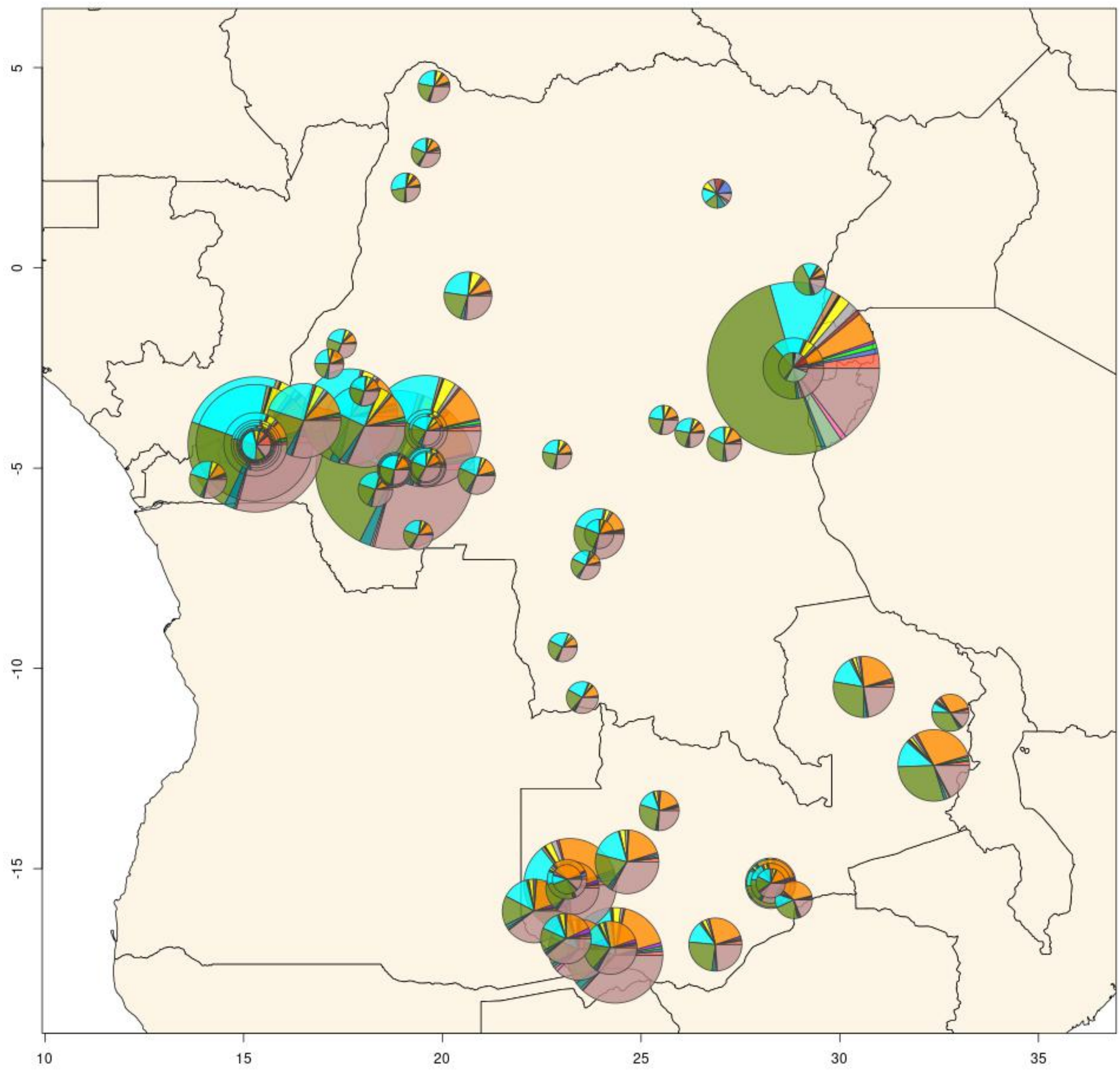


**Fig.S 3.15 | Pie charts of ADMIXTURE results for K=16 only for DRC and Zambia.**
Panel figure showing unsupervised ADMIXTURE plot at K=16 only for all the genotyped BSP from DRC and Zambia (**Table.S 4**). The smallest pie charts represent populations with only one sample while the biggest pie chart represents the Shi population with a sample size of 130 samples.

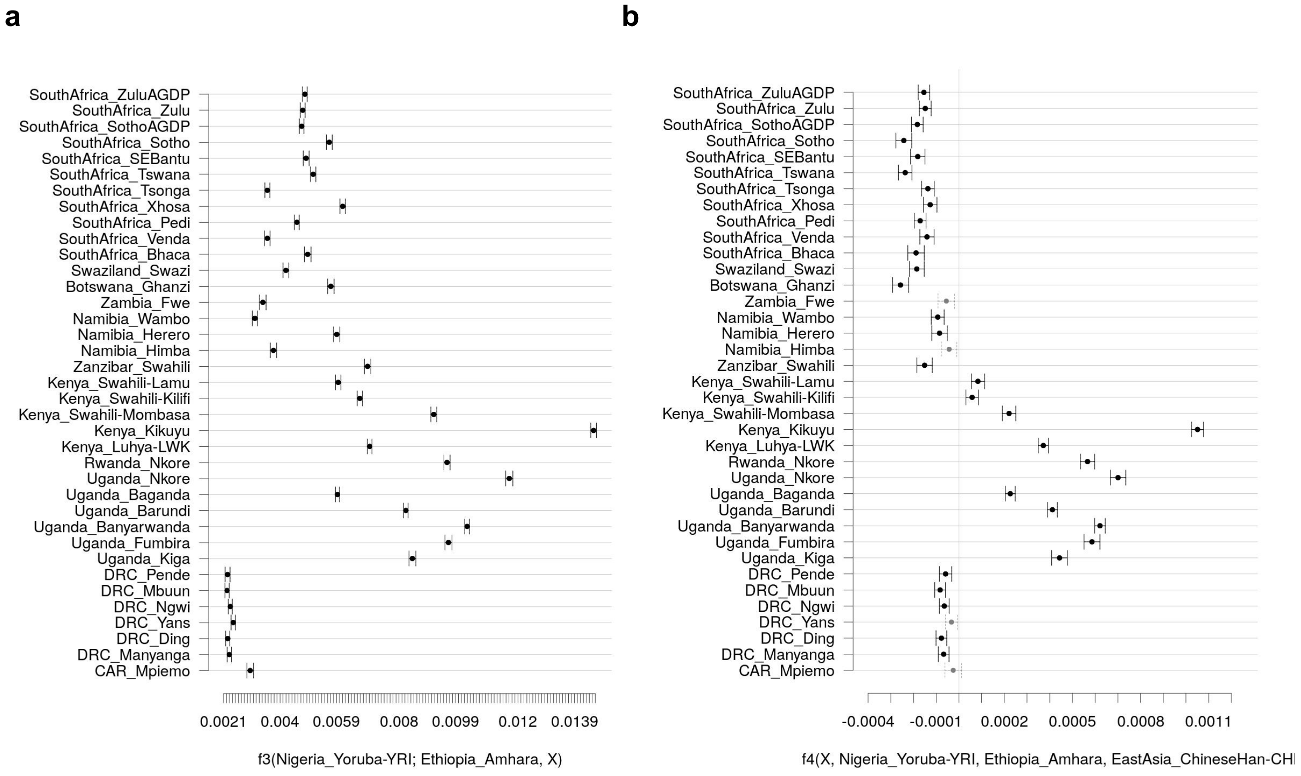


**Fig.S 3.16 | Test of Afro-Asiatic admixture in BSP estimated using f3- and f4-statistics.**
Positive values of (**a**) f3-statistics in the form f3(Yoruba; Amhara, *Target*) and (**b**) f4-statistics in the form f4(*Target*, Yoruba; Amhara, CHB) indicate increasing genetic affinity between the Amhara population in Ethiopia and each of the BSP indicated in the Y-axis. Bars indicate the standard error of the mean. Values of the f4-statistic significantly higher than zero (*P*-value < 0.05) are indicated with black color. BSP are in the same order in f3- and f4-statistics plots.

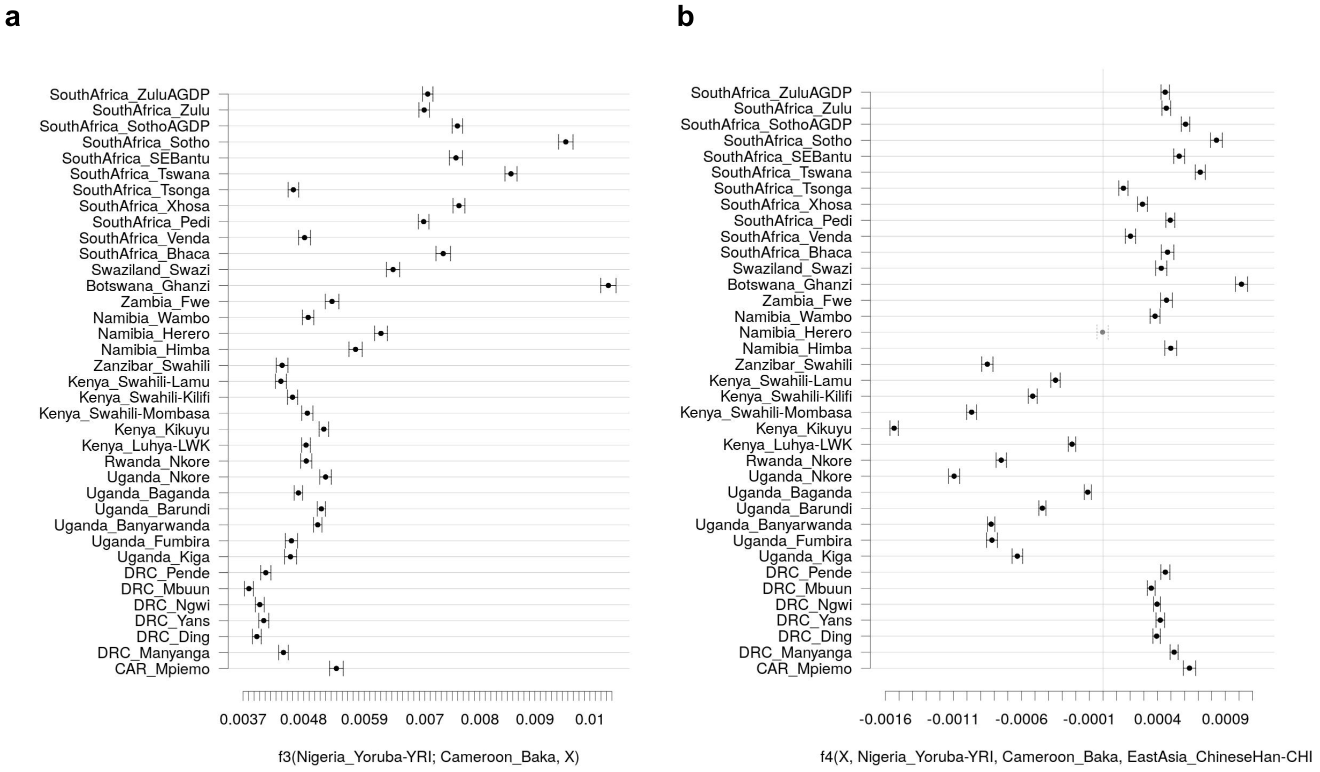


**Fig.S 3.17 | Test of western RGH admixture in BSP estimated using f3- and f4-statistics.**
Positive values of (**a**) f3-statistics in the form f3(Yoruba; Baka, *Target*) and (**b**) f4-statistics in the form f4(*Target*, Yoruba; Baka, CHB) indicate increasing genetic affinity between the wRHG Baka population to each of the studied BPS indicated in the Y-axis. Bars indicate the standard error of the mean value. Values of the statistics significantly different from zero (*P*-value <0.05) are indicated with black color. BSP are in the same order in the Y-axis of both plots. Note that two groups of BSP show affinity with western RHG, western and southern African BSP. This most likely reflects the hunter-gatherer ancestry present in southern African BSP (see Fig. S4.3). Differential affinities for western and southern African BSP are visible in Fig. S3.19.

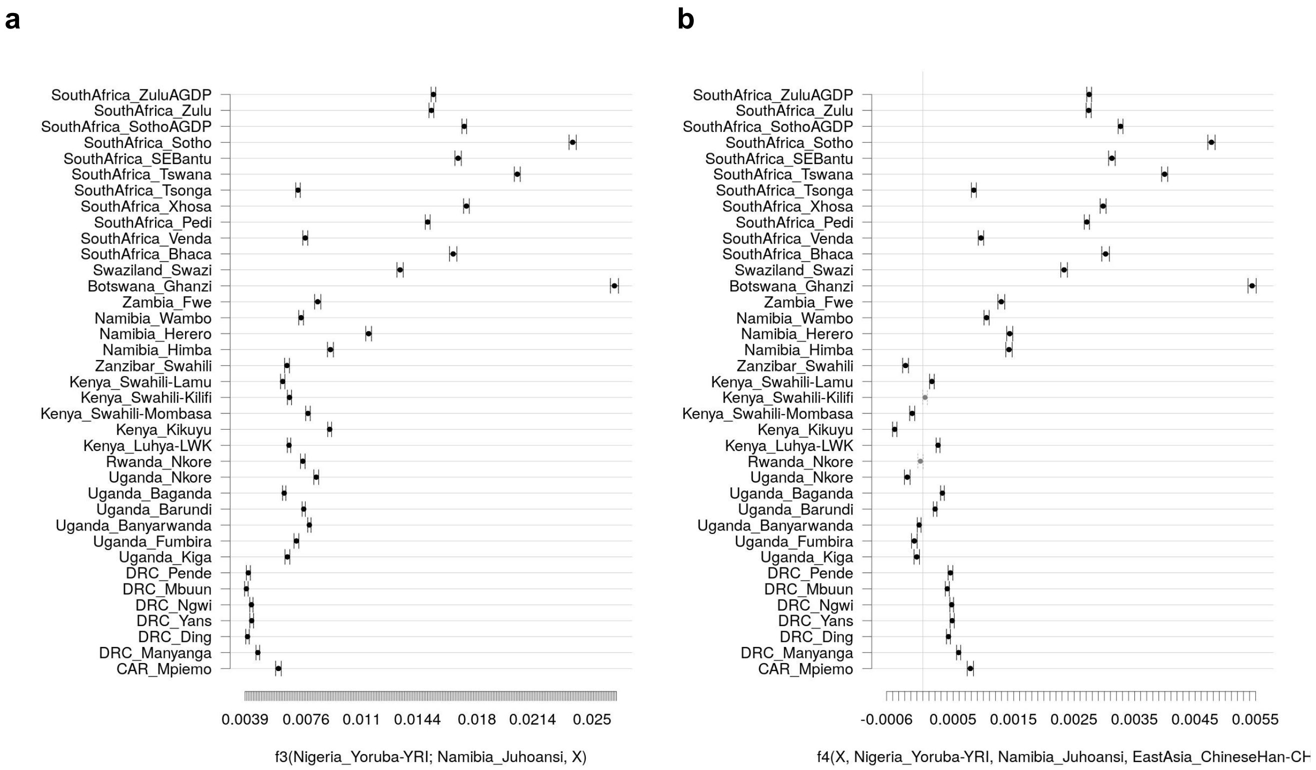


**Fig.S 3.18 | Test of Khoe-San admixture in BSP estimated using f3- and f4-statistics.**
Positive values of (**a**) f3-statistics in the form f3(Yoruba; Ju-hoansi, *Target*) and (**b**) f4-statistics in the form f4(*Target*, Yoruba; Ju-hoansi, CHB) indicate increasing genetic affinity between the Ju-hoansi population in Namibia to each of the BSP indicated in the Y-axis. Bars indicate the standard error of the mean. Values of the statistics significantly higher than zero (*P*-value < 0.05) are indicated with black color. BSP are in the same order in f3 and f4 plots.

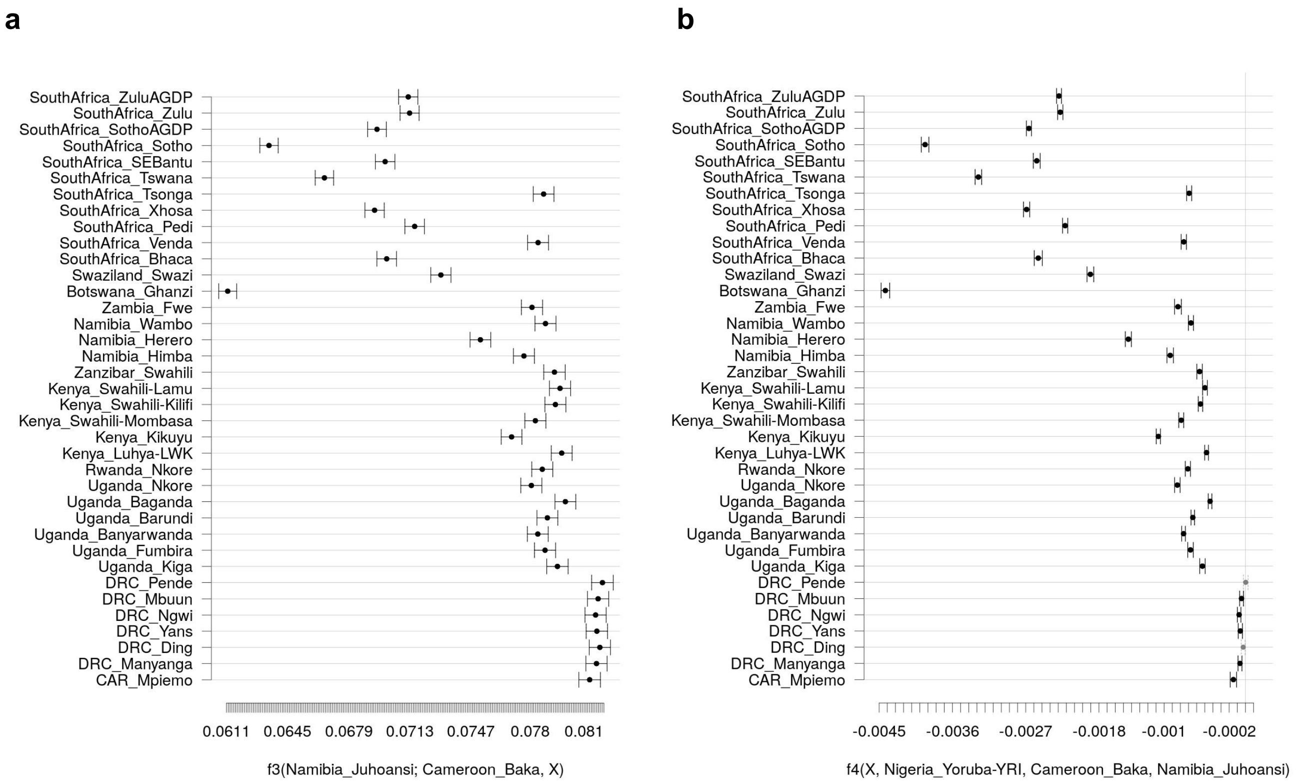


**Fig.S 3.19 | Tests of hunter-gatherer admixture in BSP estimated using f3- and f4-statistics.** To disentangle the differential contribution of different hunter-gatherer groups, we performed f3 tests in the form f3(Ju-hoansi; Baka, *Target*) (**a**) and f4 tests in the form f4(*Target*, Yoruba; Baka,Ju-hoansi) (**b**). Both tests indicate that western BSP have closer affinity to Baka groups than southern African BSP. Bars indicate the standard error of the mean. Values of the statistics significantly different from zero (*P*-value < 0.05) are indicated with black color. BSP are in the same order in f3 and f4 plots.

### 4- Ancestry-specific analyses

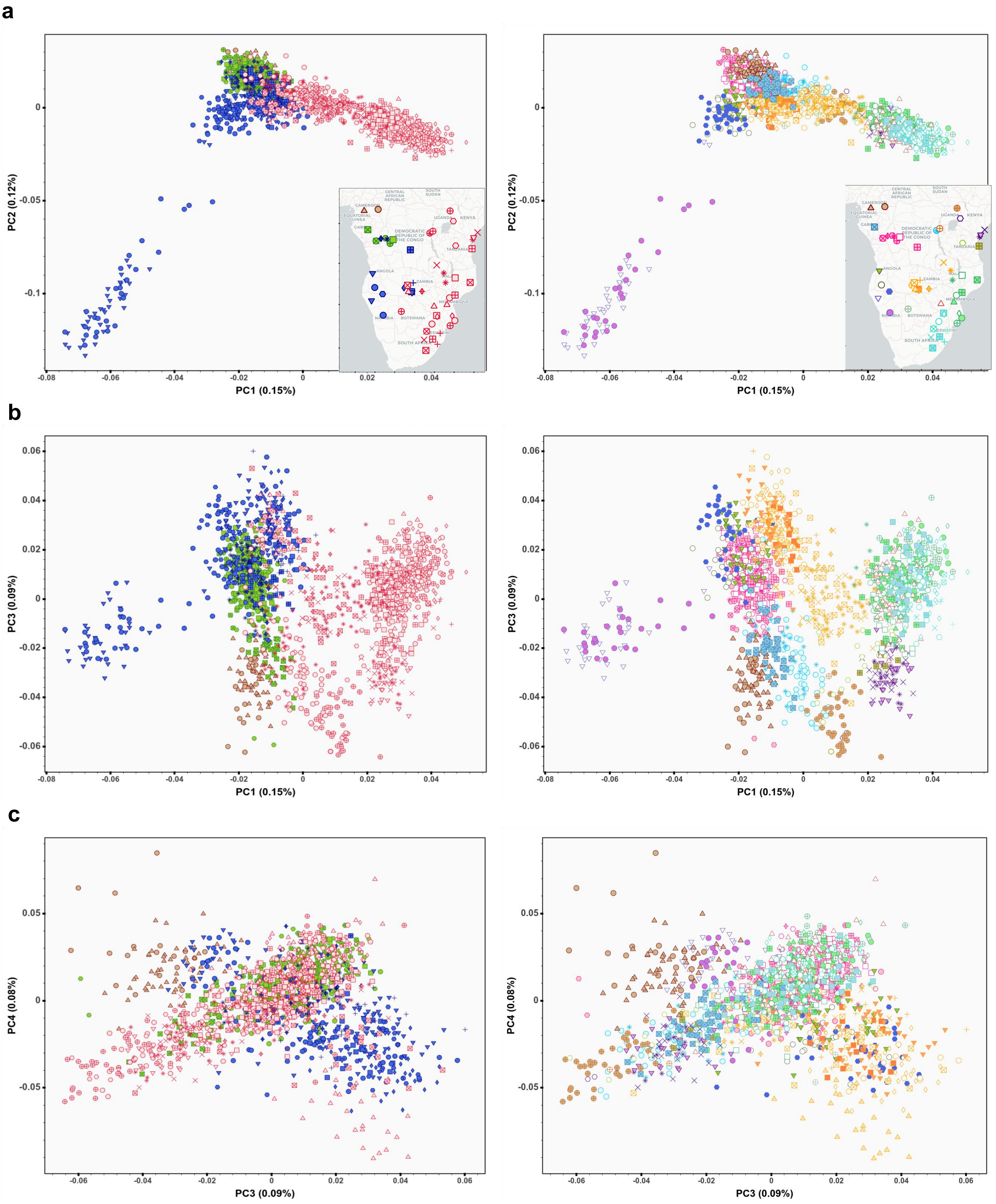


**Fig.S 4.1 | Ancestry-specific (AS-)PCA for the masked Only-BSP dataset.**
Figure showing AS-PCA plot of BSP included in the AfricanNeo dataset after masking and imputation of populations with 70% WCA-related ancestry, without using the procrustes approach. PC projections obtained for each group (left column) and for each population (right column) between: (**a**) PC1 vs PC2; (**b**) PC1 vs PC3; and (**c**) PC3 vs PC4. interactive plots were included ().

Legends and maps for each group (top) and each population (bottom) in **Fig.S 4.4.**

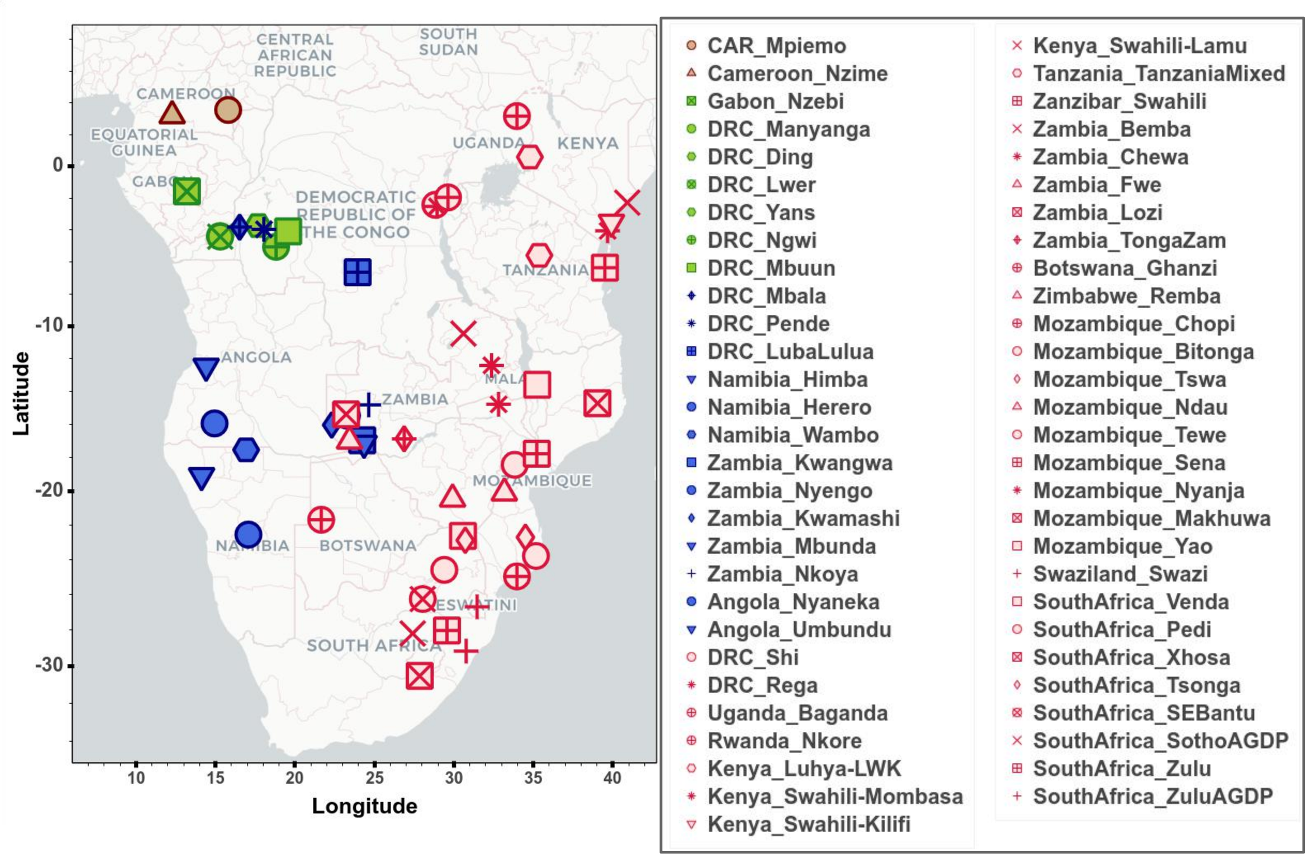


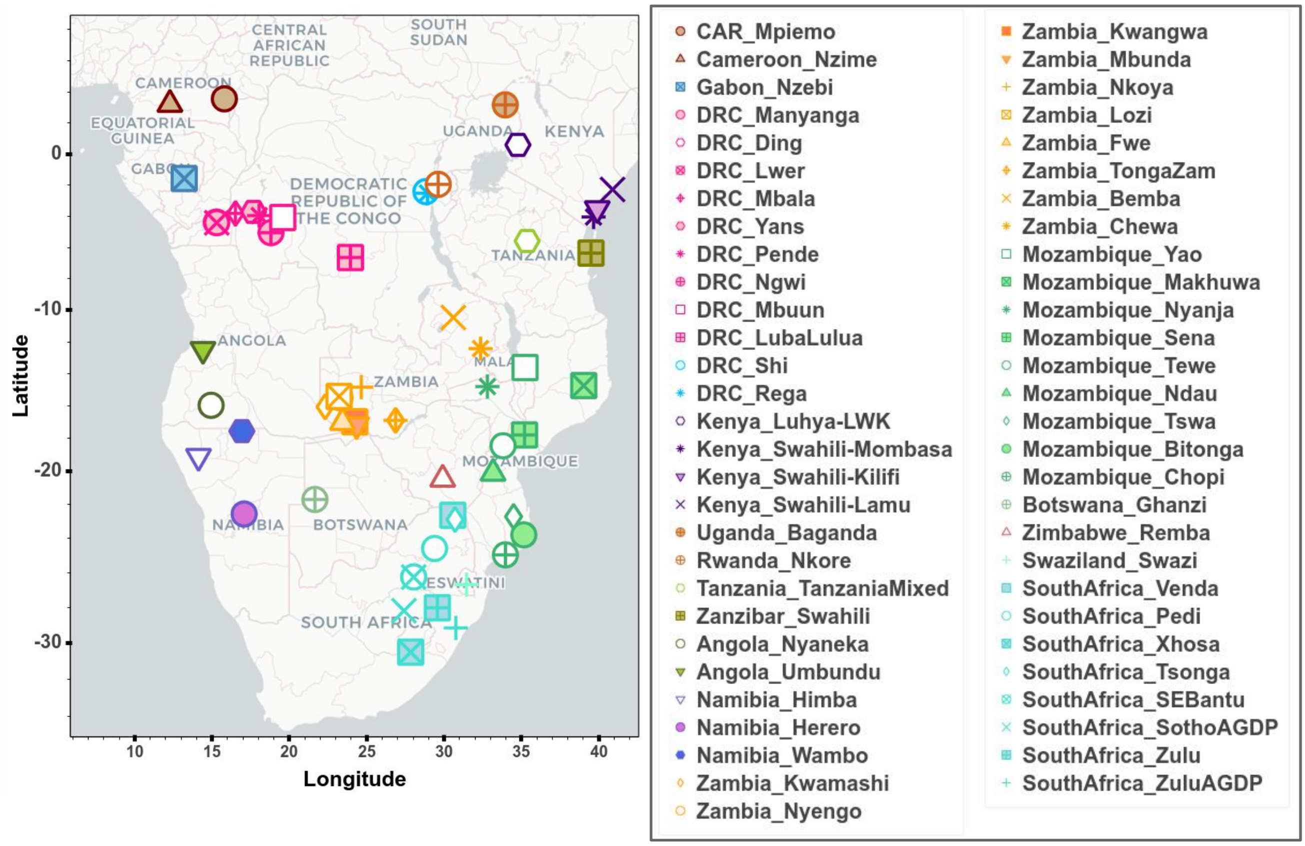

**a**

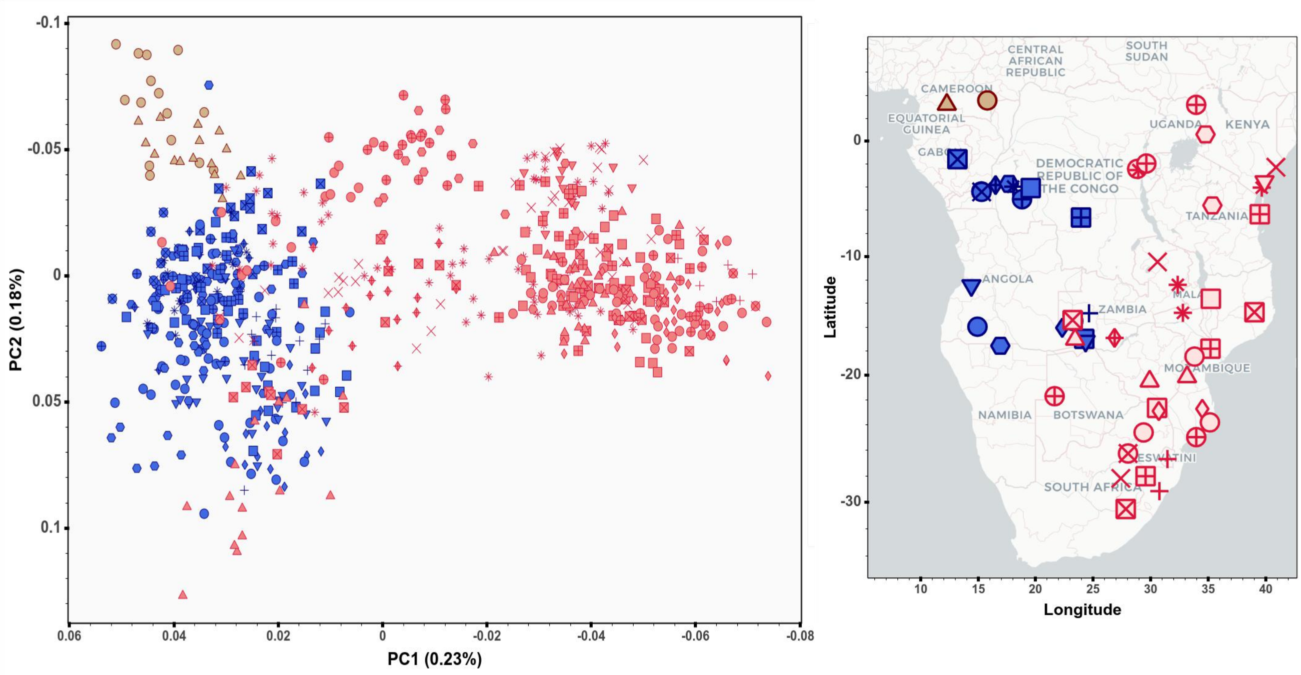


**b**

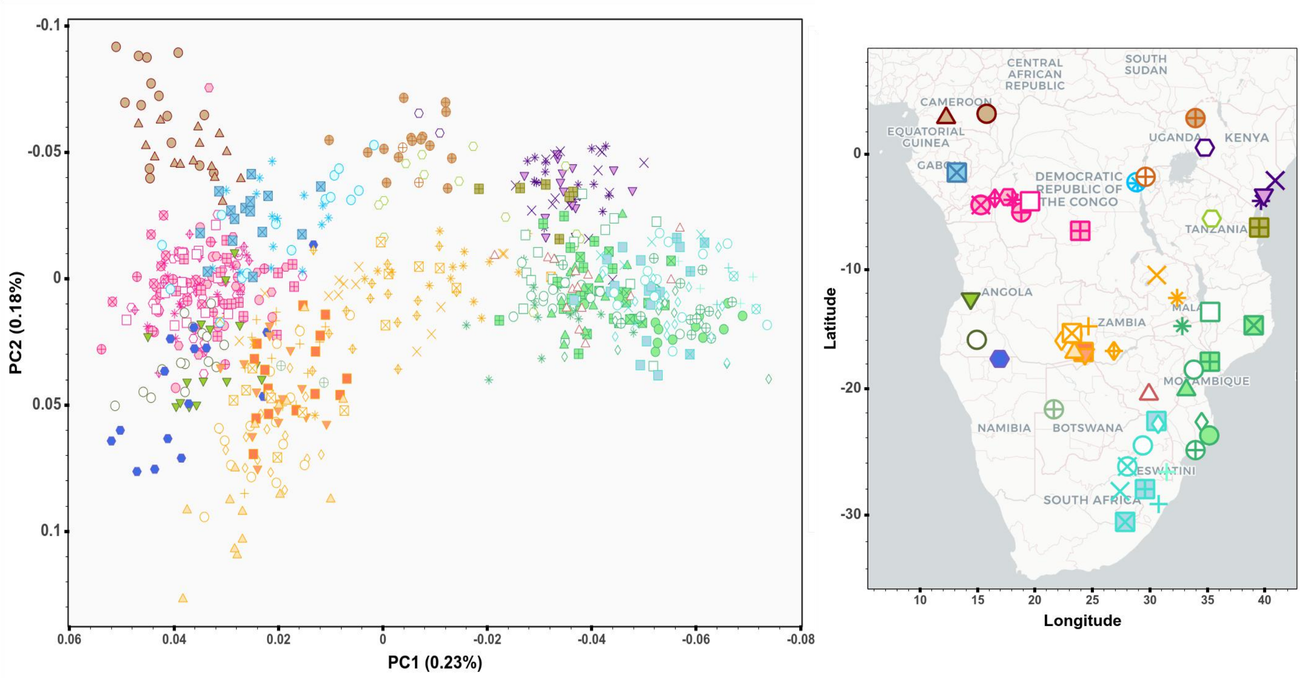


**Fig.S 4.2 | Ancestry-specific (AS-)-PCA after removing Himba and Herero.**
Figure showing AS-PCA plot of (**a**) each group and (**b**) each BSP included in the masked Only-BSP dataset after removing Himba and Herrero from the analyses. Legend is the same as in **Fig.S 4.4**, except for the Himba and Herrero populations that were not included on the maps.

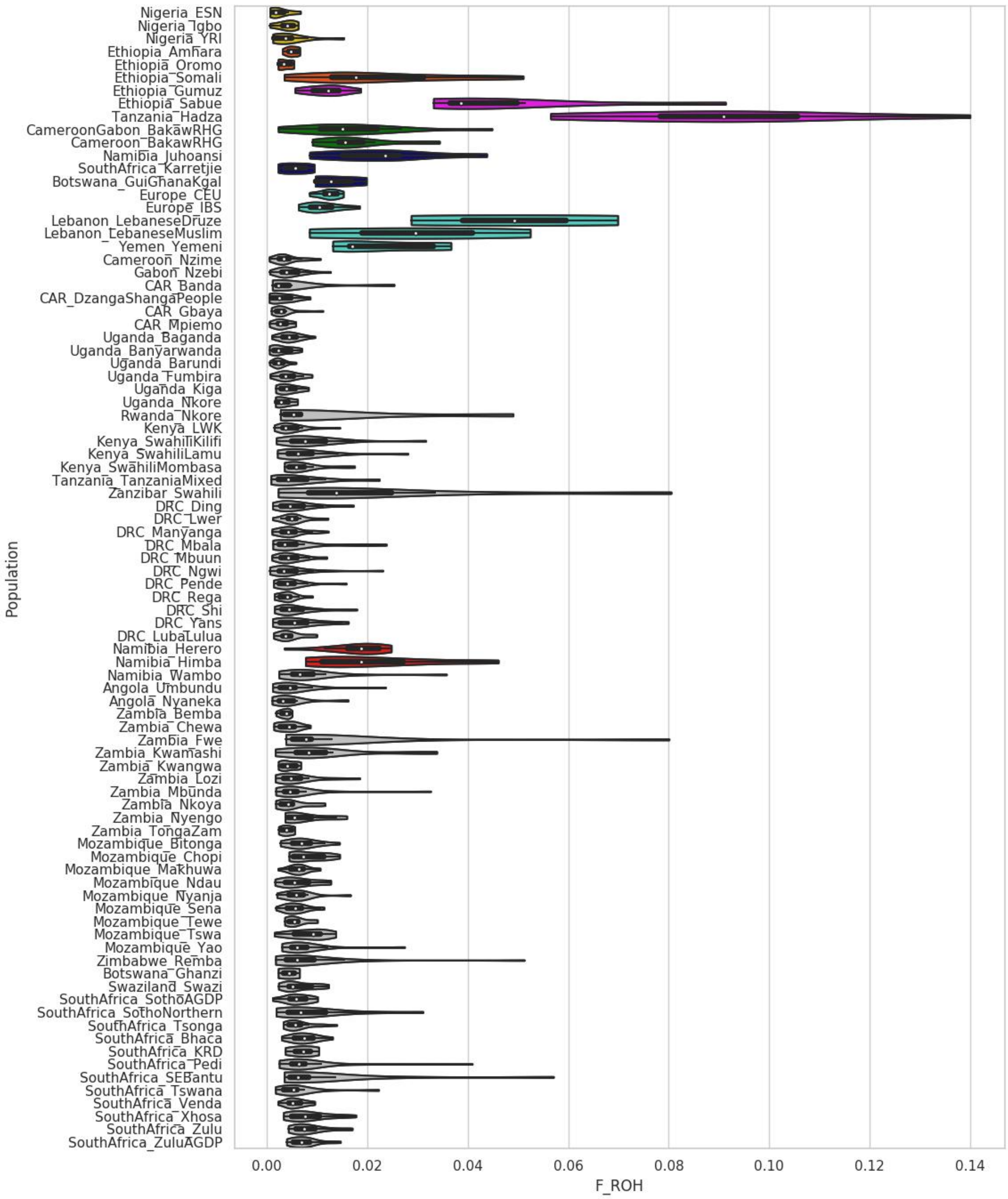


**Fig.S 4.3 | $F_{ROH}$ after masking the AfricanNeo dataset.**
Figure showing estimated ROH-based genomic inbreeding coefficient ($F_{ROH}$) of selected populations included in the AfricanNeo dataset after masking and imputation. We highlight Herero and Himba populations in red due to their higher $F_{ROH}$ values.

#### 5- Genome-wide runs of homozygosity (ROH) estimates

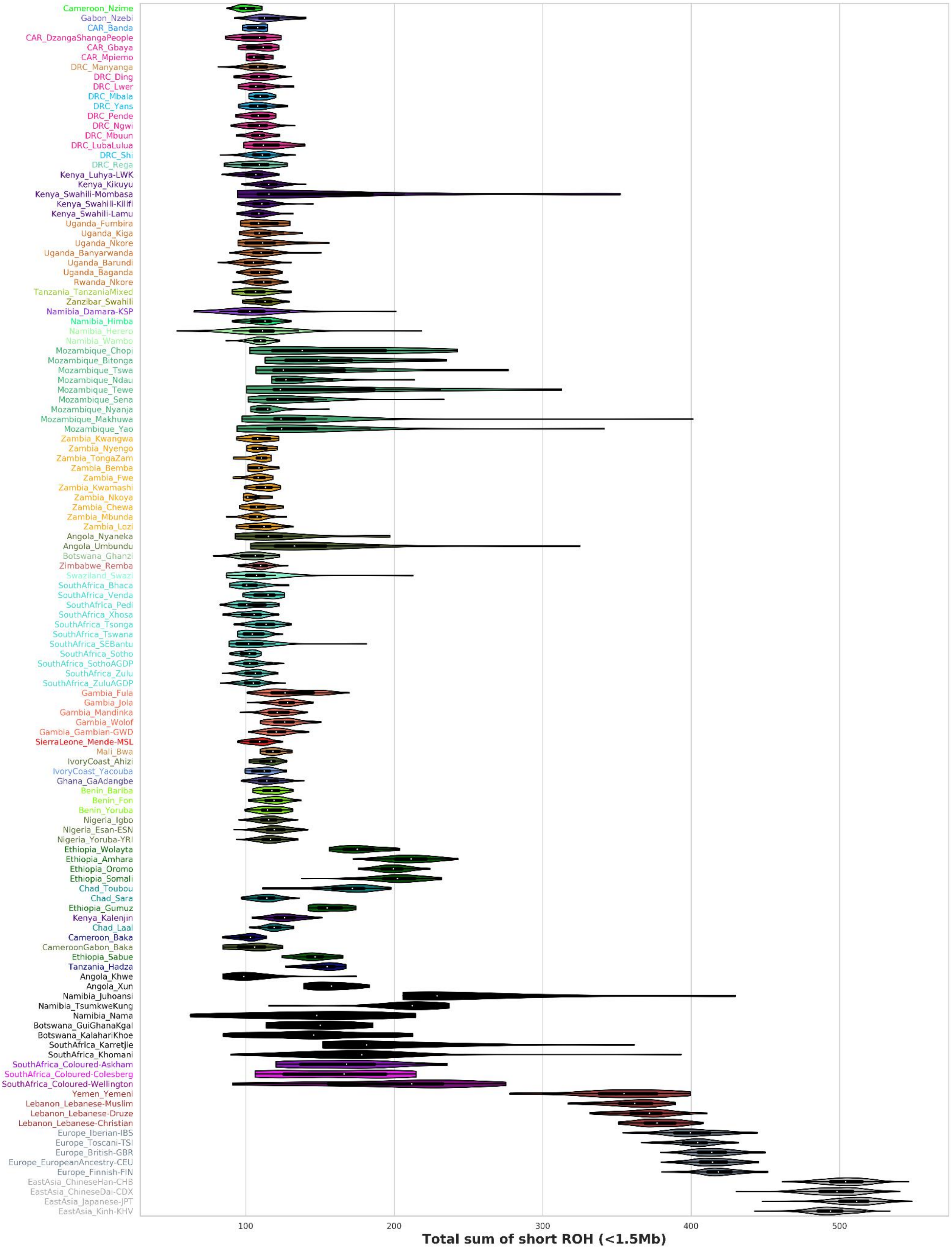


**Fig.S 5.1 | Total sum of short ROH length for the AfricanNeo dataset.**
Figure showing violin plots of the total sum of short ROH length (shorter than 1.5 Mb) in African and Eurasian populations included in the AfricanNeo dataset (see mean values in **Table.S 6**).

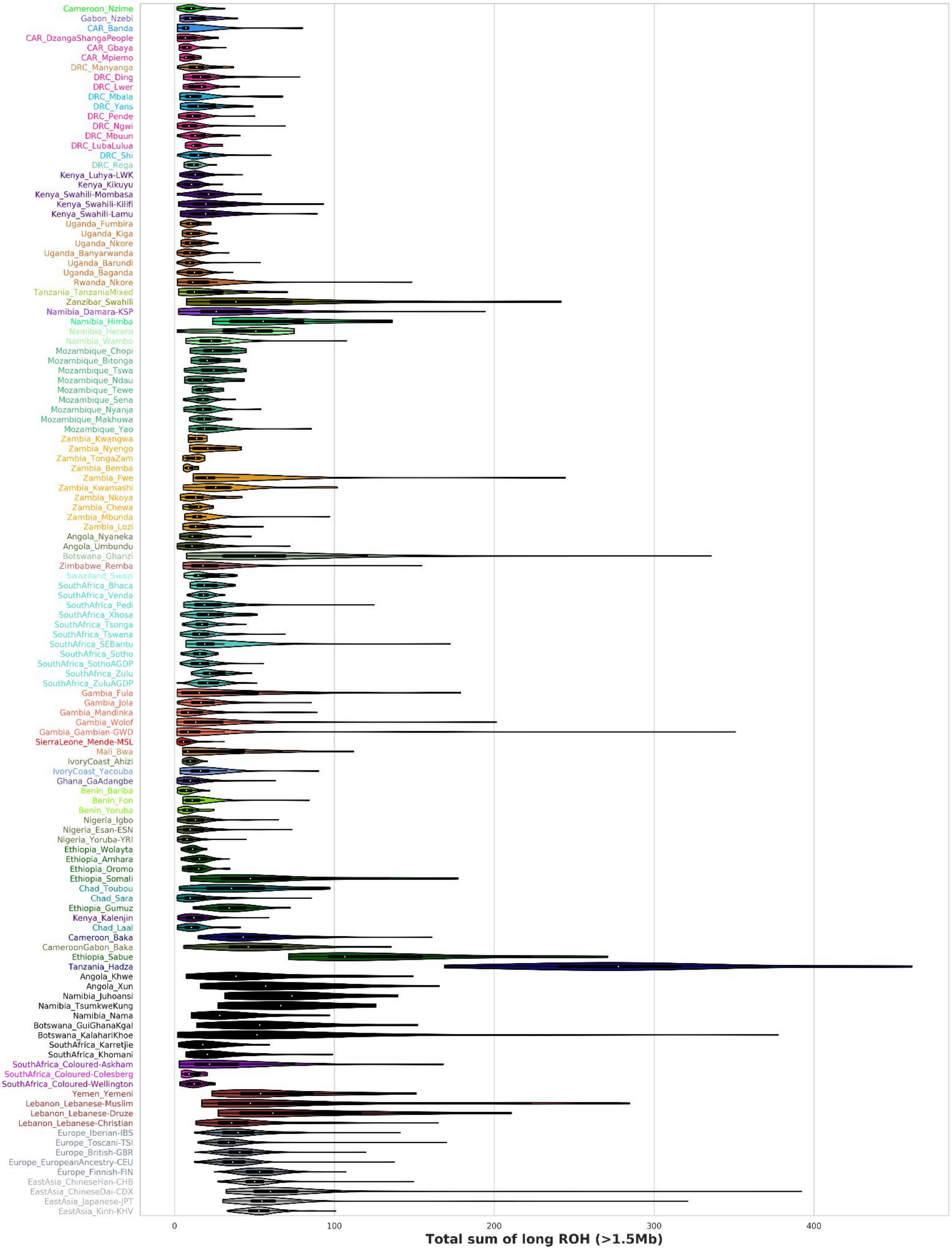


**Fig.S 5.2 | Total sum of long ROH length for the AfricanNeo dataset.**
Figure showing violin plots of the total sum of long ROH length (for segments longer than 1.5 Mb) in African and Eurasian populations included in the AfricanNeo dataset (see mean values in **Table.S 6**).

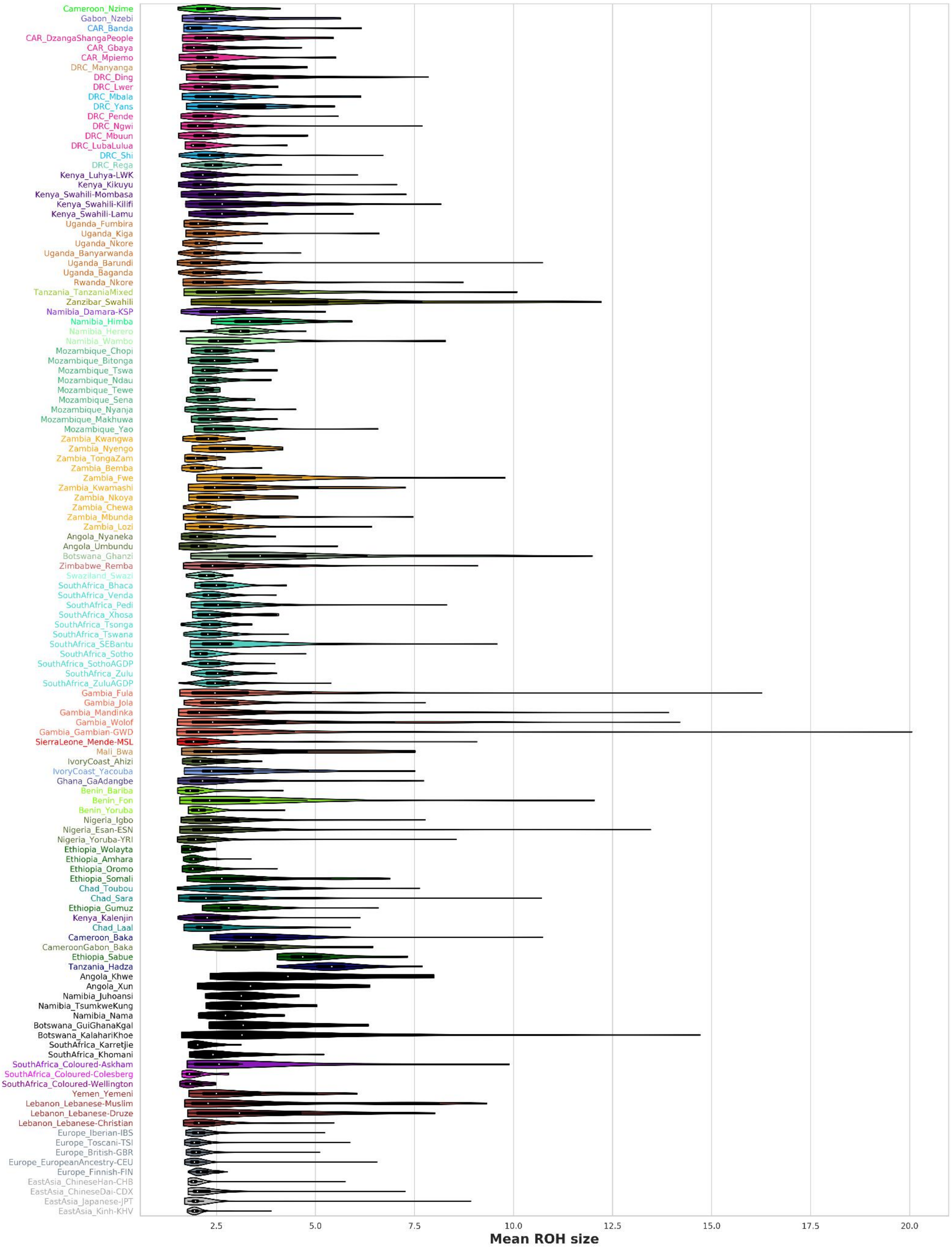


**Fig.S 5.3 | Mean of long ROH for the AfricanNeo dataset.**
Figure showing violin plots of the mean of long ROH (for segments longer than 1.5 Mb) in African and Eurasian populations included in the AfricanNeo dataset (see mean values in **Table.S 6**).

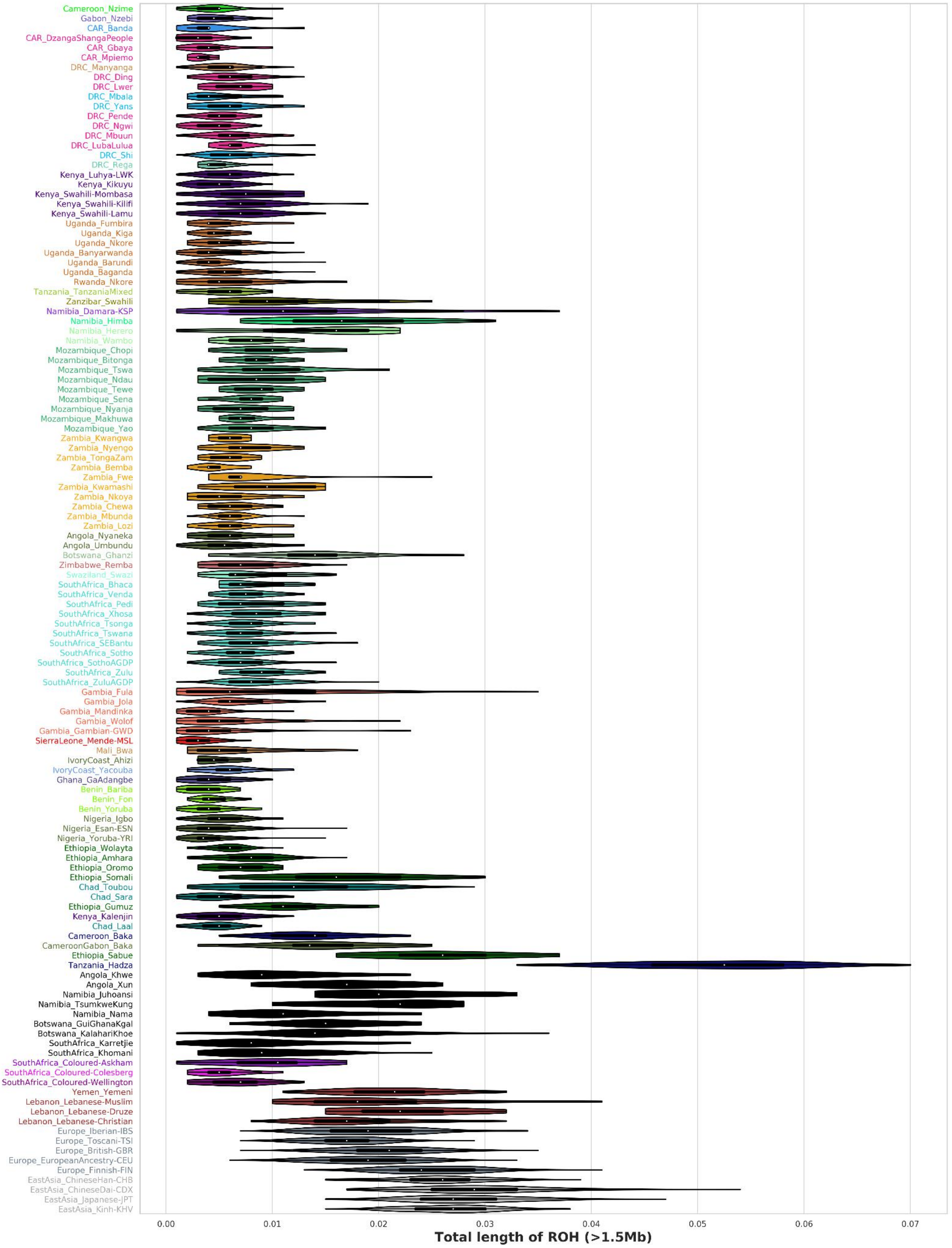


**Fig.S 5.4 | Total length of long ROH for the AfricanNeo dataset.**
Figure showing violin plots of the total length of long ROH (for segments longer than 1.5 Mb) in African and Eurasian populations included in the AfricanNeo dataset (see mean values in **Table.S 6**).

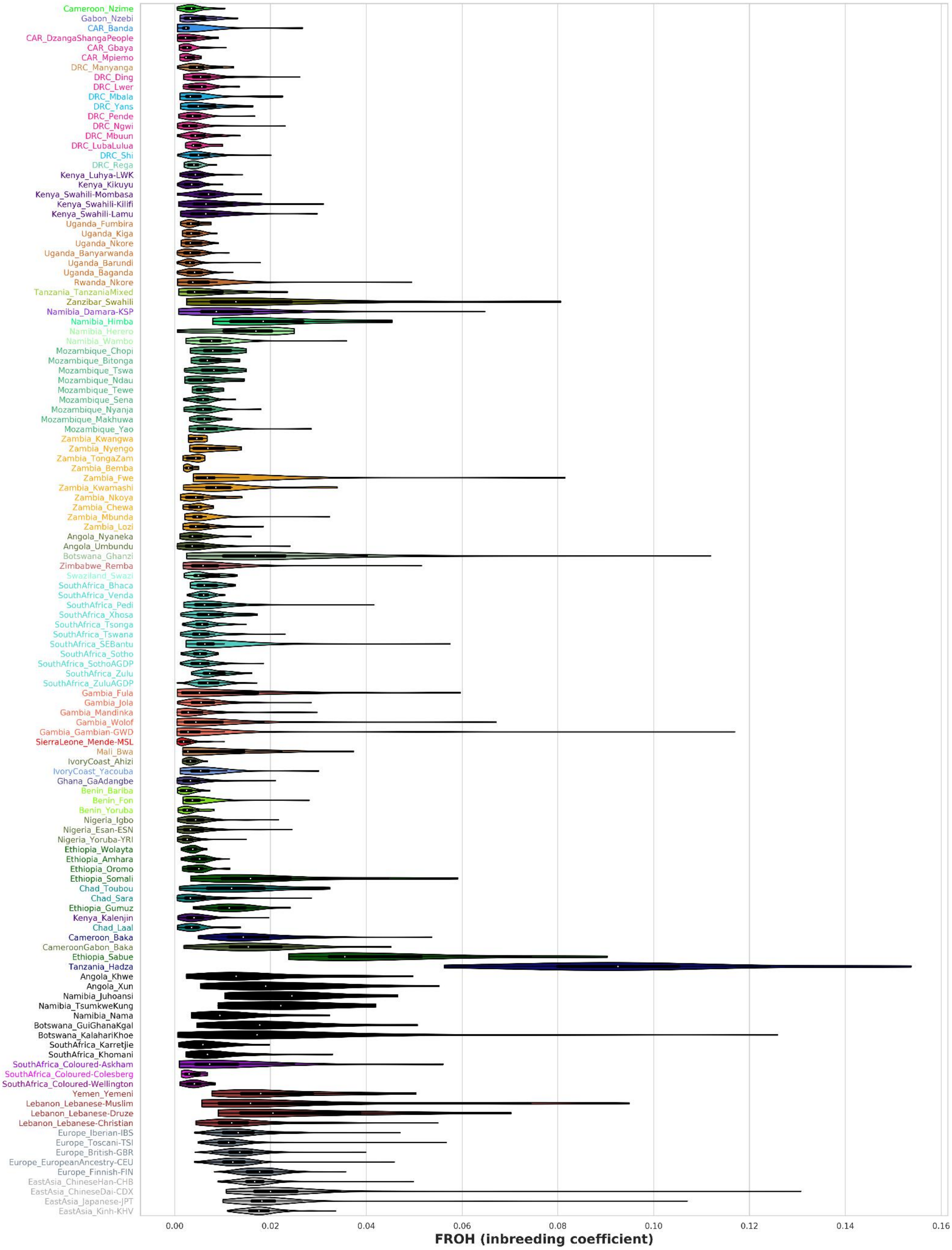


**Fig.S 5.5 | Genomic inbreeding coefficient ($F_{ROH}$) for the AfricanNeo dataset.**
Figure showing estimated genomic inbreeding coefficient ($F_{ROH}$) in African and Eurasian populations included in the AfricanNeo dataset (see mean values in **Table.S 6**).

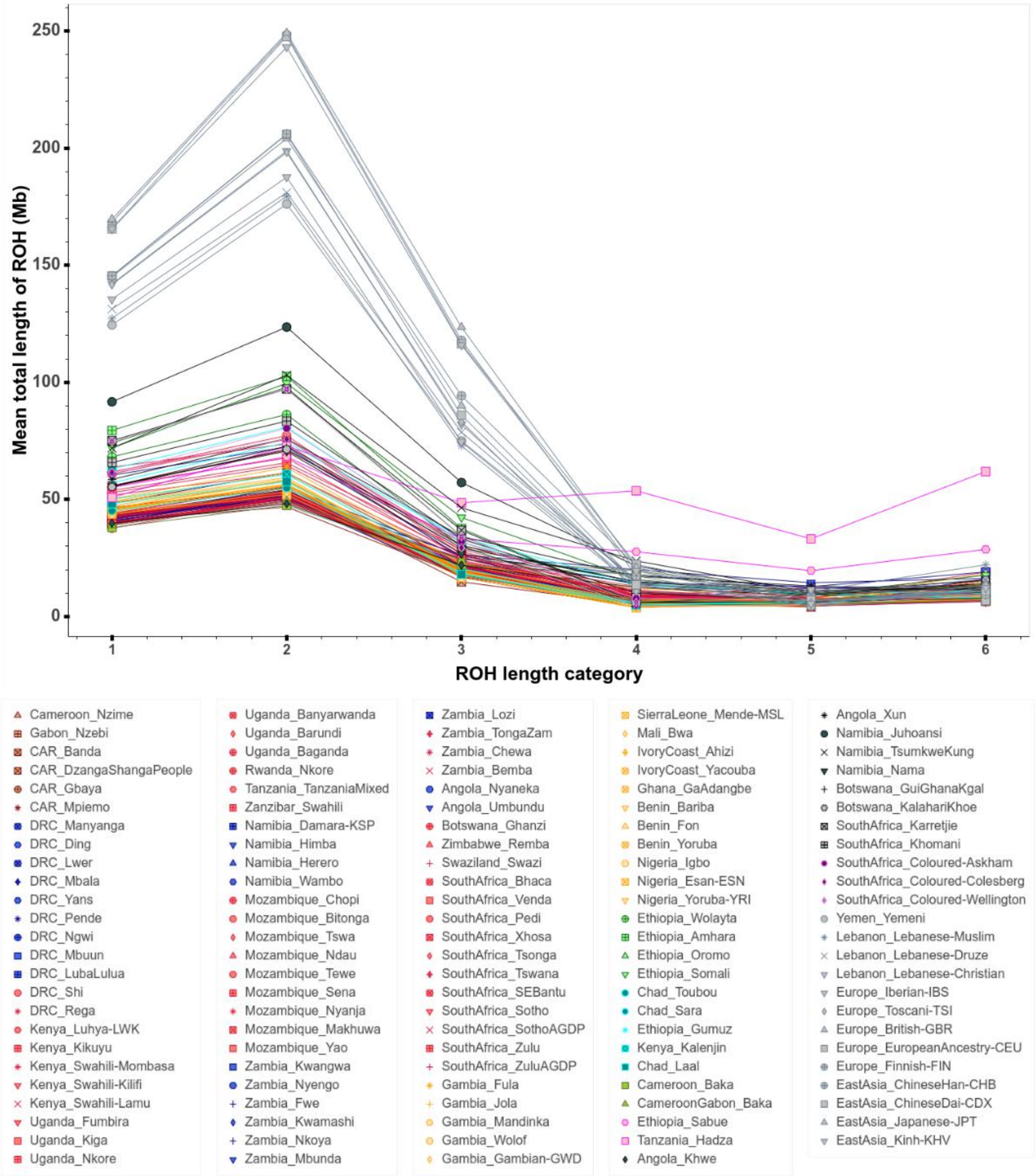


**Fig.S 5.6a | All categories of ROH length for the AfricanNeo dataset.**
Figure showing averages in each studied population for each **category of ROH length** (for segments longer than 0.3 Mb and shorter than 10 Mb) included in the AfricanNeo dataset (see mean values in **Table.S 6**). Violin plots of each population and for each category of ROH length are presented in **Fig.S 5.7–5.12**. To better visualize the results of each population, we also provide interactive plots (see **Fig.S_5.6a_ROH_categories_plot.html**). We also provide the results for only BSP in **Fig.S 5.6b.**

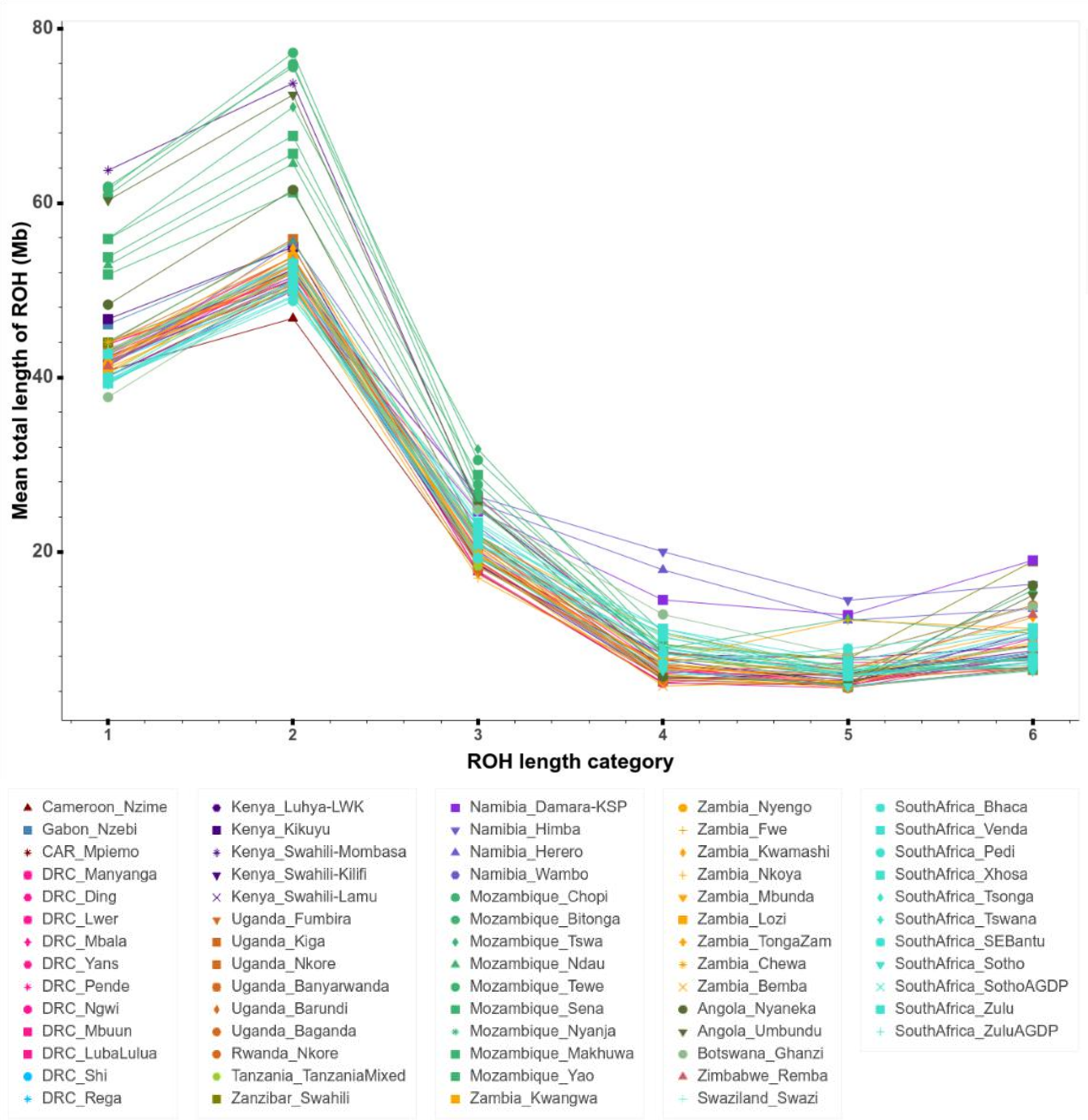


**Fig.S 5.6b | All categories of ROH length for the Only-BSP dataset.**
Figure showing averages in each BSP for each **category of ROH length** (for segments longer than 0.3 Mb and shorter than 10 Mb) (see mean values in **Table.S 6**). Violin plots of each population and for each category of ROH length are presented in **Fig.S 5.7–5.12**. To better visualize the results of each BSP, we also provide interactive plots (see **Fig.S_5.6b_ROH_categories_plot.html**).

**From Fig.S 5.7 to Fig.S 5.12 | Violin plots for six categories of ROH length.**

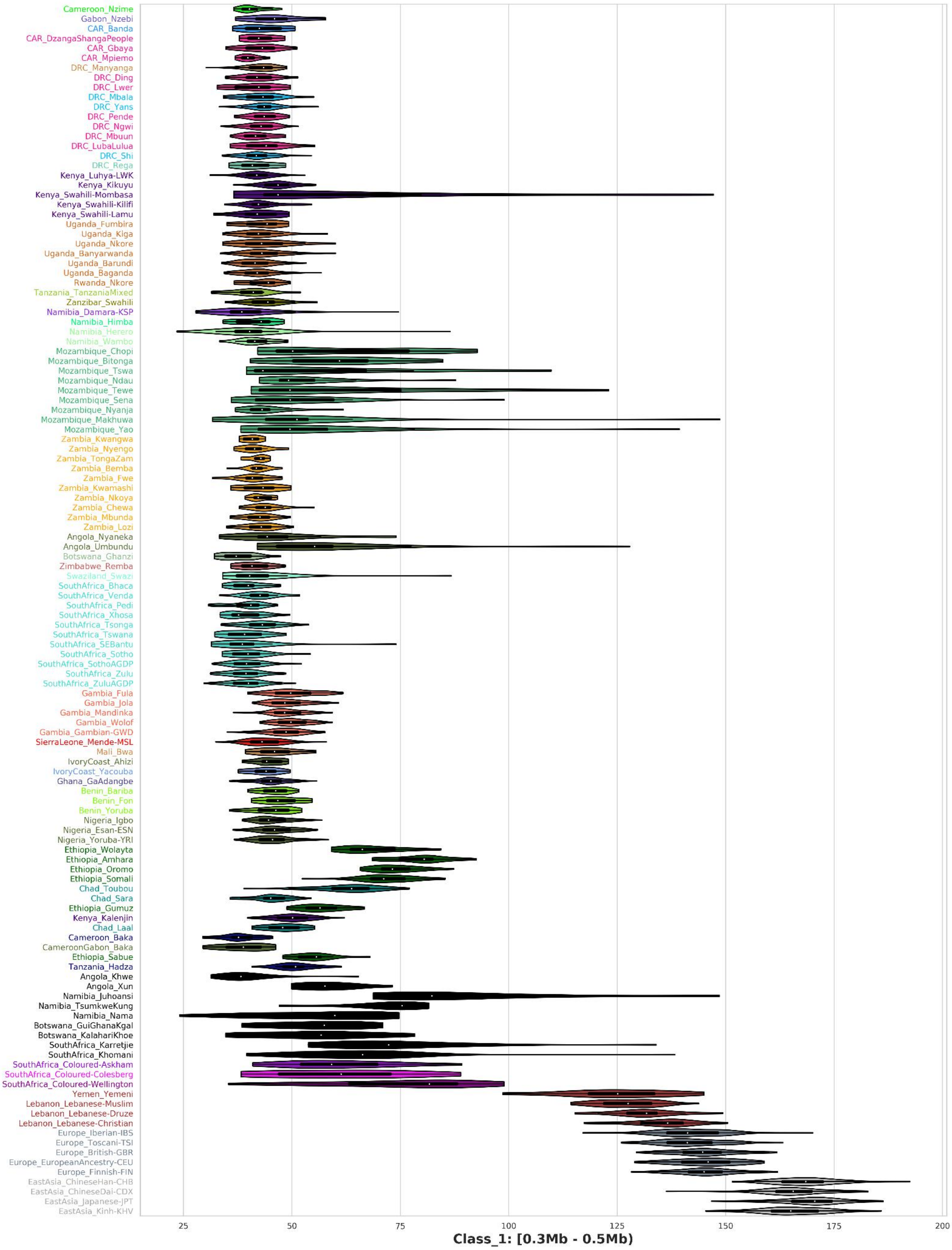


**Fig.S 5.7 | Violin plots for category 1 of ROH length** (for segments longer than 0.3 Mb and shorter than 0.5 Mb) in Eurasian populations included in the AfricanNeo dataset (mean values in **Table.S 6**).

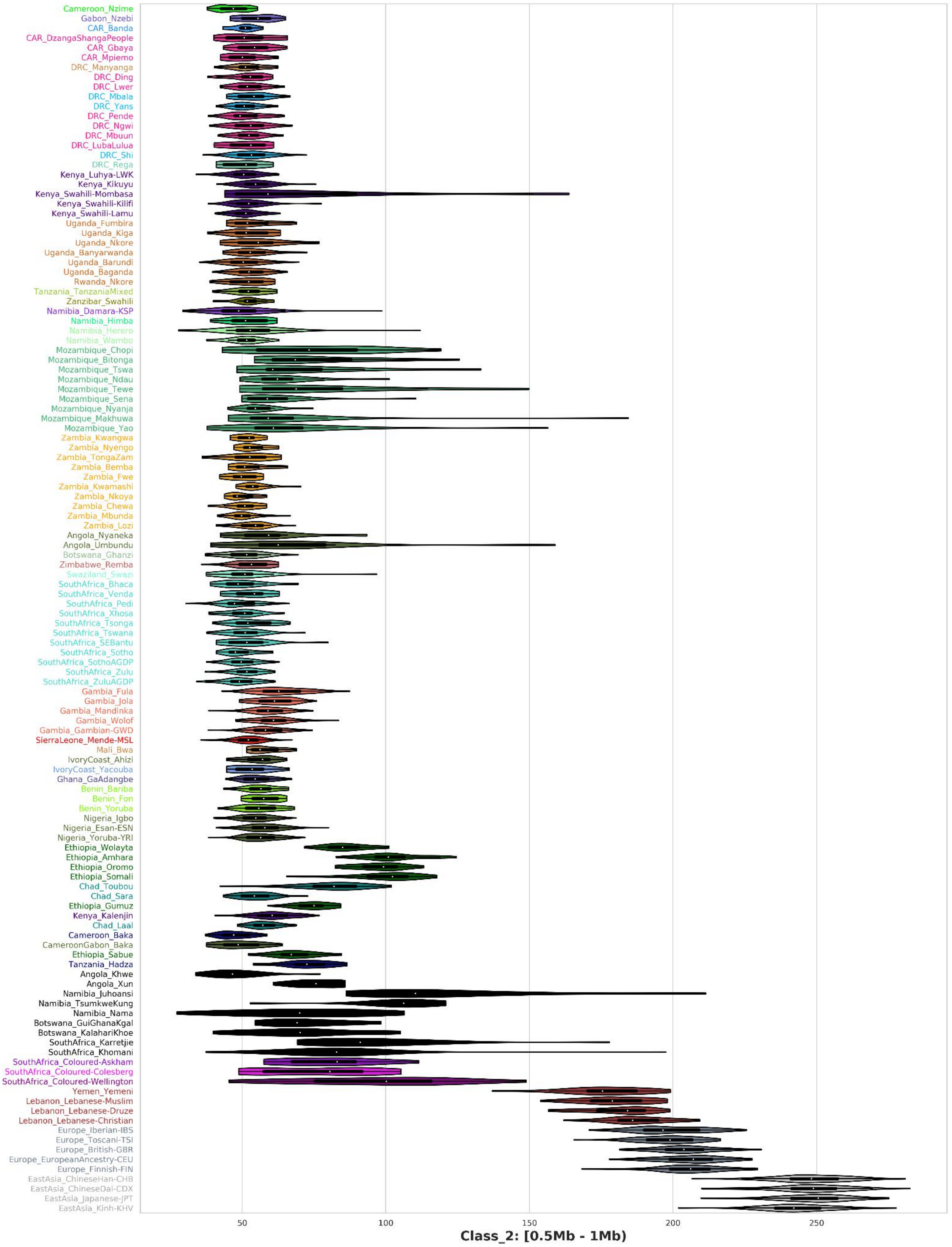


**Fig.S 5.8 | Violin plots for category 2 of ROH length** (for segments longer than 0.5 Mb and shorter than 1.0 Mb) in Eurasian populations included in the AfricanNeo dataset (see mean values in **Table.S 6**).

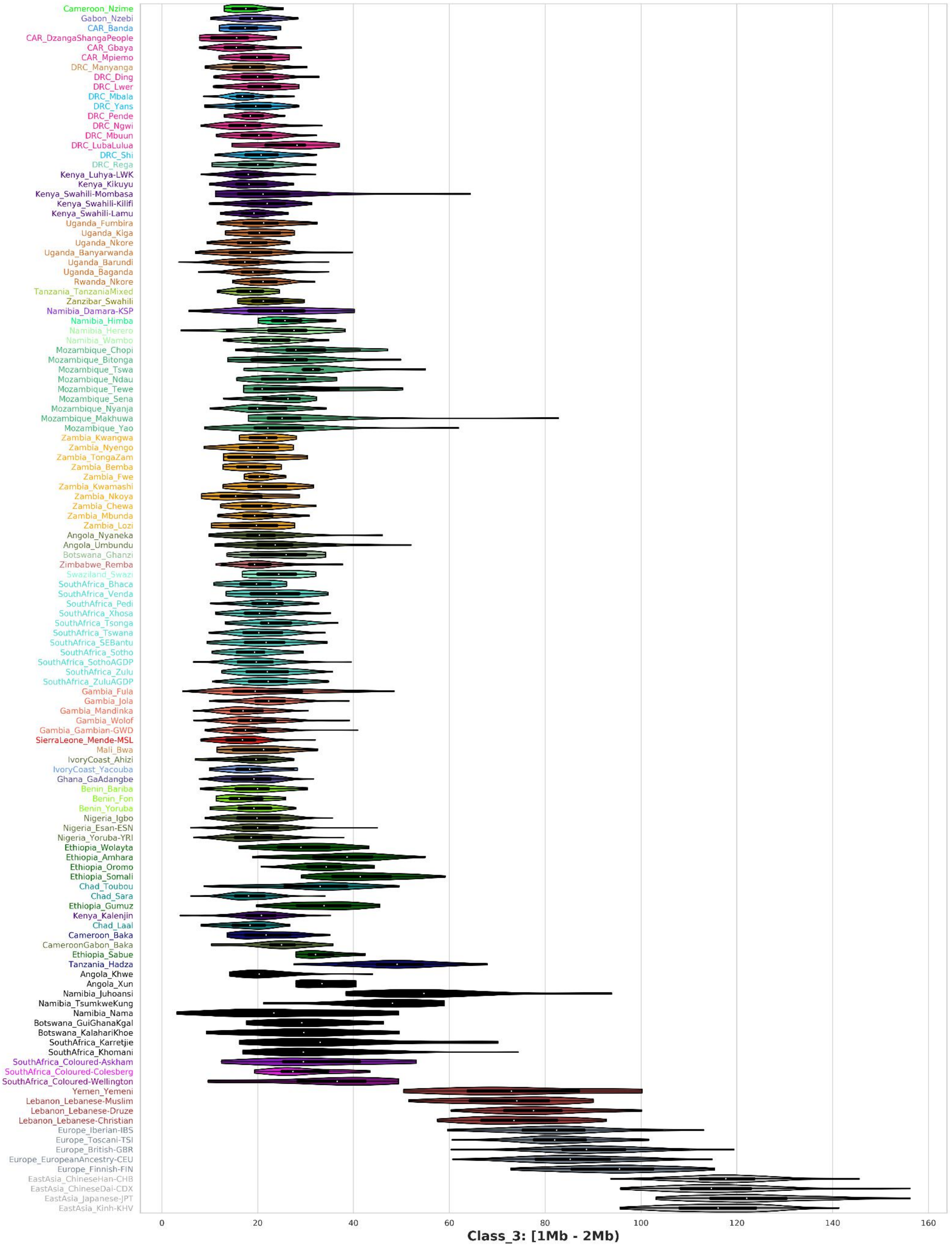


**Fig.S 5.9 | Violin plots for category 3 of ROH length** (for segments longer than 1.0 Mb and shorter than 2.0 Mb) in Eurasian populations included in the AfricanNeo dataset (see mean values in **Table.S 6**).

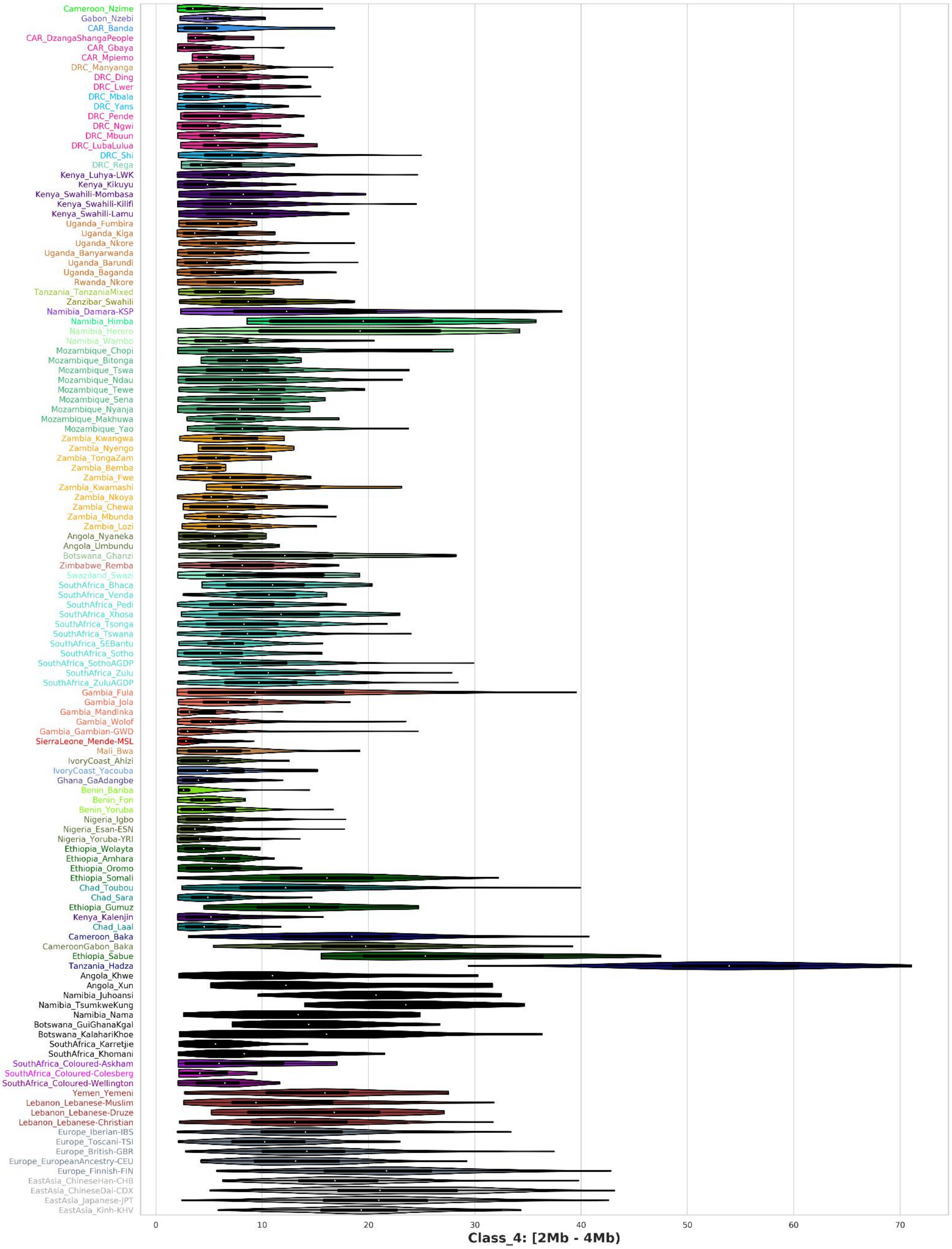


**Fig.S 5.10 | Violin plots for category 4 of ROH length** (for segments longer than 2.0 Mb and shorter than 4.0 Mb) in Eurasian populations included in the AfricanNeo dataset (see mean values in **Table.S 6**).

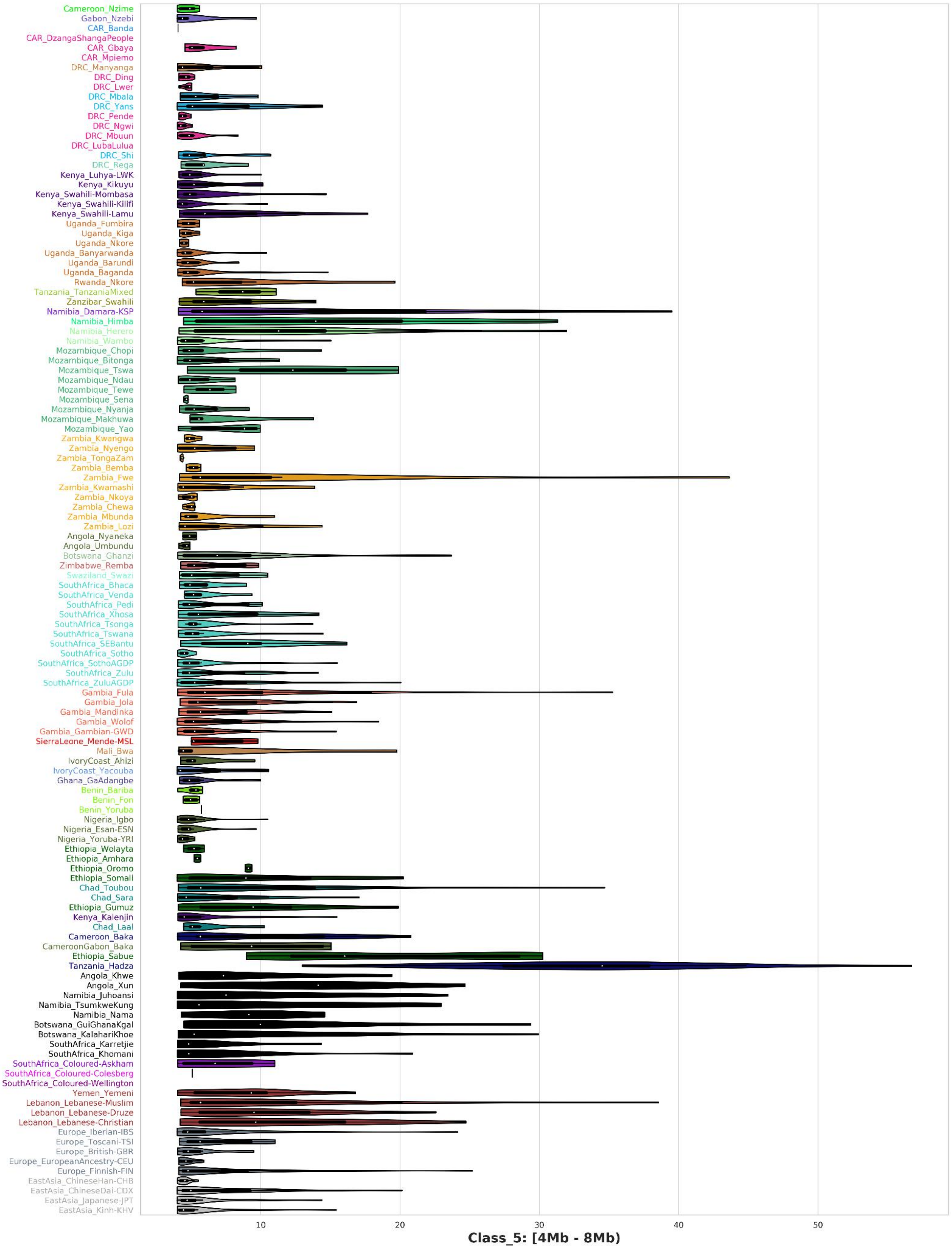


**Fig.S 5.11 | Violin plots for category 5 of ROH length** (for segments longer than 4.0 Mb and shorter than 8.0 Mb) in Eurasian populations included in the AfricanNeo dataset (see mean values in **Table.S 6**).

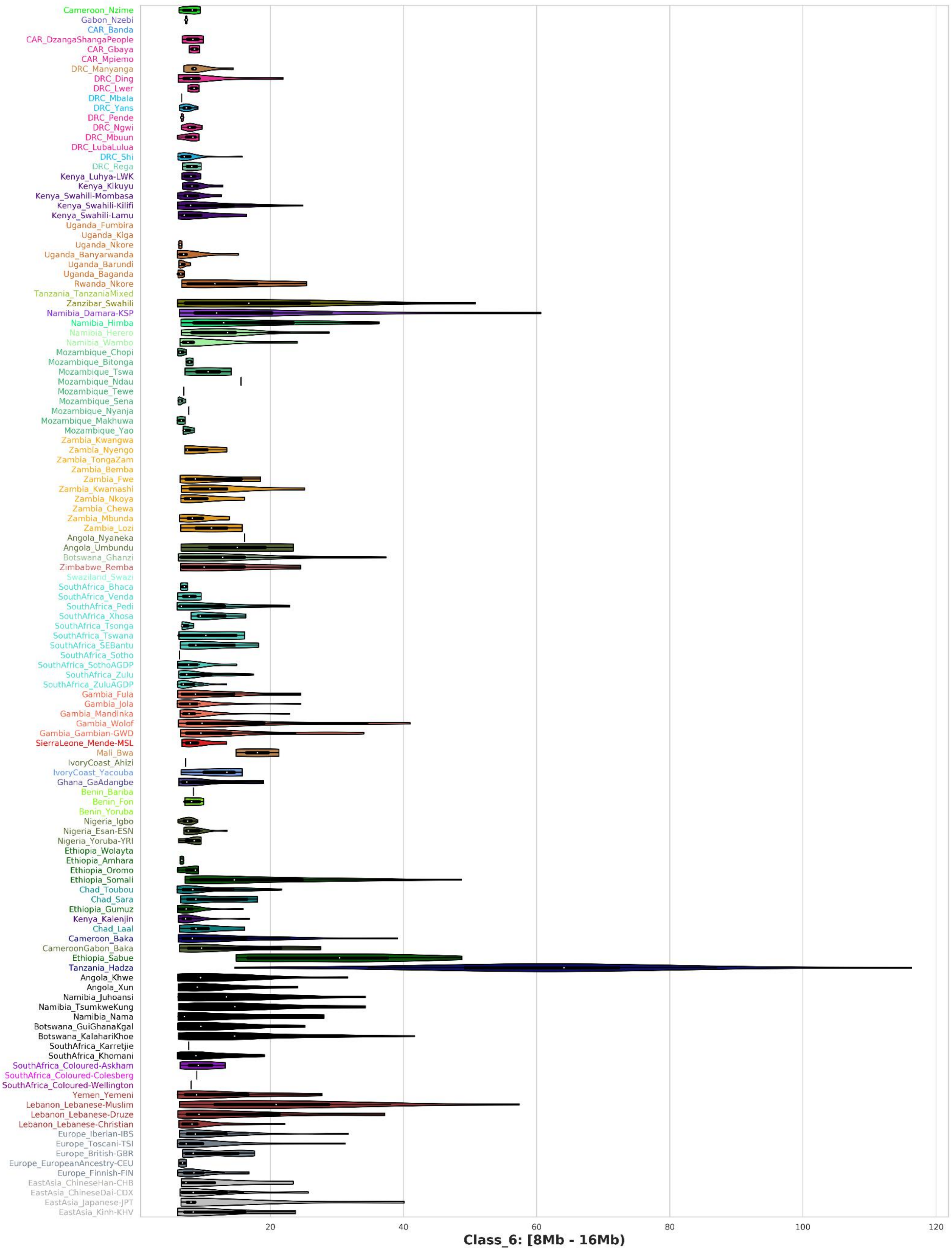


**Fig.S 5.12 | Violin plots for category 6 of ROH length** (for segments longer than 8.0 Mb and shorter than 16.0 Mb) in Eurasian populations included in the AfricanNeo dataset (see mean values in **Table.S 6**).

#### 6- Estimated effective population sizes and demographic founder events

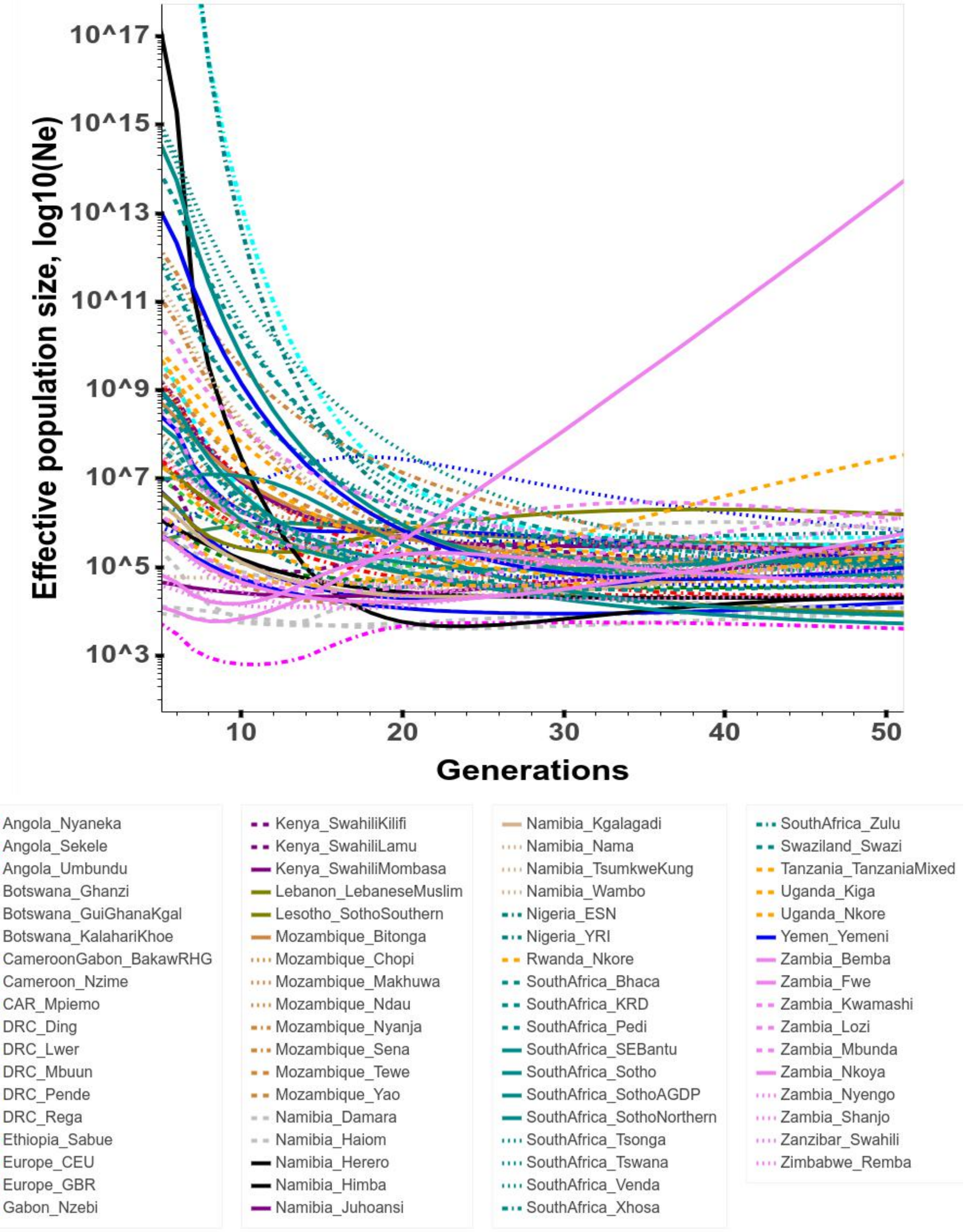


**Fig.S 6.1 | IBDNe results for the AfricanNeo dataset.**
Figure showing estimated effective population sizes ($N_e$) in studied BSP estimated using IBDNe for the last 50 generations. To better visualize the results of each population, we also provide interactive plots (see **Fig.S_6.1_IBDNe_plot.html**).

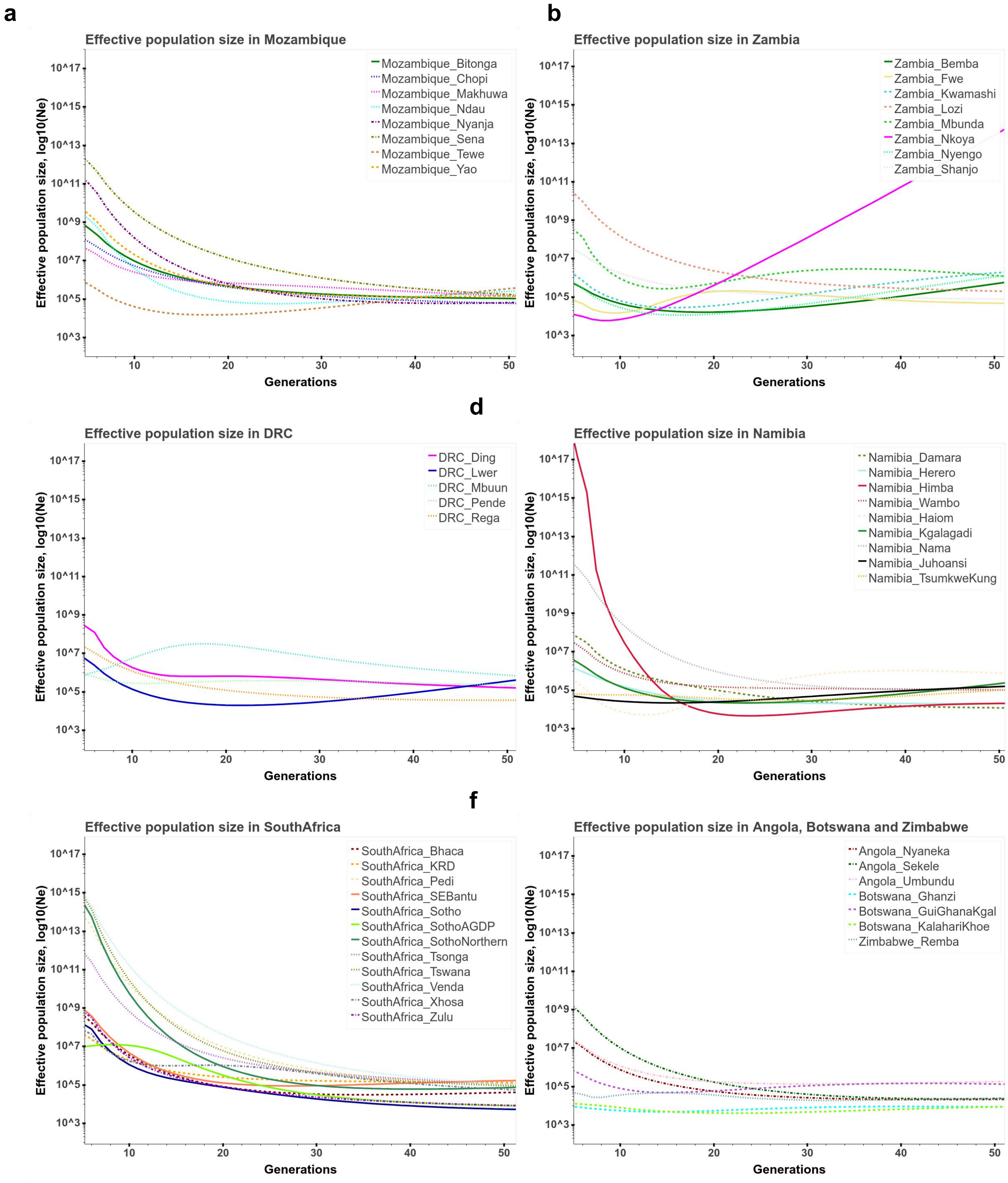


**Fig.S 6.2 | IBDNe results for BSP from six African regions.**
Figure showing estimated effective population sizes ($N_e$) of BSP from six African regions estimated using IBDNe for the last 50 generations. Figures include $N_e$ of studied BSP from (**a**) Mozambique, (**b**) Zambia, (**c**) DRC, (**d**) Namibia, (**e**) South Africa, and (**f**) Angola, Botswana and Zimbabwe.

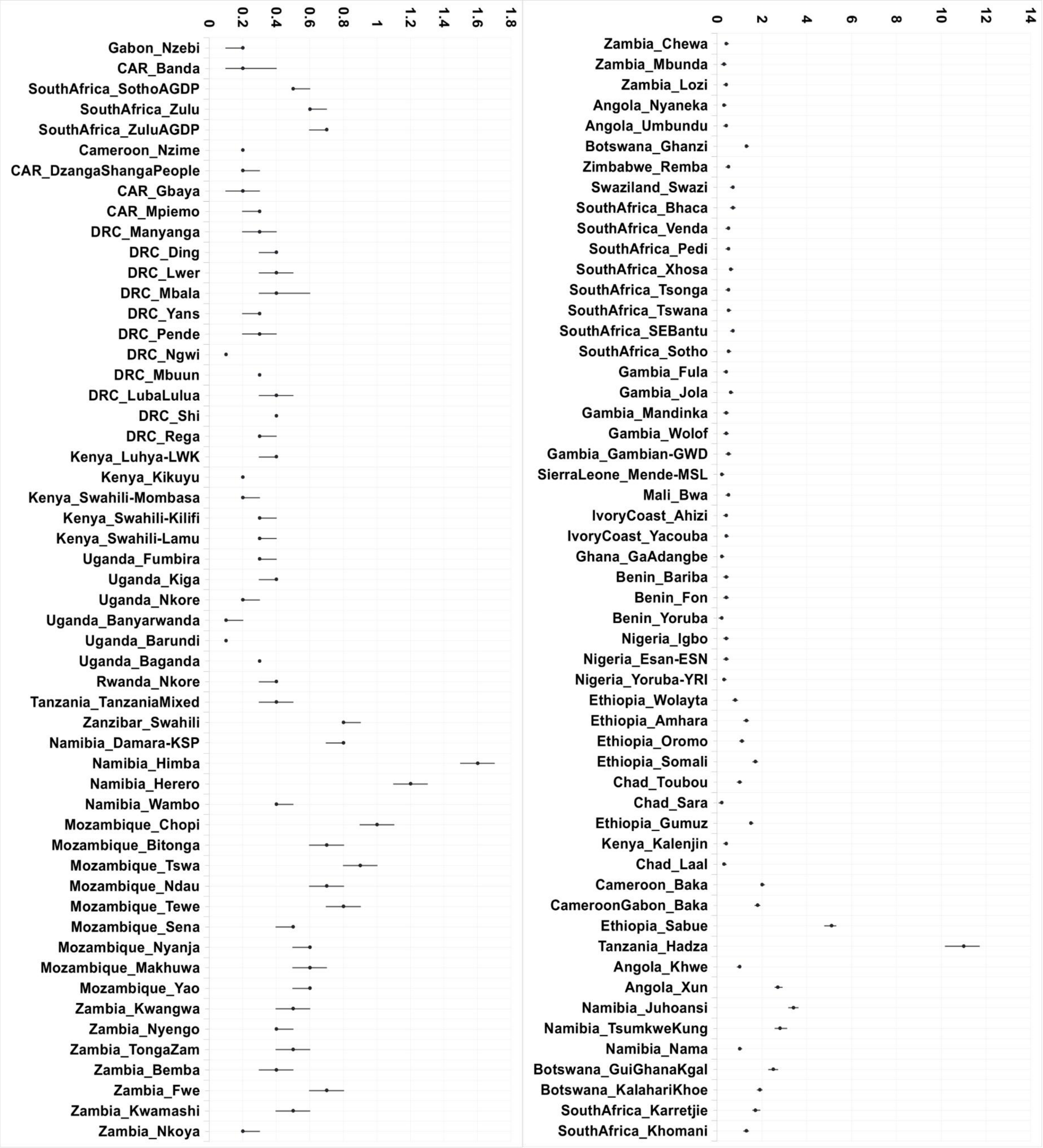


**Fig.S 6.3 | Intensity of founder events in sub-Saharan African populations.**
Figure showing the intensity of the founder event ($I_f$ in %, each dot; and 95%CI, each line) calculated for sub-Saharan African populations included in the AfricanNeo dataset using ASCEND analysis. To better see the results, the plot was divided into two plots, and the X-axes have different ranges in each plot. Further details of all the results were included in **Table.S 7**.

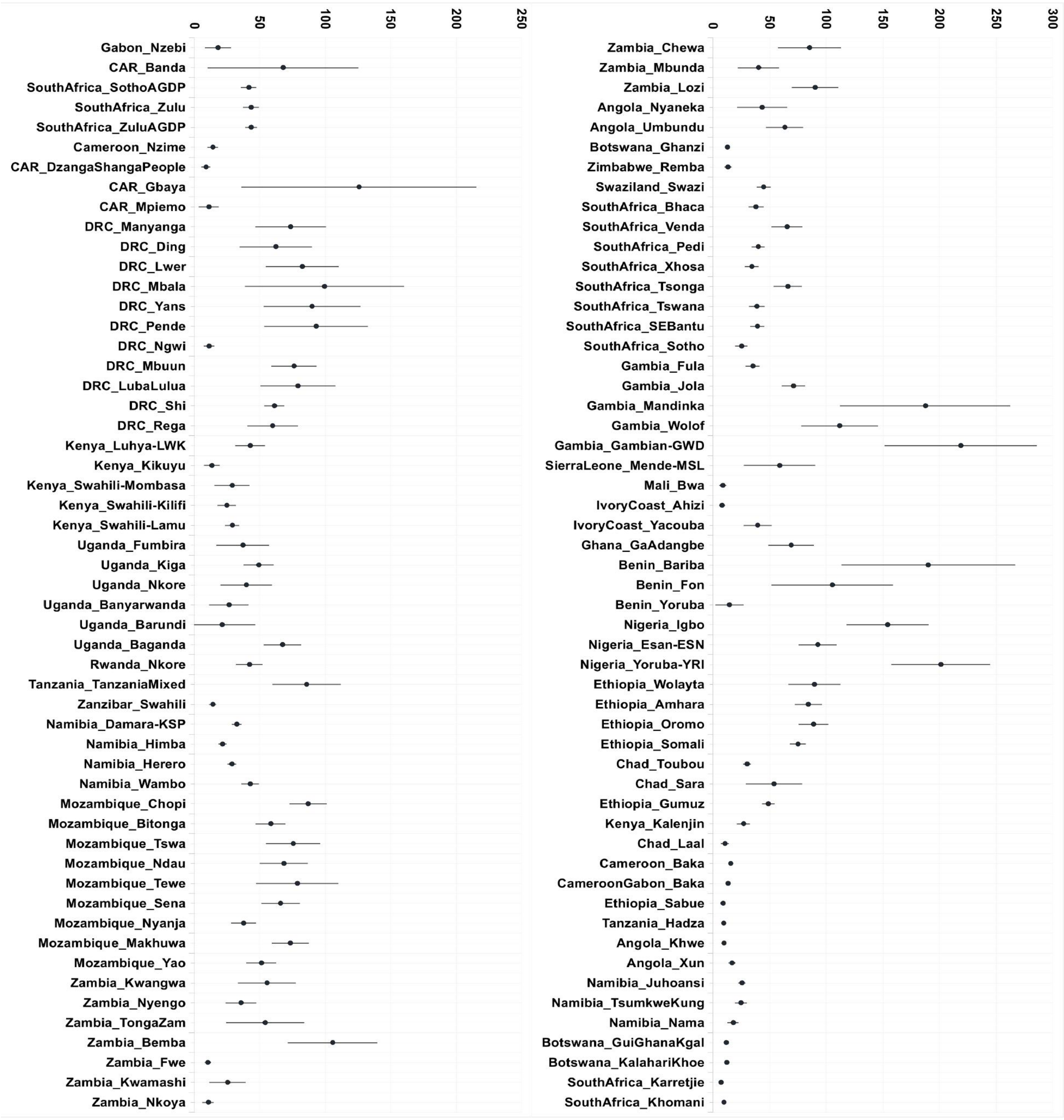


**Fig.S 6.4 | Timing of founder ages in sub-Saharan African populations.**
Figure showing the timing of the founder event ($T_f$ in generations, each dot; and 95%CI, each line) calculated for sub-Saharan African populations included in the AfricanNeo dataset using ASCEND analysis. To better see the results, the plot was divided into two plots and the x-axes have different ranges in each plot. Further details of all the results were included in **Table.S 7**.

#### 7- Patterns of isolation-by-distance for the unmasked and masked datasets

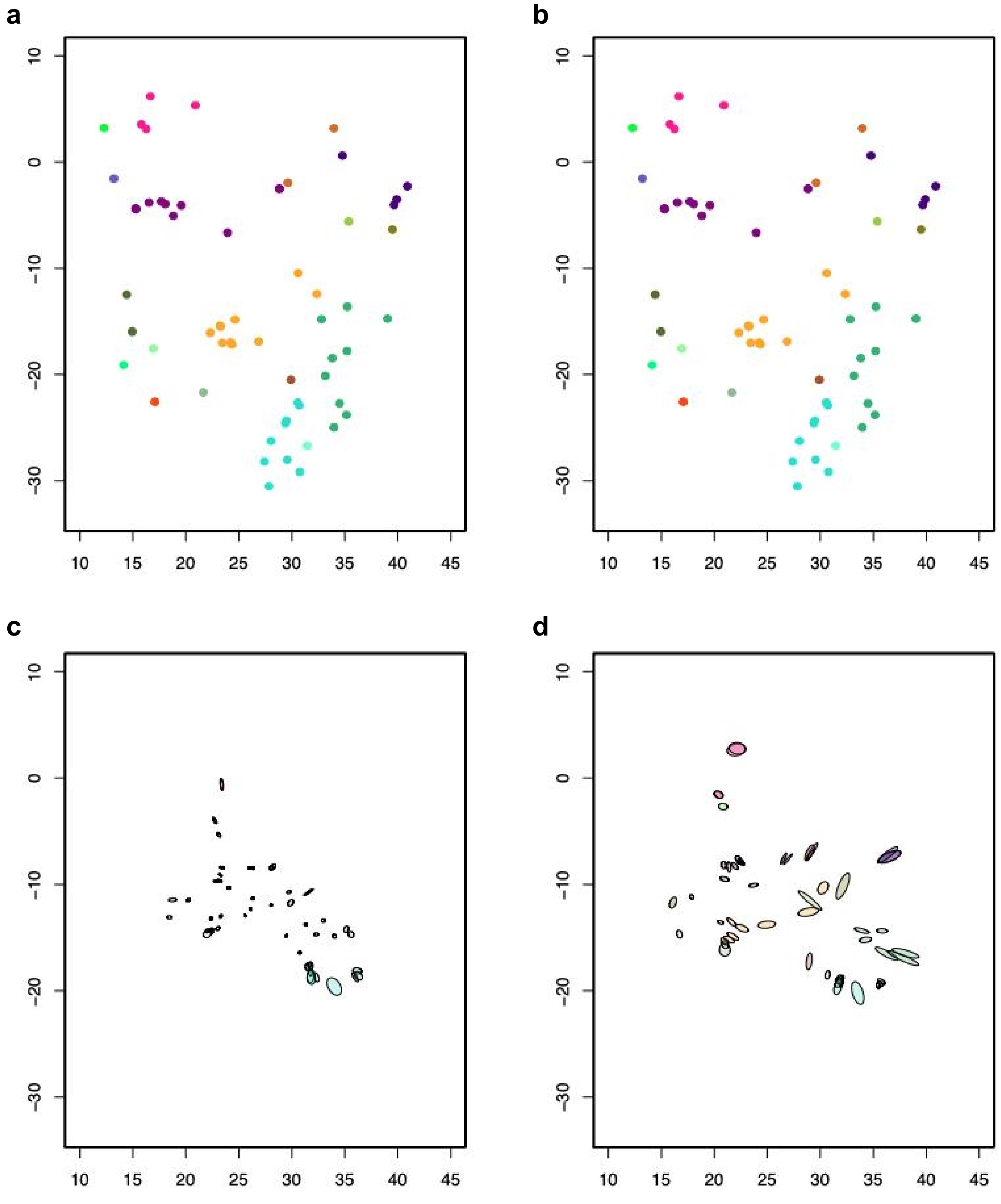


**Fig.S 7.1 | SpaceMix results with no population text overlays.**
Figure showing SpaceMix tests for the **unmasked** Only-BSP dataset, no population text overlays. X- and Y-axes are latitude and longitude in the geogenetic space, respectively. Panel showing the results of each IBD model, and the assumptions of each model were: (**a**) No migration and no admixture; (**b**) No migration but admixture; (**c**) Migration but no admixture; and (**d**) Both migration and admixture.

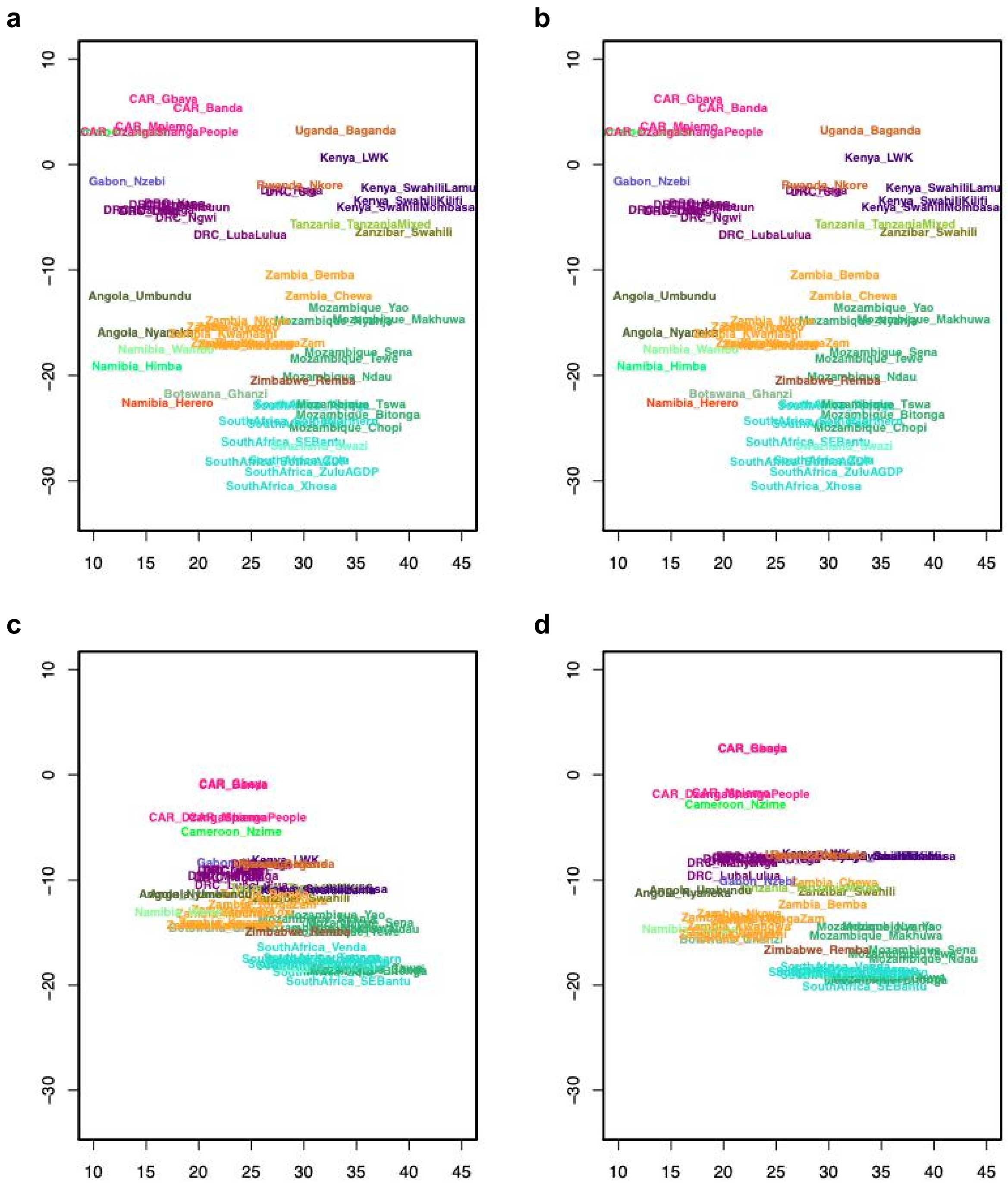


**Fig.S 7.2 | SpaceMix results with text instead of dots.**
Figure showing SpaceMix tests for the **unmasked** Only-BSP dataset, text instead of dots. X- and Y-axes are latitude and longitude in the geogenetic space, respectively. Panel showing the results of each IBD model, and the assumptions of each model were: (**a**) No migration and no admixture; (**b**) No migration but admixture; (**c**) Migration but no admixture; and (**d**) Both migration and admixture.

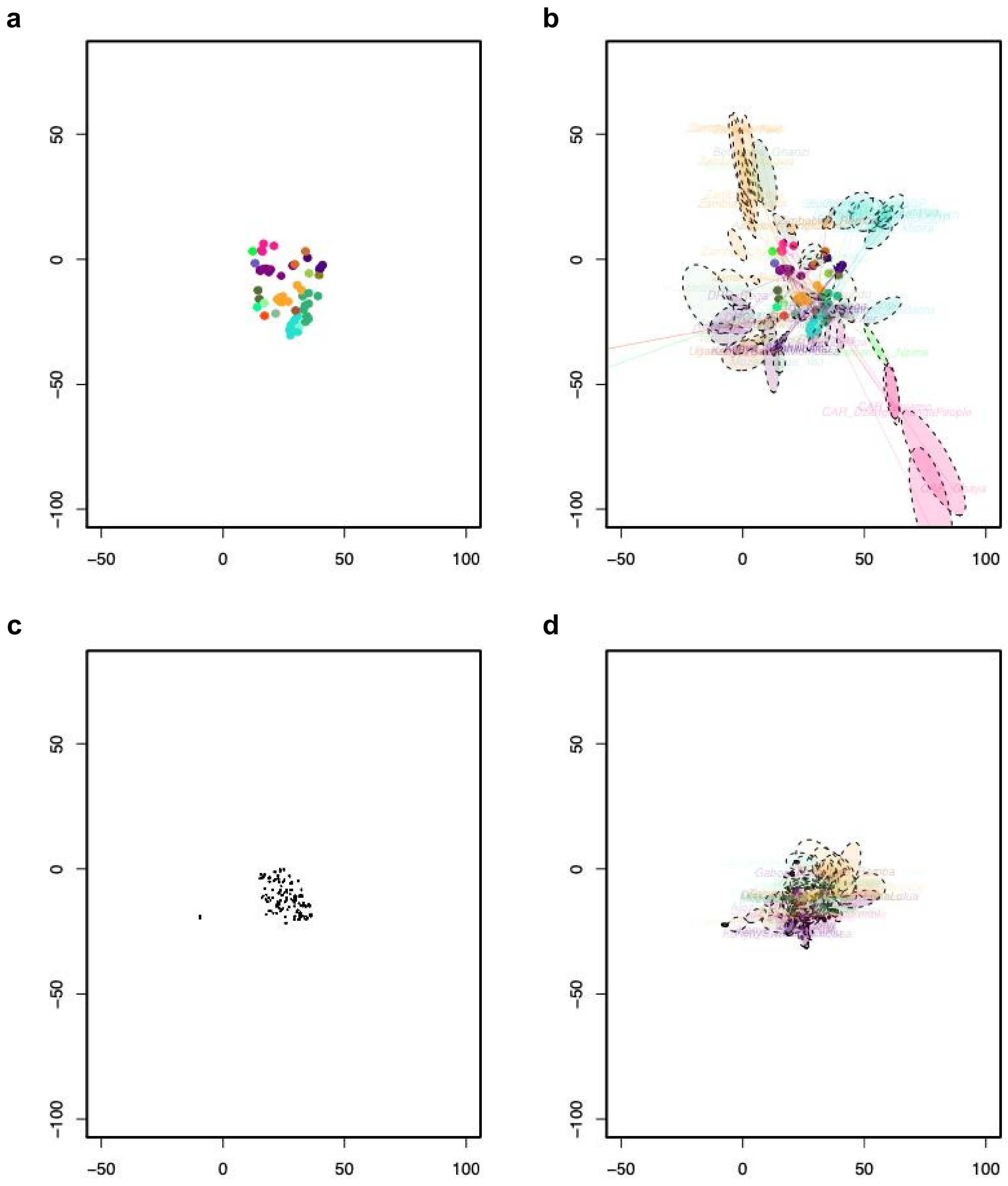

**Fig.S 7.3 | SpaceMix results with sources of admixture.**
Figure showing SpaceMix tests for the **unmasked** Only-BSP dataset, sources of admixture were indicated with a dashed ellipsis. X- and Y-axes are latitude and longitude in the geogenetic space, respectively. Panel showing the results of each IBD model, and the assumptions of each model were: (**a**) No migration and no admixture; (**b**) No migration but admixture; (**c**) Migration but no admixture; and (**d**) Both migration and admixture.

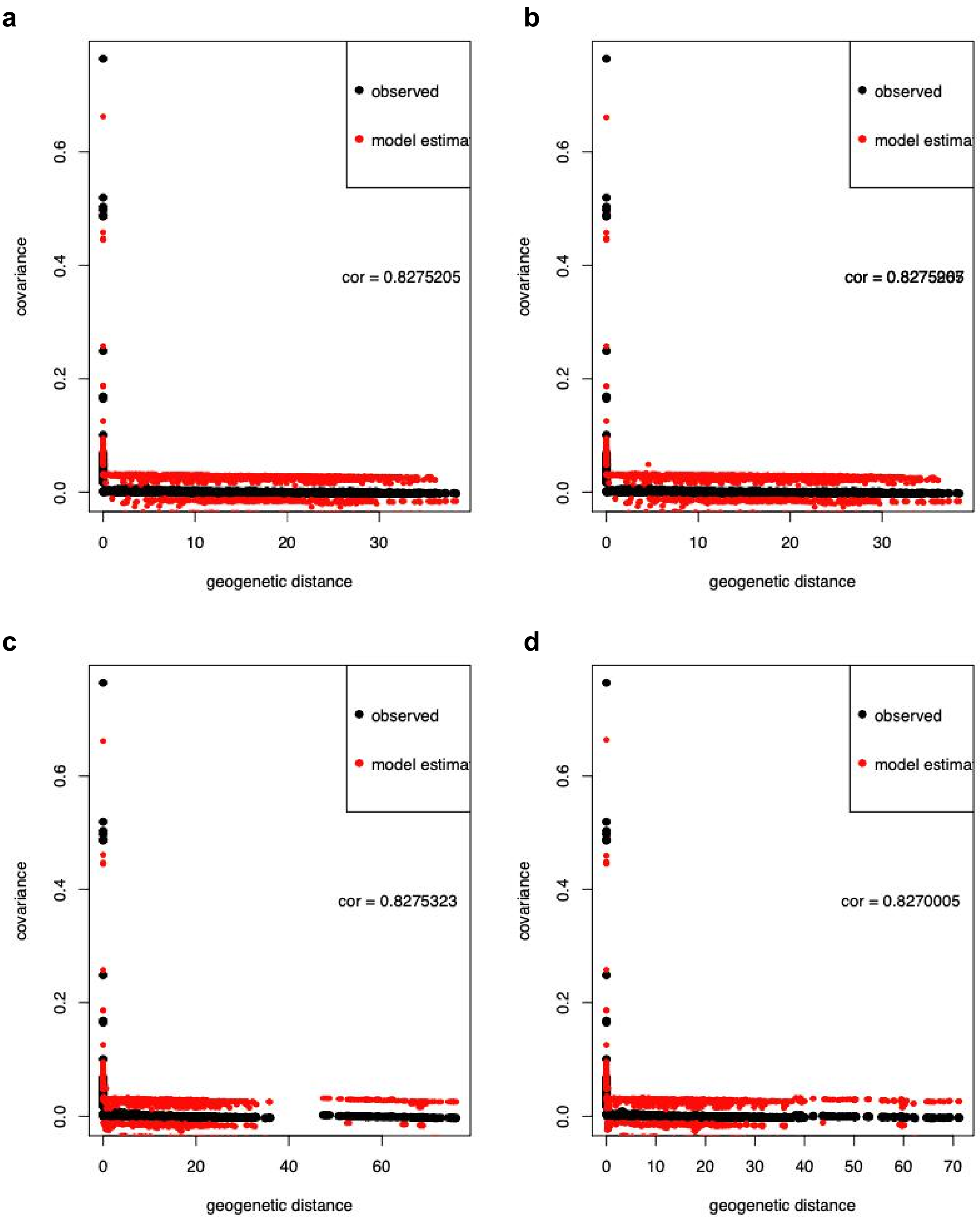


**Fig.S 7.4 | SpaceMix results with correlations of each tested model.**
Figure showing SpaceMix tests for the **unmasked** Only-BSP dataset, highlighting the correlation between the observed data and the data estimated from the IBD model. We computed the Pearson correlation between the two series (see “cor” values on each plot). Panel showing the results of each IBD model, and the assumptions of each model were: (**a**) No migration and no admixture; (**b**) No migration but admixture; (**c**) Migration but no admixture; and (**d**) Both migration and admixture.

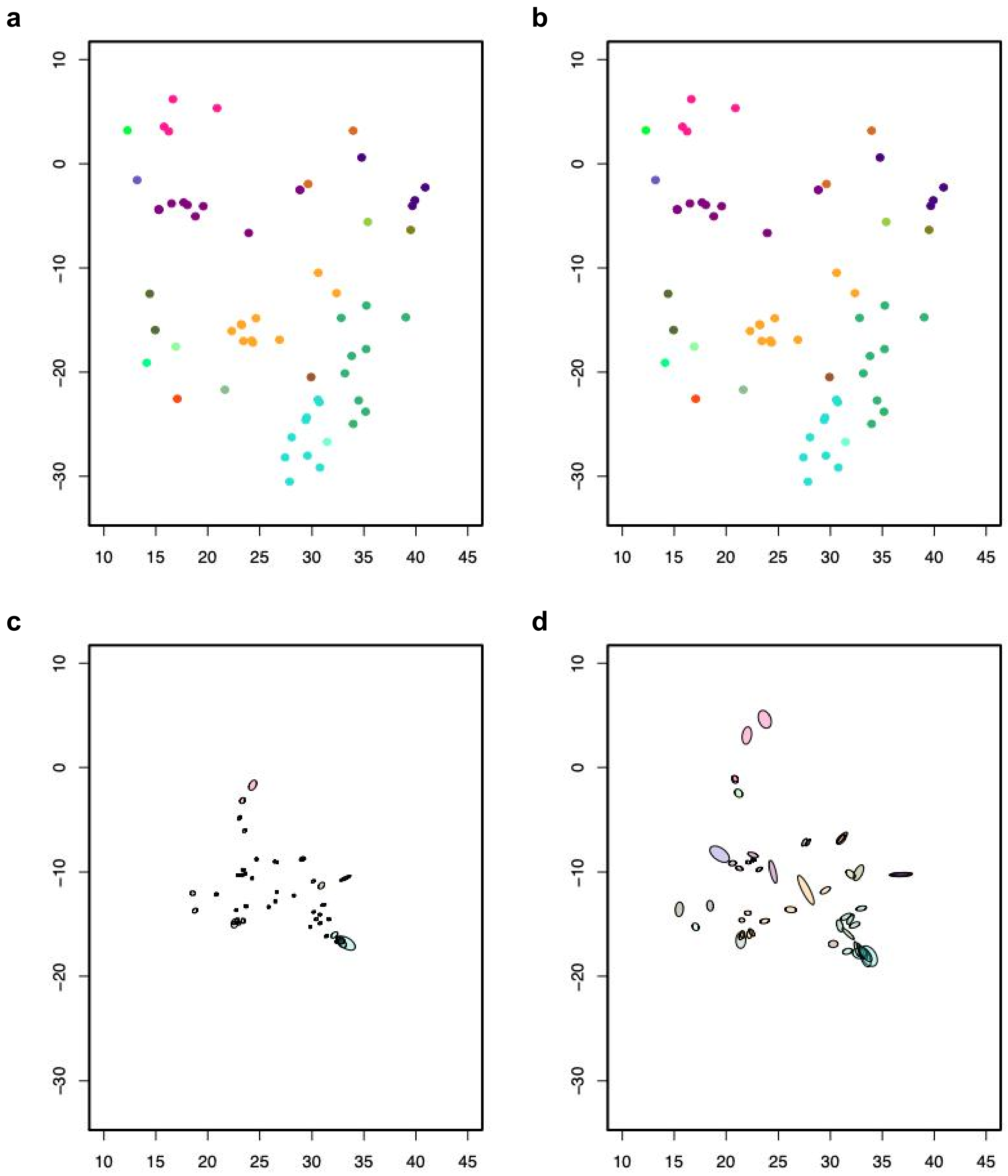


**Fig.S 7.5 | SpaceMix results with no population text overlays.**
Figure showing SpaceMix tests for the **masked** and imputed Only-BSP dataset. X- and Y-axes are latitude and longitude in the geogenetic space, respectively. Panel showing the results of each IBD model, and the assumptions of each model were: (**a**) No migration and no admixture; (**b**) No migration but admixture; (**c**) Migration but no admixture; and (**d**) Both migration and admixture.

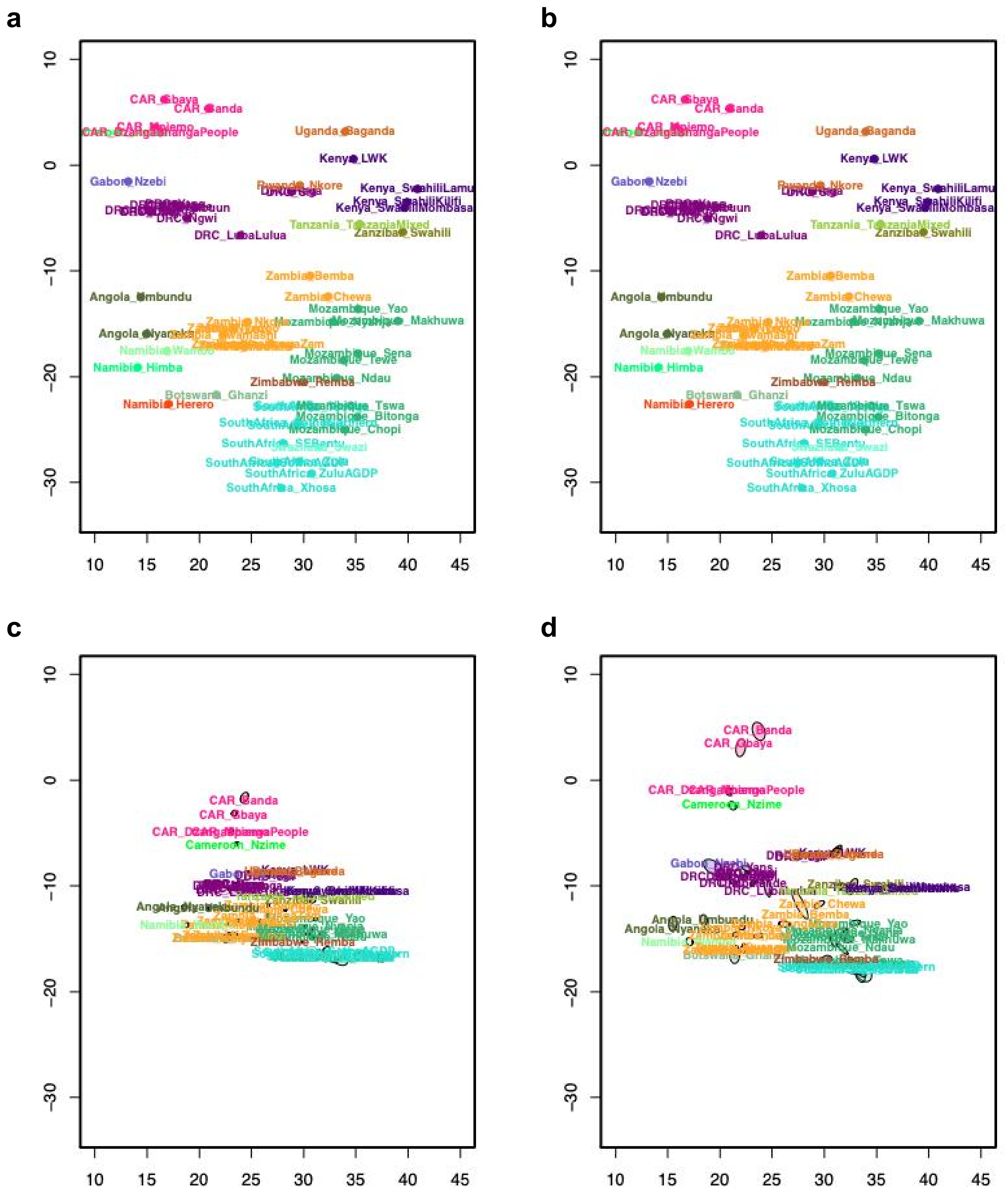


**Fig.S 7.6 | SpaceMix results with text instead of dots.**

Figure showing SpaceMix tests for the **masked** and imputed Only-BSP dataset with text overlay for each population. X- and Y-axes are latitude and longitude in geogenetic space respectively. Panel showing the results of each IBD model, and the assumptions of each model were; (**a**) No migration and no admixture; (**b**) No migration but admixture; (**c**) Migration but no admixture; and (**d**) Both migration and admixture.

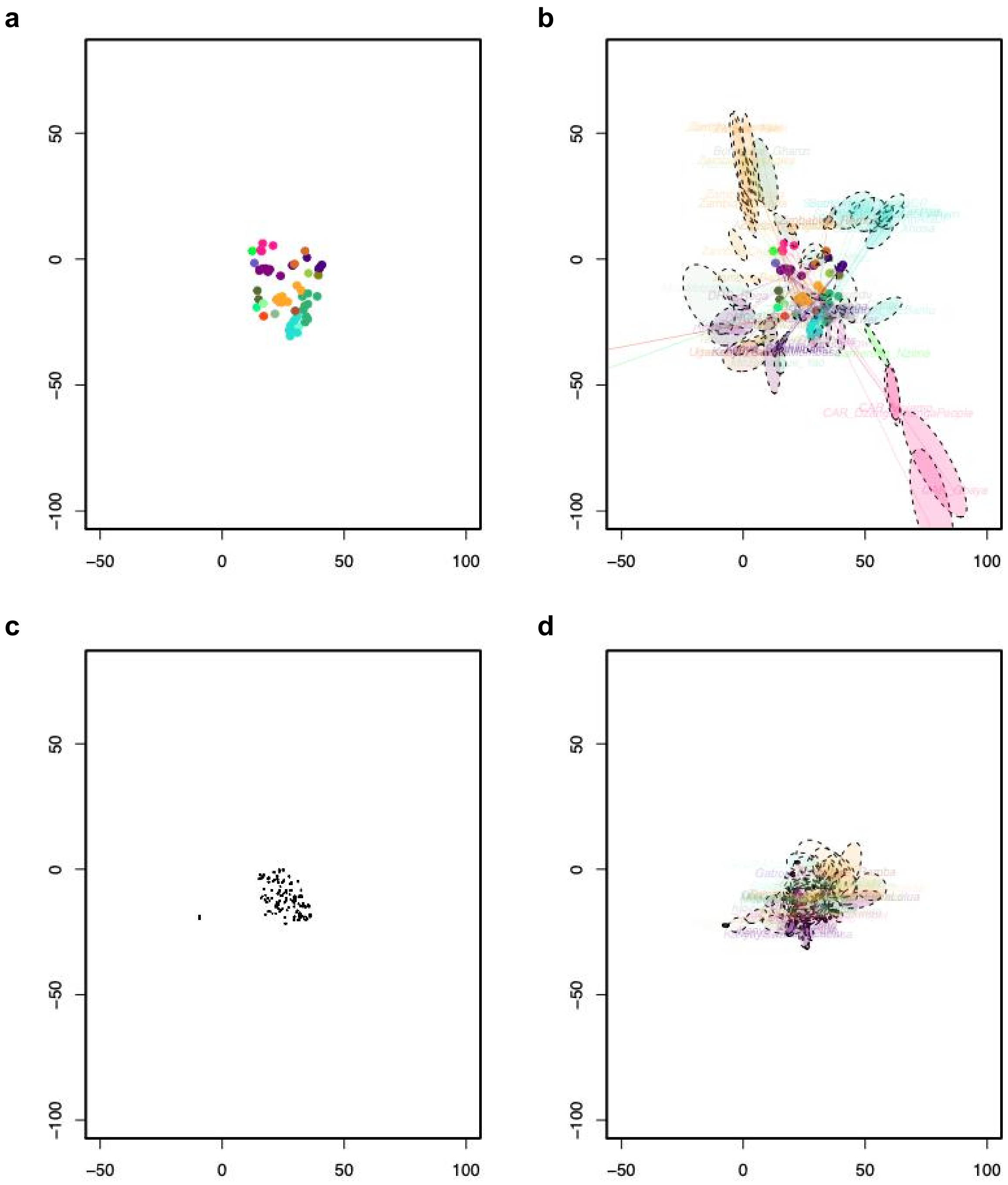


**Fig.S 7.7 | SpaceMix results with sources of admixture.**
Figure showing SpaceMix tests for the **masked** and imputed Only-BSP dataset with sources of admixture indicated with a dashed ellipsis. X- and Y-axes are latitude and longitude in the geogenetic space, respectively. Panel showing the results of each IBD model, and the assumptions of each model were: (**a**) No migration and no admixture; (**b**) No migration but admixture; (**c**) Migration but no admixture; and (**d**) Both migration and admixture.

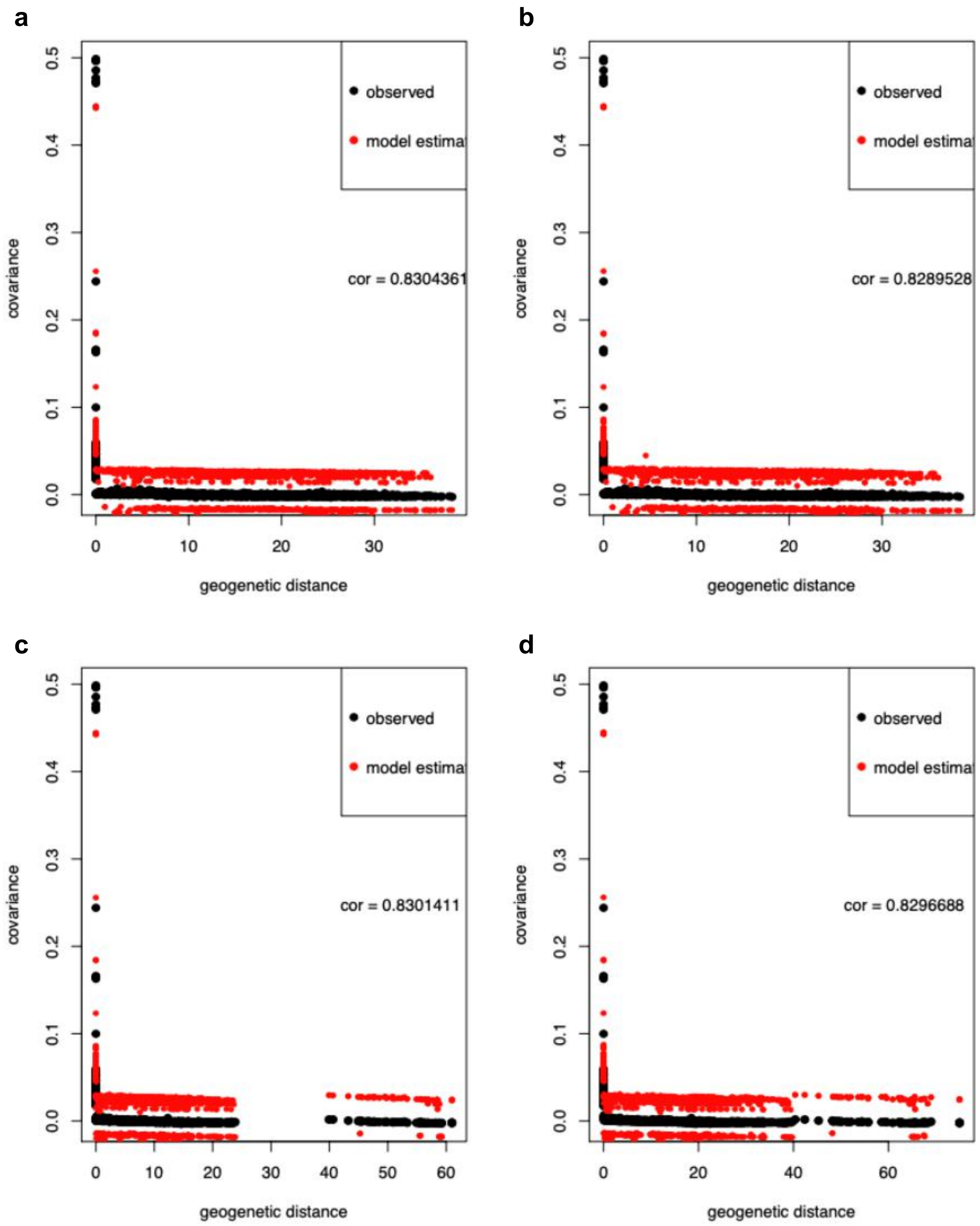


**Fig.S 7.8 | SpaceMix results with correlations of each model.**
Figure showing SpaceMix tests for the **masked** and imputed Only-BSP dataset, highlighting the correlation between the observed data and the data estimated from the model. “Cor” means Pearson correlation between the two series. Panel showing the results of each IBD model, and the assumptions of each model were: (**a**) No migration and no admixture; (**b**) No migration but admixture; (**c**) Migration but no admixture; and (**d**) Both migration and admixture.

#### 8- Patterns of haplotype diversity

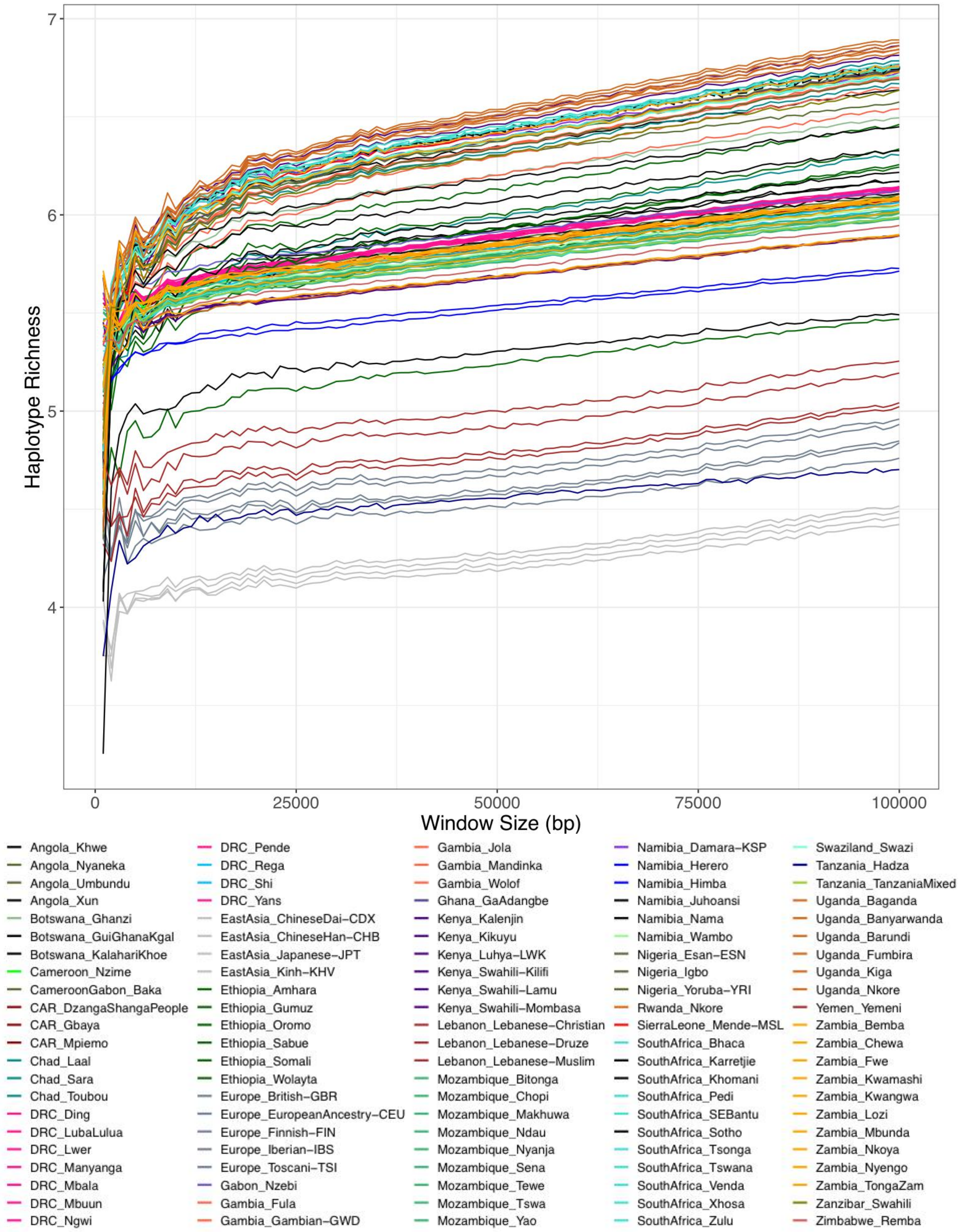


**Fig.S 8.1 | Haplotype richness (HR) for the AfricanNeo dataset.**
Figure showing estimated haplotype richness for the unmasked AfricanNeo dataset, HR for the y-axis and the window size (S) for the X-axis.

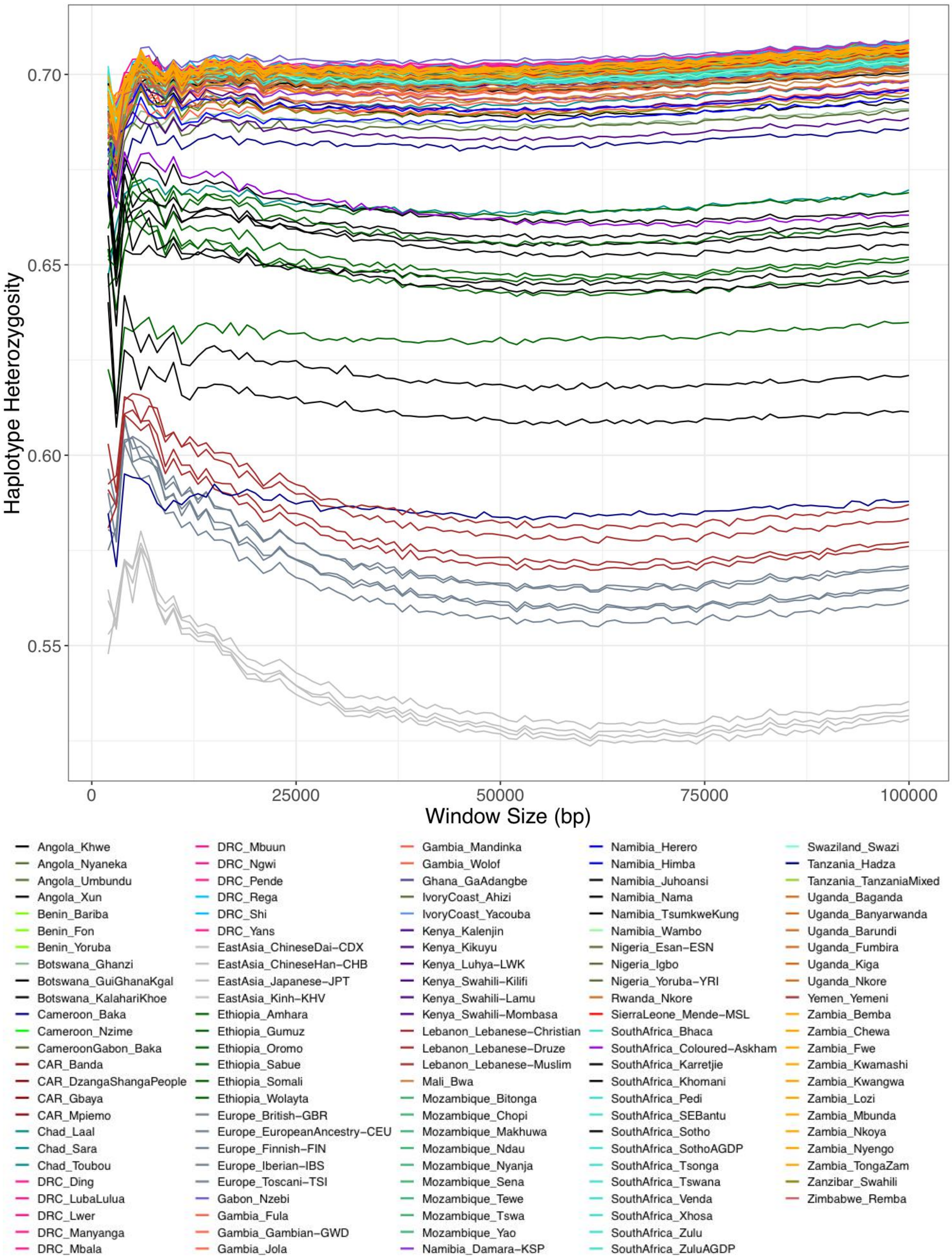


**Fig.S 8.2 | Haplotype heterozygosity (HH) for the AfricanNeo dataset.**
Figure showing estimated haplotype heterozygosity for the unmasked AfricanNeo dataset, HH on the y-axis and the window size (S) on the X-axis.

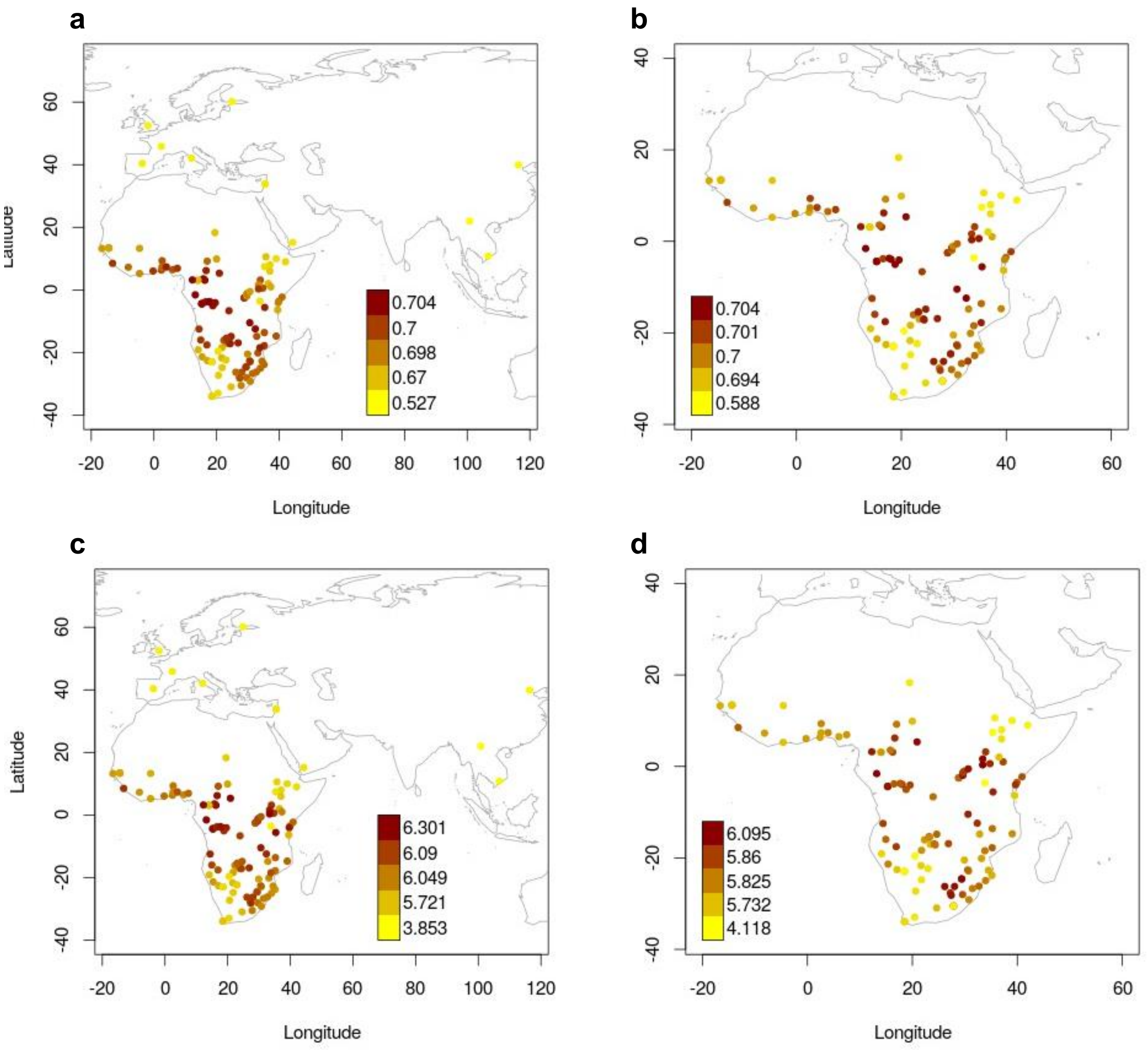


**Fig.S 8.3 | Maps of haplotype heterozygosity and haplotype richness for the AfricanNeo data.** Geographical distribution of haplotype heterozygosity and haplotype richness estimated for a window size of (S) 50 kb on the basis of populations included in the unmasked AfricanNeo dataset. Figure showing: (**a**) HH results for all studied populations included in the AfricanNeo dataset; (**b**) HH results for sub-Saharan African populations included in the Only-African dataset; (**c**) HR results for all studied populations included in the AfricanNeo dataset; and (**d**) HR results for sub-Saharan African populations included in the Only-African dataset.

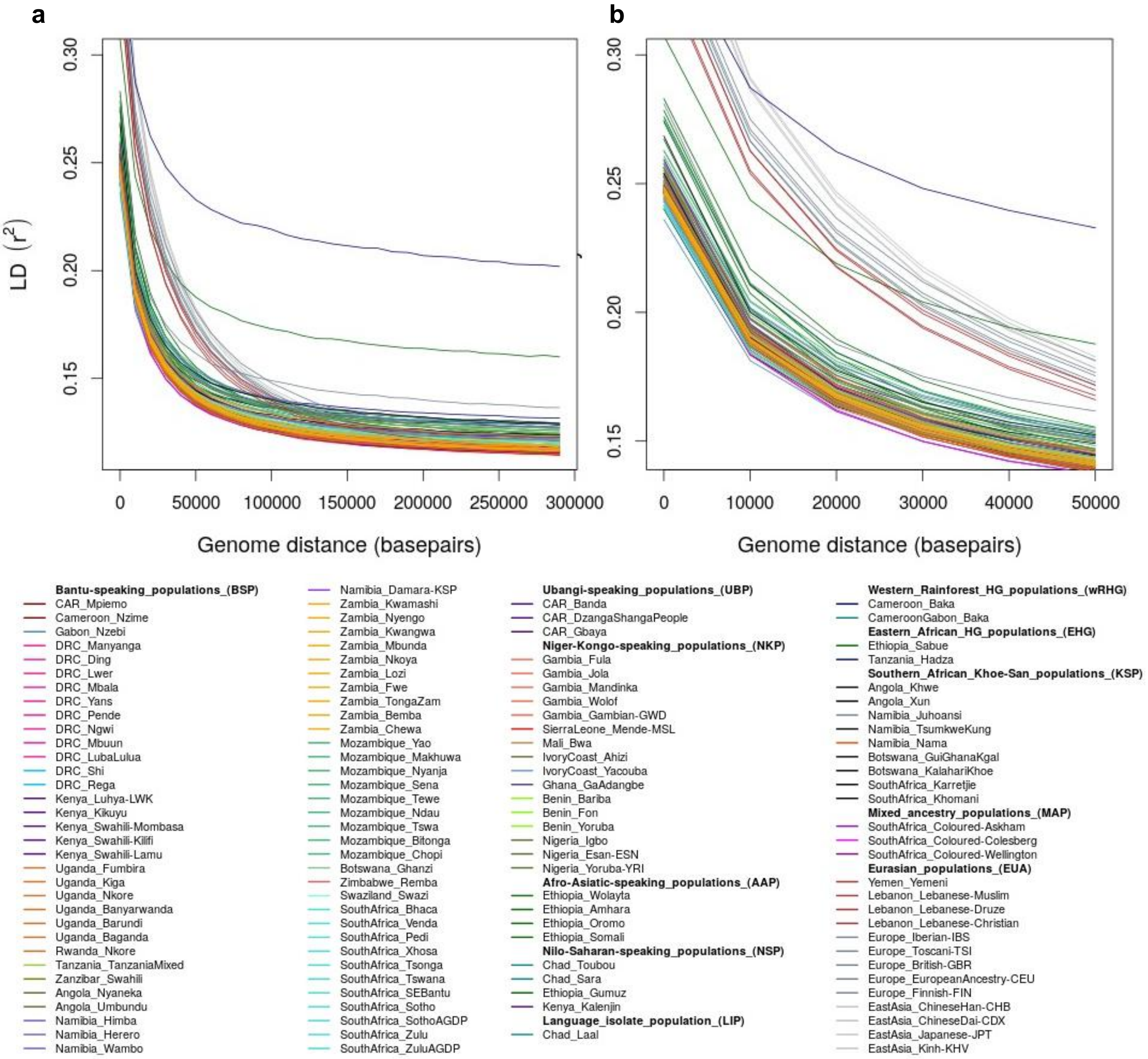


**Fig.S 8.4 | Linkage-disequilibrium (LD)-decay of the unmasked AfricanNeo dataset.**
(**a**) LD-decay with genomic distance of populations included in the unmasked AfricanNeo dataset (124 African and Eurasian populations). LD-decay figure was built from the mean value of $r^2$ for pairs of sites in 30-distance bins. Eurasian populations have lower long-distance LD than some African populations. (**b**) Figure after zooming into a short distance of LD decay of up to 50 kb.

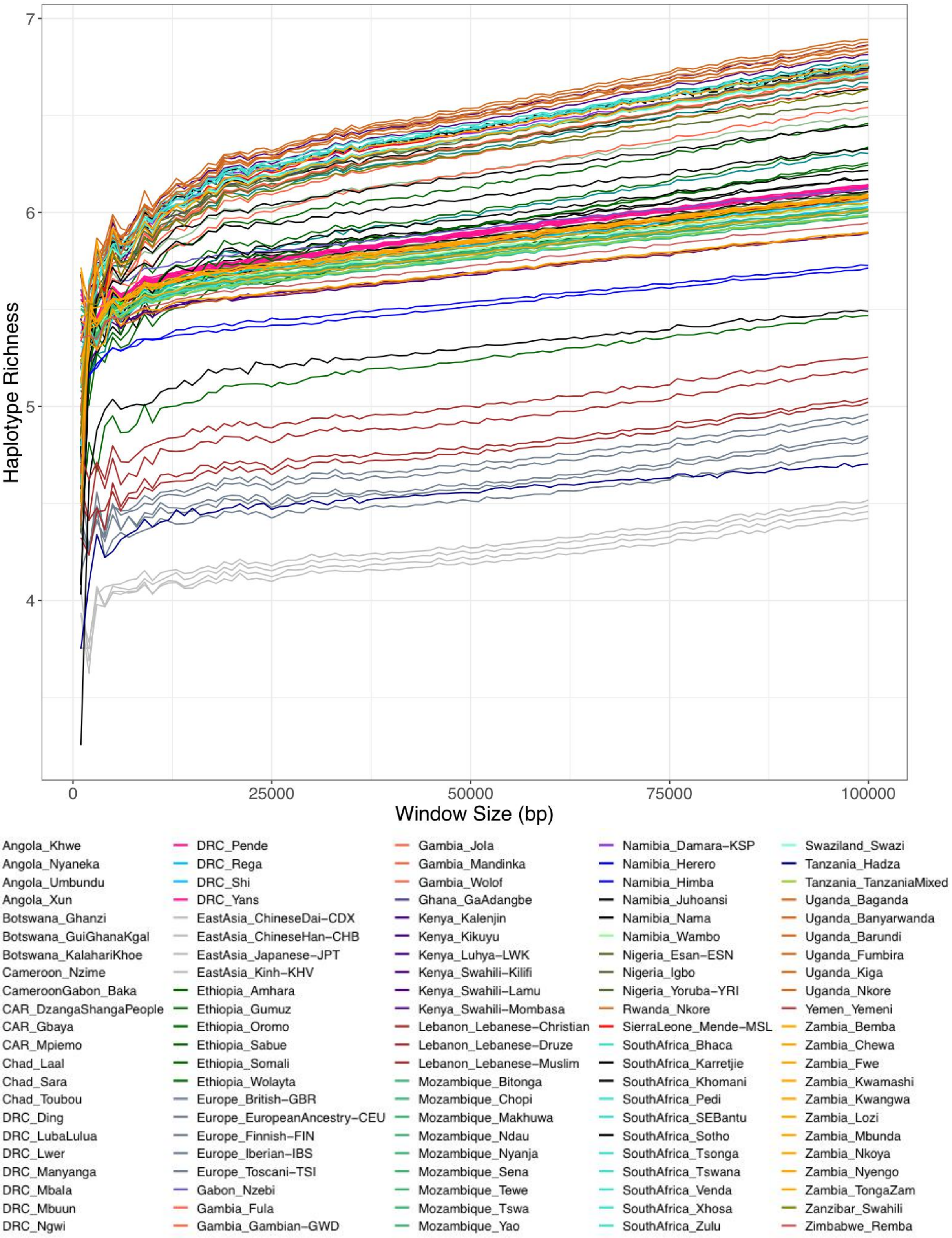


**Fig.S 8.5 | Haplotype richness (HR) with masked AfricanNeo dataset.**
Figure showing estimated haplotype richness with masked data of BSP and unmasked data of Eurasian populations included in the AfricanNeo dataset, HH on the y-axis and the window size (S) on the x-axis.

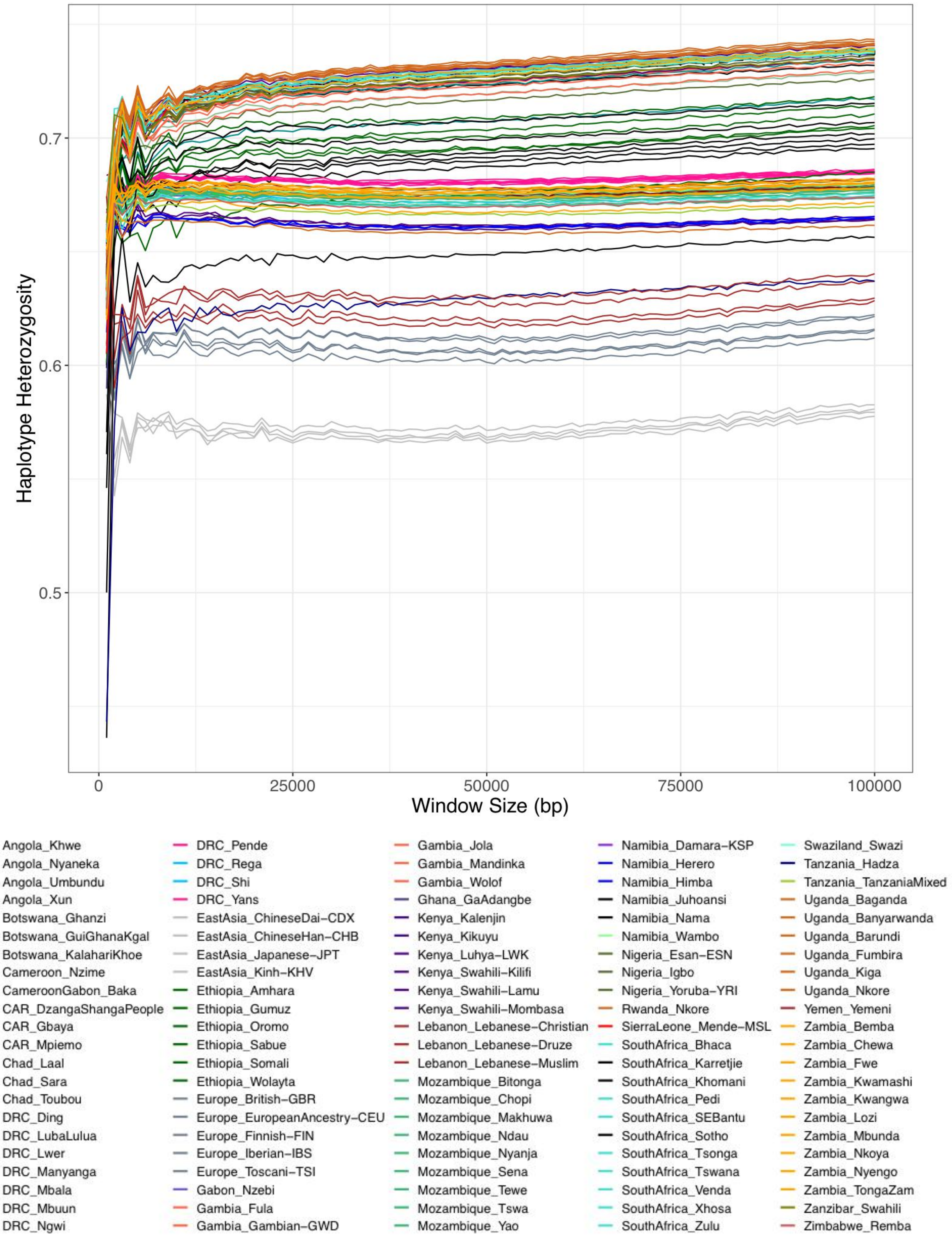


**Fig.S 8.6 | Haplotype heterozygosity (HH) with masked AfricanNeo dataset.**
Figure showing estimated haplotype heterozygosity (HH) with masked data of BSP and unmasked data of Eurasian populations included in the AfricanNeo dataset, HH on the y-axis and the window size (S) on the x-axis.

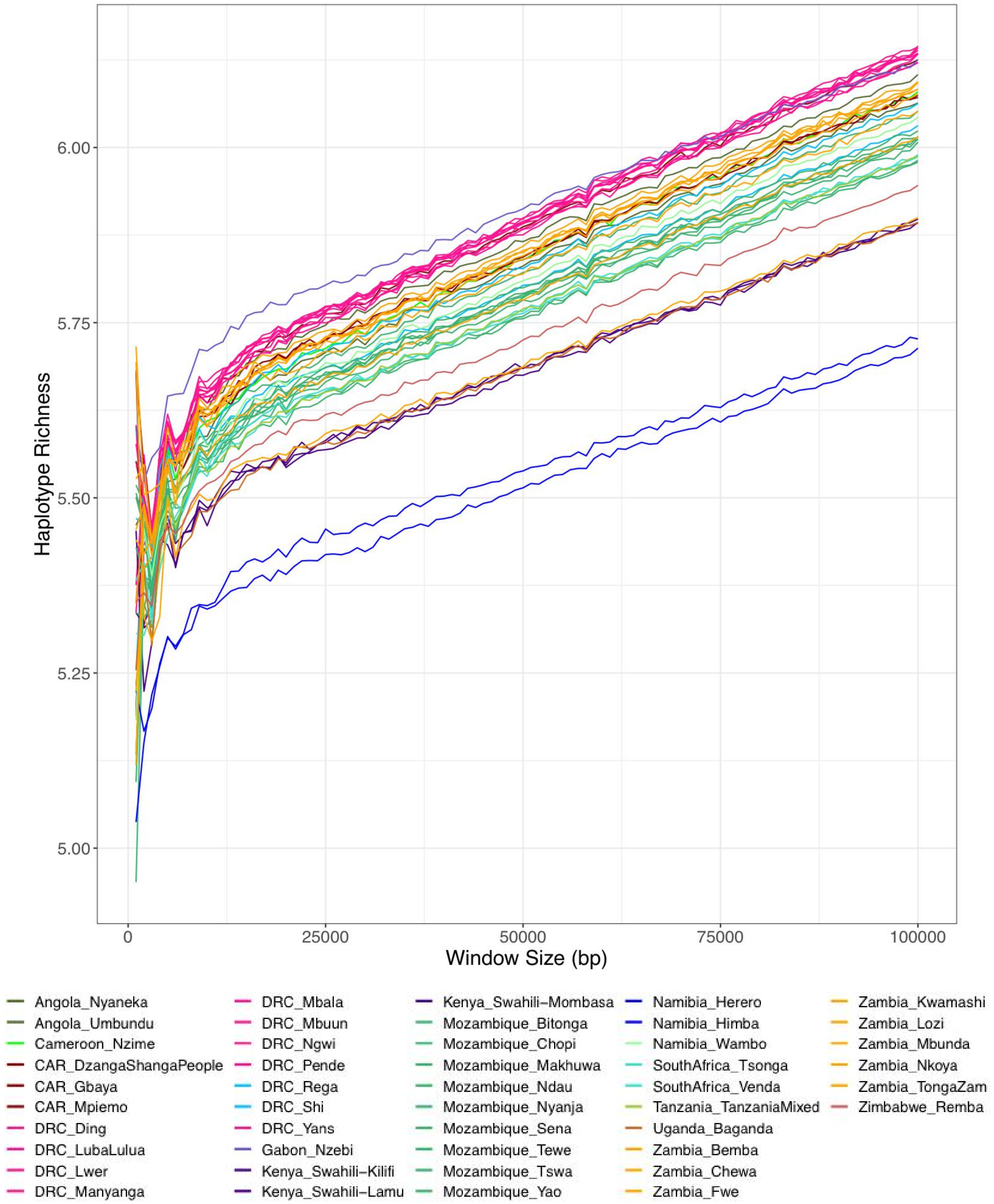


**Fig.S 8.7 | Haplotype richness (HR) for the masked Only-BSP dataset.**
Figure showing estimated haplotype richness for populations included in the masked Only-BSP dataset, HR on the y-axis and the window size (S) on the x-axis.

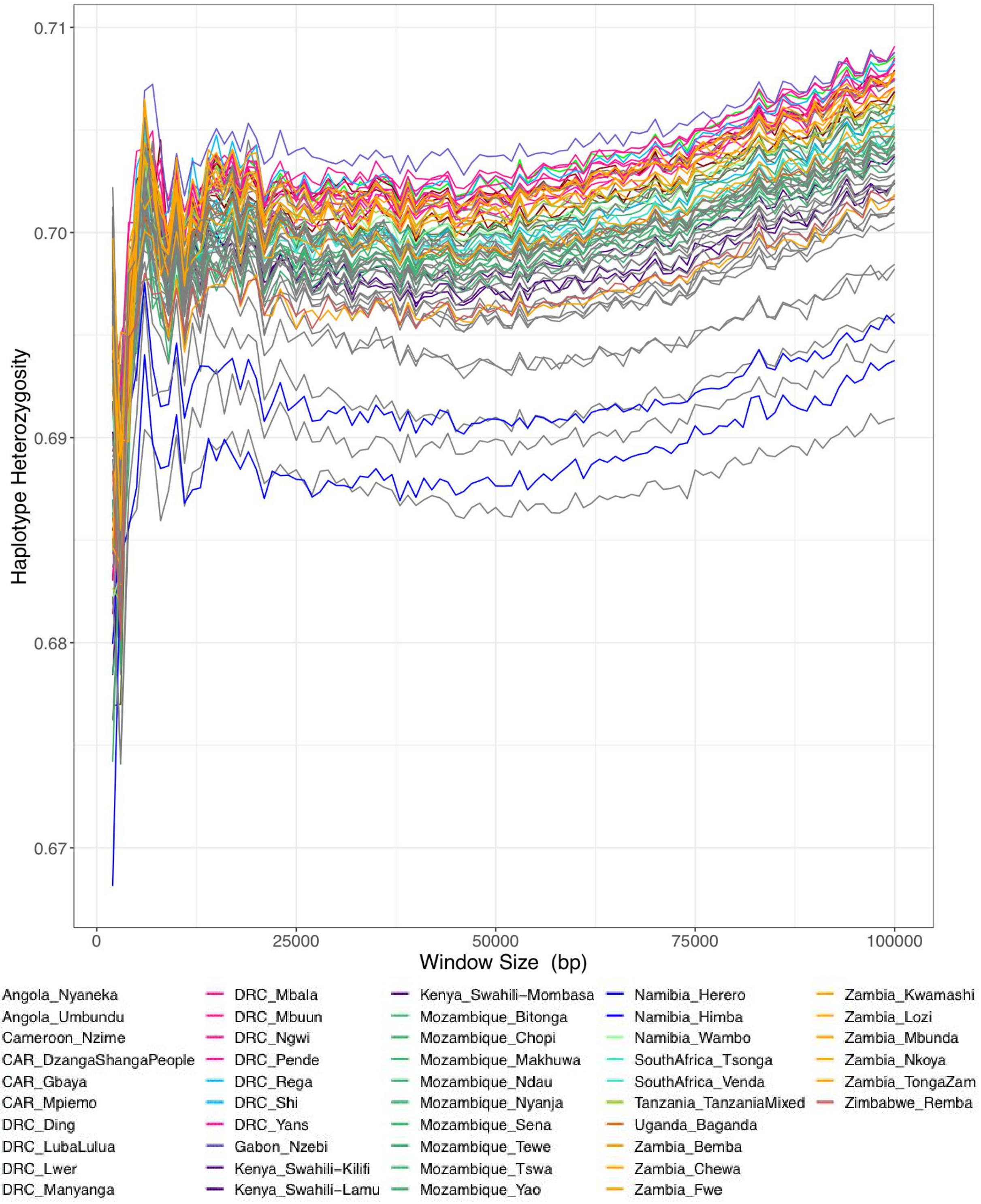


**Fig.S 8.8 | Haplotype heterozygosity (HH) for the masked Only-BSP.**
Figure showing estimated haplotype heterozygosity for populations included in the masked Only-BSP dataset, HH on the y-axis and the window size (S) on the x-axis.

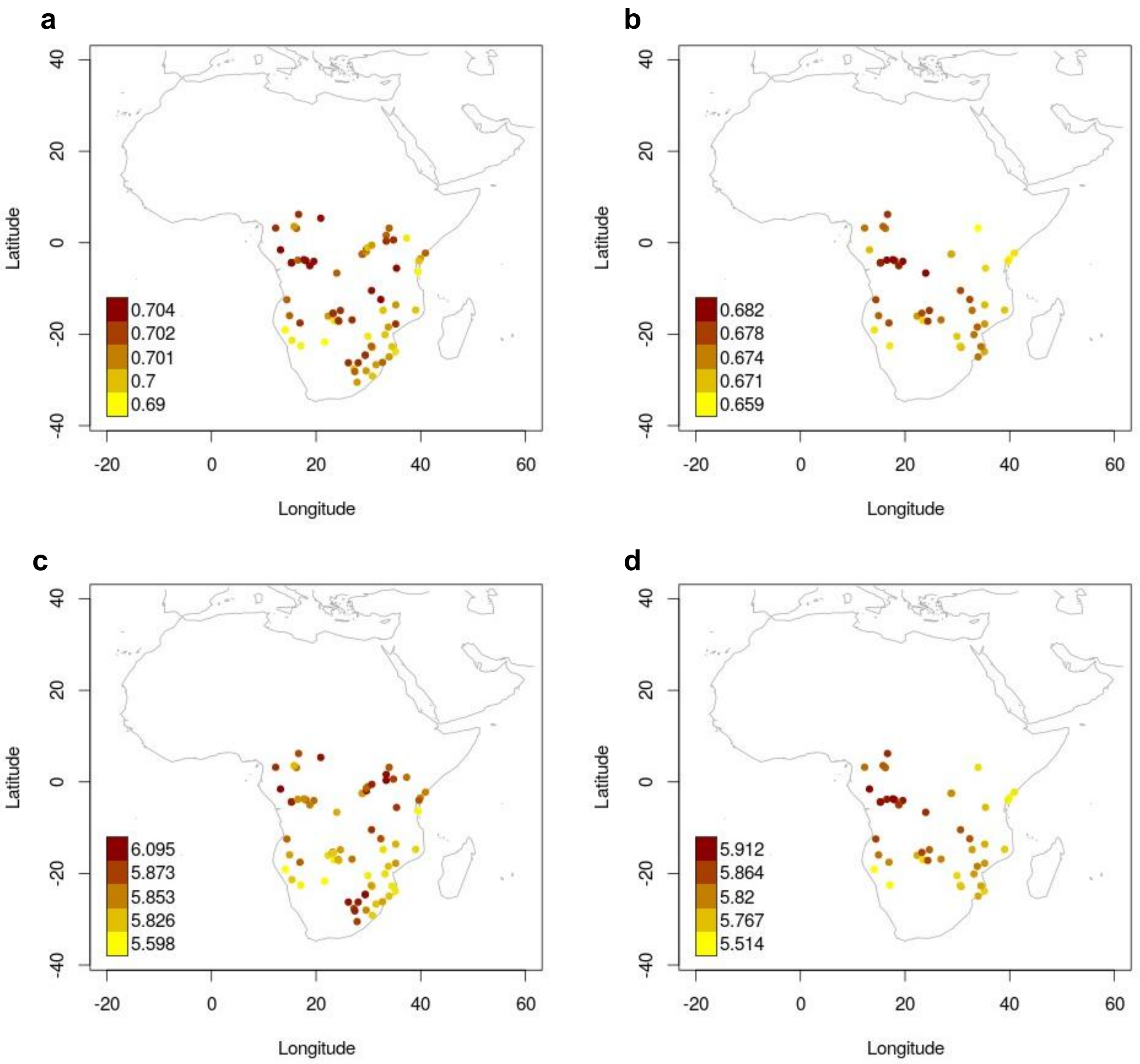


**Fig.S 8.9 | Maps of haplotype heterozygosity and haplotype richness for only BSP.**
Geographical distribution of haplotype heterozygosity and haplotype richness were estimated for BSP using a window size (S) of 50 kb. Figure showing (**A**) HH results for the unmasked Only-BSP dataset; (**B**) HH results for the masked Only-BSP dataset; (**C**) HR results for the unmasked Only-BSP dataset; and (**D**) HR results for the masked Only-BSP dataset.

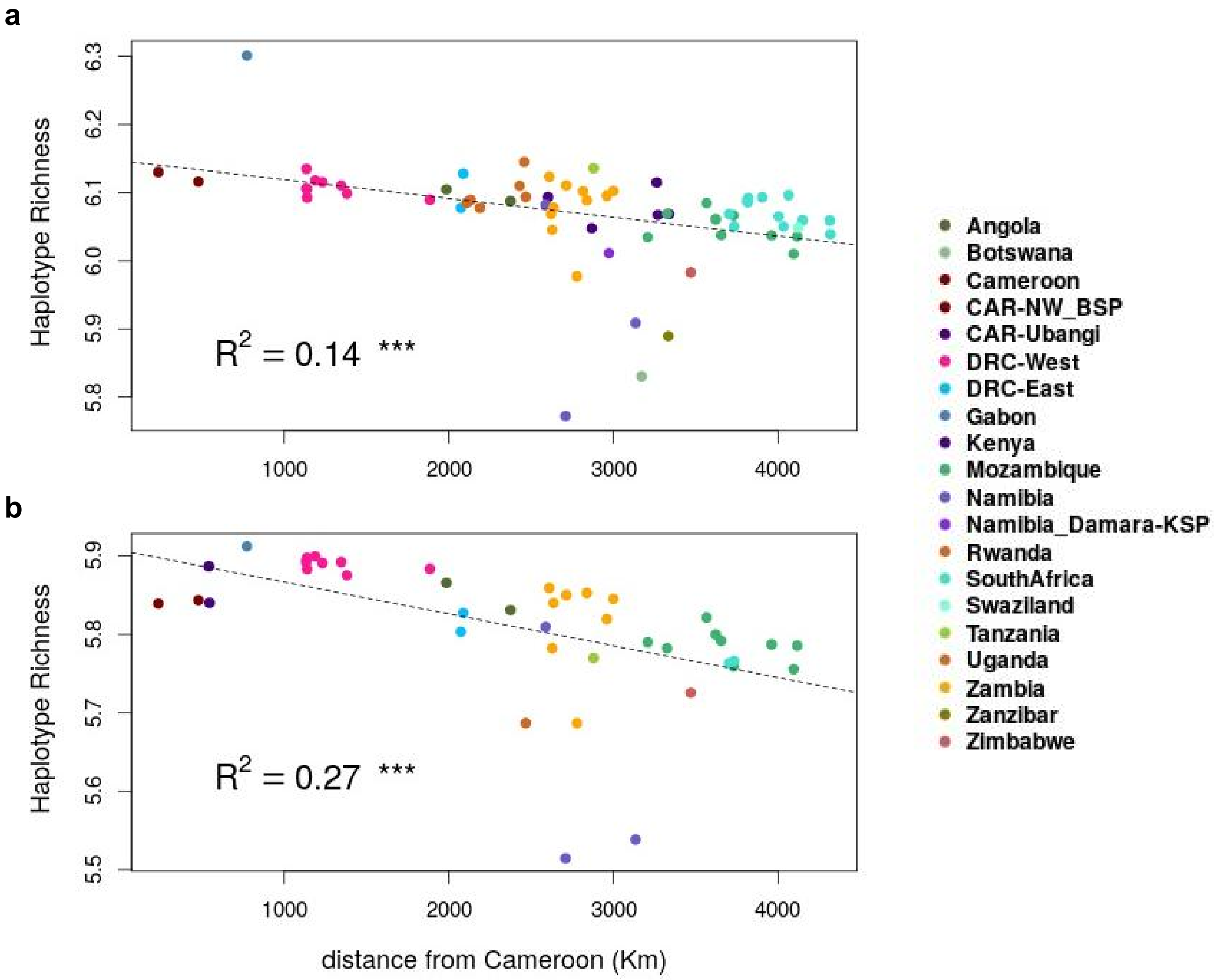


**Fig.S 8.10 | Haplotype richness plotted against distance from Cameroon.**
Figure showing haplotype richness (HR) values were plotted against the distance from Cameroon for the BSP in the dataset, calculated on the basis of (**a**) the unmasked Only-BSP dataset and (**b**) the masked Only-BSP dataset.

a

b

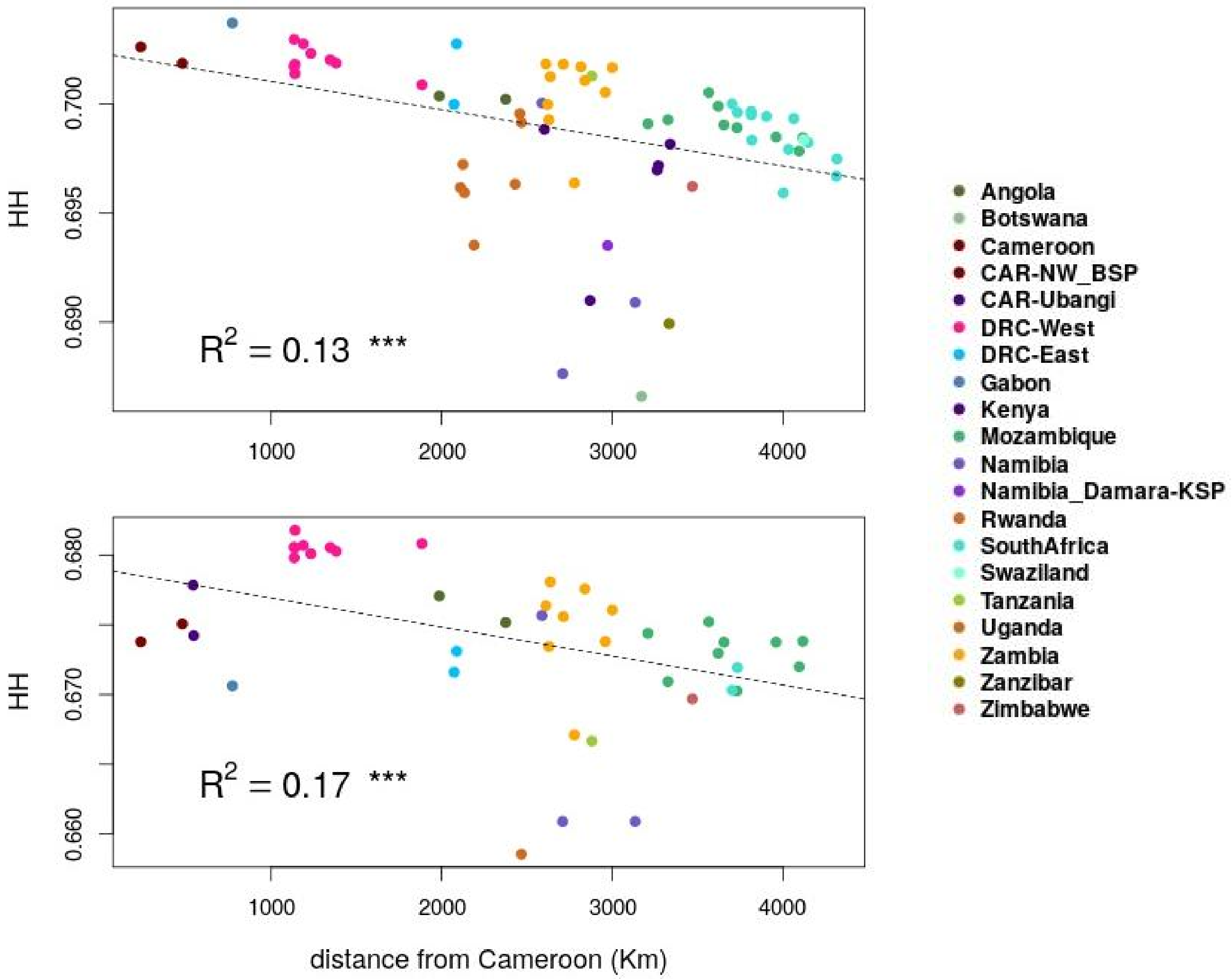


**Fig.S 8.11 | Haplotype heterozygosity plotted against distance from Cameroon.**
Figure showing haplotype heterozygosity (HH) plotted against distance from Cameroon for the BSP in the dataset, calculated on the basis of (**a**) the unmasked Only–BSP data and (**b**) the masked Only–BSP dataset.

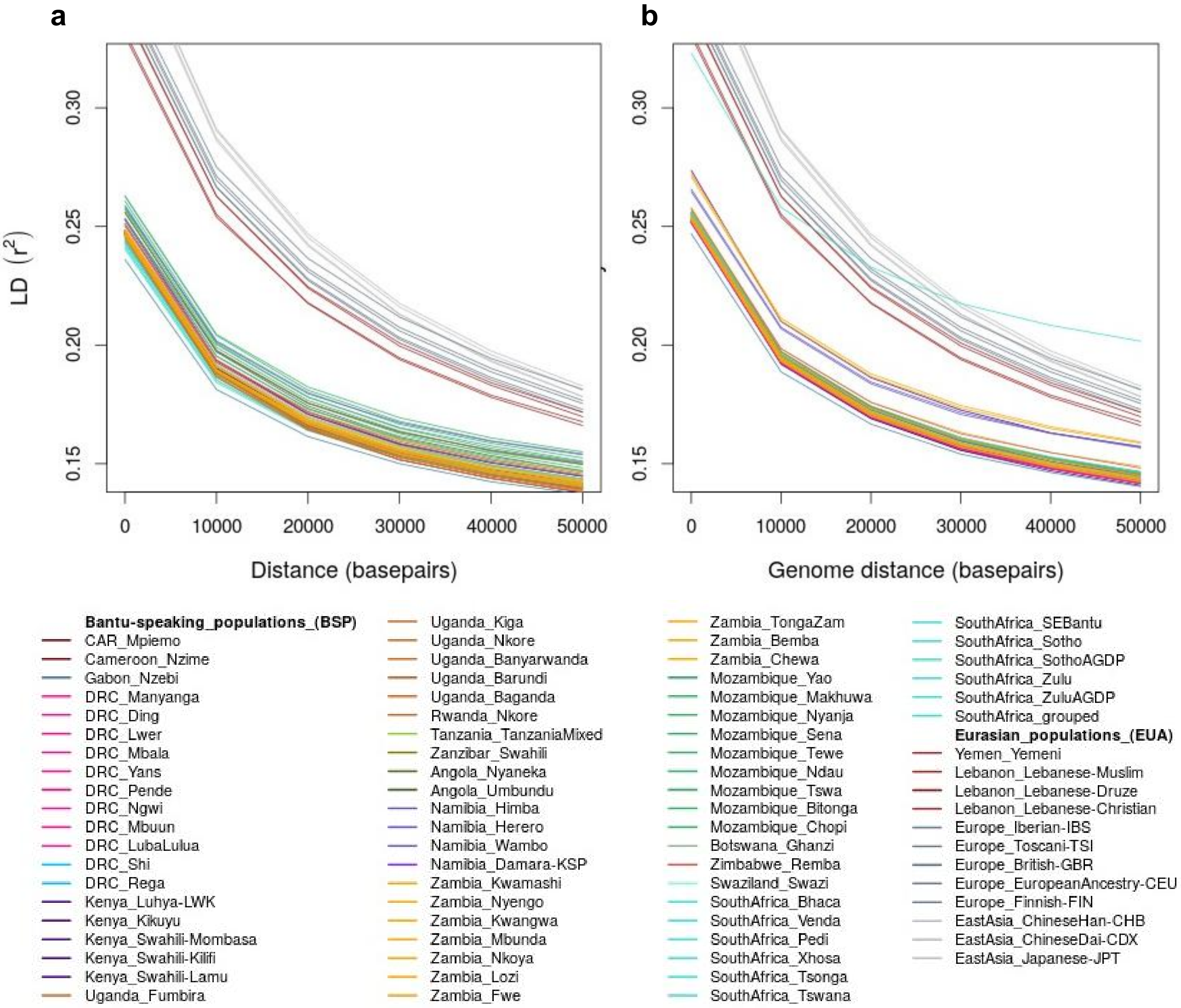


**Fig.S 8.12 | Linkage-disequilibrium (LD)-decay of Only-BSP dataset.**
LD-decay with genomic distance for BSP and Eurasian reference populations from the (**a**) unmasked Only-BSP dataset (N= 67); and (**b**) the masked Only-BSP dataset (N= 51) that includes BSP with at least 70% West-Central African (WCA) ancestry. For this plot, the minimum sample size was set to 7 individuals (e.g. Pedi population from South Africa), and South African populations with less than 3 individuals were grouped into one group called "SouthAfrica_grouped" (N= 11). Results for the masked Only-BSP dataset show slightly higher LD-decay in comparison with the unmasked Only-BSP dataset.

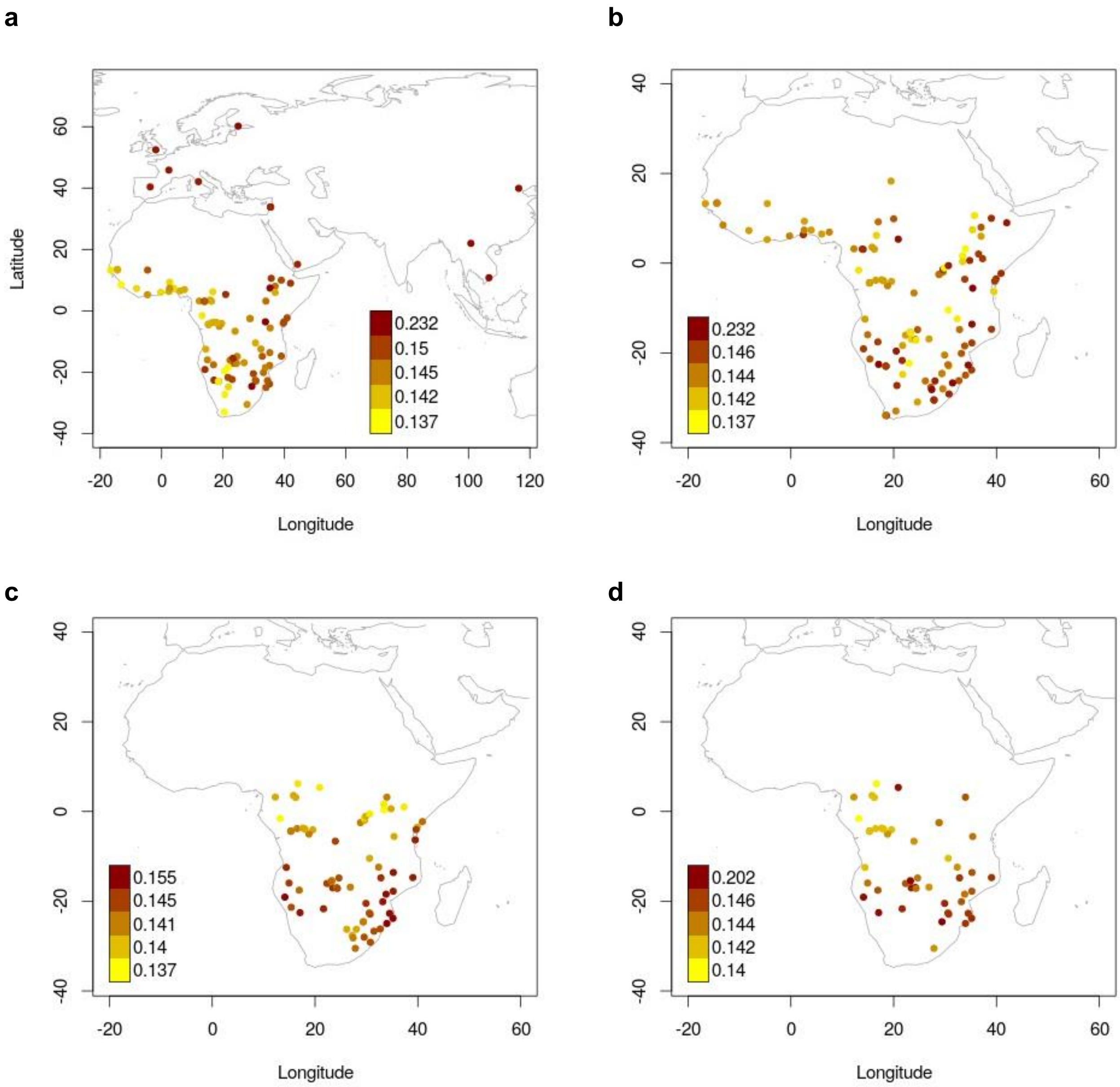


**Fig.S 8.13 | Spatial distribution of LD-decay in each studied dataset.**
Spatial distribution of linkage-disequilibrium (LD) estimates as $r^2$ at 50 Kb. Figure showing the results for (**a**) BSP and worldwide reference populations included in the unmasked AfricaNeo dataset; (**b**) Unmasked Only-African dataset; (**c**) Unmasked Only-African dataset of selected African populations (N= 70 populations; with a minimum sample size of 10 individuals) (also **Fig. 4A**). (**d**) Masked Only-BSP dataset (N= 49) that includes BSP with at least 70% of West-Central African-related ancestry (also **Fig. 4b**). For this plot, the minimum sample size was 7 individuals (SouthAfrica_Pedi) and South African populations with less than 3 samples were grouped into one group called "SouthAfrica_grouped" (N = 11; and the position of this group was set as latitude= -30.51 and longitude= 27.84). Values of the colour scale are the quantiles 0.0, 0.25, 0.50, 0.75, and 1.0 of $r^2$ distribution in each set of populations.

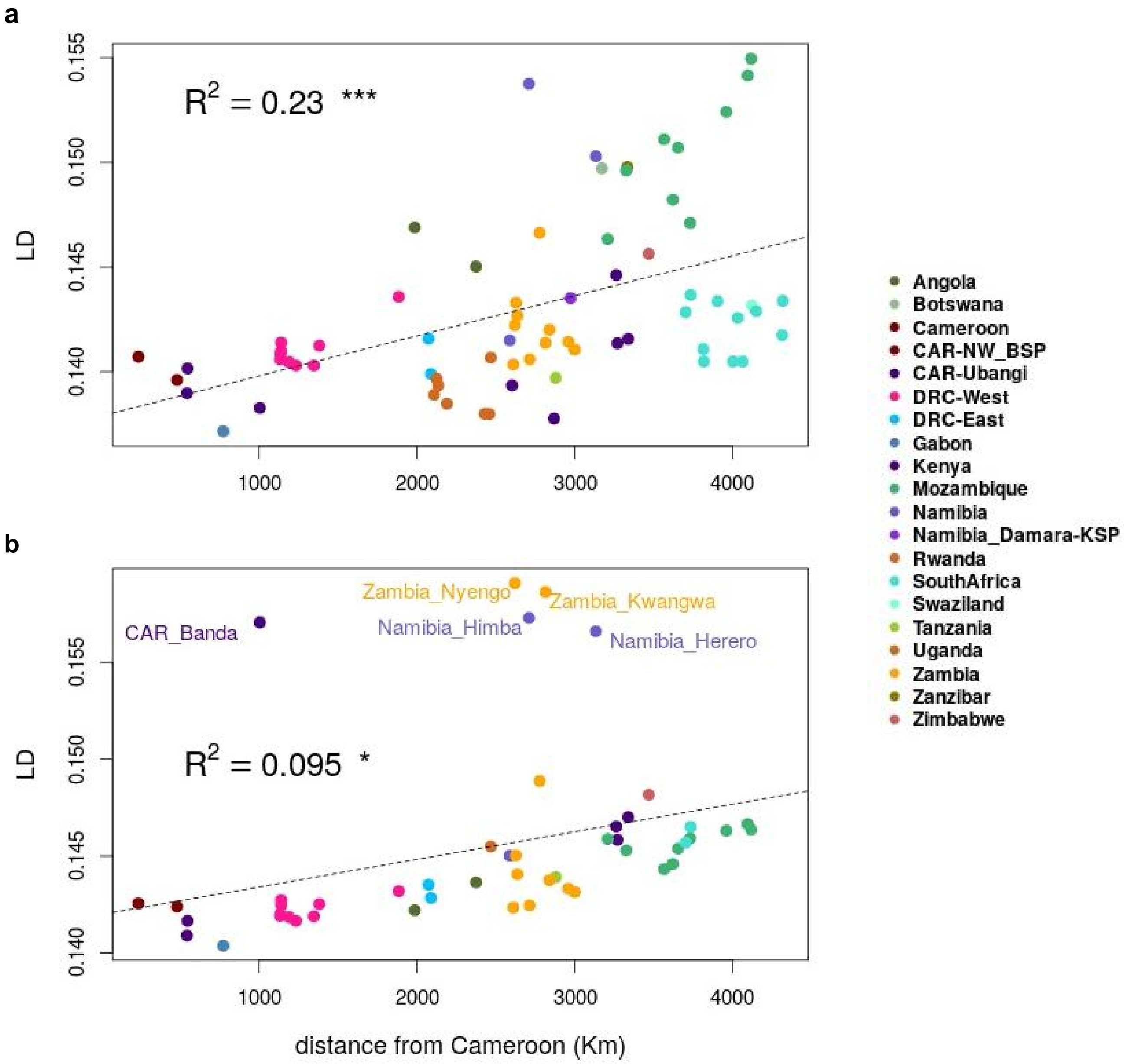


**Fig.S 8.14 | Increase of LD-decay with geographical distances in studied populations.**
Increase of LD estimates with geographical distance from Cameroon to the sampling location of each BSP and three Ubangi-speaking populations. (**a**) Unmasked dataset of selected populations (N= 70). (**b**) Masked dataset of selected populations (N= 49; SoutAfrica_Pedi with LD > 0.27 was excluded from this analysis). For each studied population, the LD value at 50kb was used in the correlation (see **Suppl. Materials**). Spatial distances were calculated as the spherical distance from each population and a centroid position located in the center of Cameroon. The dotted line represents the linear relationship between LD estimates and geographical distances.

**9- Maximum likelihood trees based on population allele frequencies**

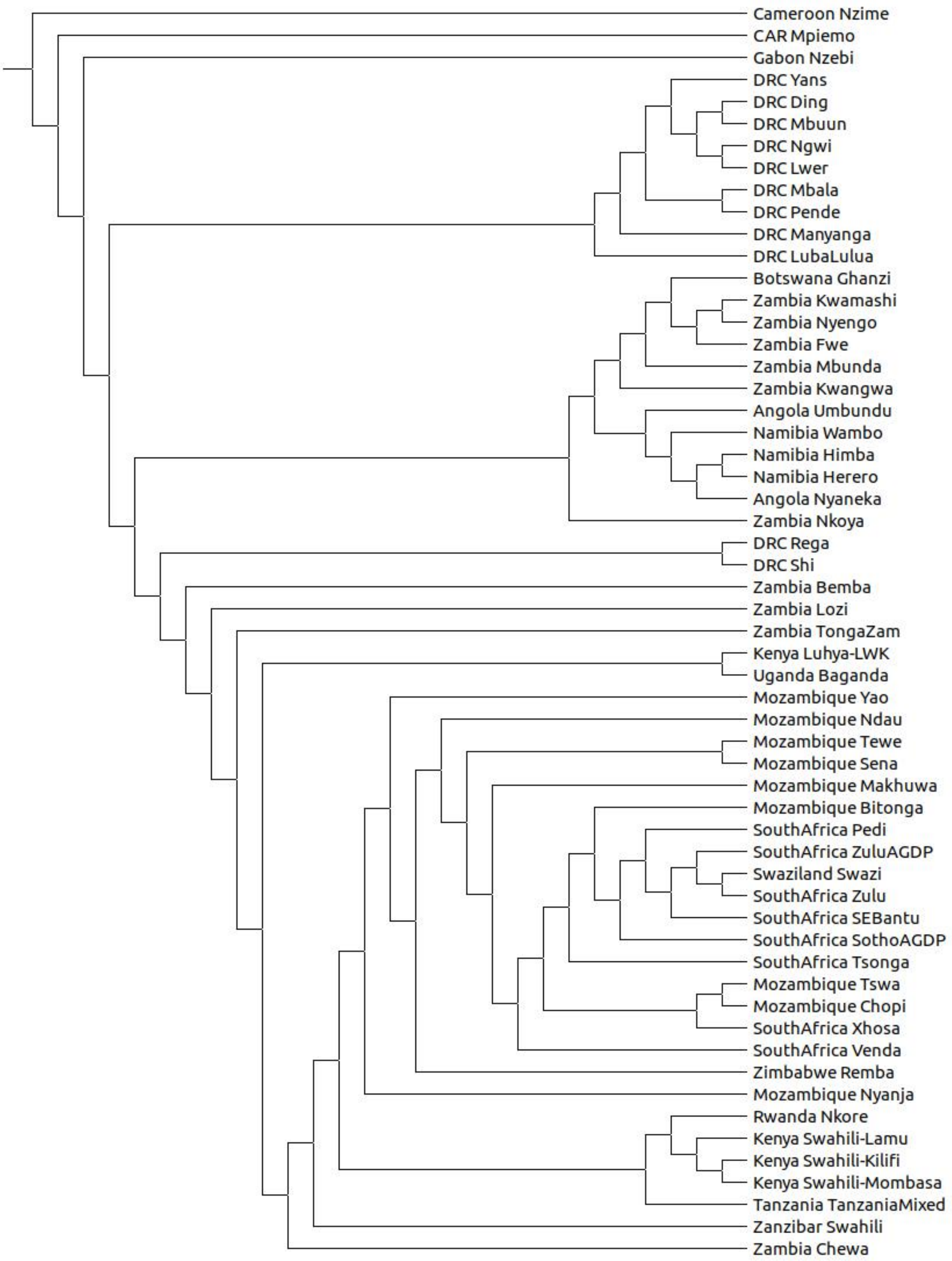


**Fig.S 9.1a | TreeMix for the masked Only-BSP dataset in rectangular shape.**
Figure showing population tree results on the basis of the masked and imputed Only-BSP dataset. Figure showing a tree in a rectangular shape.

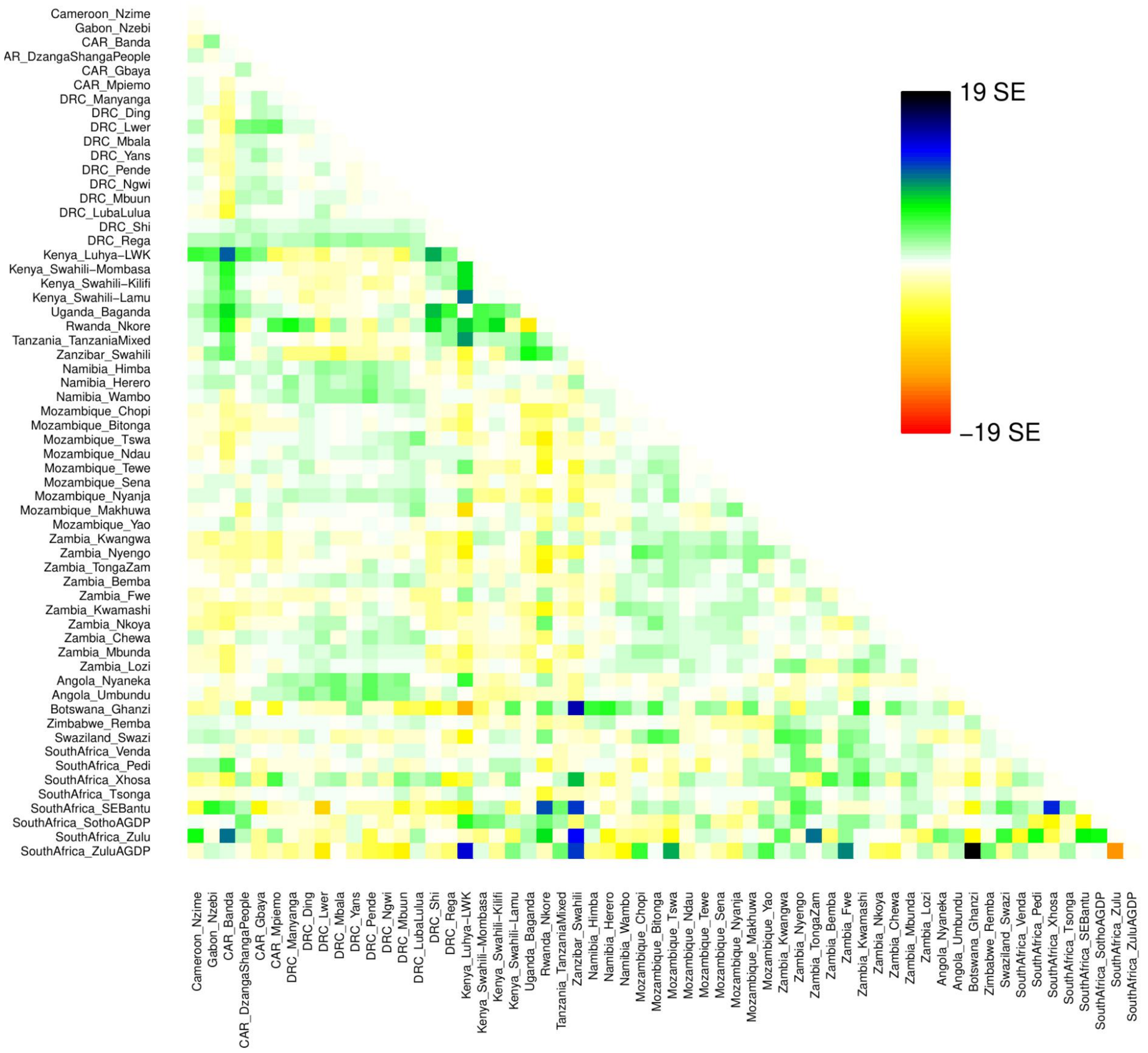


**Fig.S 9.1b | Coancestry matrix for the masked and imputed Only-BSP dataset.**
Figure showing coancestry matrix summarizing genetic differences between pairwise populations included in the masked Only-BSP dataset obtained using TreeMix default options for plotting.

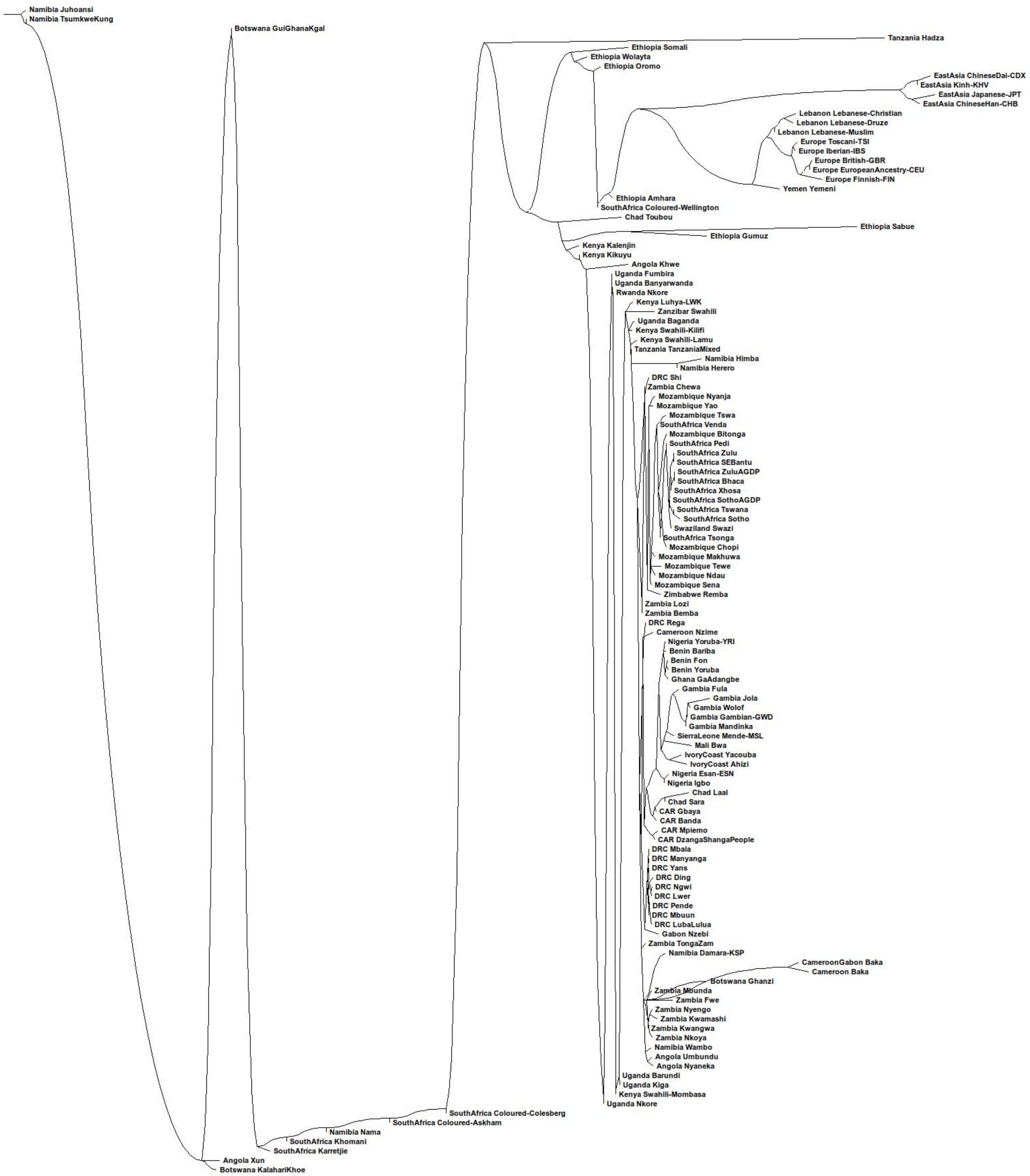


**Fig.S 9.2a | Population tree of the unmasked AfricanNeo dataset.**
Figure showing the tree was built from the covariance matrix of the population allele frequencies with TreeMix including all the populations of the unmasked AfricanNeo dataset.

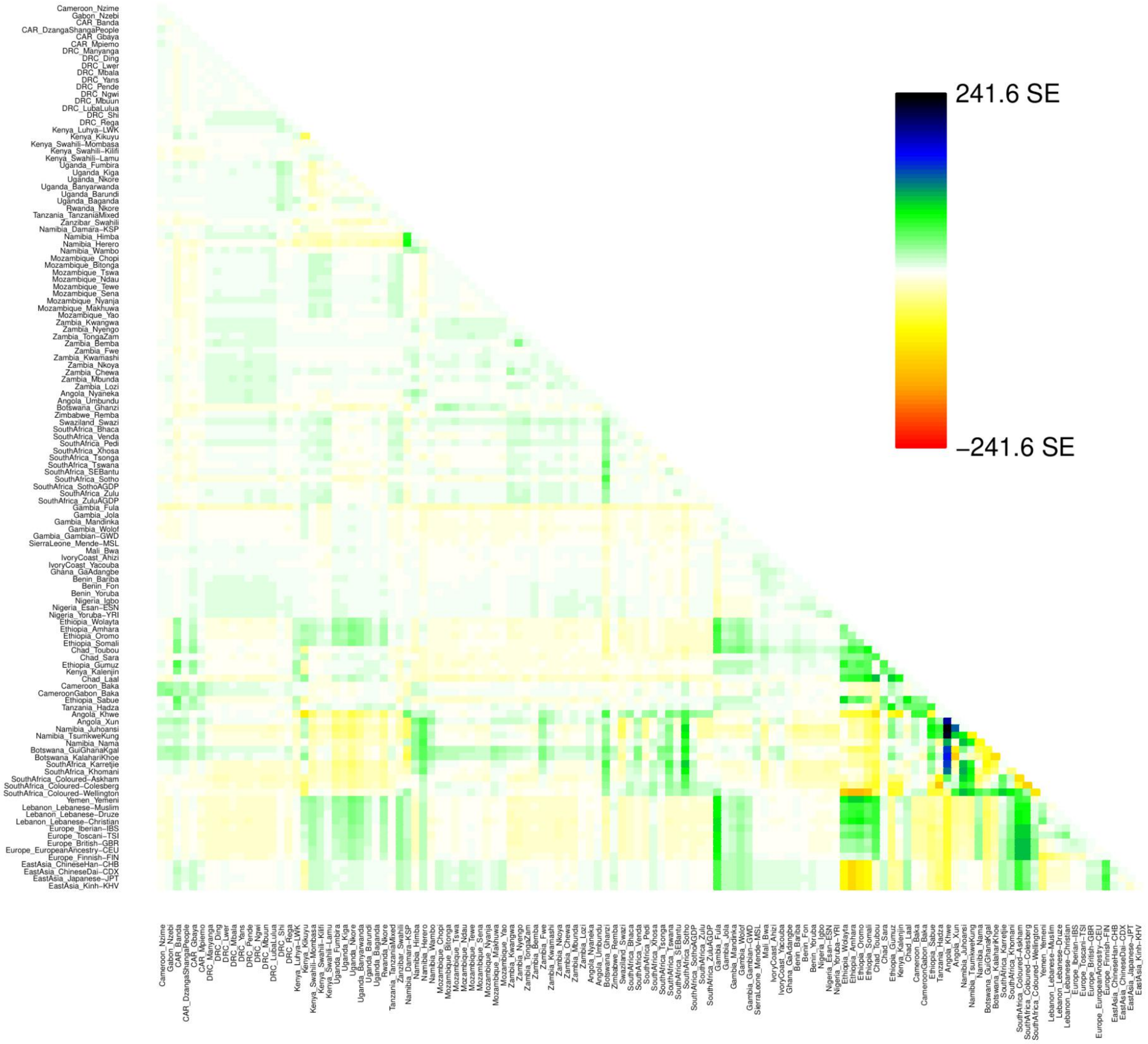


**Fig.S 9.2b | TreeMix results for the unmasked AfricanNeo dataset.**
Figure showing population tree results were plotted using default options in TreeMix.

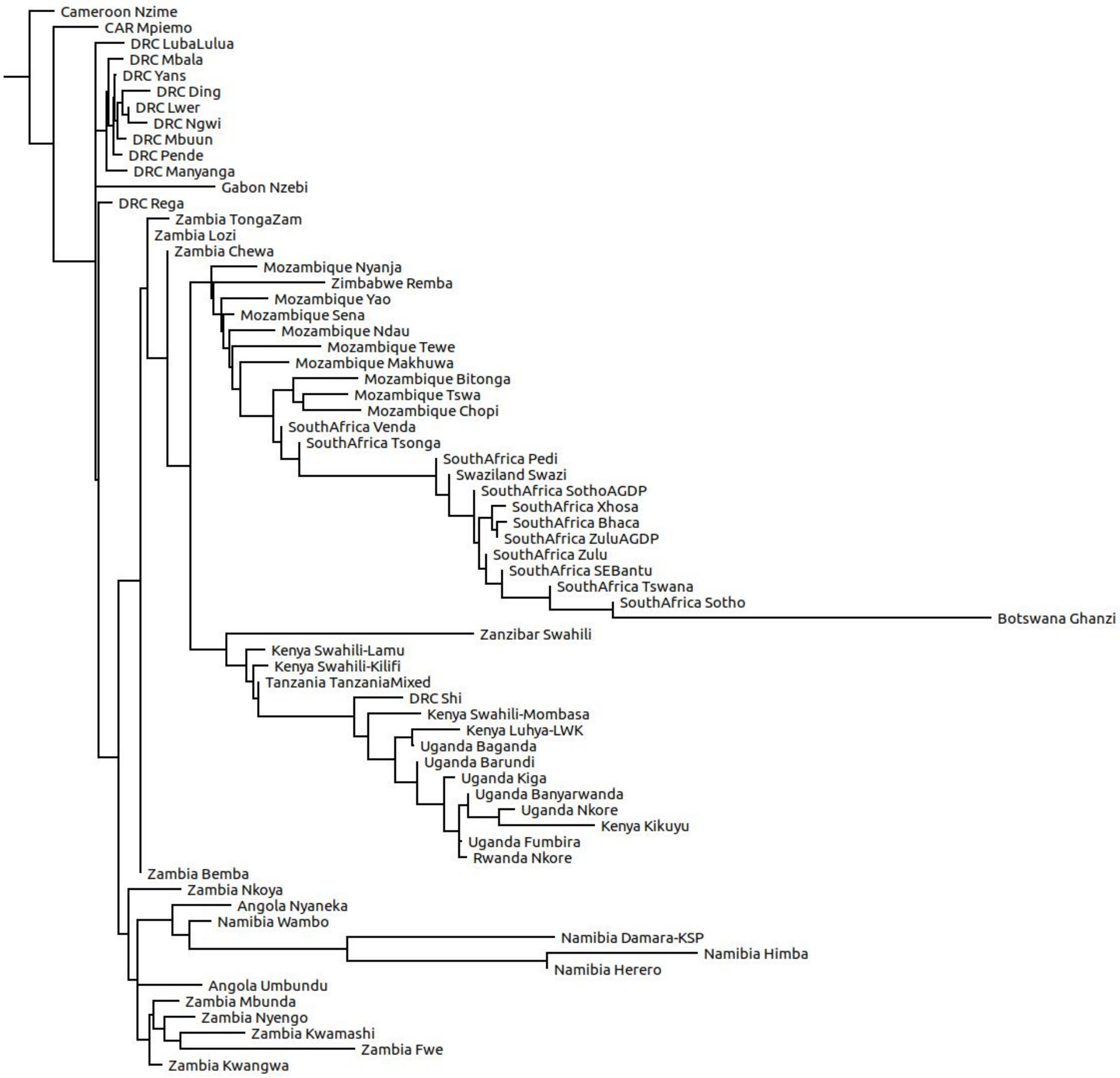


**Fig.S 9.3a | TreeMix for the unmasked Only-BSP dataset in rectangular shape.**
Figure showing population tree results for the unmasked Only-BSP dataset. Figure showing a tree in a rectangular shape.

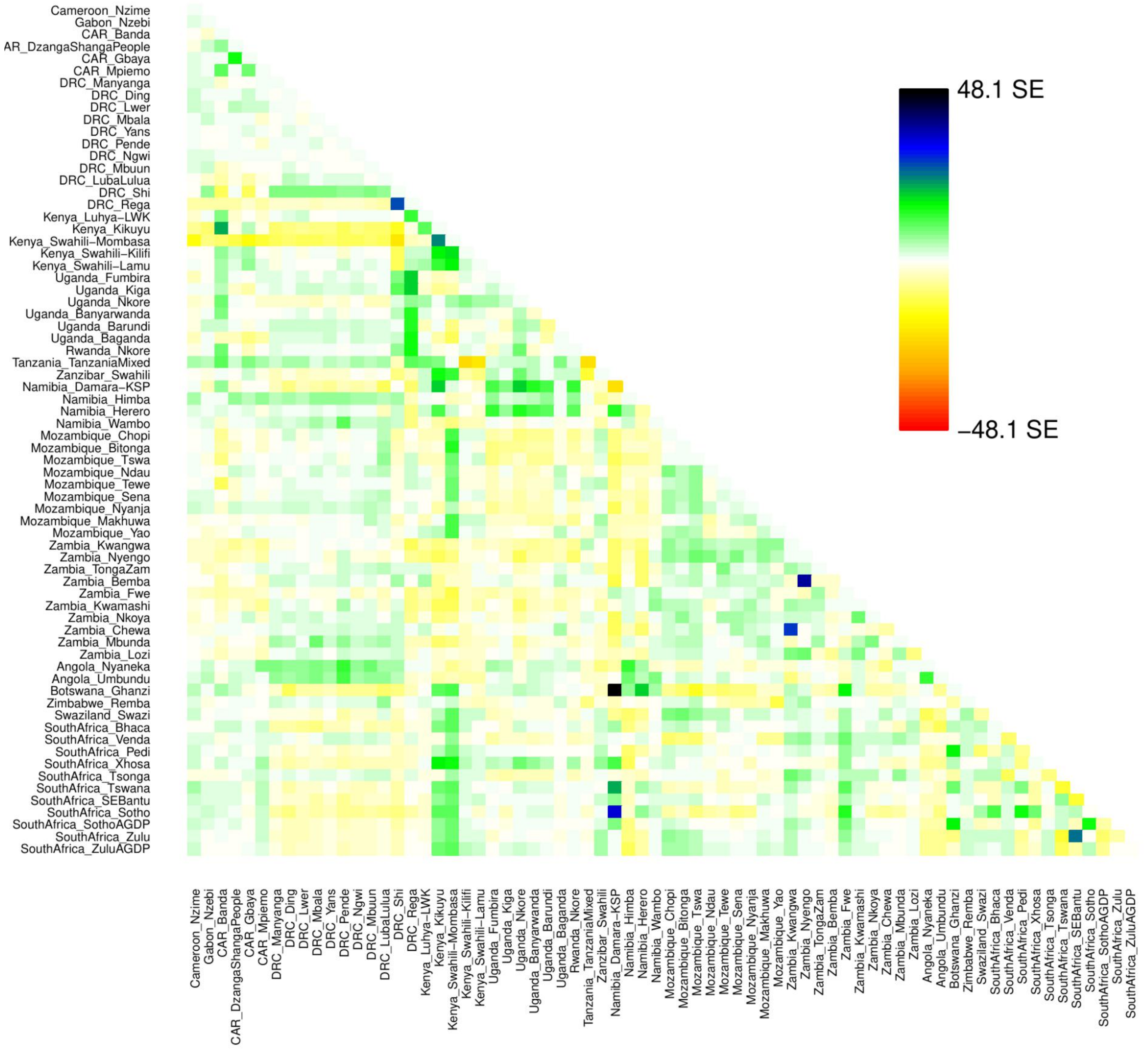


**Fig.S 9.3b | Coancestry matrix for the unmasked Only-BSP dataset.**
Figure showing coancestry matrix summarizing genetic differences between pairwise populations included in the unmasked Only-BSP dataset obtained using TreeMix default options for plotting.

**10- Visualizing migration routes in sub-Saharan Africa**

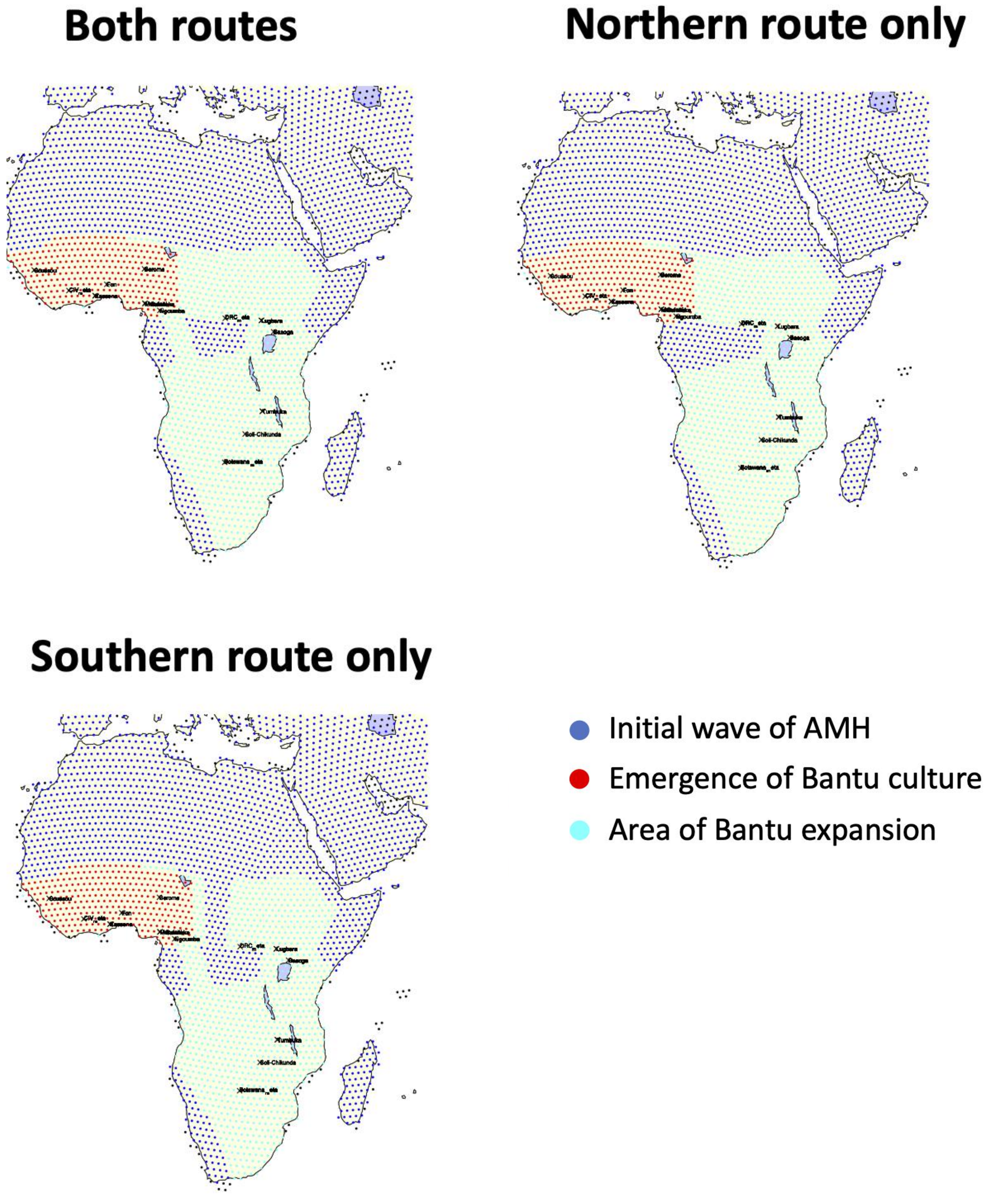


**Fig.S 10.1 | Spatially explicit framework**
Figure showing the three demographic scenarios considered in the spatially explicit framework. Each dot represents a local population, and the color indicates whether the deme is part of the initial wave of anatomically modern humans (AMH) alone (in blue), part of the area where Bantu culture emerged (in red) and subsequently spread (in cyan), shown against the continental contours and major water bodies for reference. Black dots and labels indicate the location and name of the populations with whole genome sequences included in the analysis.

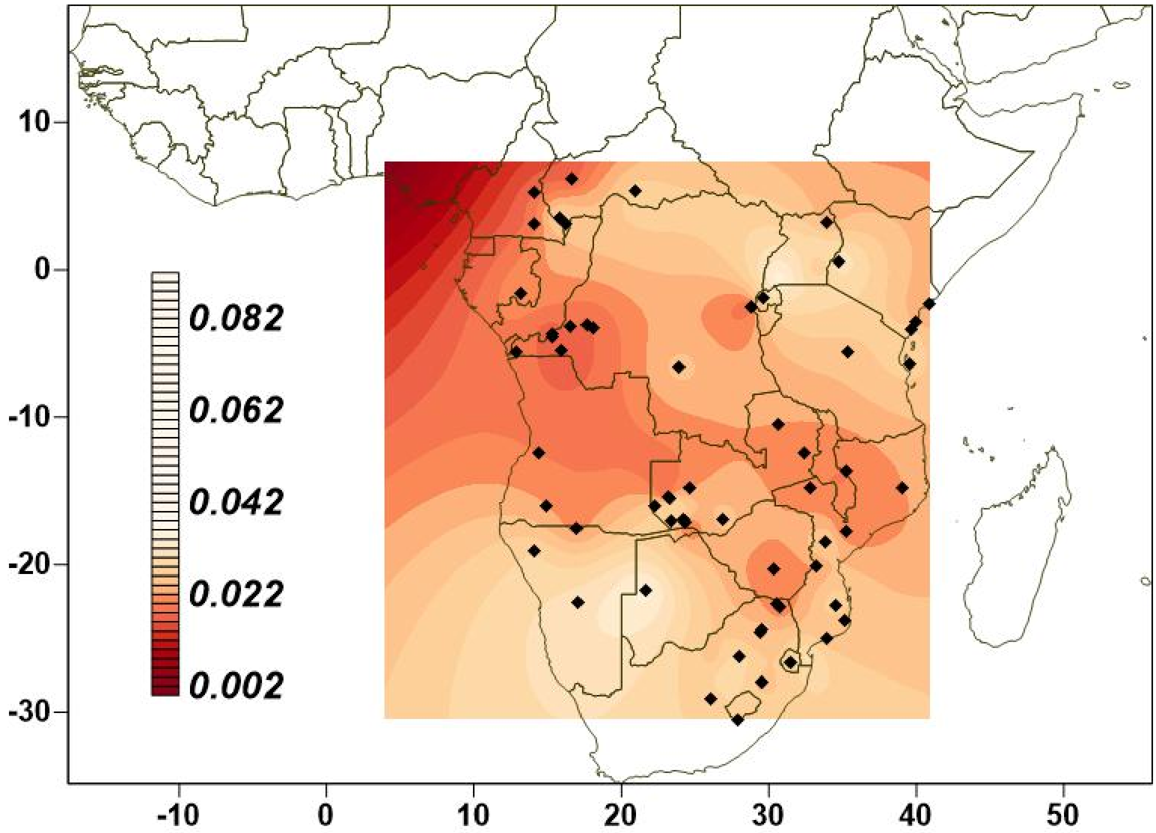


**Fig.S 10.2 | $F_{ST}$ matrix for the masked Only-BSP dataset.**
Figure showing $F_{ST}$ distances of one population in Cameroon and one BSP (black diamonds) from the masked and imputed Only-BSP dataset.

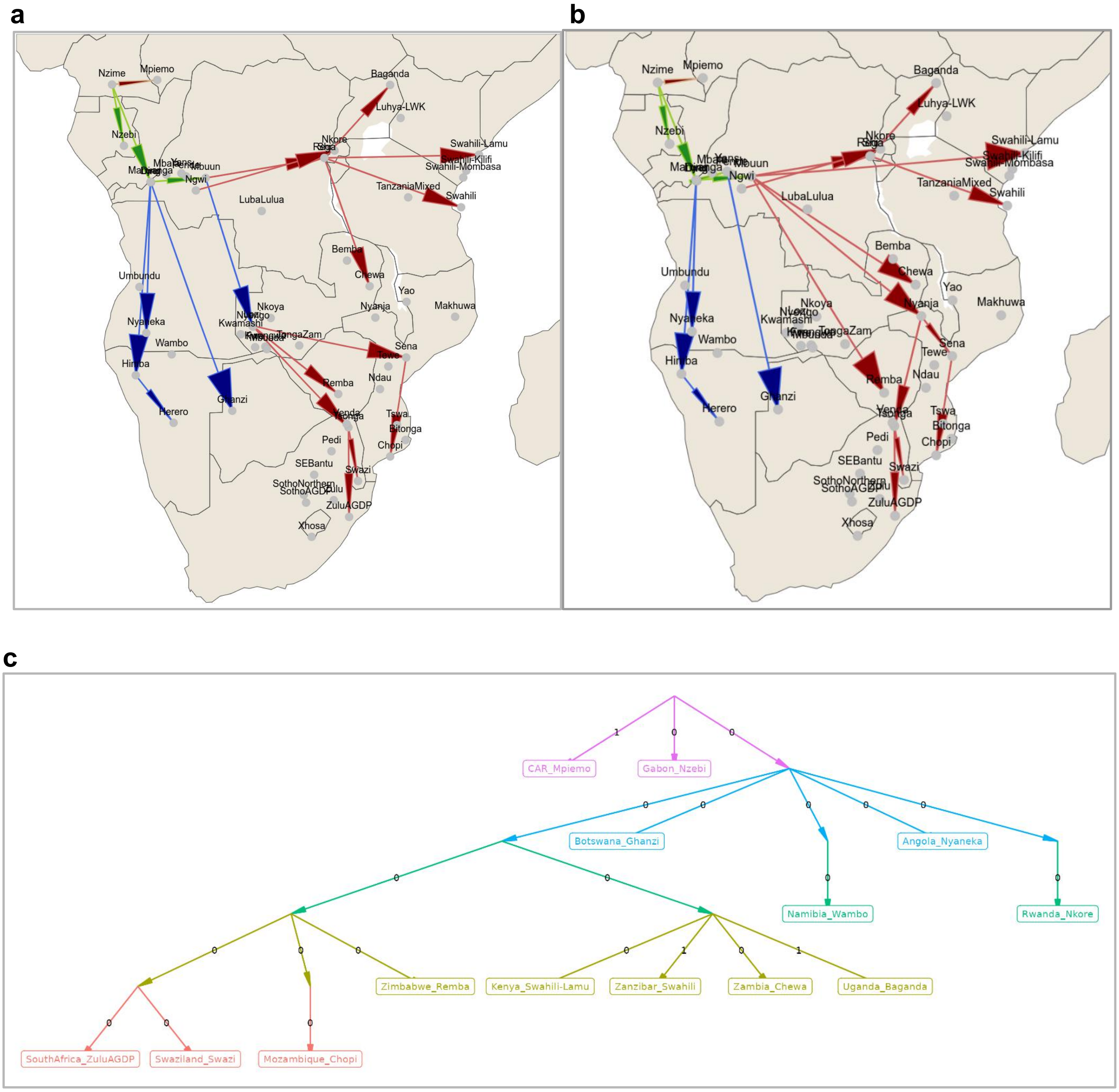


**Fig.S 10.3 | $F_{ST}$ map for the masked Only-BSP dataset.**

$F_{ST}$ values were estimated on the basis of the masked Only-BSP dataset. (**a**) Figure showing $F_{ST}$ map for all the BSP included in the AfricanNeo dataset (also **Fig. 5a**), and (**b**) for the BSP included in the AfricanNeo dataset except for the Lozi population from Zambia (also **Fig. 5b**). Arrow colors correspond to studied four Bantu-speaking groups: north-western Bantu (in brown), west-western Bantu (in green), south-western Bantu (in dark blue), and eastern Bantu speakers (in red). (**c**) Admixture graph for the masked Only-BSP dataset. Figure showing admixture graph created using qpGraph for the masked Only-BSP dataset including populations analyzed in **Fig.S 10.2a**.

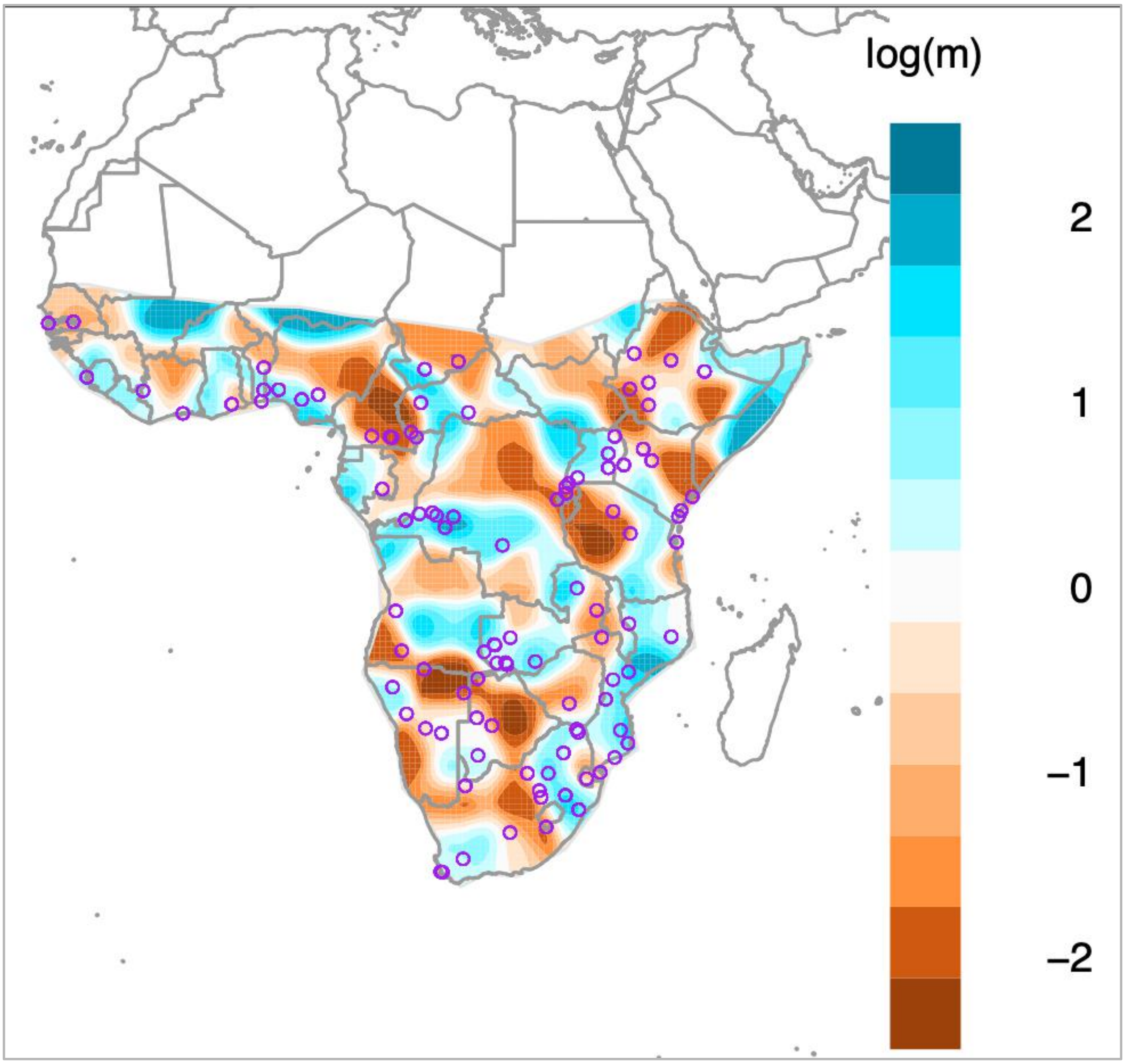


**Fig.S 10.4 | EEMS for the Only-African dataset.**
Figure showing EEMS results on the basis of unmasked Only-African dataset using 200 demes (also **Fig. 2c**). Blue areas are regions with high effective migration rates within the dataset, whilst brown areas indicate regions of inferred low effective migration rates (i.e. patterns of historic barriers to human migration).

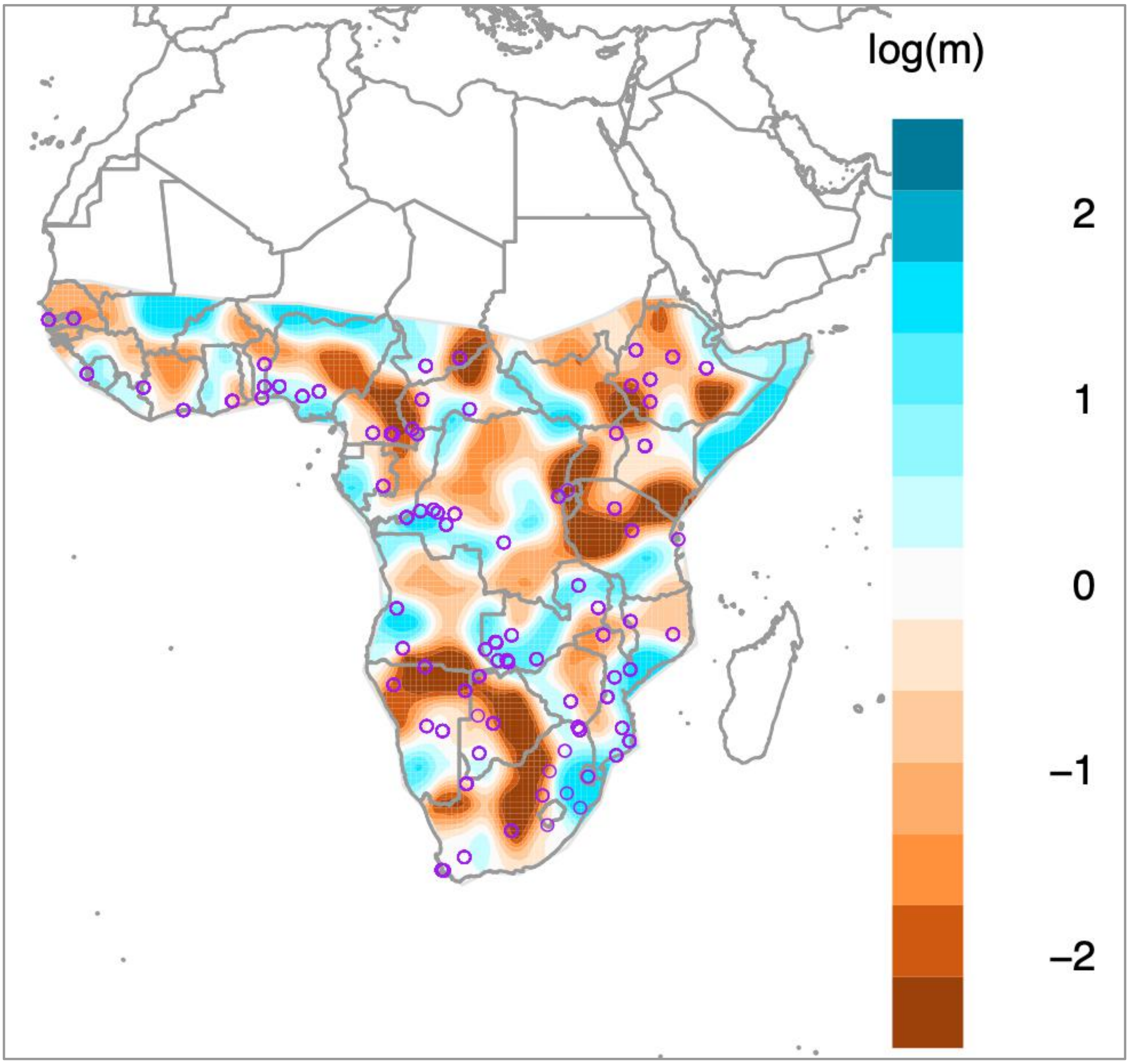


**Fig.S 10.5 | EEMS for the Only-African dataset after masking data of BSP.**
Blue areas indicate regions with high effective migration rates within the dataset whilst brown areas indicate regions of inferred low effective migration rates (i.e. patterns of historic barriers to human migration).

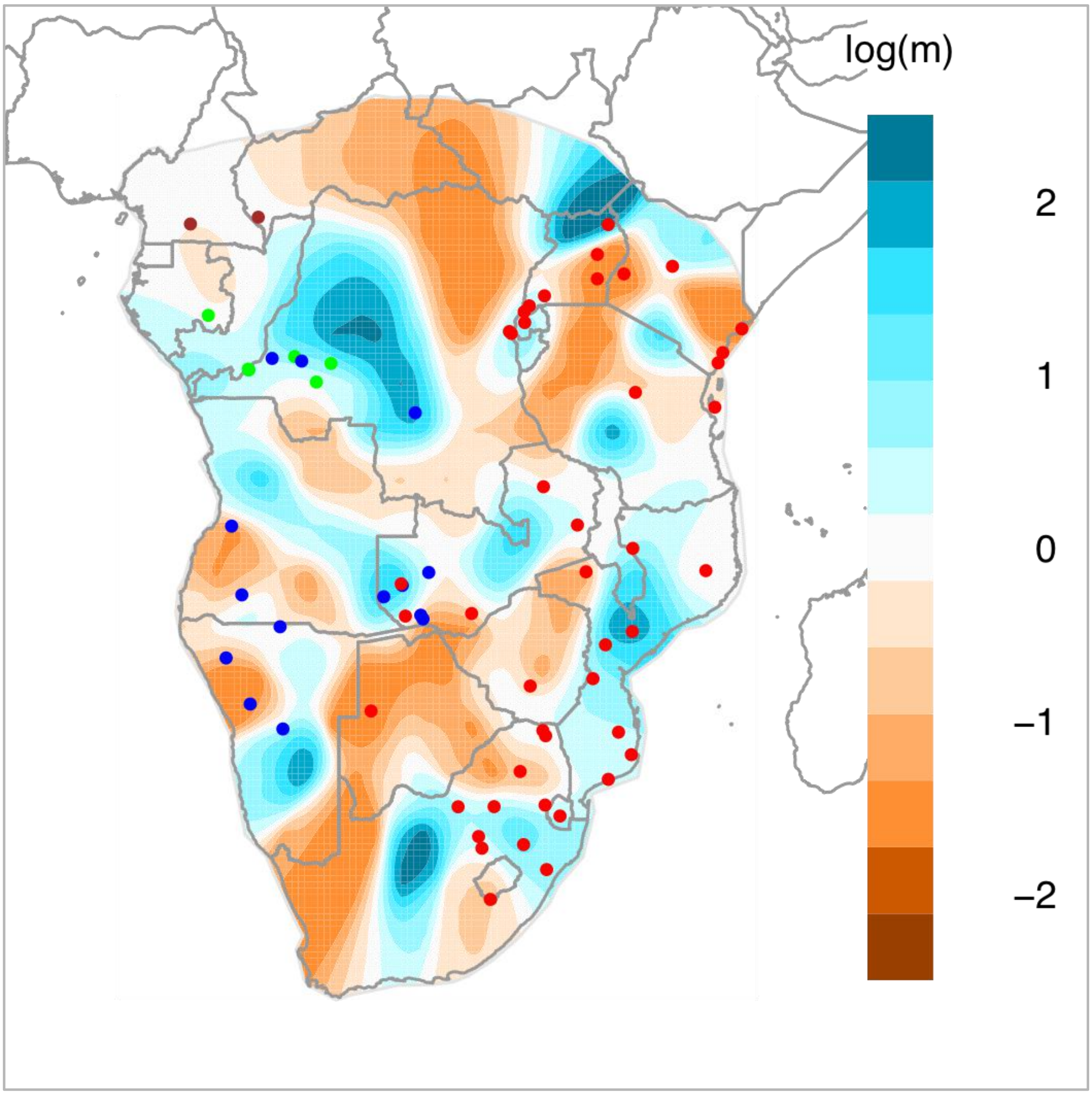


**Fig.S 10.6 | EEMS for the unmasked Only-BSP dataset.**
Figure showing EEMS results on the basis of unmasked Only-BSP dataset using 200 demes (also **Fig. 5c**). Purple circles indicate the sampling locations of each studied population. Blue areas are regions with high effective migration rates within the dataset whilst brown areas indicate regions of inferred low effective migration rates (i.e. patterns of historic barriers to human migration). Each dot war coloured according to the following classification: north-western Bantu 2 (in brown), west-western Bantu (in green), south-western Bantu (in dark blue), and eastern Bantu speakers (in red).

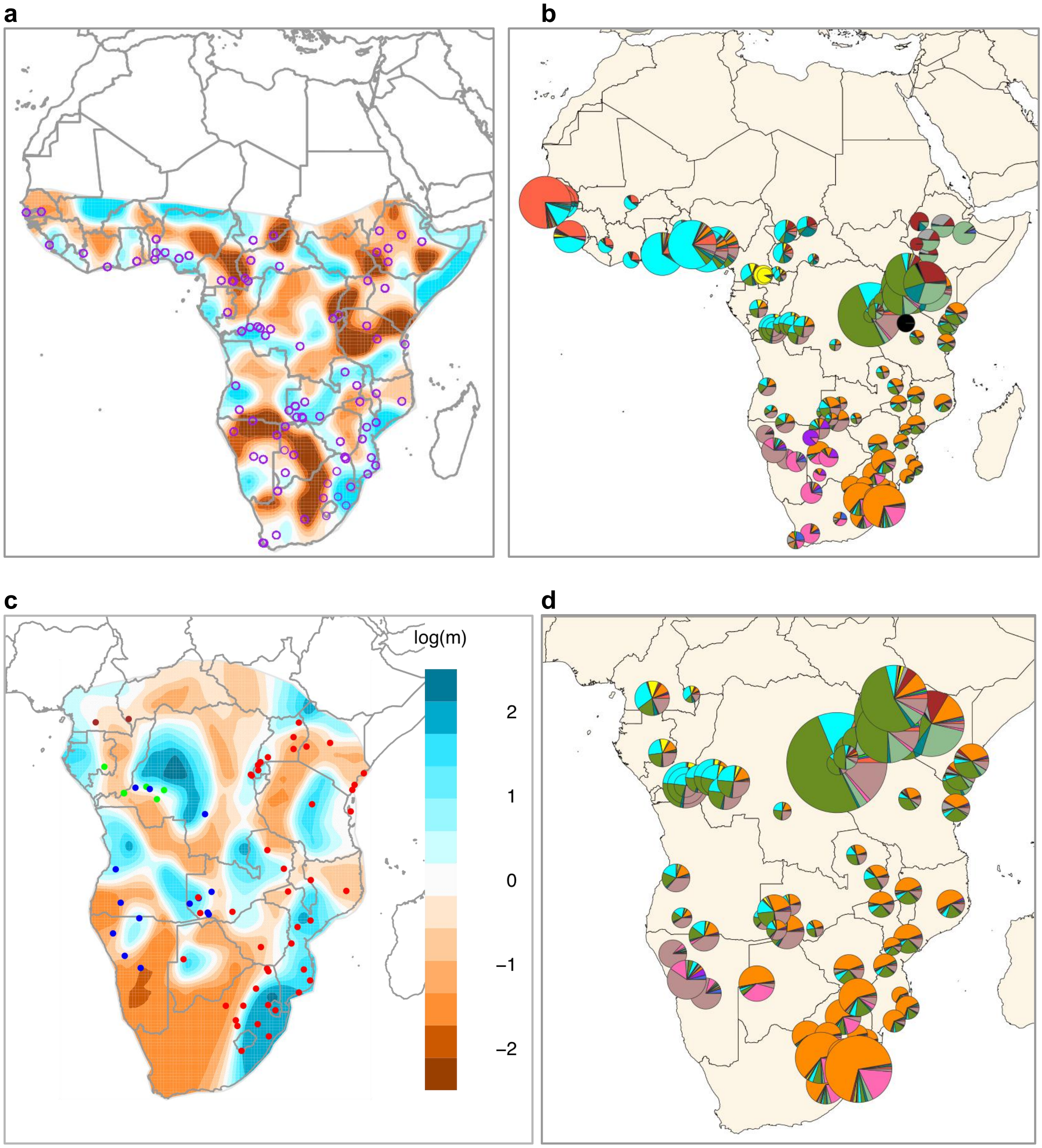


**Fig.S 10.7 | Comparisons between EEMS and ADMIXTURE results**
Figure comparing results obtained using EEMS and ADMIXTURE results for different unmasked datasets for selected sub-Saharan African populations: (**a**) EEMS results on the basis of the AfricanNeo dataset (also in **Fig.S 13.1**); (**b**) ADMIXTURE results at K=16 on the basis of the AfricanNeo dataset (also in **Fig.S 3.2**); (**c**) EEMS results on the basis of the Only-BSP dataset (also in **Fig.S 13.3**); and (**d**) ADMIXTURE results at K=16 on the basis of the Only-BSP dataset (also in **Fig.S 3.4**).

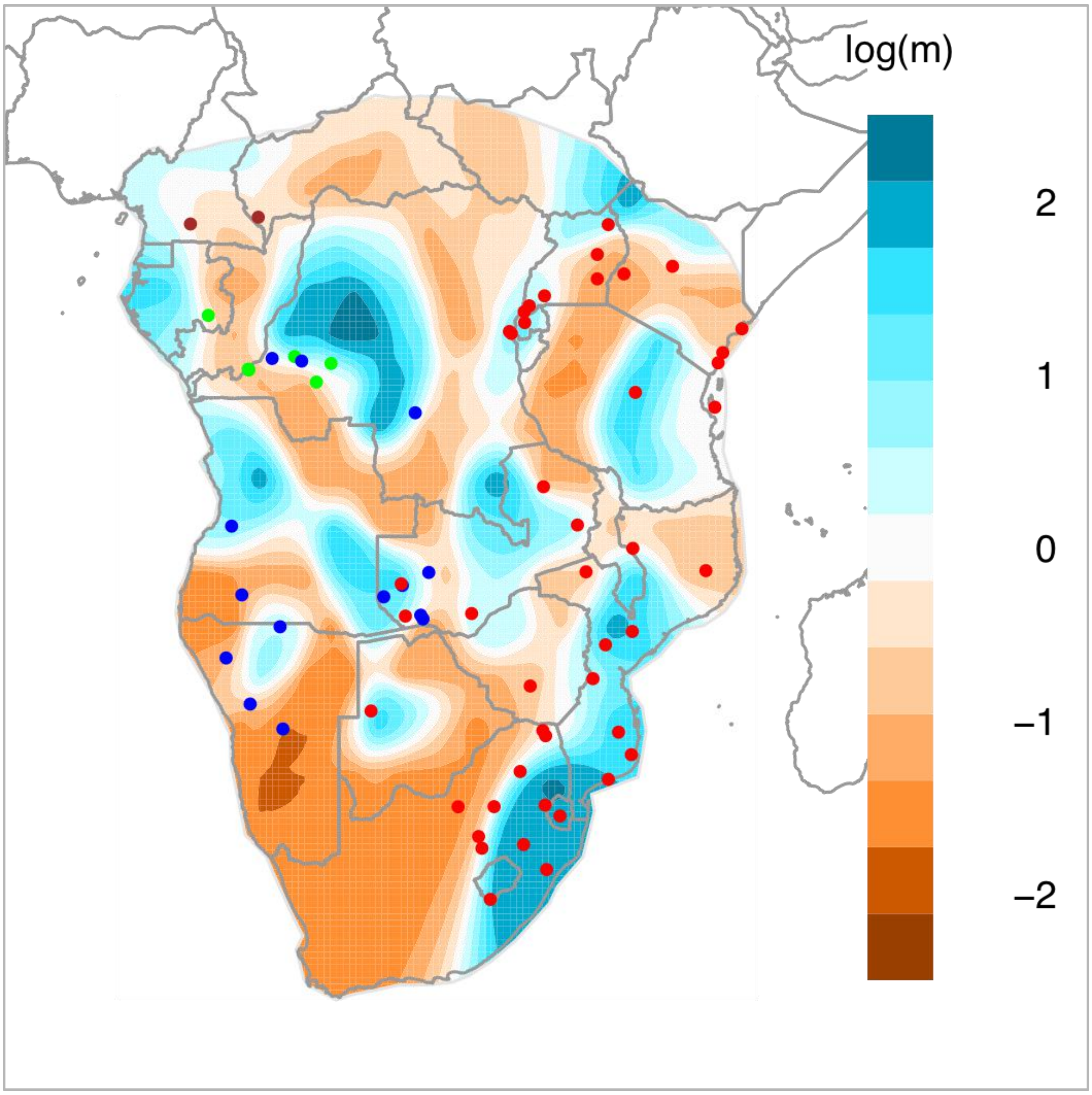


**Fig.S 10.8 | EEMS on the basis of the masked Only-Bantu dataset.**
Figure showing EEMS results on the basis of only masked and imputed data of BSP. Purple circles indicate the sampling locations of each studied population. Blue areas are regions with high effective migration rates within the dataset whilst brown areas indicate regions of inferred low effective migration rates (i.e. patterns of historic barriers to migration).

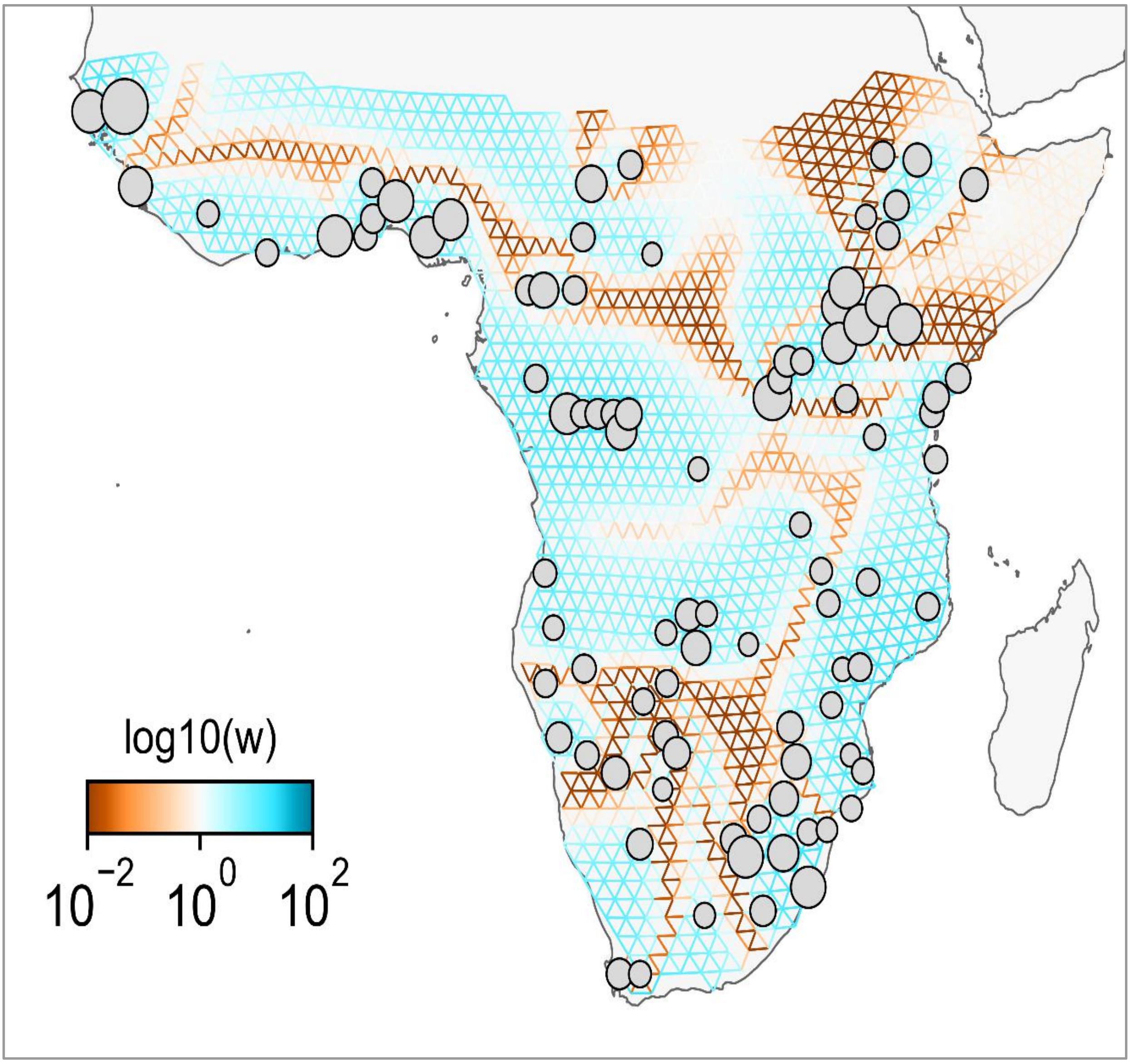


**Fig.S 10.9 | FEEMS of the unmasked AfricanNeo dataset.**
Figure showing FEEMS results on the basis of the full list of sub-Saharan African populations included in the AfricanNeo dataset. Each studied population is represented with one dot, and the size of the dots is in relation to the sample size of each population.

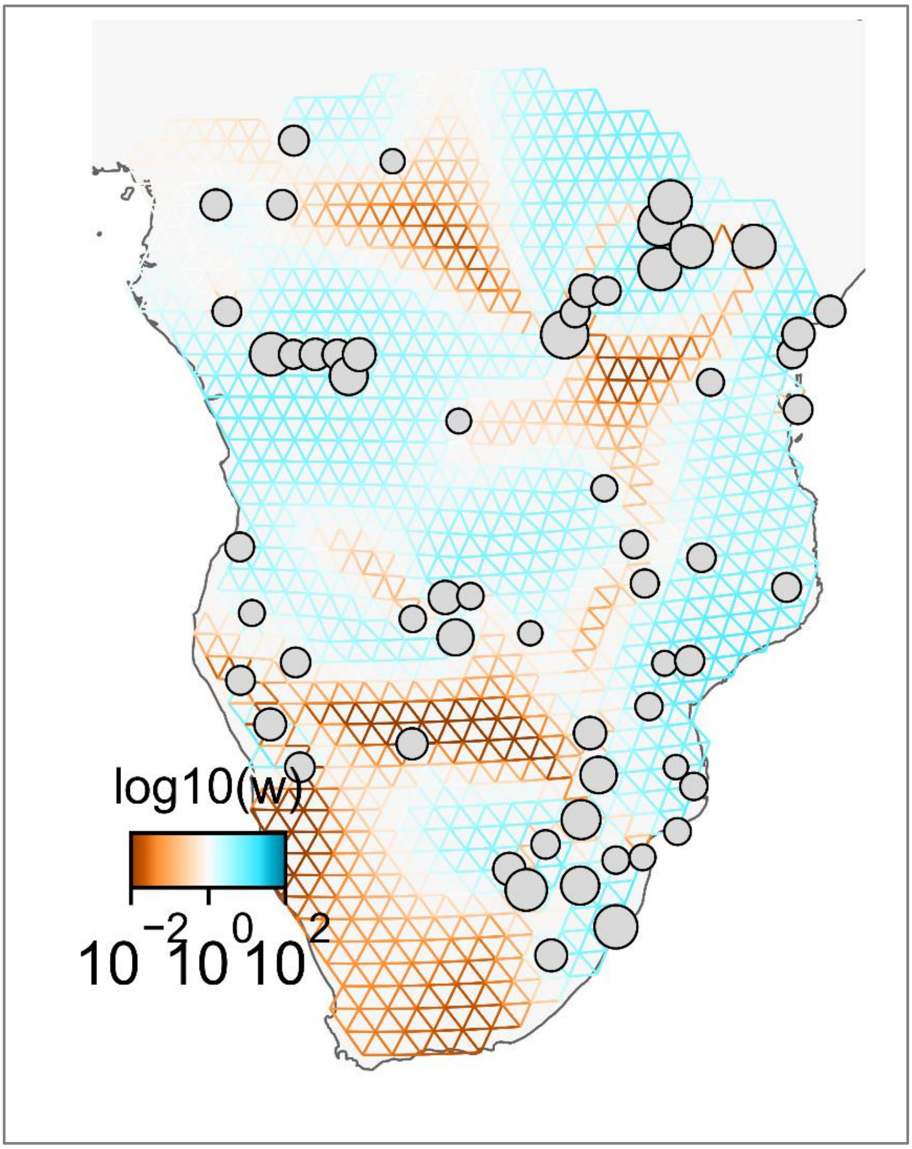


**Fig.S 10.10 | FEEMS on the basis of the unmasked Only-BSP dataset.**
Figure showing FEEMS results on the basis of the unmasked Only-BSP dataset. Each studied population is represented with one dot, and the size of the dots is in relation to the sample size of each population.

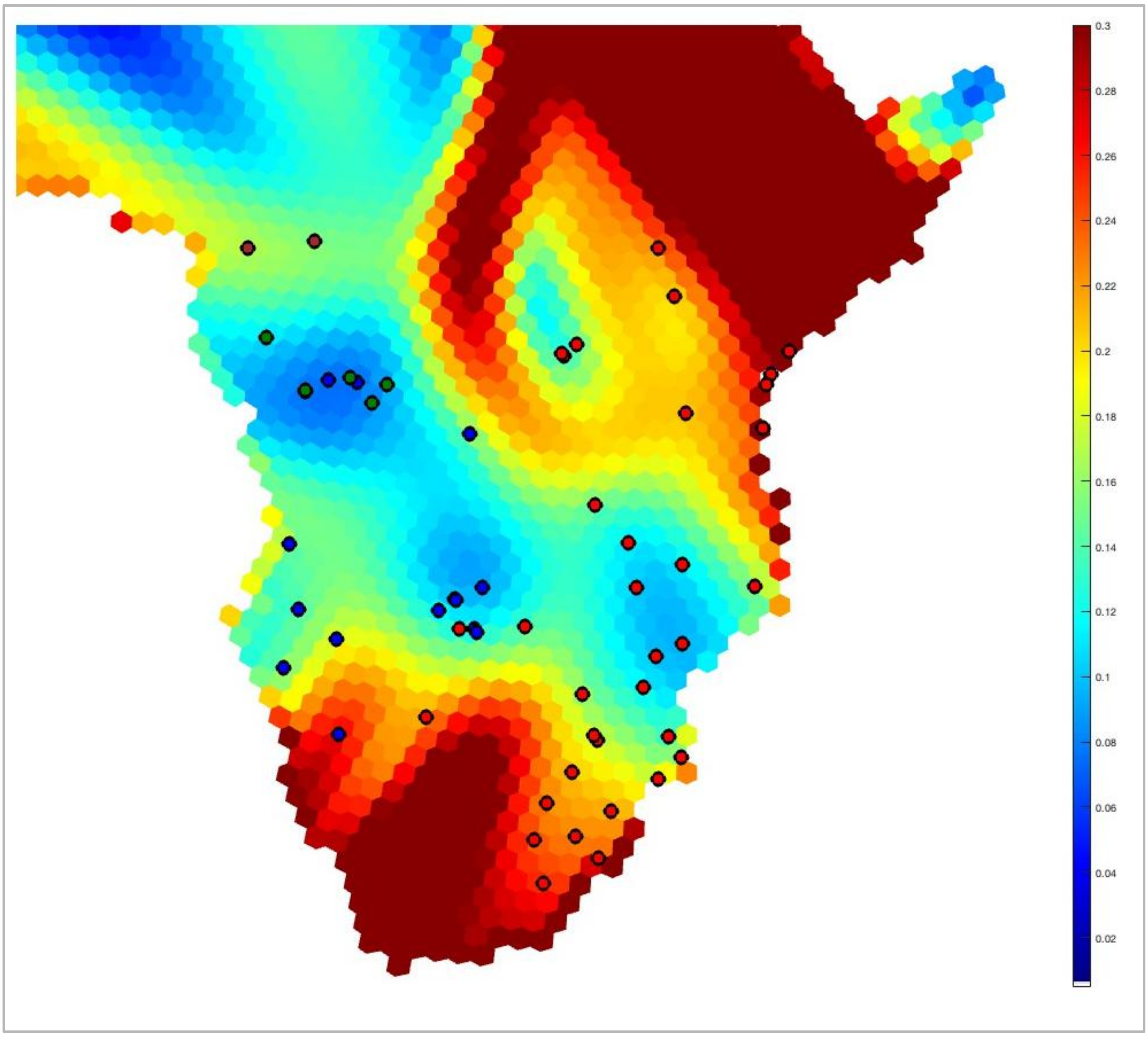


**Fig.S 10.11 | Spatial visualization of genetic barriers analysis on a grid.**
Figure showing barriers to migration using $F_{ST}$ as the distance for the masked and imputed BSP dataset. High values (in red) indicate sharp changes in alleles between populations, and thus are indicative of barriers to gene flow whilst low values (in blue) indicate the opposite. Hexagons of the grid were plotted with a color scale representing the $F_{ST}$ gradient.

#### 11- Estimated admixture dates in BSP

a

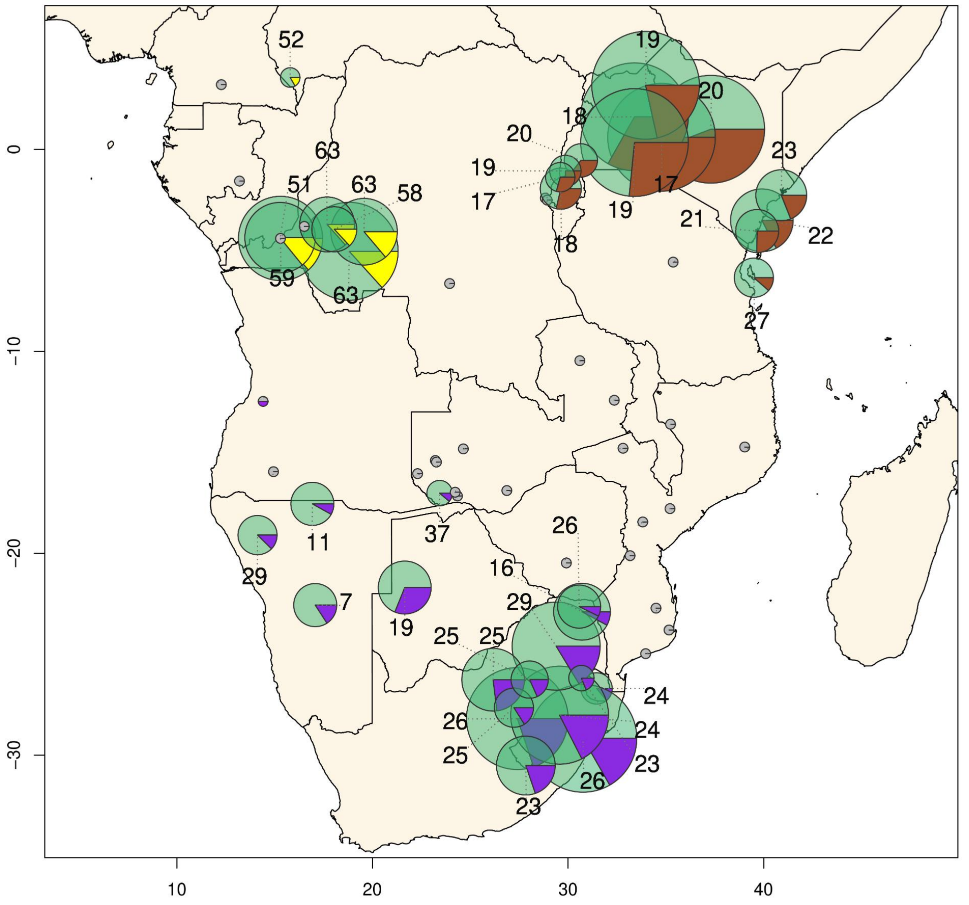

b

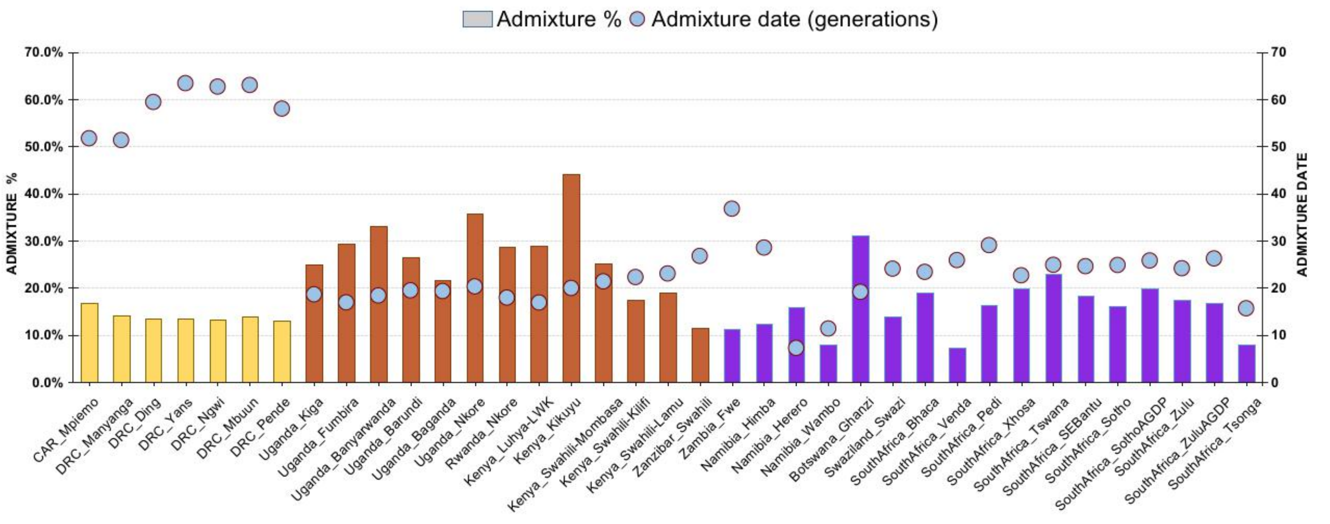


**Fig.S 11.1 | MOSAIC results for BSP with admixture.**
Figure showing MOSAIC results for all studied BSP computed using a two-way admixture model (also **Fig. 2b**). Figure showing (**a**) pie charts and locations of BSP with inferred admixture and (**b**) bar plots for the admixture proportions and dates (red squares). Each ancestry has different colors: west-central African-related ancestry in green; western rainforest hunter-gatherer ancestry in yellow; Afro-Asiatic-speaking ancestry in brown; and Khoe-San speaking ancestry in purple. The size of the charts is in relation to the sample size of each BSP.

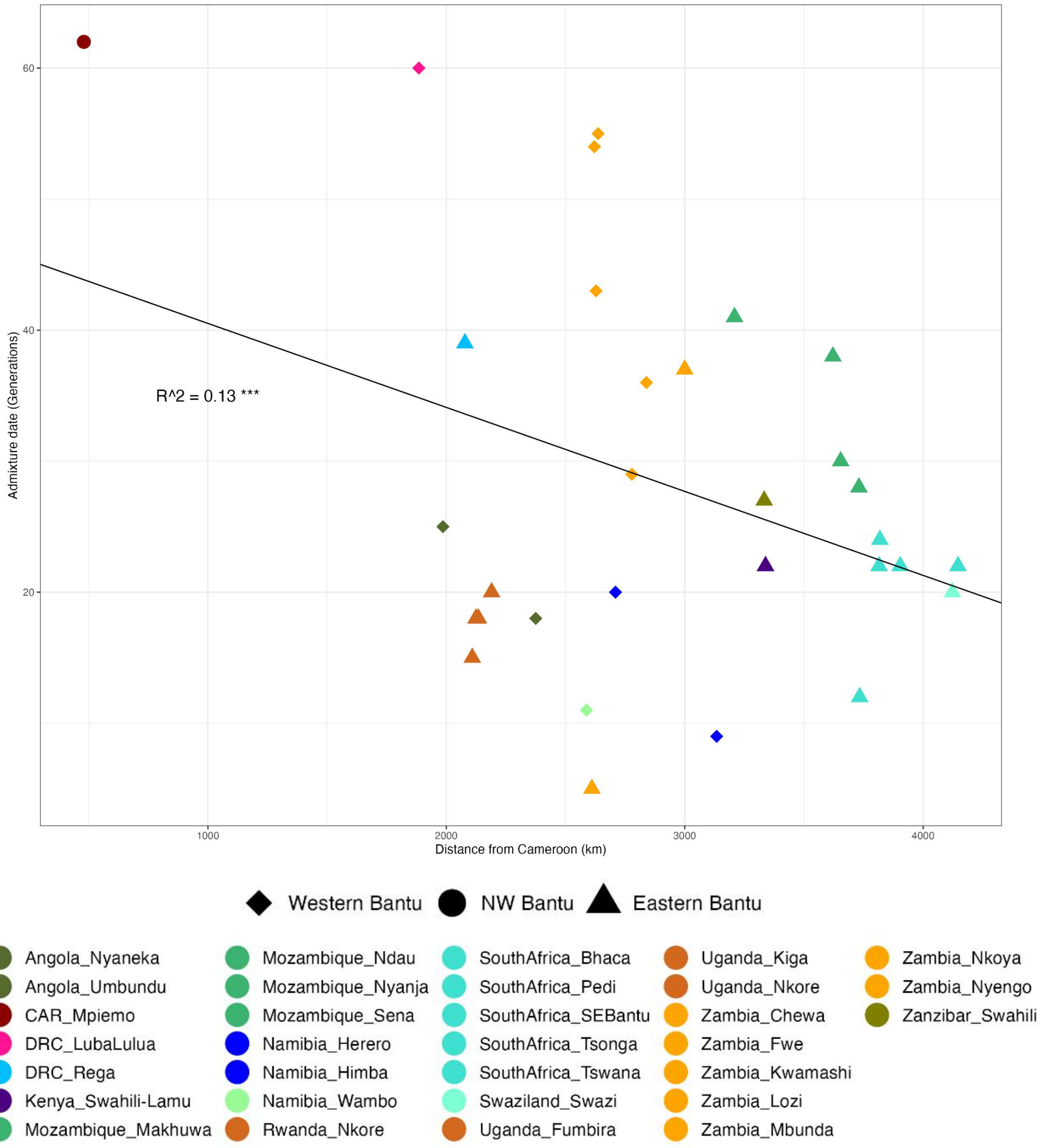


**Fig.S 11.2 | Admixture dates versus geographical distances from Cameroon.**
Figure showing admixture dates estimated using MOSAIC analyses for BSP from **Fig.S 11.1** plotted against geographical distances from Cameroon (also **Fig. 2C**). The markers have different shapes for each Bantu-speaking group: north-western (a circle), western (a diamond), and eastern (triangle) group.

#### 12- Comparisons between aDNA samples and modern-day African populations

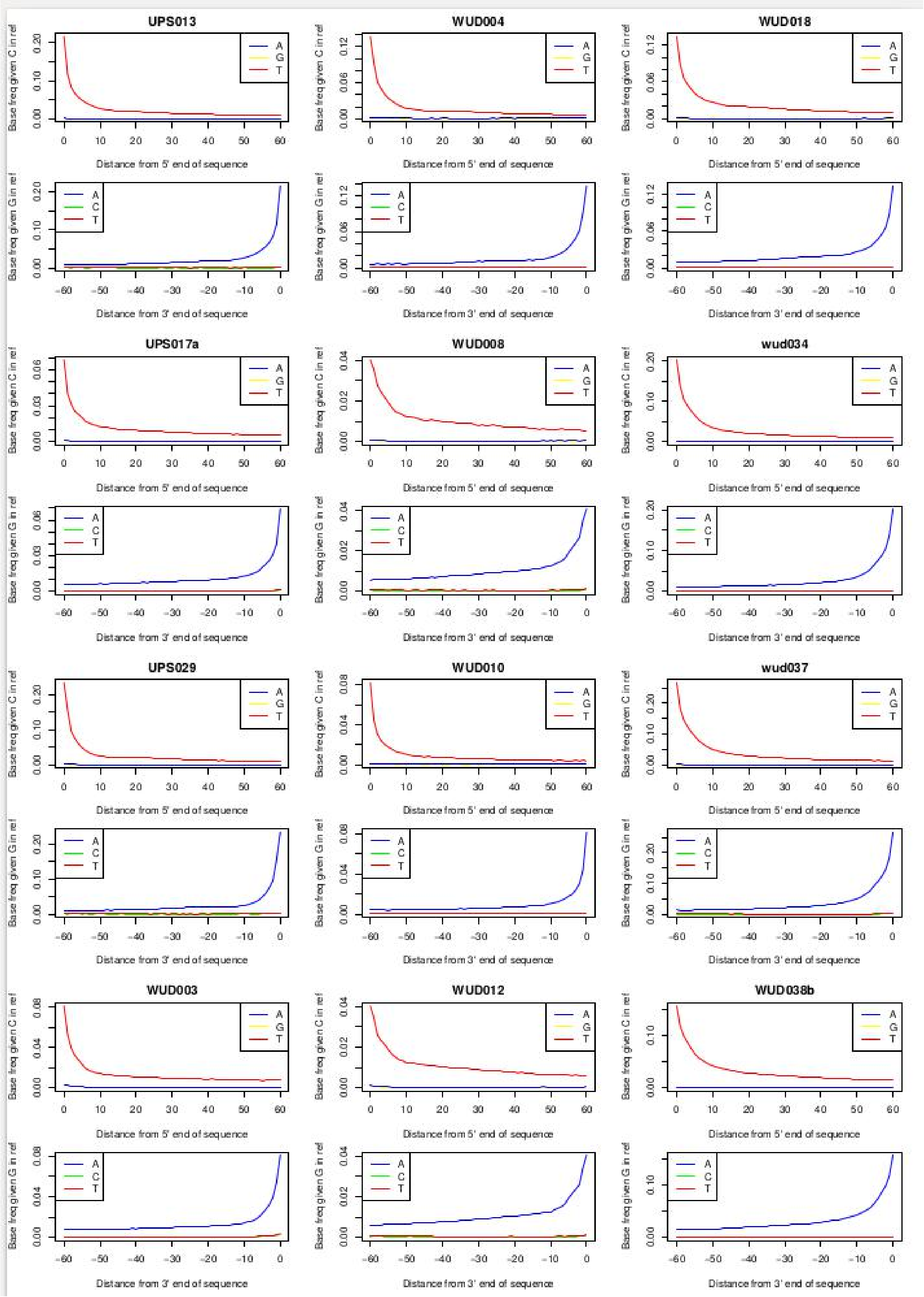


**Fig.S 12.0 | Misincorporation patterns for the twelve ancient samples newly sequenced.** The magnitude and the smoothness of the curves suggest non-damage in the aDNA sample.

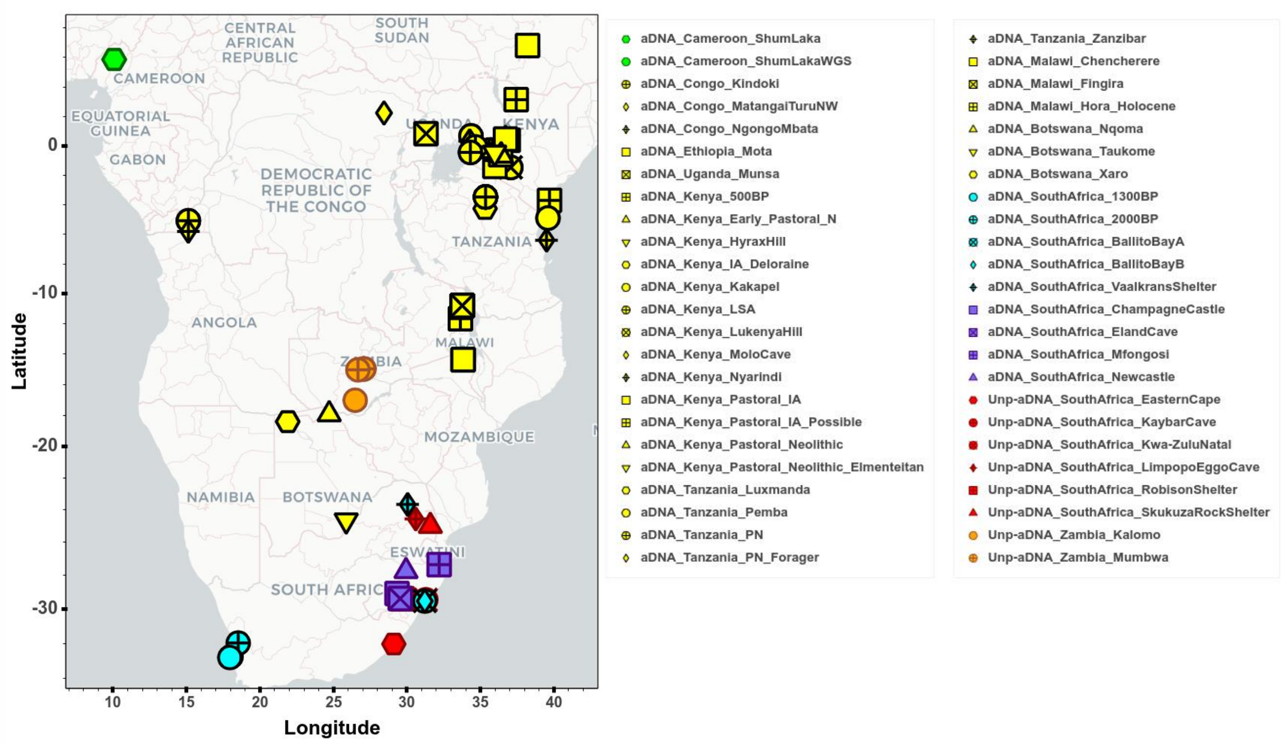


**Fig.S 12.1 | Geographical locations of aDNA individuals included in this study.**
Figure showing the locations of the 95 aDNA individuals analyzed in this study (**Table.S 5**). To better visualize the figure, we highlighted new aDNA individuals from Zambia in orange, new aDNA individuals from South Africa in red, aDNA individuals from previous studies in South Africa in cyan (Schlebusch et al. 2017) and purple (Skoglund et al. 2017), aDNA individuals from Shum Laka from Cameroon in green, and other aDNA individuals from previous studies in yellow. To better visualize the locations and ID of each aDNA individual we created an interactive plot (see **Fig.S_12.1_aDNA_Map.html**).

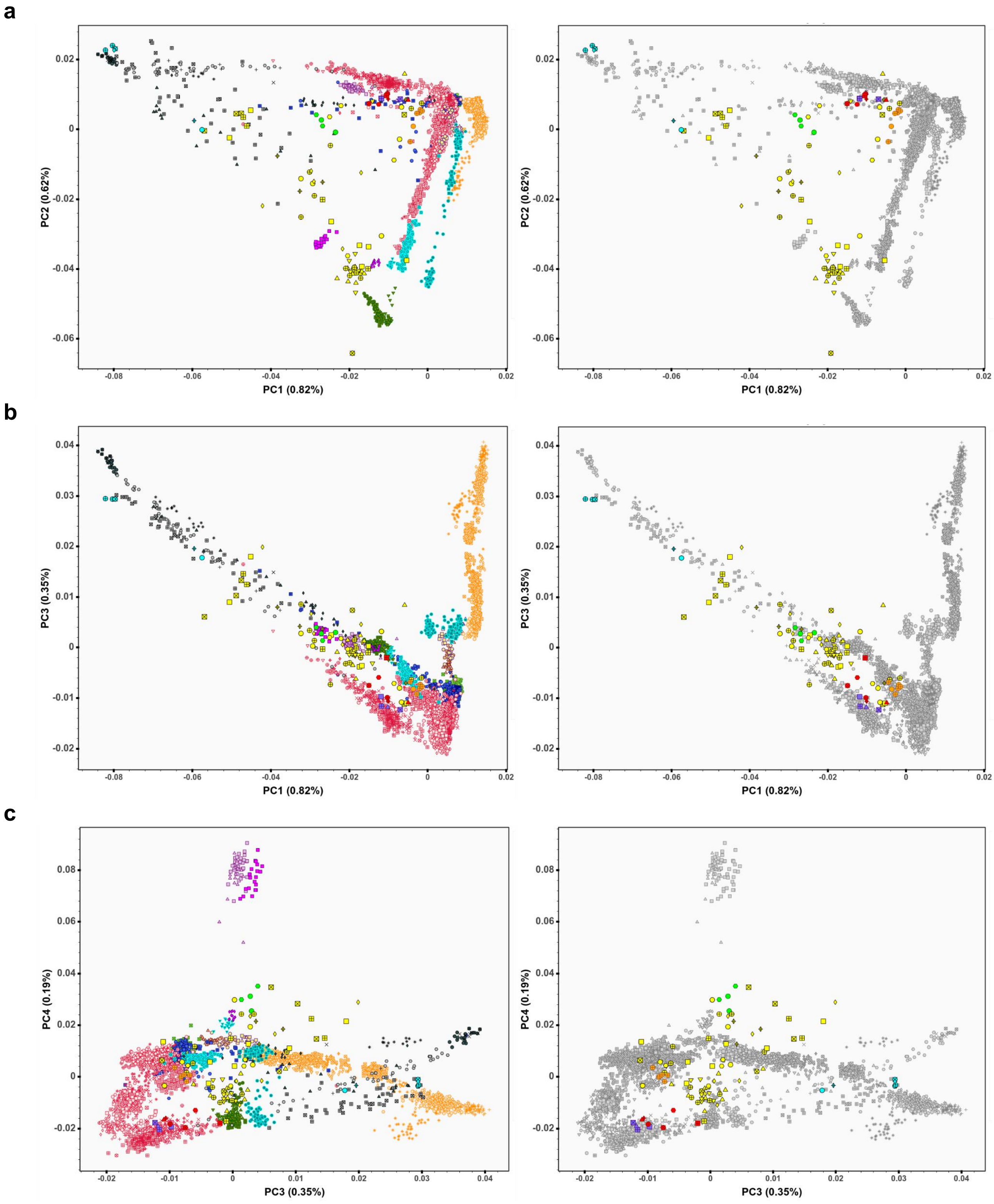


**Fig.S 12.2 | PCA of aDNA and modern African populations.**
Figure showing PCA plot of ancient DNA samples with a background of present-day populations included in the Only-African dataset. (**a**) PC1 vs PC2; (**b**) PC1 vs PC3; and (**c**) PC3 vs PC4. In the left column, groups of present-day populations were highlighted with colors (the same as in **Fig.S 1.5**), and in the right column, present-day samples were highlighted in gray. Ancient DNA samples were highlighted with the colors presented in **Fig.S 12.1** (see interactive plot **Fig.S_12.2*_PCA_aDNA_plot.html**).

Legend for the left column in **Fig.S 12.2**.

CAR_Mplemo
Cameroon_Nzime
Gabon_Nzebi
DRC_Manyanga
DRC_Ding
DRC_Lwer
DRC_Yans
DRC_Ngwi
DRC_Mbuun
DRC_Mbala
DRC_Pende
DRC_LubaLulua
Namibia_Himba
Namibia_Herero
Namibia_Wambo
Namibia_Damara-KSP
Zambia_Kwangwa
Zambia_Nyengo
Zambia_Kwamashi
Zambia_Mbunda
Zambia_Nkoya
Angola_Nyaneka
Angola_Umbundu
DRC_Shi
DRC_Rega
Uganda_Kiga
Uganda_Fumbira
Uganda_Banyarwanda
Uganda_Barundi
Uganda_Baganda
Uganda_Nkore

Rwanda_Nkore
Kenya_Luhya-LWK
Kenya_Kikuyu
Kenya_Swahili-Mombasa
Kenya_Swahili-Kilifi
Kenya_Swahili-Lamu
Tanzania_TanzaniaMixed
Zanzibar_Swahili
Zambia_Bemba
Zambia_Chewa
Zambia_Fwe
Zambia_Lozi
Zambia_TongaZam
Botswana_Ghanzi
Zimbabwe_Remba
Mozambique_Chopi
Mozambique_Bitonga
Mozambique_Tswa
Mozambique_Ndau
Mozambique_Tewe
Mozambique_Sena
Mozambique_Nyanja
Mozambique_Makhuwa
Mozambique_Yao
Swaziland_Swazi
SouthAfrica_Bhaca
SouthAfrica_Venda
SouthAfrica_Pedi
SouthAfrica_Xhosa
SouthAfrica_Tsonga
SouthAfrica_Tswana

SouthAfrica_SEBantu
SouthAfrica_Sotho
SouthAfrica_SothoAGDP
SouthAfrica_Zulu
SouthAfrica_ZuluAGDP
CAR_Banda
CAR_DzangaShangaPeople
CAR_Gbaya
Gambia_Fula
Gambia_Jola
Gambia_Mandinka
Gambia_Wolof
Gambia_Gambian-GWD
SierraLeone_Mende-MSL
Mali_Bwa
IvoryCoast_Ahizi
IvoryCoast_Yacouba
Ghana_GaAdangbe
Benin_Bariba
Benin_Fon
Benin_Yoruba
Nigeria_Igbo
Nigeria_Esan-ESN
Nigeria_Yoruba-YRI
Ethiopia_Wolayta
Ethiopia_Amhara
Ethiopia_Oromo
Ethiopia_Somali
Chad_Toubou
Chad_Sara
Ethiopia_Gumuz

Kenya_Kalenjin
Chad_Laal
Cameroon_Baka
CameroonGabon_Baka
Ethiopia_Sabue
Tanzania_Hadza
Angola_Khwe
Angola_Xun
Namibia_Juhoansi
Namibia_TsumkweKung
Namibia_Nama
Botswana_GuiGhanaKgal
Botswana_KalahariKhoe
SouthAfrica_Karretjie
SouthAfrica_Khomani
aDNA_Cameroon_ShumLaka
aDNA_Cameroon_ShumLakaWGS
aDNA_Congo_Kindoki
aDNA_Congo_MatangaiTuruNW
aDNA_Congo_NgongoMbata
aDNA_Ethiopia_Mota
aDNA_Uganda_Munsa
aDNA_Kenya_500BP
aDNA_Kenya_Early_Pastoral_N
aDNA_Kenya_HyraxHill
aDNA_Kenya_IA_Deloraine
aDNA_Kenya_Kakapel
aDNA_Kenya_LSA
aDNA_Kenya_LukenyaHill
aDNA_Kenya_MoloCave
aDNA_Kenya_Nyarindi

aDNA_Kenya_Pastoral_IA
aDNA_Kenya_Pastoral_IA_Possible
aDNA_Kenya_Pastoral_Neolithic
aDNA_Kenya_Pastoral_Neolithic_Elmenteitan
aDNA_Tanzania_Luxmanda
aDNA_Tanzania_Pemba
aDNA_Tanzania_PN
aDNA_Tanzania_PN_Forager
aDNA_Tanzania_Zanzibar
aDNA_Malawi_Chencherere
aDNA_Malawi_Fingira
aDNA_Malawi_Hora_Holocene
aDNA_Botswana_Nqoma
aDNA_Botswana_Taukome
aDNA_Botswana_Xaro
aDNA_SouthAfrica_1300BP
aDNA_SouthAfrica_2000BP
aDNA_SouthAfrica_BallitoBayA
aDNA_SouthAfrica_BallitoBayB
aDNA_SouthAfrica_VaalkransShelter
aDNA_SouthAfrica_ChampagneCastle
aDNA_SouthAfrica_ElandCave
aDNA_SouthAfrica_Mfongosi
aDNA_SouthAfrica_Newcastle
Unp-aDNA_SouthAfrica_EasternCape
Unp-aDNA_SouthAfrica_KaybarCave
Unp-aDNA_SouthAfrica_Kwa-ZuluNatal
Unp-aDNA_SouthAfrica_LimpopoEggoCave
Unp-aDNA_SouthAfrica_RobisonShelter
Unp-aDNA_SouthAfrica_SkukuzaRockShelter
Unp-aDNA_Zambia_Kalomo
Unp-aDNA_Zambia_Mumbwa

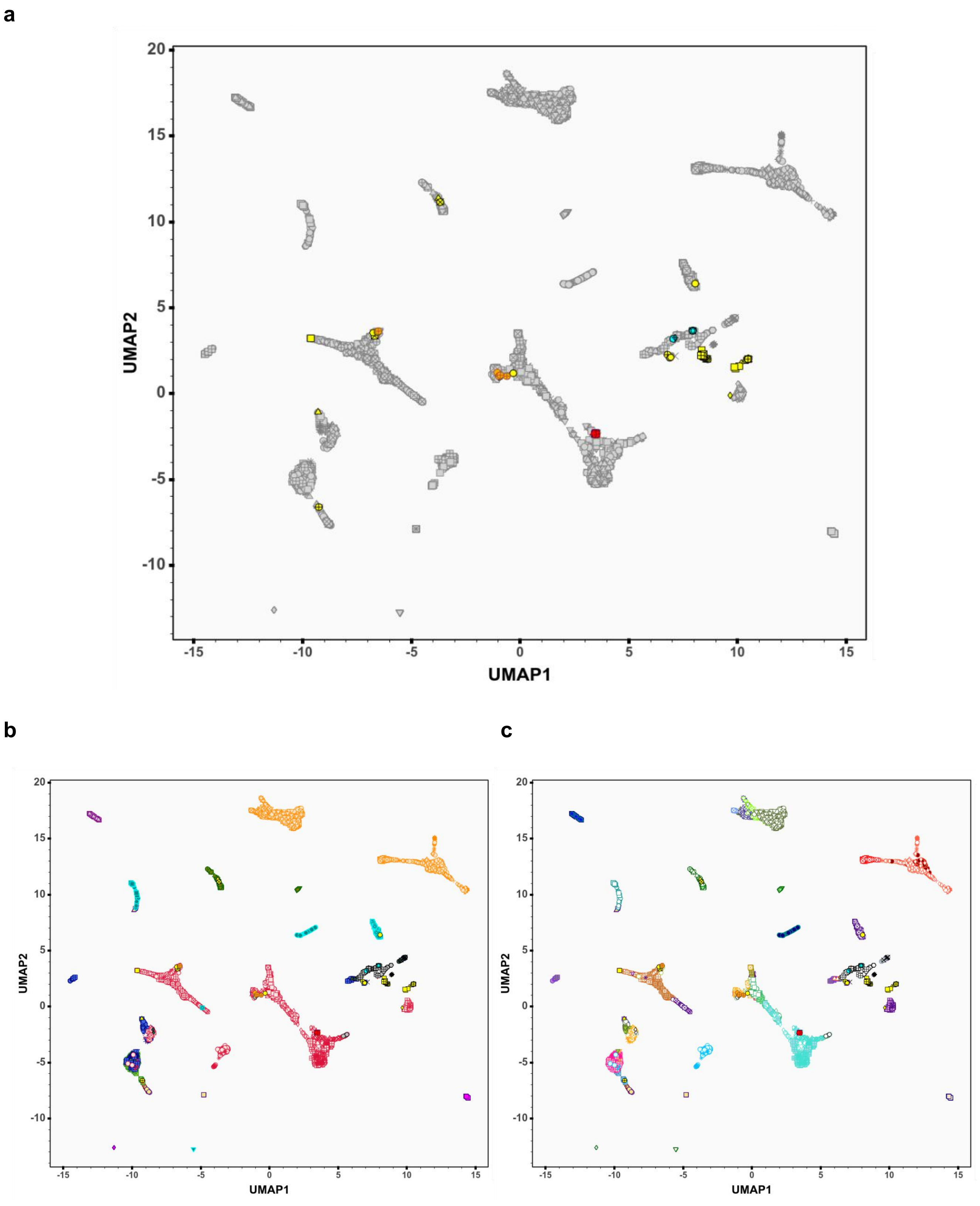


**Fig.S 12.3 | PCA-UMAP of aDNA individuals and present-day African populations.**
PCA-UMAP plot of ancient DNA samples with a background of sub-Saharan African populations included in the AfricanNeo dataset (**a**) in gray on the top, and on the bottom with colors (**b**) for each group and (**c**) for each population. Legends are the same as in **Fig.S 12.2** (see interactive plot **Fig.S_12.3*_PCA_aDNA_plot.html**).

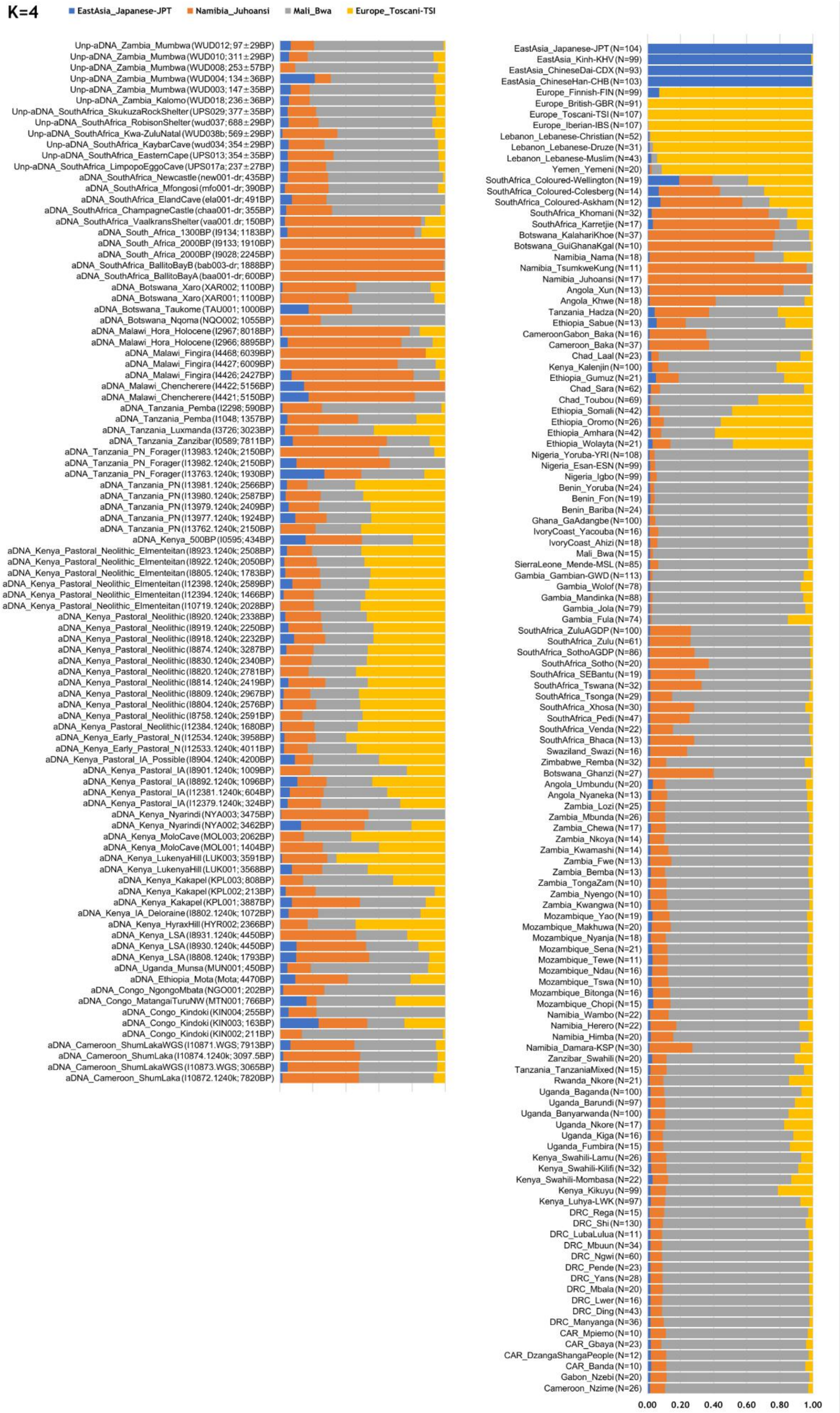


**Fig.S 12.4 | ADMIXTURE results at K=4 of ancient and modern African populations.**
Figure showing ADMIXTURE results at K=4 of ancient and modern African and Eurasian populations. For a better comparison, figure showing the averages for each aDNA individual and for each modern population (sample sizes in parenthesis).

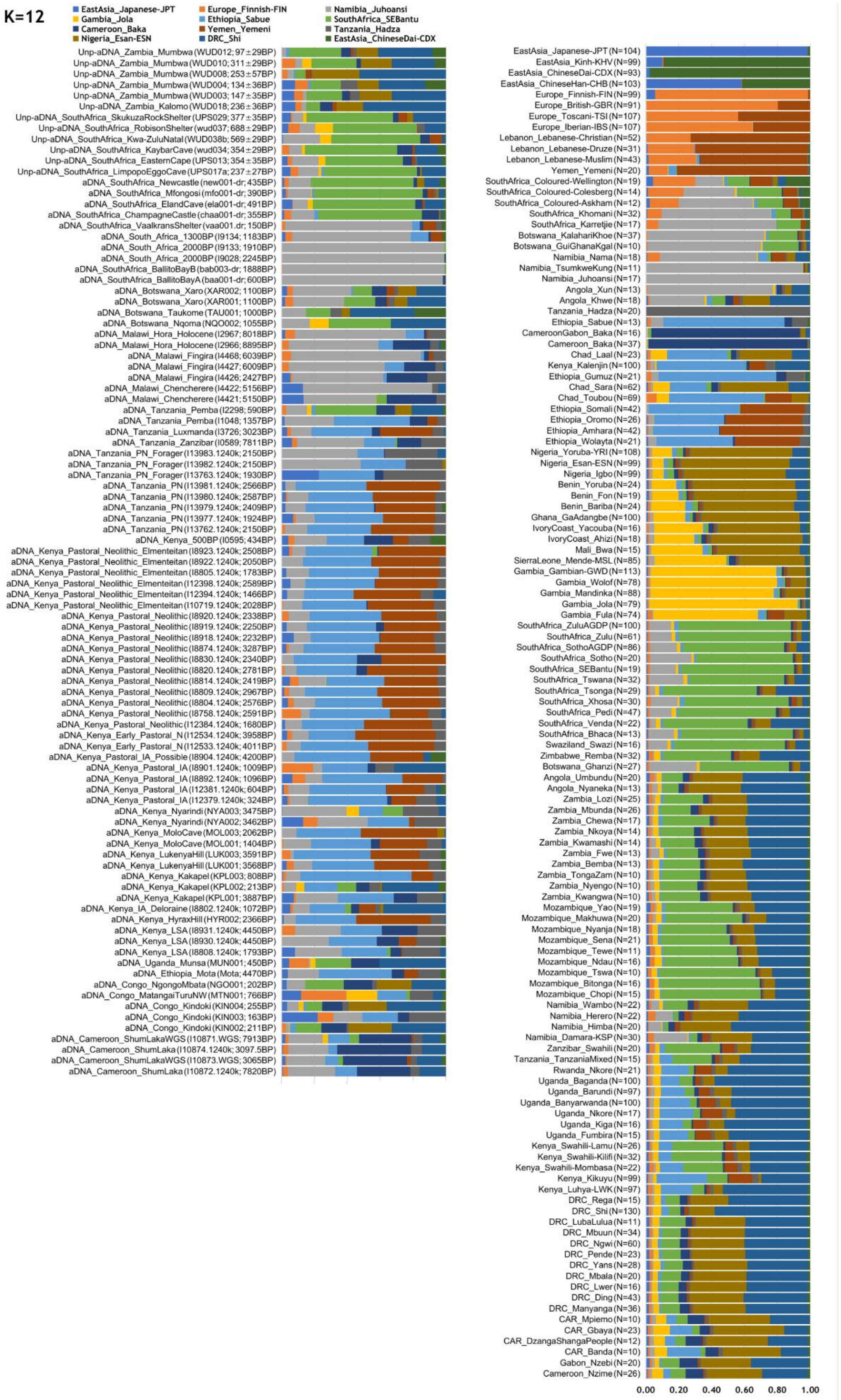


**Fig.S 12.5 | ADMIXTURE results at K=12 of ancient and modern African populations.**
Figure showing ADMIXTURE results at K=12 of ancient and modern African and Eurasian populations. For a better comparison, figure showing the averages for each aDNA individual and for modern populations (sample sizes in parenthesis).

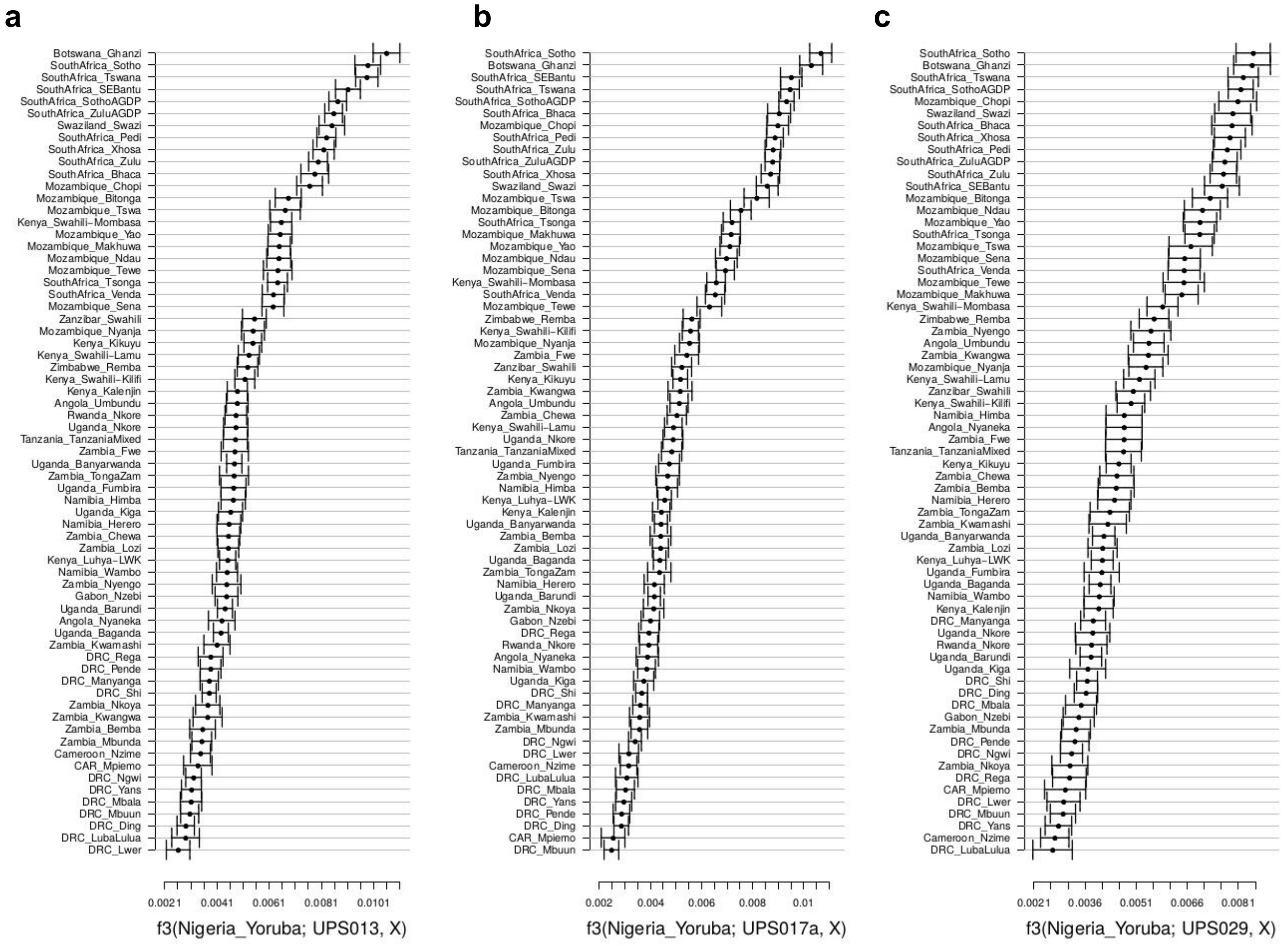


**Fig.S 12.6 | Genetic affinity of ancient samples UPS013, UPS017a, UPS029 to modern BSP estimated using the f3-statistics.**
Positive values of f3-statistics in the form f3(Yoruba; ancient-sample, BSP) indicate increasing genetic affinity between modern BSP and three ancient samples: (**a**) UPS013, (**b**) UPS017a, and (**c**) UPS029. Bars indicate the standard error of the mean. Values of the statistic significantly higher than zero (*P*-value < 0.05) are indicated with black color. In each plot, BSP indicated in the Y-axis are ordered according to decreasing values of the f3-statistics. All three samples originate from current-day South Africa. For the three ancient samples (UPS013, UPS017a, and UPS029) the radio-carbon calibrated ages are 1465-1648 CE, 1649-1806 CE, and 1461-1634 CE, respectively, and the genome coverage percentages are 0.484, 0.942, and 0.367, respectively (**Table.S 11**).

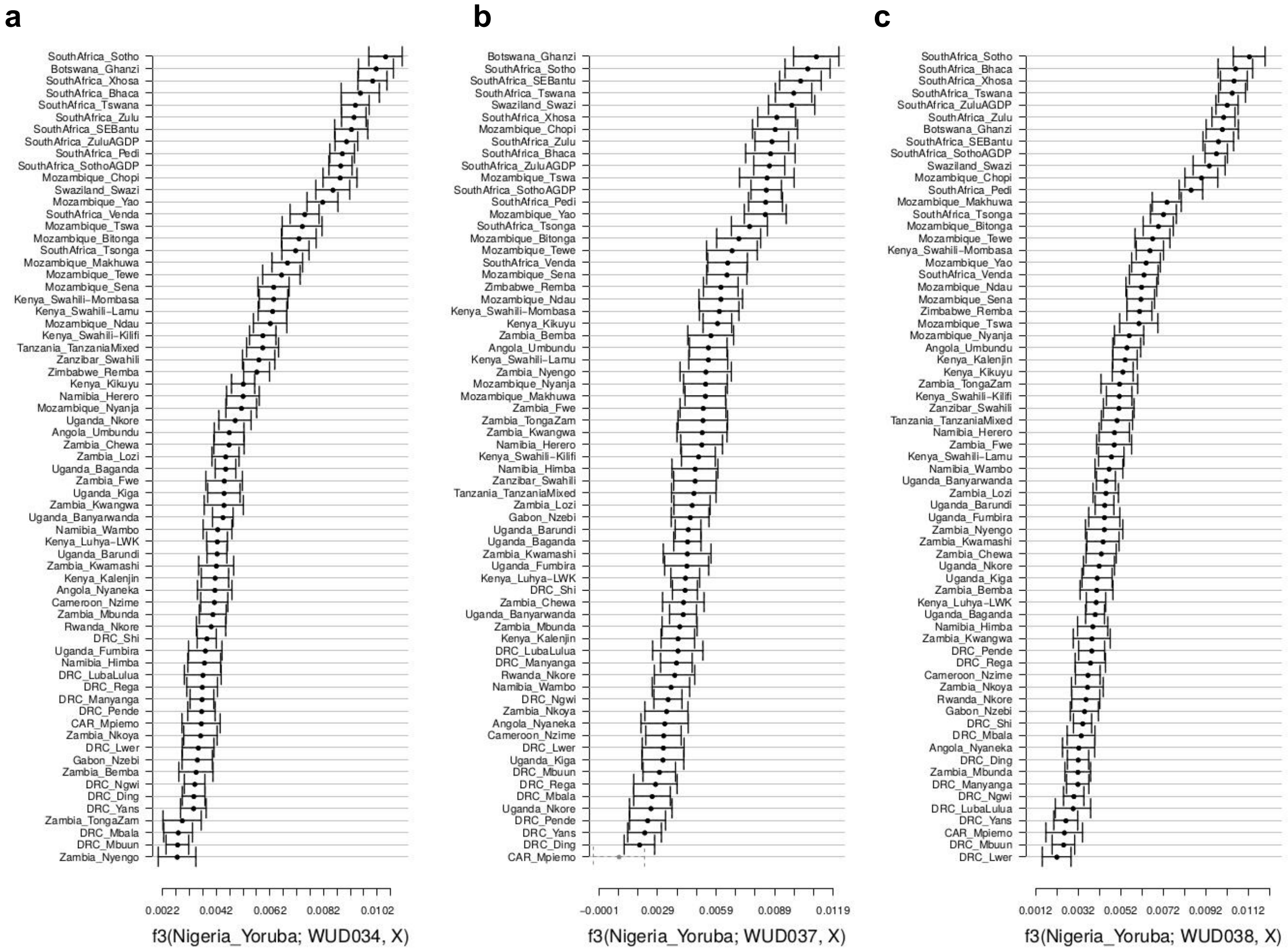


**Fig.S 12.7 | Genetic affinity of ancient samples WUD034, WUD037, WUD038b to modern BSP estimated using the f3-statistics.**

Positive values of f3-statistics in the form f3(Yoruba; ancient-sample, BSP) indicate increasing genetic affinity between modern BSP and three ancient samples: (**a**) WUD034, (**b**) WUD037, and (**c**) WUD038b. Bars indicate the standard error of the mean. Values of the statistic significantly higher than zero (*P*-value < 0.05) are indicated with black color. In each plot, BSP indicated in the Y-axis are ordered according to decreasing values of the f3-statistics. All three samples originate from current-day South Africa. For the three ancient samples (WUD034, WUD037, and WUD038b) the radio-carbon calibrated ages are 1487–1646 CE, 1289-1394 CE, and 1327-1445 CE, respectively, and the genome coverage percentages are 0.220, 0.055, and 0.144, respectively (**Table.S 11**).

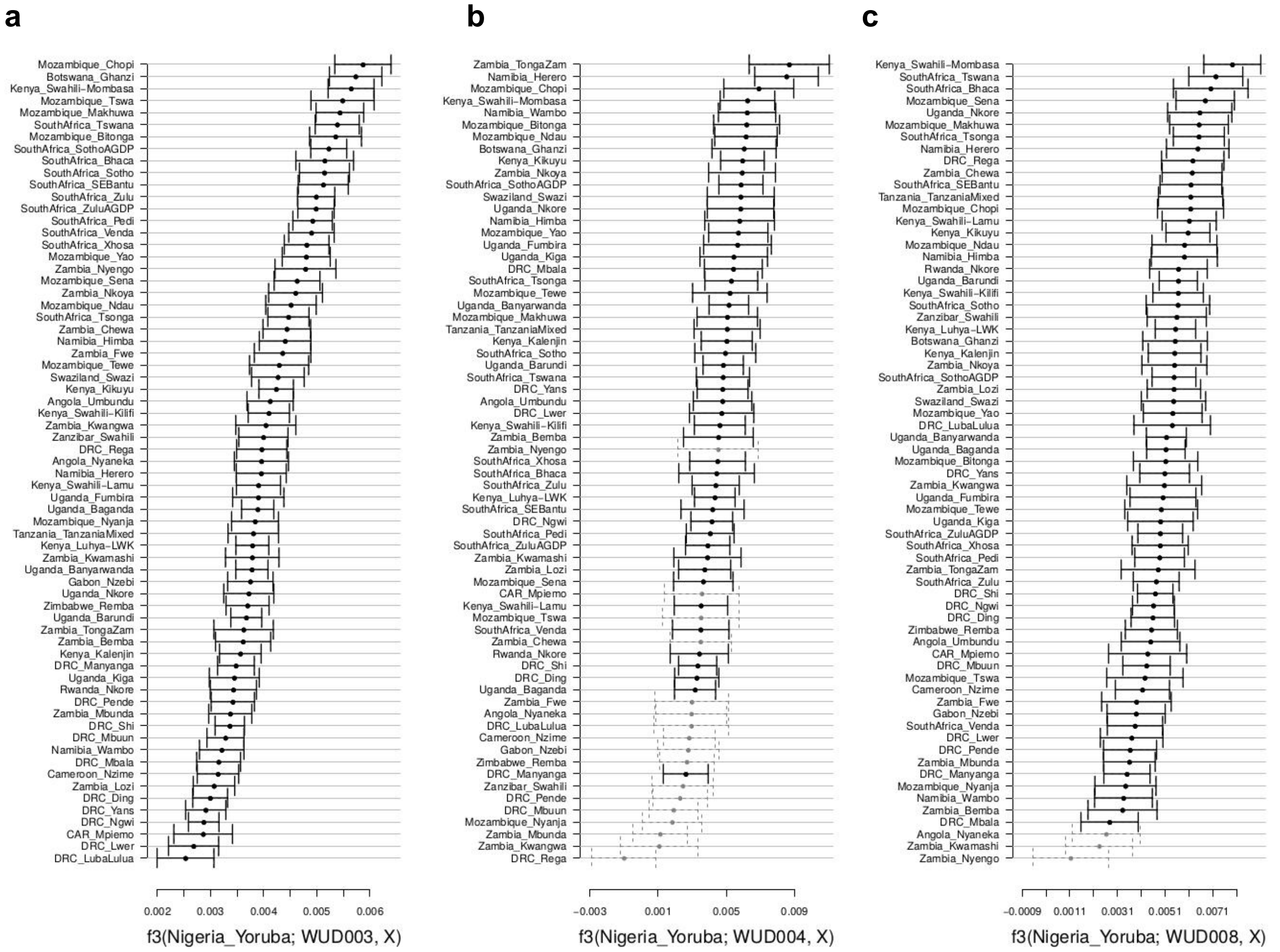


**Fig.S 12.8 | Genetic affinity of ancient samples WUD003, WUD004, WUD008 to modern BSP estimated using the f3-statistics.**

Positive values of f3-statistics in the form f3(Yoruba; ancient-sample, BSP) indicate increasing genetic affinity between modern BSP and three ancient samples: (**a**) WUD003, (**b**) WUD004, and (**c**) WUD008. Bars indicate the standard error of the mean. Values of the statistic significantly higher than zero (*P*-value < 0.05) are indicated with black color. In each plot, BPS indicated in the Y-axis are ordered according to decreasing values of the f3-statistics. All three samples originate from current-day Zambia. For the three ancient samples (WUD003, WUD004, and WUD008) radio-carbon calibrated age is 1673–1913 CE, 1675–1916 CE and 1506–1880 CE, respectively, and genome coverage percentage is 0.469, 0.020, and 0.036, respectively (**Table.S 11**).

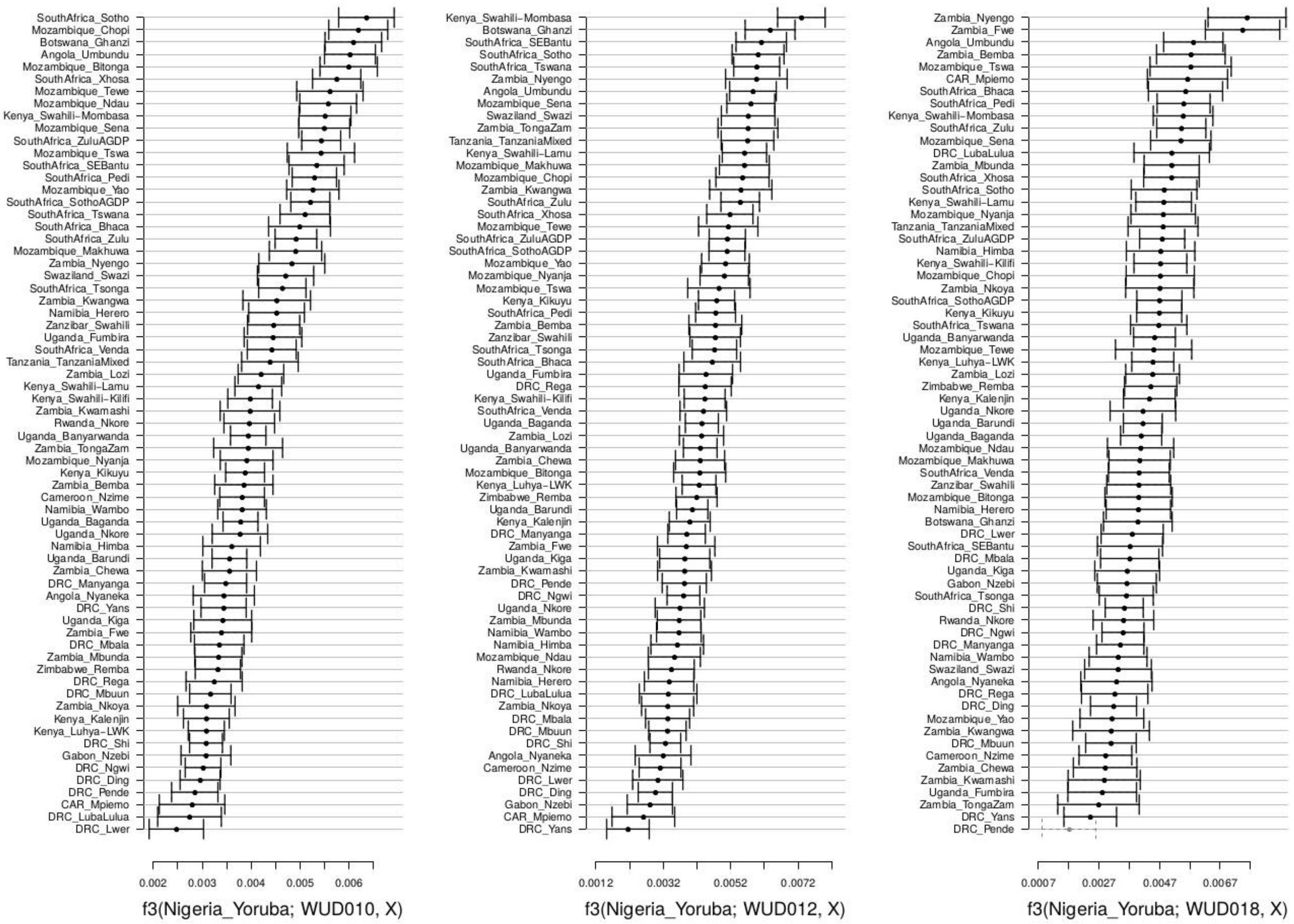


**Fig.S 12.9 | Genetic affinity of ancient samples WUD010, WUD012, WUD018 to modern BSP estimated using the f3-statistics.**

Positive values of f3-statistics in the form f3(Yoruba; ancient-sample, BSP) indicate increasing genetic affinity between the ancient sample to each of the BPS indicated in the Y-axis. Bars indicate the standard error of the mean. Values of the statistic significantly higher than zero (*P*-value < 0.05) are indicated with black color. In each plot, BSP are ordered according to decreasing values of the f3-statistics. All three samples originate from current-day Zambia. For WUD010, WUD012, WUD018 radio-carbon calibrated age is 1505–1668 CE, 1698-1950 CE and 1635-1950 CE, respectively, and genome coverage percentage is 0.270, 0.130, and 0.058, respectively (**Table.S 11**).

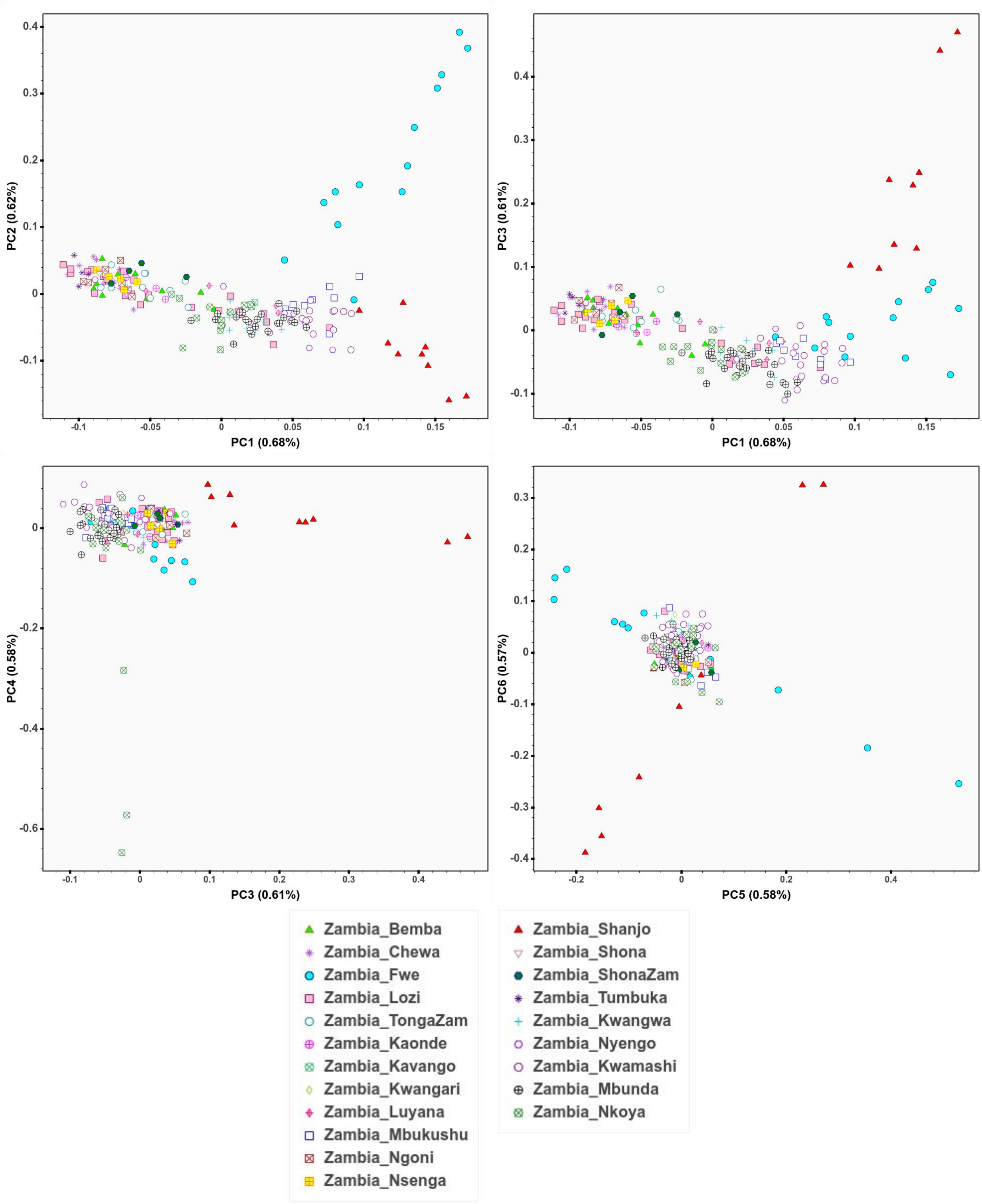


**Fig.S 12.11 | PCA of Only-Zambia database**

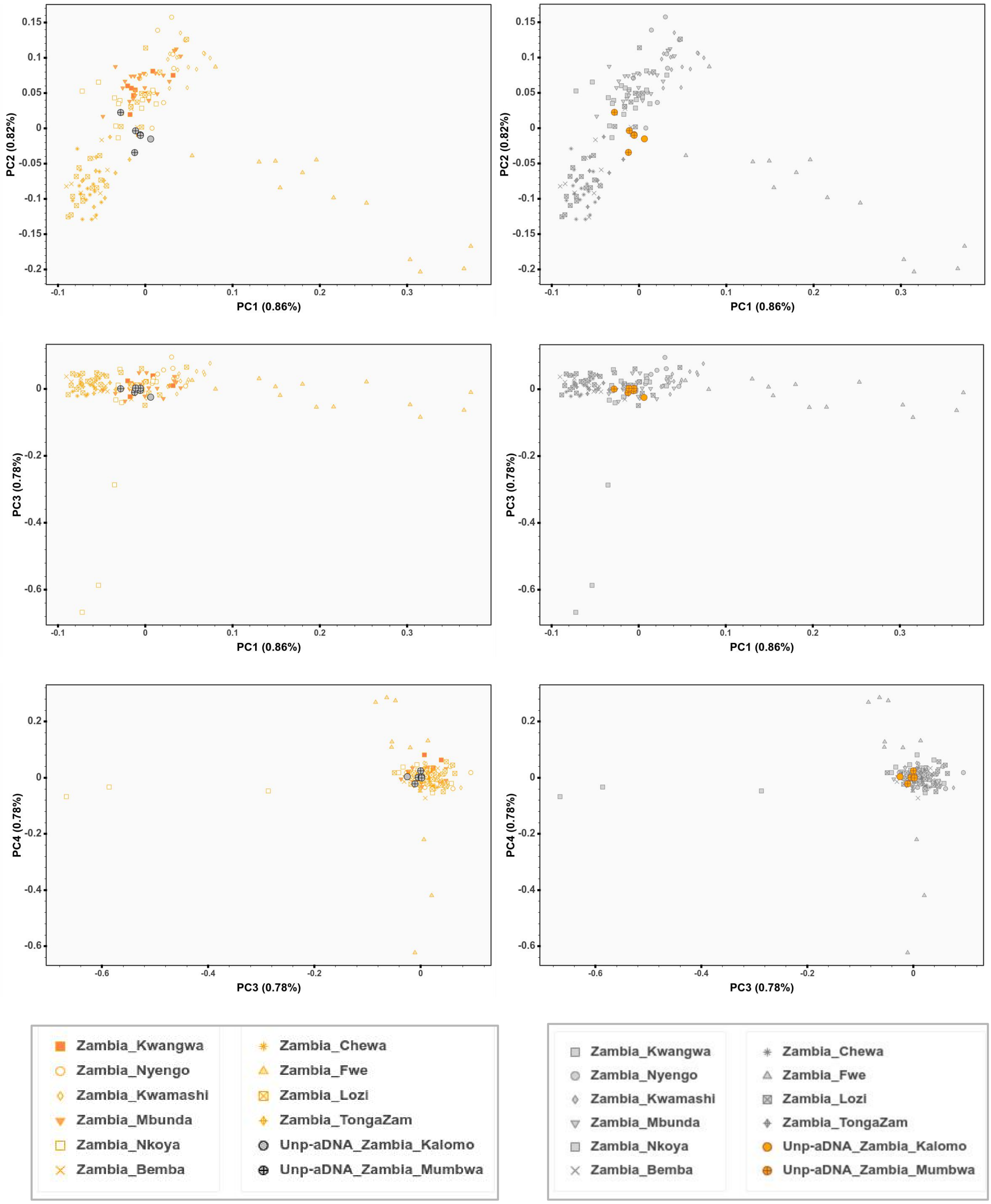


**Fig.S 12.12 | PCA of ancient samples and modern BSP from Zambia.**
Figure showing PCA plot of ancient DNA samples onto a background of BSP from Zambia. Figure showing in the left column modern BSP in orange and aDNA individuals in gray, and in the right column modern BSP in gray and aDNA individuals in orange.

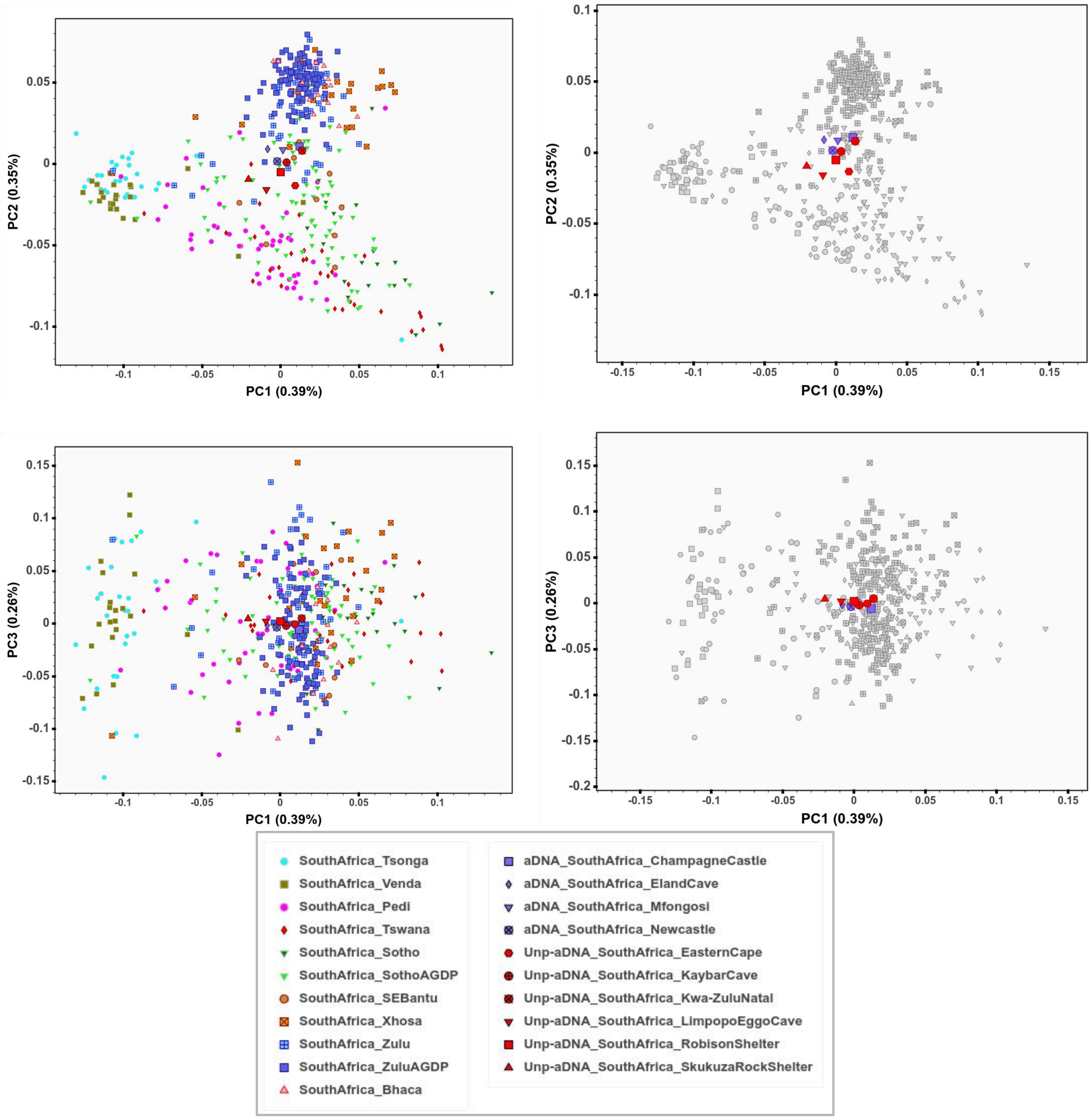


**Fig.S 12.13 | PCA of ancient samples and modern BSP from South Africa.**
Figure showing PCA plot of ancient DNA samples onto a background of BSP from South Africa. Figure showing in the left column modern BSP with colors and aDNA individuals in red and purple, and in the right column modern BSP in gray and aDNA individuals in red and purple.

**a**

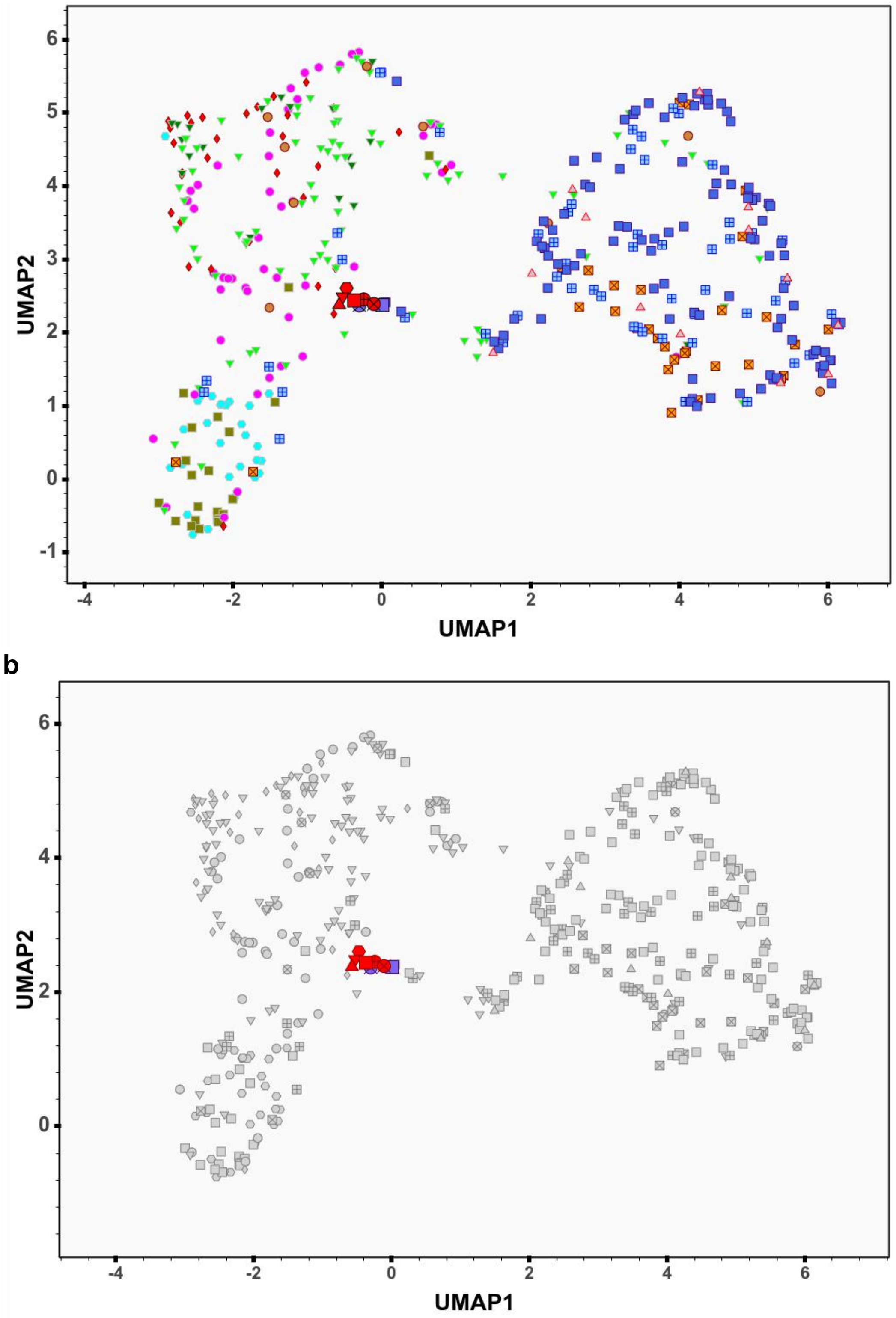


**Fig.S 12.14 | PCA-UMAP of ancient samples and modern BSP from South Africa.**
Figure showing PCA-UMAP plot of ancient DNA samples (red and purple markers) onto a background of BSP from South Africa (**a**) in different colors and (**b**) in gray.

#### 13- PCA results after masking datasets

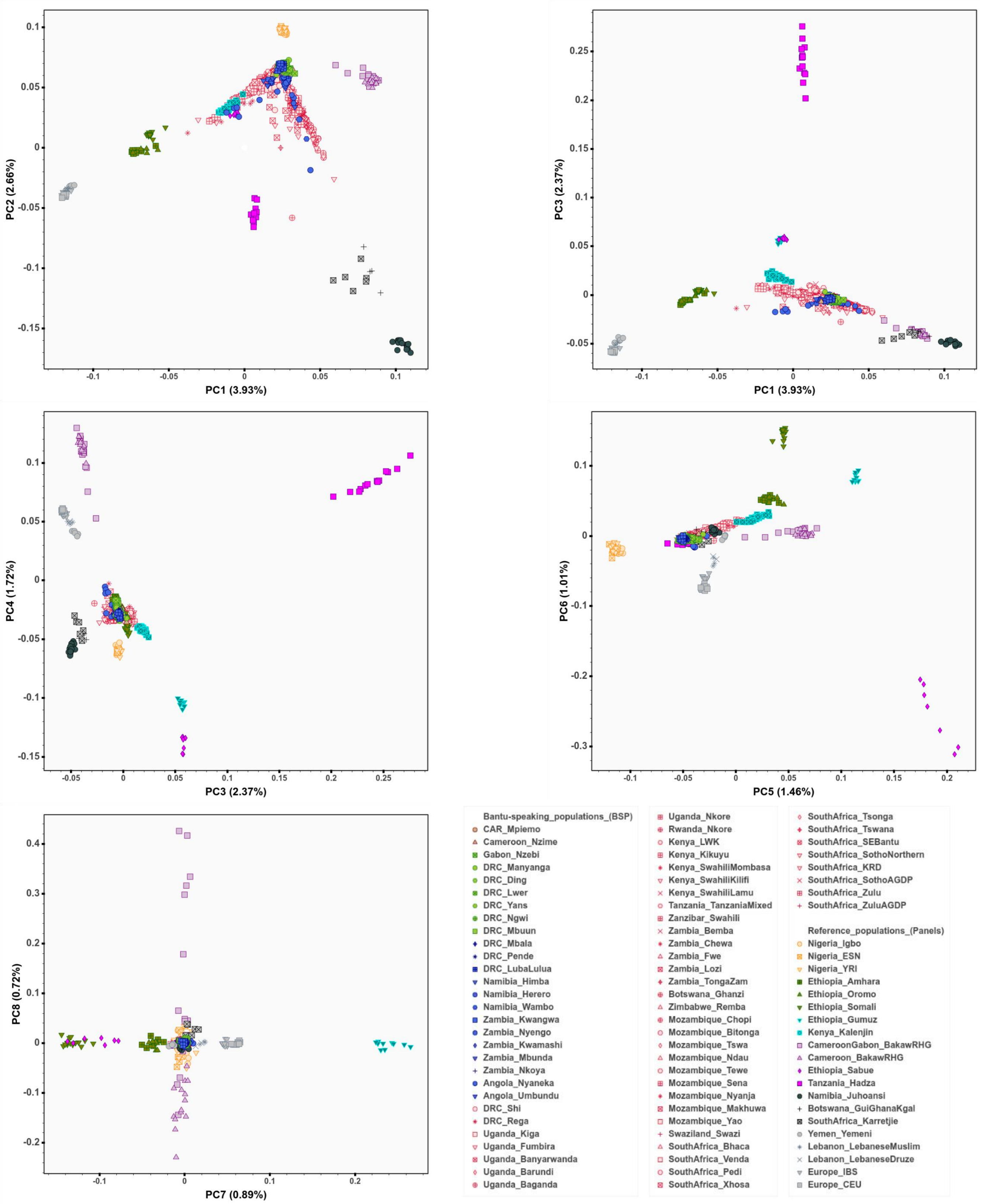


**Fig.S 13.1 | PCA of BSP and six selected reference panels.**
PCA plot of unmasked BSP and unmasked data of selected six reference panels.

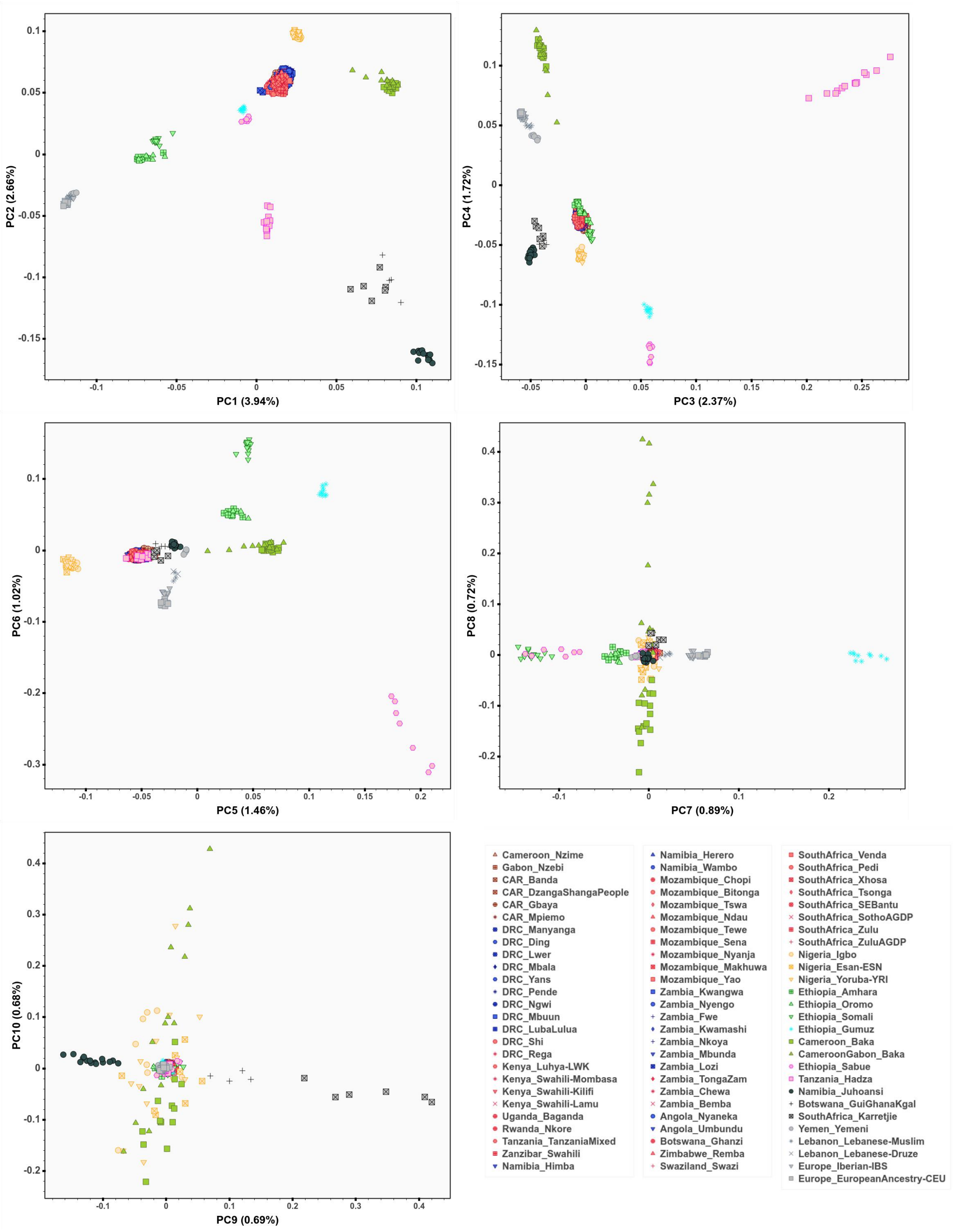


**Fig.S 13.2 | Ancestry-specific (AS-)PCA of BSP and six selected reference panels.**
AS-PCA plot of masked BSP dataset and unmasked data of selected six reference panels. We used masking phasing and imputation to analyze BSP with at least 70% WCA-related ancestry.

#### 14. List of Supplementary Tables included in the Excel file

**Table.S 1 |** All the samples that were genotyped for this study (1,740 individuals) in the Illumina H3Africa array.

**Table.S 2 |** Populations and groups that were included in each assembled dataset.

**Table.S 3 |** ADMIXTURE results at K4 on the basis of the AfricanNeo dataset. Table showing estimated averages for each K-group in each population. Each K-group was labelled with the population that has the highest values for that K-group.

**Table.S 4 |** ADMIXTURE results at K16 on the basis of the AfricanNeo dataset. Table showing estimated averages for each K-group in each population. Each K-group was labelled with the population that has the highest values for that K-group.

**Table.S 5 |** Ancient DNA samples from previous publications that were included in this study.

**Table.S 6 |** Averages (and standard deviation) for eleven parameters of runs of homozygosity (ROH) estimated in each population included in the AfricanNeo dataset.

**Table.S 7 |** Dates since the founder event (Tf, in generations before present), strength of the intensity of the bottleneck events (If in %) and normalized root mean squared deviation (NRMSD) between the empirical decay curve and the theoretical decay curve in each sub-Saharan African population that was included in the AfricanNeo dataset estimated using ASCEND. Populations with significant founder events were highlighted in bold.

**Table.S 8 |** Admixture dates (in generations) estimated using MOSAIC for selected BSP using a two-way admixture model.

**Table.S 9 |** Correlations between genetic, linguistic and geographical distances. Correlations (r-statistics and P-values) were estimated on the basis of the masked and imputed Only-Bantu dataset using partial Mantel test between matrices of genetic distances (pairwise FST matrix between BSP), linguistic distances (uncorrected linguistic distances from the multistate matrix) and geographical distances (geographical matrix). To compute Pearson's r-statistics, we used 100,000 permutations.

**Table.S 10 |** H3Africa populations used in the spatially explicit framework.

**Table.S 11 |** Statistics of ancient DNA samples presented in this study for the first time.

**Table.S 12 |** Linguistic data analyzed in the present study.
